# Single cell derived mRNA signals across human kidney tumors

**DOI:** 10.1101/2020.03.19.998815

**Authors:** Matthew D Young, Thomas J Mitchell, Lars Custers, Thanasis Margaritis, Francisco Morales, Kwasi Kwakwa, Eleonora Khabirova, Gerda Kildisiute, Thomas RW Oliver, Ronald R. de Krijger, Marry M. van den Heuvel-Eibrink, Federico Comitani, Alice Piapi, Eva Bugallo-Blanco, Christine Thevanesan, Christina Burke, Elena Prigmore, Kirsty Ambridge, Kenny Roberts, Felipe A Vieira Braga, Tim HH Coorens, Ignacio Del Valle, Anna Wilbrey-Clark, Lira Mamanova, Grant D Stewart, Vincent J Gnanapragasam, Dyanne Rampling, Neil Sebire, Nicholas Coleman, Liz Hook, Anne Warren, Muzlifah Haniffa, Marcel Kool, Stefan M Pfister, John C Achermann, Xiaoling He, Roger A Barker, Adam Shlien, Omer A Bayraktar, Sarah Teichmann, Frank C. Holstege, Kerstin B Meyer, Jarno Drost, Karin Straathof, Sam Behjati

**Affiliations:** Wellcome Sanger Institute, Hinxton CB10 1SA, UK; Cambridge University Hospitals NHS Foundation Trust, Cambridge, CB2 0QQ, UK; Department of Surgery, University of Cambridge, Cambridge, CB2 0QQ, UK; Princess Máxima Center for Pediatric Oncology, 3584 CS Utrecht, The Netherlands; Oncode institute, 3584 CS Utrecht, The Netherlands; Department of Pathology, University of Cambridge, Cambridge, CB2 1QP, UK; Program in Genetics and Genome Biology, The Hospital for Sick Children, Toronto, Ontario, Canada; Department of Laboratory Medicine and Pathobiology, University of Toronto, Toronto, Ontario, Canada; Department of Paediatric Laboratory Medicine, The Hospital for Sick Children, Toronto, Ontario, Canada; UCL Great Ormond Street Hospital Institute of Child Health, London WC1N 1EH, UK; Amsterdam UMC, University of Amsterdam, Amsterdam, The Netherlands; Great Ormond Street Hospital for Children NHS Foundation Trust, London WC1N 3JH, UK; Cambridge Urology Translational Research and Clinical Trials office, Cambridge Biomedical Campus Cambridge CB2 0QQ University of Cambridge; Department of Dermatology and NIHR Newcastle Biomedical Research Centre, Newcastle Hospitals NHS Foundation Trust, Newcastle upon Tyne, NE1 4LP, UK; Intitute of Cellular Medicine, Newcastle University, Newcastle upon Tyne, NE1 4HH, UK; Hopp Children’s Cancer Center Heidelberg (KiTZ), 69120 Heidelberg Germany; German Cancer Research Center (DKFZ) and German Cancer Consortium (DKTK), Division of Pediatric Neurooncology, 69120 Heidelberg, Germany; Heidelberg University Hospital, Department of Pediatric Hematology and Oncology, 69120 Heidelberg, Germany; MRC-WT Cambridge Stem Cell Institute, University of Cambridge, CB2 0QQ, Cambridge, UK; Department of Clinical Neuroscience, University of Cambridge, Cambridge, CB2 0QQ, UK; Department of Paediatrics, University of Cambridge, Cambridge, CB2 0QQ, UK

## Abstract

The cellular transcriptome may provide clues into the differentiation state and origin of human cancer, as tumor cells may retain patterns of gene expression similar to the cell they derive from. Here, we studied the differentiation state and cellular origin of human kidney tumors, by assessing mRNA signals in 1,300 childhood and adult renal tumors, spanning seven different tumor types. Using single cell mRNA reference maps of normal tissues generated by the *Human Cell Atlas* project, we measured the abundance of reference “cellular signals” in each tumor. Quantifying global differentiation states, we found that, irrespective of tumor type, childhood tumors exhibited fetal cellular signals, thus replacing the long-held presumption of “fetalness” with a precise, quantitative readout of immaturity. By contrast, in adult cancers our assessment refuted the suggestion of dedifferentiation towards a fetal state in the overwhelming majority of cases, with the exception of lethal variants of clear cell renal cell carcinoma. Examining the specific cellular phenotype of each tumor type revealed an intimate connection between the different mesenchymal populations of the developing kidney and childhood renal tumors, whereas adult tumors mostly represented specific mature tubular cell types. RNA signals of each tumor type were remarkably uniform and specific, indicating a possible therapeutic and diagnostic utility. We demonstrated this utility with a case study of a cryptic renal tumor. Whilst not classifiable by clinical pathological work-up, mRNA signals revealed the diagnosis. Our findings provide a cellular definition of human renal tumors through an approach that is broadly applicable to human cancer.

## Introduction

As cancer cells evolve from normal cells, they may retain patterns of messenger RNA (mRNA) characteristic of the cell of origin. In such cases, the cancer cell transcriptome may contain information that can identify the cancer cell of origin, its differentiation state, or trajectory towards a cancer cell. It is therefore conceivable that tumor transcriptomes can be used to identify the cells from which tumors arise and test fundamental hypotheses regarding tumor’s differentiation states, such as the “fetalness” of childhood tumors or the dedifferentiation of adult tumors towards a fetal state.

Single cell transcriptomics allows for a direct quantitative comparison to be made between single tumor and relevant normal cell transcriptomes. For example, single cell transcriptomes identified that a specific subtype of proximal tubular cells are the normal cell correlate of clear cell renal cell carcinoma (ccRCC) cells^1^. Such experiments can also reveal more precise information about normal cells within the tumor microenvironment. However, the high resource requirements of single cell transcriptomics preclude investigations of large patient cohorts, which are required to study rare subtypes, test the generalizability of such signals and determine associations with clinical parameters. An alternative approach is to identify the presence of single cell derived mRNA signals in bulk tumor transcriptomes, utilizing normal single cell transcriptomes as a reference. Smaller numbers of single cancer cell experiments can then be used to validate cellular signals identified.

Tumor bulk transcriptomes for most types of human cancer have been generated in the context of cancer genomics efforts of recent years, such as those conducted by the *International Cancer Genome Consortium* (ICGC) and *The Cancer Genome Atlas* (TCGA)^2,3^. Single cell reference data, generated by efforts collectively known as the *Human Cell Atlas*^4,5^, have begun to provide quantitative transcriptional definitions of the normal cells that constitute the developing and mature human kidneys^1,6–10^. By combining these bulk tumor transcriptome databases with single cell reference data, we may therefore be able to identify single cell signals in bulk transcriptomes across large cohorts of kidney tumors.

Here, we studied normal single cell mRNA signals in bulk kidney tumor transcriptomes (n=1,258; **Fig. 1A, Table S1**) and validated our findings using targeted single cell experiments (n=10, **Fig. 1A, Table S2**). There were three central aims of our analyses. Firstly, we tested the fundamental presumption that childhood renal tumors exhibit fetal cell signals whilst adult tumors dedifferentiate towards a fetal state. Next, we defined for each tumor type its normal cell correlate which may represent its cell of origin and provide diagnostic cues. Finally, we explored the tumor micro-environment across different tumor types.

**Figure 1 –.**
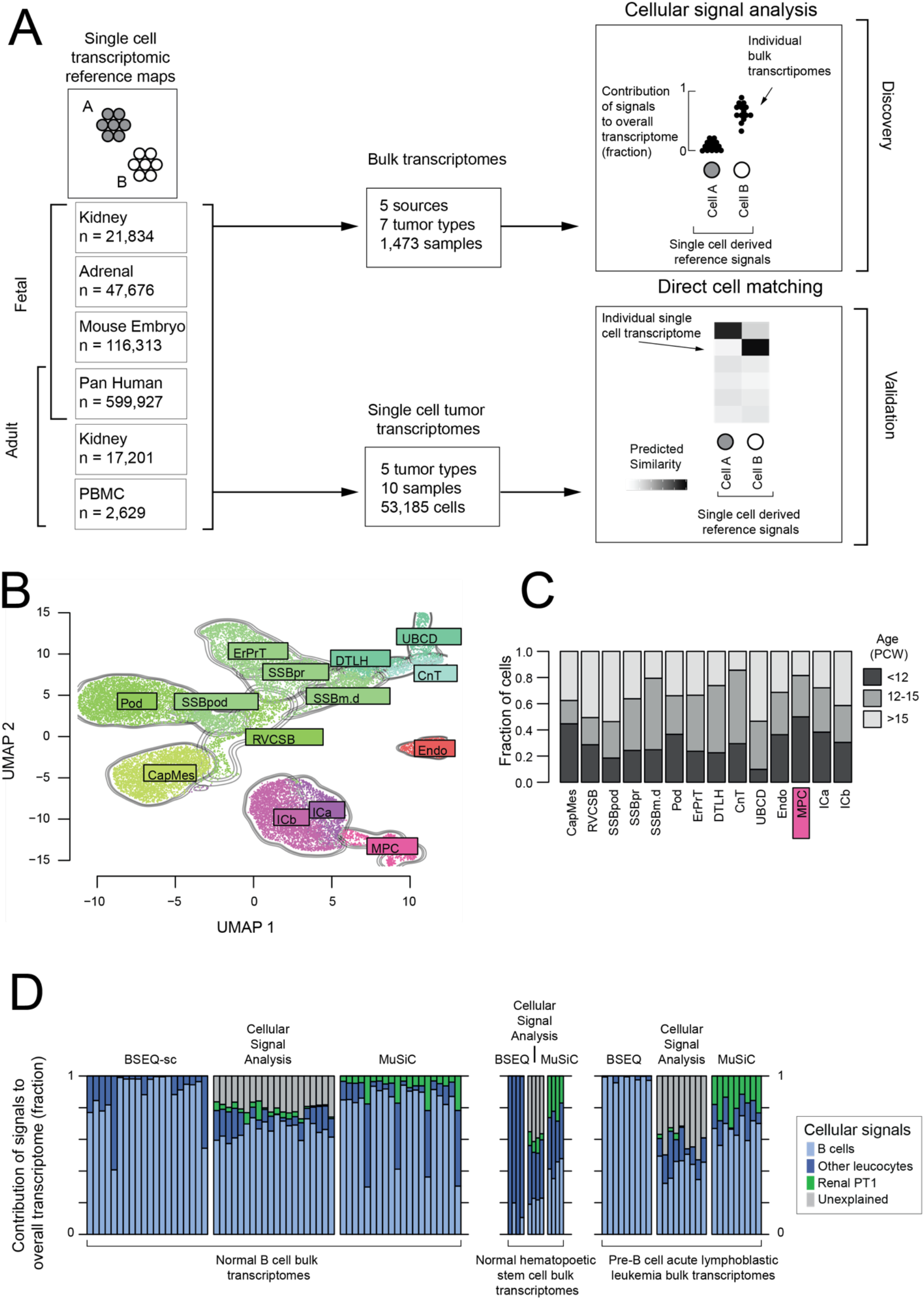
Methodology overview and validation. **A. Schematic of methodology:** Overview of the method used in this study. Single cell reference signals are defined from reference single cell atlases (left). These are then compared to bulk transcriptomes (top, **Table S1**) to discover the dominant normal signal contribution and validated using single cell transcriptomes (bottom, **Table S2**). Bulk transcriptomes are combined with reference transcriptomes to calculate the relative contribution of each reference signal in explaining the bulk transcriptome (see **Methods**). The strength of the signals are normalized to sum to 1, so each signal (corresponding to a population of single cells in a reference single cell RNA-seq dataset) is given a score between 0 and 1 for each bulk transcriptome, representing the relative strength of this signal in the bulk transcriptome in question (top right). For single cell transcriptomic validation, logistic regression is used to calculate a similarity score for each single cell transcriptome to each reference population (bottom right). **B. Combined fetal kidney reference map:** UMAP representation of fetal kidney reference map combining previously published and newly generated data. Contours and colors indicate the labelled cell type. Abbreviations: CapMes – Cap Mesenchyme, RVCSB – Renal vesicle and comma-shaped body, SSBpod – S-shaped body podocyte, SSBpr – S-shaped body proximal tubules, SSBm.d – S-shaped body medial and distal, Pod – Podocytes, ErPrT – Early proximal tubules, DTLH – Distal tubule and loop of Henle, UBCD – Ureteric Bud and collecting duct, CnT – Connecting tubules, Endo – Endothelium, ICa – Interstitial cells a (smooth muscle), ICb – Interstitial cells b (stromal), MPC – Mesenchymal progenitor cells. **C. Age distribution of fetal kidney populations:** Bar heights indicate the fraction of cells from fetal kidney populations in **B** derived from different aged fetuses as indicated by the color scale. Age is given in post conception weeks. **D. Method comparison on flow sorted immune-cells and ALL:** Comparison of two widely used “bulk deconvolution” methods (BSEQ-sc and MuSiC) to cellular signal analysis. Each bar represents a bulk transcriptome from a population of flow sorted adult B-cells (left), hematopoietic progenitors (middle), and pre-B acute lymphoblastic leukemia (right). The signal contribution is calculated for each sample using a reference signal set consisting of immune cells from adult peripheral blood mononuclear cells and proximal tubular cells from the mature (pediatric and adult) kidney (PT1 cells, included as a negative control). The relative contribution of each of these reference signals (plus an “unexplained signal” component where appropriate) is shown by the size and color of the stacked bars for each sample, as indicated by the legend on the far right.

## Results

### An integrated single cell reference map of the kidney

The nephron is the functional unit of the kidney and together with its associated vasculature and support cells make up the majority of kidney cells. The nephron is derived from the mesoderm and forms from a combination of mesenchymal cell populations that mature into the epithelial cells of the nephron via mesenchymal to epithelial transition (MET)^11^. To precisely define these mesenchymal populations and the populations they mature into, we created a refined fetal kidney reference map combining previously generated ^1,8^ and newly generated human fetal kidney single cell data (**Fig. 1B, S1**).

This reference revealed 4 key mesenchymal populations: mesenchymal progenitor cells (MPCs), cap mesenchyme (CM), and two populations of specialized interstitial cells: smooth muscle-like cells (ICa), and cortical stromal cells (ICb) (**Fig. 1B, S1A-B**)^8,11^. The cap mesenchyme condenses on the ureteric bud and forms the tubular structures of the nephron via mesenchymal to epithelial transition. The mesenchymal cells which do not condense into cap mesenchyme and remain in the interstitial space form interstitial support cells for the nephron, such as mesangial cells. The final mesenchymal population, which we termed mesenchymal progenitor cells, was not present in sufficient numbers to be reported in earlier single cell transcriptomic studies of the developing kidney. These MPCs are enriched for early time points (**Fig. 1C**), strongly resemble mesenchymal cells in the fetal adrenal (**Fig. S1D**)^12^, and both populations resemble primitive mesodermal populations in the post gastrulation mouse embryo (**Fig. S1D**)^13^. Developmentally, both the adrenal cortex and the kidney are derived from the same mesodermal lineage.

We combined this refined map of the developing kidney with previously generated maps of the mature kidney^1^, the developing adrenal gland^12^, and the post-gastrulation mouse^13^ (**Fig. 1A**). Together these provide a complete single cell reference map of the kidney across developmental time.

### Quantification of reference cellular mRNA signals in bulk transcriptomes

Our single cell reference map of the kidney provides a cellular mRNA signal of each population of cells. To measure the abundance of these reference cellular signals in bulk tumor transcriptomes, we devised a method that fits raw bulk mRNA counts for the entire transcriptome – not just marker genes – to a weighted linear combination of transcriptomic signals derived from reference single cell data.

A number of bulk deconvolution tools exist that aim to identify the cellular composition of bulk tissues using a single cell reference^14–16^. However, the aim of our analysis was not to identify and quantify the number of cells present in the microenvironment, but to identify the major cellular signals (or transcriptional programs) used by tumor cells. As such, we do not expect any of our single cell reference populations to exactly match the tumor cells’ transcriptome. We therefore designed our method to identify the major transcriptional signals (defined using single cell data) present in bulk transcriptomic data. We term this approach “cellular signal analysis” to differentiate it from “deconvolution analysis”, the inference of cellular composition of bulk transcriptomes.

We applied cellular signal analysis to 766 ccRCC transcriptomes from The Cancer Genome Atlas^17^ to assess whether the known cellular identity of these cancer cells could be identified. Our method correctly identified the signal of a specific proximal tubular cell population as the predominant cell signal in ccRCC bulk cancer transcriptomes (**Fig. S2**). We next applied cellular signal analysis and published deconvolution methods, MuSiC^15^ and BSeq-SC^14^ to bulk mRNA of purified normal human B-cells, human pre-B cell leukemia, and hematopoietic stem cells (**Fig. 1D**)^18,19^. For this comparison we used a reference that combined single cell transcriptomes from peripheral blood cells with a negative control population; proximal tubular kidney cells. BSeq-SC was unable to differentiate between normal mature B cells and cancerous pre-B cell leukemia cells, giving both a mature B cell label. MuSiC found an implausible renal tubular signal in both hematopoietic stem cells and leukemia.

Cellular signal analysis instead explained most of the difference between the tumor and reference transcriptomes through an “unexplained signal”. Mathematically, this “unexplained signal” represents an intercept term, included to limit the assignment of spurious signals when a bulk transcriptome differs from all signals in the reference (see **Methods**). Taken together, these comparisons demonstrate the need for a bespoke analysis tool to perform cellular signal analysis and that the unexplained signal metric of our method uniquely highlights when reference data is inappropriate.

### Childhood tumors, but not adult tumors, exhibit a fetal transcriptome

For each tumor, we determined whether it exhibited a fetal or mature (i.e. post-natal) transcriptome, to guide the choice of reference in subsequent analyses. This analysis also enabled us to test two fundamental hypotheses about the differentiation state of tumors – that childhood tumors represent fetal cell types and that adult cancers, especially epithelial malignancies, dedifferentiate towards a fetal state.

We calculated the immaturity by fitting each bulk transcriptome to a combined reference set composed of cellular signals from both mature and fetal kidney reference populations. The immaturity score was the fractional contribution of the developmental signals to the bulk transcriptome. Using this approach, we established a reference range of mature normal kidneys (**Fig. 2A**). We demonstrated the validity of this range by scoring fetal kidney transcriptomes which lay significantly outside the mature range (p=0.015, Wilcoxon rank sum test).

**Figure 2 –.**
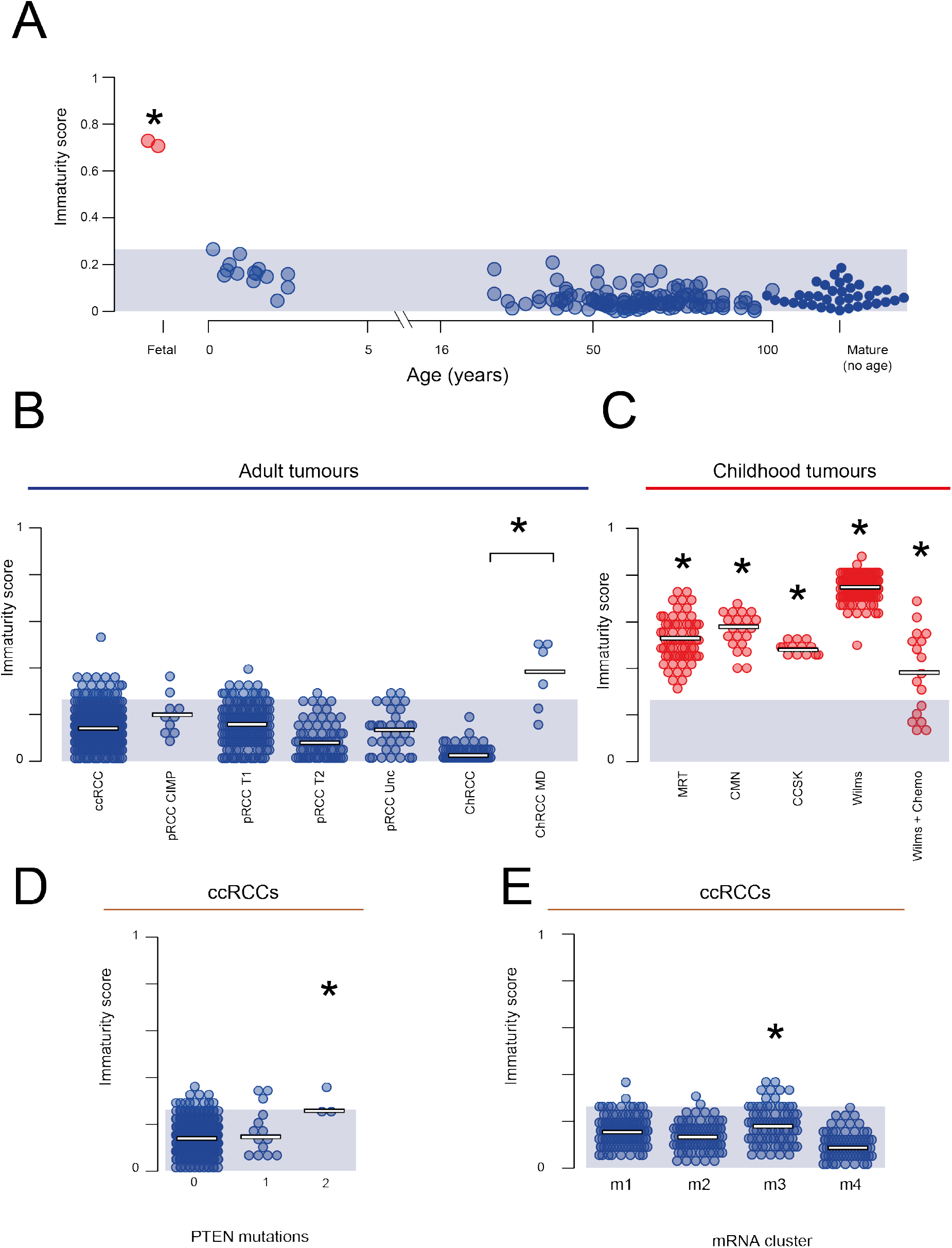
Immaturity score. **A. Immaturity score for normal renal bulk transcriptomes:** For 201 bulk transcriptomes derived from normal kidney biopsies across the human life span, an immaturity score was calculated. This score was calculated by fitting each bulk transcriptome using a combined reference set consisting of signals from all cells in the mature and fetal kidney. The immaturity score is then the total normalized signal contribution from fetal kidney in each bulk transcriptome (shown on the y-axis). Samples are shown by age when known (x-axis), with samples for which no age information was available shown on the right. The shaded blue area indicates the range of maturity scores across all normal tissue post-natal transcriptomes. Bulk fetal transcriptomes are shown on the left in red, with a star indicating that their maturity score is significantly higher than that of the normal samples (p=0.015, Wilcoxon rank sum test). **B. Immaturity score for adult renal tumor transcriptomes:** The immaturity score is calculated as in **A.** for 853 adult renal tumors. The normal immaturity score range is shown by the blue shaded region. The metabolically divergent subtype of Chromophobe renal cell carcinomas are shown separately from other chromophobe RCCs. These metabolically divergent tumors have a significantly different maturity score (p<10^-4^, Wilcoxon rank sum test). **C. Immaturity score for childhood renal tumor transcriptomes**: The immaturity score is calculated as in **A.** for 287 childhood renal tumors. The normal immaturity score range is shown by the blue shaded region. Each type of childhood tumor had a significantly different maturity score than post-natal normal tissue kidneys (p<10^-4^, Wilcoxon rank sum test). **D. Immaturity score for ccRCCs by PTEN mutation status:** The immaturity score for clear cell renal cell carcinomas as calculated in **A**, broken down by PTEN mutation status with 0 indicating wild type, 1 mono-allelic loss and 2 bi-allelic loss. The star indicates that bi-allelic loss is a significant predictor of higher immaturity score (p<0.01). **E. Immaturity score for ccRCCs by transcriptional group:** The immaturity score for clear cell renal cell carcinomas as calculated in **A**, broken down by the transcriptomic subgroups identified in^20^. The star indicates that samples in m3 have a significantly lower immaturity score (p<0.01).

We next calculated the same maturity score for individual tumors, which showed a clear signal of “fetalness” across all types of childhood kidney tumors (**Fig. 2B-C**). Although all childhood kidney tumors had a significant enrichment for developmental cellular signals, pretreated Wilms tumor had a significantly lower score than other childhood kidney tumors, including untreated Wilms. The comparison between treated and untreated Wilms suggests that chemotherapy reduces the developmental signal in Wilms tumor, a notion we explore in detail in a later section.

A significant developmental signal was absent from almost all adult tumors (**Fig. 2B**). This suggests that global “dedifferentiation” to a developmental state does not occur in adult kidney tumors. One obvious exception to the ubiquitous lack of a strong developmental signal in adult tumors (p<10^-4^, Wilcoxon rank sum test) was a cohort of lethal chromophobe RCC, classified previously as metabolically divergent due to their comparatively low expression of genes associated with the Krebs cycle, electron transport chain, and the AMPK pathway^17^.

Motivated by this observation, we tested whether other clinical markers such as somatic genotype, morphology or molecularly defined subgroup were predictive of immaturity score. We found that clear cell renal cell carcinomas with two independent somatic mutations in *PTEN* had a significantly higher immaturity score (**Fig. 2D;** t-test, FDR<0.01). As with lethal chromophobe tumors, *PTEN* mutated ccRCCs conferred a far worse prognosis, with all samples belonging to the TCGA defined m3/CCB subgroup with the worst prognostic outcome of all groups^20^. Investigating further, we found an association between immaturity score and the m3 transcriptional subgroup (**Fig. 2E**; t-test, FDR<0.01). No other clinical covariate had a statistically significant association with immaturity score at a 1% significance level (**Table S3-4**).

### Congenital mesoblastic nephroma resembles mesenchymal progenitor cells

Congenital mesoblastic nephroma (CMN) is a renal tumor of infants that has low metastatic potential. There are two morphological subtypes of CMN, classical and cellular variants^21^. Cell signal analysis in CMN bulk transcriptomes (n=18) revealed a uniform signal of mesenchymal progenitor cells across tumors (**Fig. 3A, S3**), irrespective of morphological subtype. Of note, these mesenchymal progenitor cells were characterized by expression of *NTRK3* and *EGFR* genes (**Fig. 3B**), the principal oncogenes that drive CMN through activating structural variants^22^. To verify that this signal was not a generic consequence of fibroblast like cells, we repeated the analysis of bulk CMN transcriptomes using a developmental reference combined with mature fibroblasts. This comparison revealed the same match to mesenchymal progenitor cells, with a low contribution from mature fibroblasts (**Fig. 3C**).

**Figure 3 –.**
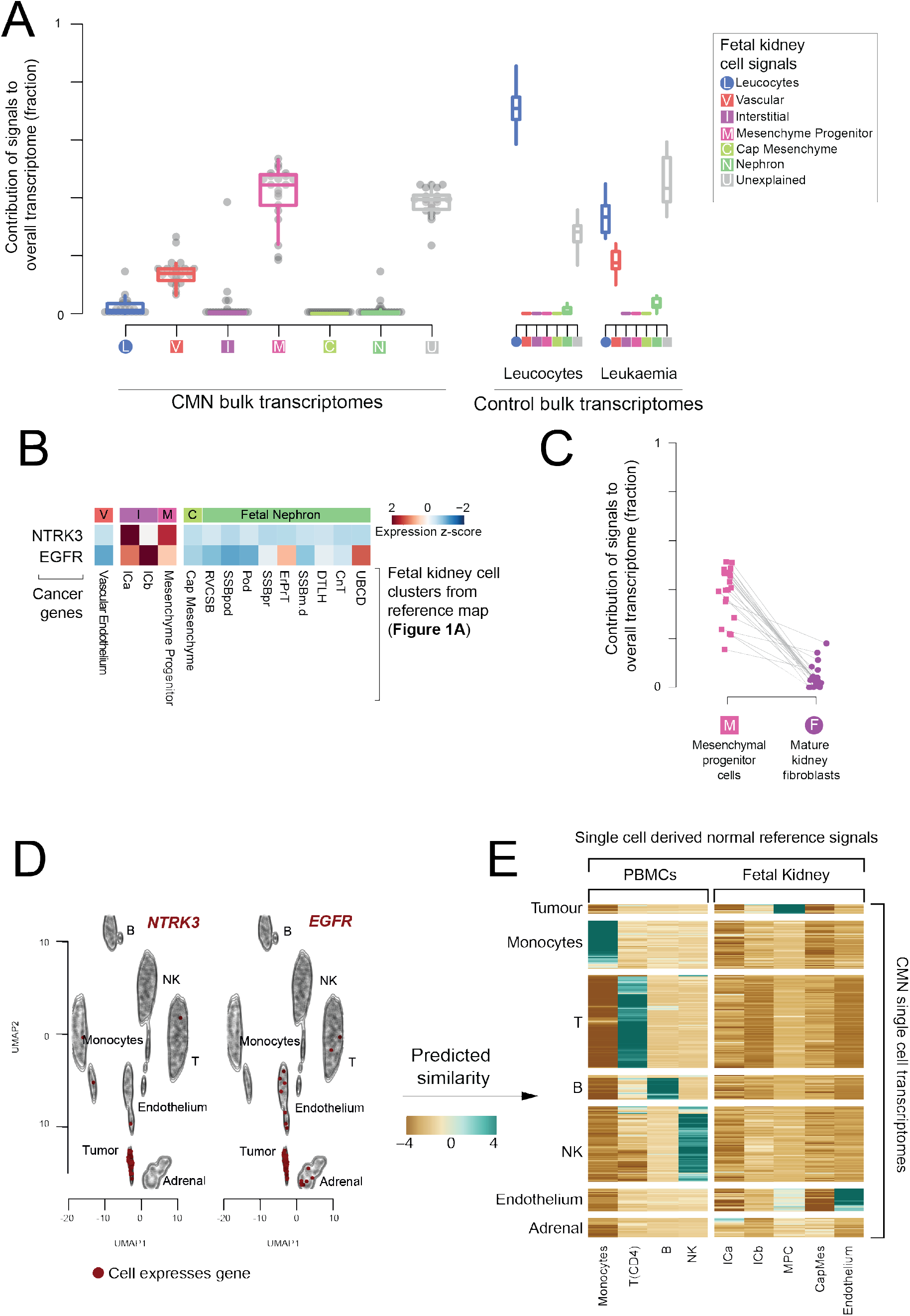
Congenital mesoblastic nephromas. **A. Composition of bulk CMNs:** The relative contribution of single cell derived signals from fetal kidney in explaining the bulk transcriptomes of 18 congenital mesoblastic nephromas (CMNs) along with control leucocyte and ALL populations. The relative contribution of each signal to each bulk RNA-seq sample is shown by the y-axis. Each signal/sample combination is represented by a single point and the distribution of relative signal contributions to the bulk transcriptomes are summarized with boxplots. Each signal type is labelled with an abbreviation and colored as shown by the legend on the right. Signals are marked with a square for fetal kidney and a circle for mature. CMN samples are shown in the block on the left, while the two groups of control samples are shown on the right. The samples marked “Leukocytes” are bulk transcriptomes from flow sorted leukocytes, while the samples marked “Leukemia” represent B-precursor acute lymphoblastic leukemia. **B. Expression of CMN cancer genes in fetal kidney:** Expression of CMN driver genes (rows) in reference fetal kidney single cell RNA-seq populations (columns). The data has been scaled to have mean 0 and a standard deviation of 1 in each row (i.e., z-transformed). **C. Comparing mesenchymal progenitor cell signals to mature fibroblasts:** All 18 CMN bulk transcriptomes were analyzed using a reference signal set including both fetal kidney cells and the fibroblasts from mature kidney. This figure shows the comparison of their inferred contribution to each transcriptome for each sample (y-axis), with lines joining points representing the same sample. **D. Expression of CMN marker genes:** tSNE map of single cell transcriptomes of 4,416 cells derived from a CMN biopsy. Cells belonging to the same cluster are indicated by contours and are labelled by the cell type they represent. Cells positive for *NTRK3* (left) and *EGFR* (right) are colored red. Abbreviations: B = B cell; T = T cell; DC = dendritic cell; NK = NK cell; NKT = NKT cell. **E. Comparison of single cell CMN to fetal kidney:** Comparison of clusters of cells for which single cell transcriptomes were obtained from the CMN biopsy (rows) with fetal kidney and leucocyte reference populations (columns). For each CMN cluster/reference population pair a log-similarity score was calculated using logistic regression (see Methods). Positive log-similarity scores represent a high probability of similarity between the reference and test cluster.

To validate this mesenchymal stem cell signal in CMN, we subjected cells dissociated from a fresh CMN tumor specimen, to single cell mRNA sequencing using the Chromium 10x platform. We annotated single cells based on literature derived marker genes (**Fig. 3D, S4**) and compared to single cell clusters of normal fetal kidneys using previously developed quantitative approaches^1^. This comparison revealed that CMN tumor cells matched the same mesenchymal progenitor cell population, validating the cell signal seen in bulk tumor tissue (**Fig. 3E**).

### Wilms tumour, clear cell sarcoma of the kidney and the effect of treatment

Wilms tumor is the most common childhood kidney cancer and is thought to arise from aberrant cells of the developing nephron. Clear cell sarcoma of the kidney (CCSK) is a rare, at times aggressive childhood renal cancer that is treated as a high risk Wilms tumor in clinical practice^23^. We assessed the cellular signals in bulk transcriptomes from treatment-naïve CCSK, high risk treatment-naive Wilms, and intermediate risk Wilms post chemotherapy. Cellular signal analysis revealed a largely uniform early nephron signal (cap mesenchyme, comma shaped body, S-shaped body) in the treatment-naïve Wilms cohort (**Fig. 4A, S5**). By comparison, the post-treatment cohort had a much reduced contribution from the early nephron, instead of containing a mixture of tubular, early nephron and mesenchymal signals with a relatively high unexplained signal fraction (**Fig. 4A, S5**). Previous work utilizing single cell data from post-chemotherapy Wilms tumors identified the same lack of cap mesenchyme signal identified by our analysis of bulk transcriptomes^1^. The CCSK transcriptomes showed a mixture of mesenchymal and early nephron signals, with an extremely high unexplained signal fraction (**Fig. 4A, S5**).

**Figure 4 –.**
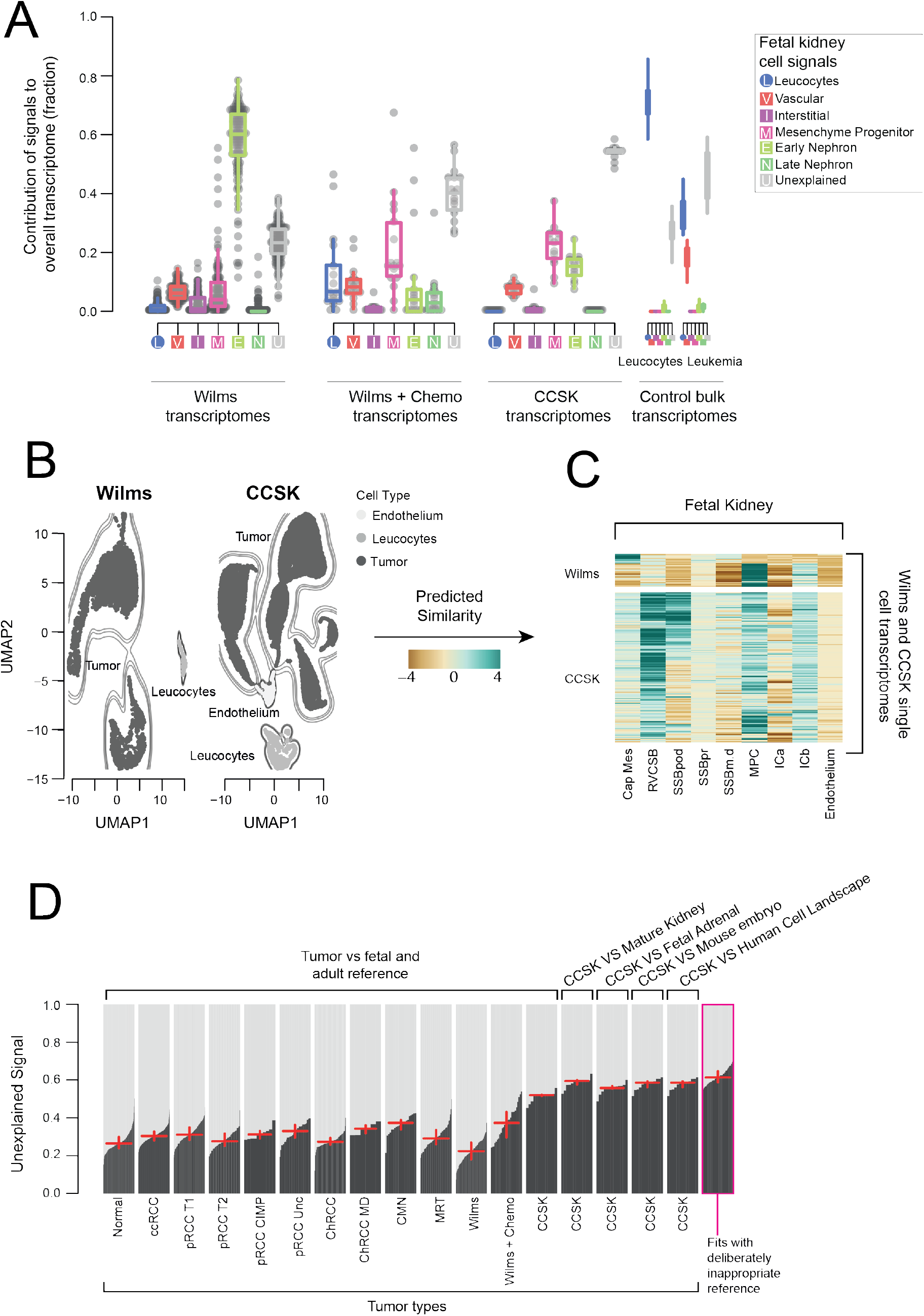
Wilms tumor and clear cell sarcoma of the kidney. **A. Bulk Wilms tumor and CCSK compared to fetal kidney:** The relative contribution of single cell derived signals from the fetal kidney in explaining the bulk transcriptomes of 137 nephroblastomas (Wilms tumors) and 13 CCSKs along with control leucocyte and ALL populations. The signal intensity assigned to each sample is shown by the y-axis, normalized so that the sum of all signal contributions is 1 for each sample. The relative contribution of each signal to each bulk RNA-seq sample is shown by the y-axis. Each signal/sample combination is represented by a single point and the distribution of relative signal contributions to the bulk transcriptomes are summarized with boxplots. Each signal type is labelled with an abbreviation and colored as shown by the legend on the right. Signals are marked with a square for fetal kidney and circle for mature. Wilms tumor samples are shown in the block on the left, while the two groups of control samples are shown on the right. The samples marked “Leucocytes” are bulk transcriptomes from flow sorted leukocytes, while the samples marked “Leukemia” represent B-precursor acute lymphoblastic leukemia. **B. UMAP of CCSK and Wilms single cell transcriptomes:** Each point represents a single transcriptome from 1 Wilms tumor (left) or 3 CCSK (right) single cell transcriptome samples. Shading and contours indicate the cell type, as also indicated by labels. **C. Comparison of CCSK and Wilms transcriptomes to reference signals:** The similarity of single cell transcriptomes of tumor populations in **B** compared to fetal kidney reference signals indicated on the x-axis. Each row represents a single transcriptome and the color indicates the logit similarity. **D. Comparison of unexplained signal contribution to CCSKs and other tumor types:** Comparison of the fraction of the bulk transcriptomes attributed to the “unexplained signal” in clear cell sarcoma of the kidney samples (CCSK) and other groups of samples. For each group of samples, the unexplained signal is calculated for each sample individually, using the reference set of signals given at the top of the plot (e.g., fetal kidney). The unexplained signal fractions are then shown by black bars, sorted in increasing order, with the red horizontal line showing the median value and the vertical line the range between the 25^th^ and 75^th^ percentiles. CCSK samples were fitted using 5 different reference sets (fetal and mature kidney, mature kidney only, fetal adrenal, mouse embryo, and the pan-tissue human cell landscape). The final group on the right, represents samples fitted using inappropriate references. This population serves as a calibration of the expected level of unexplained signal when the bulk transcriptome is not explained by any of the provided reference signals.

To validate the cap mesenchyme signal in treatment-naïve Wilms, we generated single cell mRNA transcriptomes from one fresh sample. Annotation of this data revealed two proliferating populations (**Fig. 4B, S6**). Comparison to fetal kidney showed that one of these populations strongly matched the cap mesenchyme, validating its presence in treatment-naïve Wilms tumor (**Fig. 4C**). The second population exhibited a strong match to mesenchymal progenitor cells (**Fig. 4C**).

To further investigate the origins of CCSK we generated single nuclear transcriptomes from 2 archival samples and single cell transcriptomes from one fresh sample (**Fig. 4B, S7**). In contrast to Wilms tumor, all CCSK tumor cells matched multiple mesenchymal and early nephron populations (**Fig. 4C**). Although the matching populations were consistent with the results of cell signal analysis on bulk CCSK transcriptomes (**Fig. 4A**), the match to multiple reference populations at the single cell level suggests that CCSK transcriptomes represent a transcriptional state that is intermediate between multiple mesenchymal populations in the developing kidney. To test the possibility that the true normal cell correlate for CCSKs was not in the fetal kidney, we next matched CCSK bulk transcriptomes against mature kidney, fetal adrenal, developing mouse, and the pan-tissue human cell landscape^24^. In each of these comparisons, the unexplained signal explained at least 50% of the CCSK bulk transcriptomes, a much higher fraction than any other tumor type (**Fig. 4D**). This unexplained signal fraction was comparable to the level obtained from a deliberately inappropriate comparison of flow sorted B cell bulk transcriptomes compared to the non-immune developing kidney (**Fig. 4D**). In aggregate, these data suggest that CCSKs represent transcriptionally grossly distorted renal mesenchymal cells.

### Malignant rhabdoid tumors exhibit signals of neural crest and early mesenchyme

Malignant rhabdoid tumor (MRT) is an aggressive, often fatal childhood cancer, that typically affects the kidney but may also occur at other sites. It is considered to be the extracranial counterpart of the CNS tumor, atypical teratoid/rhabdoid tumor (AT/RT). The principal, usually sole, driver event in MRT and AT/RT is biallelic inactivation of *SMARCB1*. In previous analyses of microRNA profiles, MRTs co-clustered with a range of tissues: neural crest derived tumors, cerebellum, and synovial sarcoma^25^.

Assessing fetal renal single cell signals in 65 MRT bulk transcriptomes yielded a mesenchymal progenitor cell signal (**Fig. 5A, S8**). However, the nephron and unexplained signal fractions were also high, indicating that tumor cells only moderately resemble this reference population. To investigate further, we studied MRT single cell transcriptomes, derived from an MRT expanded by a primary organoid culture (see **Methods**), from nuclear mRNA sequencing, and from fresh tissue MRT cells (**Fig. 5B, S9**). Comparison to our fetal kidney reference revealed that MRT cell transcriptomes did not show any consistent match (**Fig. 5C**). This may indicate that the mesenchymal progenitor cell signal obtained in bulk represents a signal of the broad embryological lineage of the tumor, rather than a cell type.

**Figure 5 –.**
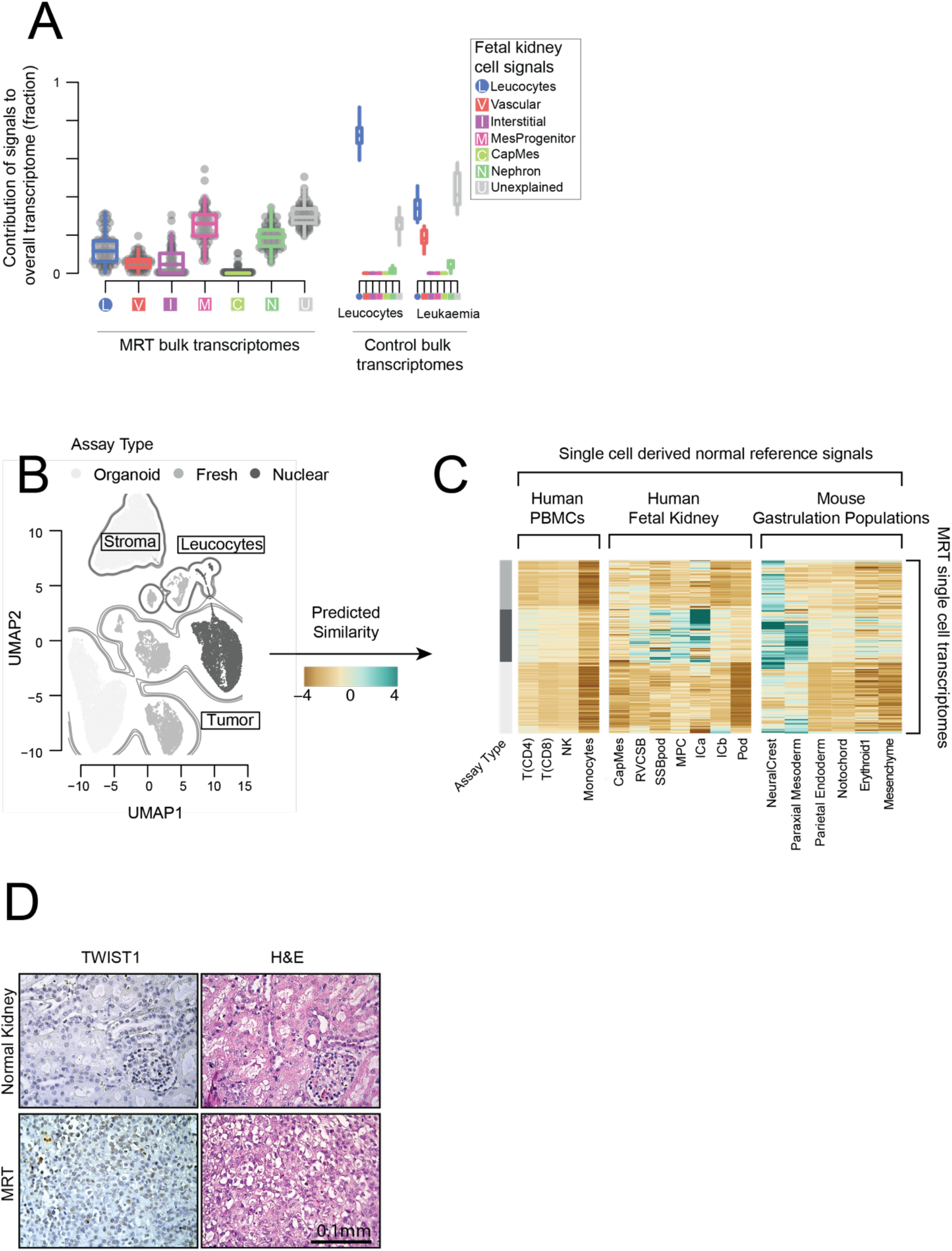
Malignant rhabdoid tumors. **A. Bulk MRTs compared to fetal kidney:** The relative contribution of single cell derived signals from fetal kidney in explaining the bulk transcriptomes of 65 malignant rhabdoid tumors (MRTs) along with control leukocyte and ALL populations. The relative contribution of each signal to each bulk RNA-seq sample is shown by the y-axis. Each signal/sample combination is represented by a single point and the distribution of relative signal contributions to the bulk transcriptomes are summarized with boxplots. Each signal type is labelled with an abbreviation and colored as shown by the legend on the right. Signals are marked with a square for fetal kidney and circle for mature. MRT samples are shown in the block on the left, while the two groups of control samples are shown on the right. The samples marked “Leucocytes” are bulk transcriptomes from flow sorted leucocytes, while the samples marked “Leukemia” represent B-precursor acute lymphoblastic leukemia. **B. UMAP of single cell MRT transcriptomes:** Each dot represents a single transcriptome from either tumor/tubular derived organoid cells (white), fresh tissue MRTs cells (grey) or archival MRT nuclei (black). Contours indicate tumor cells, stroma, and leucocytes as labelled. **C. Log similarity of single cell MRT cells to fetal kidney and developing mouse:** Comparison of the transcriptomes in **B** to cellular signals defined from single cell reference transcriptomes. The reference population is indicated on the x-axis and the grey bar on the left indicates the technology each cell was derived from. Each row corresponds to a single transcriptome from **B**. The color scheme encodes the logit similarity score for each cell against each reference population (see **Methods**). **D. Immunohistochemistry of *TWIST1* in MRT and normal kidney:** Staining of a region of normal kidney and MRT tissue for *TWIST1*. The MRT image shows a part of the tissue selected for being *TWIST1* positive, there were large sections of tumor tissue that were also *TWIST1* negative. All normal kidney tissue was *TWIST1* negative.

We therefore compared MRT cells against published reference cell populations of gastrulation embryos generated from mice^13^, a developmental stage that is not accessible to study in humans. Although there were differences between and within samples, all produced a match to neural crest and/or early mesodermal/mesenchymal populations (**Fig. 5C**). To validate this early mesodermal signal, we performed immunohistochemistry for the presence of a protein specific to paraxial mesoderm, TWIST1. Consistent with its expression in a subset of cells by single cell mRNA sequencing, occasional MRT cells exhibited *TWIST1* staining, whilst no protein was detected in normal kidney (**Fig. 5D, S10**). Overall our data show that MRTs do not exclusively exhibit mRNA signals of either neural crest or mesenchyme cells. Instead, our findings point at a hybrid state of MRTs, representing mRNA features of both, neural crest and mesenchyme, suggesting that MRTs may come from early mesoderm or form along the differentiation trajectory of neural crest to mesenchyme.

### Adult tumors represent specific tubular cells

As discussed above, our analyses confirmed a previous finding that the predominant single cell signal in the most common types of adult renal cancer, clear cell RCC (ccRCC) and papillary RCC (pRCC), was derived from a specific subtype of proximal tubular cells, termed PT1 cell (**Fig. S2**)^1^. In addition, cell signal analysis also revealed some properties of the tumor microenvironment. We found a prominent vascular endothelial signal in ccRCCs (**Fig. 6A, S11**), but not in pRCCs. The downstream effects in RCC of inactivation of the von Hippel-Lindau gene and upregulation of vascular endothelial growth factors are well documented^26^. The prominent difference in the endothelial signal provides a read-out of this pathway, further explaining why anti-angiogenic treatments appear to be more effective in ccRCCs than in pRCCs^27^.

**Figure 6 –.**
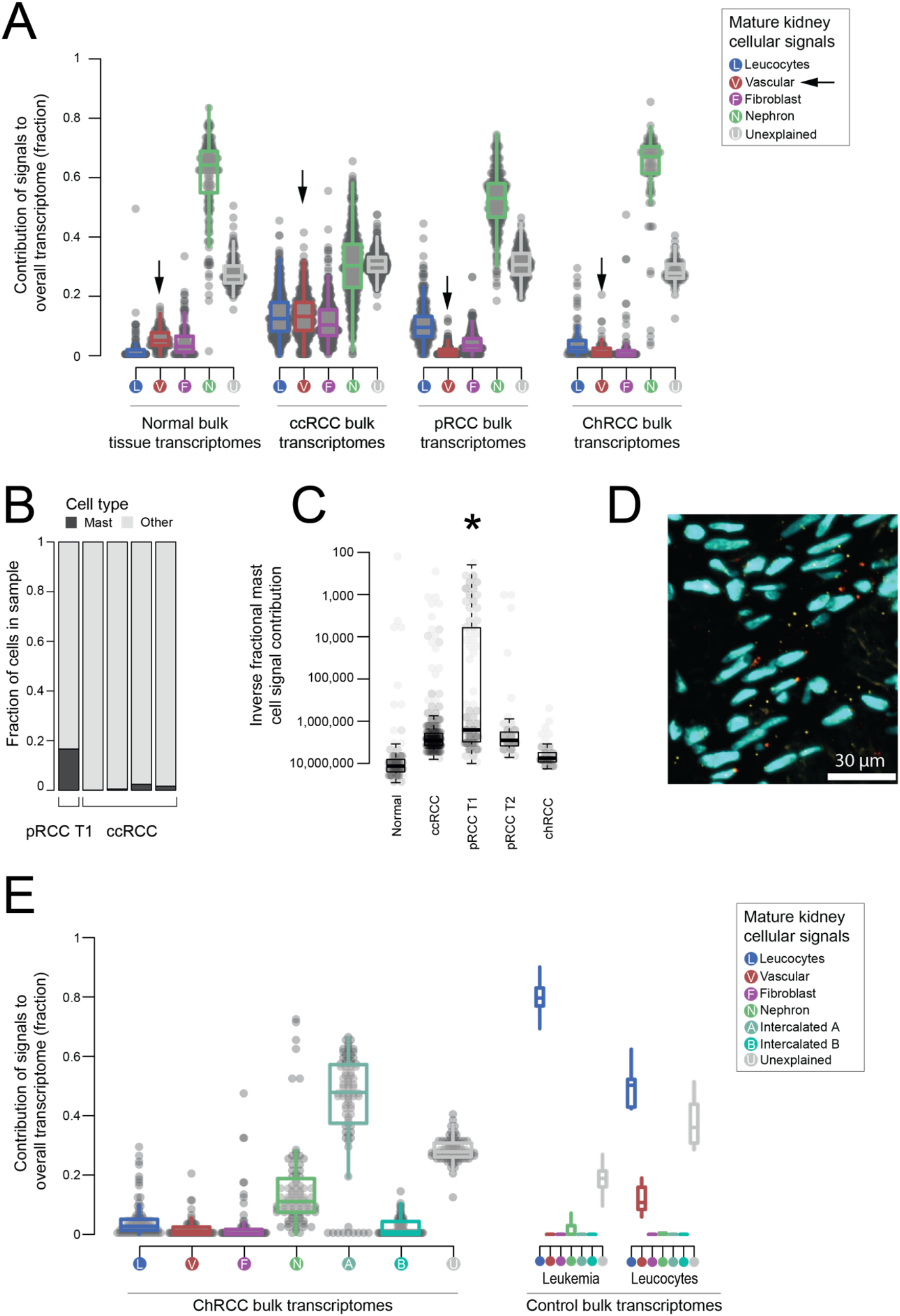
Adult kidney tumors. **A. Bulk renal cell carcinomas compared to mature kidney:** The relative contribution of single cell derived signals from mature kidneys in explaining the bulk transcriptomes of 171 normal kidney biopsies, 500 clear cell renal cell carcinomas (ccRCC), 274 papillary renal cell carcinomas (pRCC), and 81 chromophobe renal cell carcinomas (ChRCC). The relative contribution of each signal to each bulk RNA-seq sample is shown by the y-axis. Each signal/sample combination is represented by a single point and the distributions of relative signal contributions to the bulk transcriptomes are summarized with boxplots. Each signal type is labelled with an abbreviation and colored as shown by the legend on the right. Signals are marked with a circle for mature kidneys. The contribution to the bulk transcriptomes for all leucocyte, vascular, and nephron signals are aggregated together. **B. Mast cell fraction in single cell RCC samples:** Bar height indicates the fraction of cells that are mast cells (black) or other (grey) in 5 renal cell carcinoma single cell transcriptomic experiments (x-axis labels). **C. Mast cell signals in bulk RCC transcriptomes:** Each dot represents a bulk transcriptome of type indicated on the x-axis. The y-axis indicates the inverse of the mast cell signal for each bulk transcriptome. Boxplots show the distribution of mast cell signals across each sample type and the star indicates that mast cell signals are significantly higher in pRCC T1 type tumors (Wilcoxon rank-sum test, p< 1e-4). **D. smFISH validation:** An example section of single molecule fluorescence in-situ hybridization imaging of a pRCC T1 tumor section. Nuclei are stained blue with dapi and expression of the tumor marker *MET* is shown in yellow and the mast cell marker *TPSB2* in red. See **Table S5** for a quantification of smFISH applied to pRCC T1/T2 and ccRCC tumours. **E. Bulk chromophobe renal cell carcinomas compared to mature kidney:** The relative contribution of single cell derived signals from the mature kidney in explaining the bulk transcriptomes of 81 chromophobe renal cell carcinomas (ChRCC), along with control leucocyte and ALL populations. The relative contribution of each signal to each bulk RNA-seq sample is shown by the y-axis. Each signal/sample combination is represented by a single point and the distribution of relative signal contributions to the bulk transcriptomes are summarized with boxplots. Each signal type is labelled with an abbreviation and colored as shown by the legend on the right. Signals are marked with a circle for mature kidneys. ChRCC samples are shown in the blocks on the left, while the two groups of control samples are shown on the right. The samples marked “Leucocyte” are bulk transcriptomes from flow sorted leucocytes, while the samples marked “Leukemia” represent B-precursor acute lymphoblastic leukemia.

Continuing our investigation of the tumour microenvironment, we observed mast cells to be over-represented in single cell data derived from pRCCs (**Fig. 6B**). Performing cellular signal analysis revealed a high contribution of mast cell signal in a subset of tumors, significantly enriched for type 1 pRCC tumors (p<1e-4, Wilcoxon rank-sum test; **Fig. 6C**). This finding was further validated by single molecule fluorescence in-situ hybridization (smFISH), which found a higher fraction of mast cells a type 1 pRCC sample, than type 2 or ccRCC (**Fig. 6D, Table S5**).

Previous analyses of chromophobe cell renal cell carcinoma (ChRCC) have shown that ChRCC exhibit expression profiles of collecting duct cells^28^. Controversy exists as to whether the normal cell correlate of ChRCC is the type A or type B intercalated cells^29^. This is in part due to ChRCC retaining expression of both canonical markers of intercalated cells, SLC4A1 and SLC26A4 respectively (**Fig. S12**). Using cell signal analysis, which considers the entire transcriptomes of type A and type B cells, rather than just two markers, revealed a uniform type A signal across all chromophobe tumors (**Fig. 6E, S13**), bar the lethal variant of so-called metabolically divergent tumors (**Fig. 2B, S13**). The proliferation and active remodeling of type A cells has been demonstrated under conditions of systemic acidosis^30^, lending further credence to their possible status as the cell of origin for ChRCCs.

### Single cell signals provide diagnostic clues

An overarching finding of our study was that each tumor type possesses a particular pattern of cellular signals that were uniform in, and specific to, bulk transcriptomes from individual tumor types (**Fig. 7A-B, S14**). Accordingly, cellular signal assessment of bulk transcriptomes may provide sensitive and specific diagnostic clues. To test this proposition, we examined cellular signals in the bulk transcriptome of a histologically undefinable metastatic primary renal tumor from an 11 year old boy. Following resection, the tumor was examined histologically locally and by international reference renal pathologists (**Fig. 7C**). A definitive diagnosis could not be reached although an adult type renal cell carcinoma was favored. Nevertheless, the child was treated as a Wilms-like tumor, with cytotoxic chemotherapy and radiotherapy, following nephrectomy. He remains in complete remission two years following diagnosis, thus retrospectively suggesting a diagnosis of a Wilms-like tumor, as adult type kidney carcinomas do not respond to cytotoxic treatment.

**Figure 7 –.**
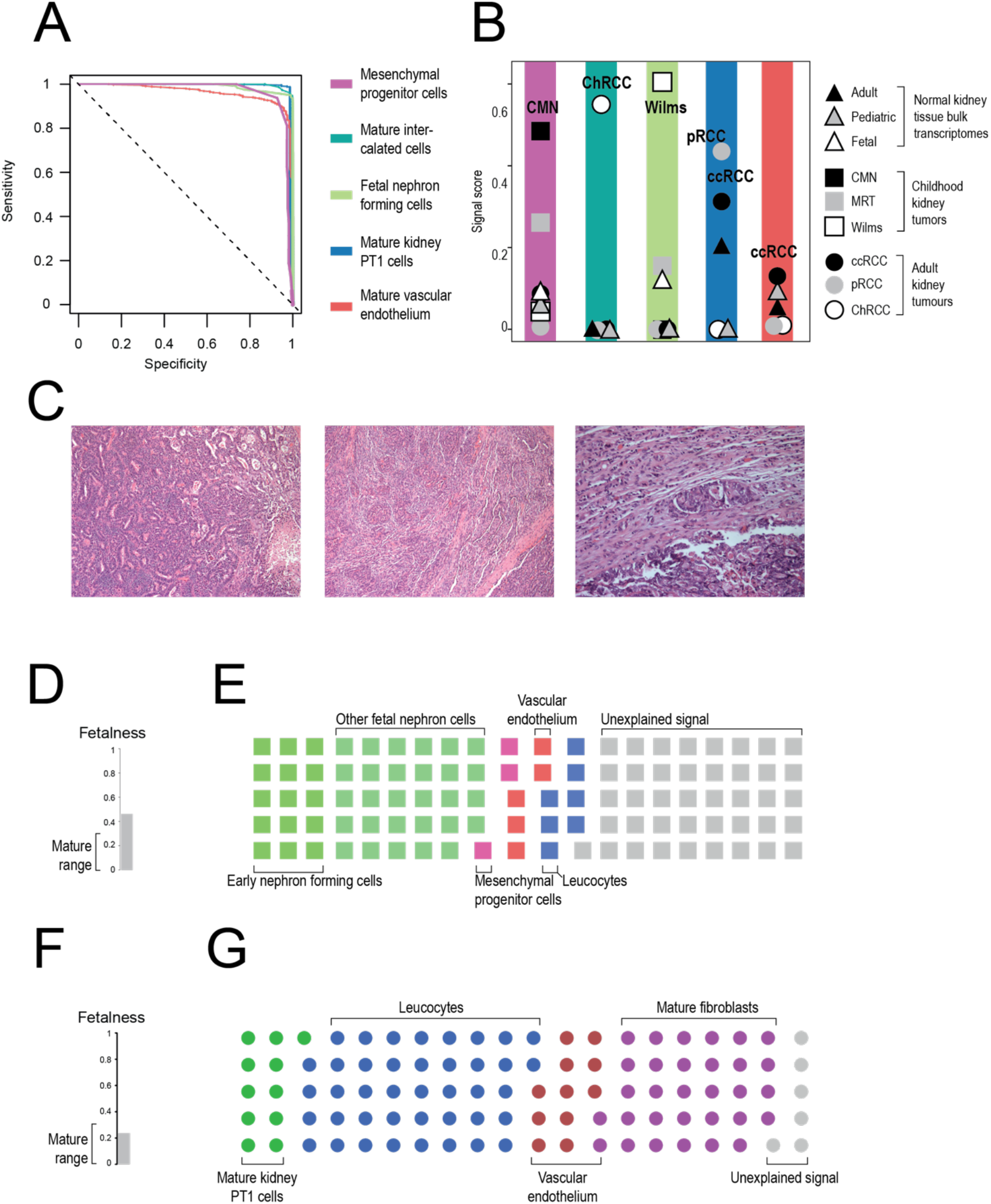
Clinical utility of cellular signal analysis. **A. Sensitivity/Specificity of signals in classifying tumor types:** Curves showing the sensitivity and specificity of using the scores defined by the color scheme to classify tumors by type at different cut-offs. The different score and tumor type pairs are: fetal interstitial cells and CMN (light blue), intercalated cells and ChRCCs (dark blue), the developing nephron and Nephroblastoma (light green), PT1 and ccRCCs/pRCCs (dark green), and mature vascular and ccRCCs (red). **B. Median reference contribution by tumor type:** Each point represents the median score for the group of samples indicated by the combination of shape and shading, as explained by the legend on the right, for the score type specified by the background shading. Score types are the same as in **A**. **C. Histology image of unclassified childhood renal tumor:** The tumor mostly compromised pleomorphic epithelioid cells that formed tubules, papillae, glands and nests, as well as more solid areas with spindled cells and clefting similar to that of synovial sarcoma. Patchy tumor necrosis was apparent. Some areas showed smaller, more uniform cells lining narrow tubular structures, resembling adenomatous perilobar nephrogenic rests. Overall, the morphology and ancillary tests were inconclusive. **D. Immaturity score for unclassified childhood renal tumor:** The immaturity score for the unclassified childhood renal tumor, calculated as in **Fig. 2**. The range of immaturity scores found in the normal mature kidney are indicated with the label on the left. **E. Summary of signal contribution from fetal and mature kidney to unclassified childhood renal tumor:** The relative contribution of single cell derived signals from the fetal kidney in explaining the bulk transcriptomes of the unclassified tumor from **C**. The 100 squares are colored so the fraction of squares of each color matches the fractional contribution of each fetal kidney signal contributes to explaining the bulk transcriptome of this sample. The labels above and below indicate what each color represents. **F. Immaturity score for childhood renal cell carcinoma:** The immaturity score for a childhood renal cell carcinoma, calculated as in **Fig. 2**. The range of immaturity scores found in normal mature kidney are indicated with the label on the left. **G. Summary of signal contribution from fetal and mature kidney to childhood renal cell carcinoma:** As in **D/E** but for a transcriptome derived from renal cell carcinoma fit using a mature kidney signal set.

We performed bulk mRNA sequencing on tumor specimens from this patient. Assessment of mRNA signals in bulk tissue suggested that the tumor exhibited a fetal transcriptome with cellular signals consistent with a Wilms-like tumor (**Fig. 7D-E**). The transcriptional diagnosis of a Wilms-like tumor was further substantiated by analyses of whole genome sequences. The tumor harbored classical somatic changes of Wilms, namely canonical *CTNNB1* and *KRAS* hotspot mutations and uniparental disomy of 11p (**Fig. S15**). By comparison, when we assessed single cell signals of an adult-type ccRCC that developed in a 15 year old adolescent, we found an overall mature transcriptome. Furthermore, the tumor exhibited the PT1 signal of ccRCC as well as a stark vascular endothelial typical of ccRCC (**Fig. 7F-G**).

## Discussion

We have determined normal cell signals in the major types of human renal tumors. This has enabled us to replace the approximate notion of the “fetalness” of childhood renal tumors with quantitative transcriptional evidence that the entire spectrum of pediatric renal tumors represent an aberrant developmental state. At the same time, our analyses question the suggestion that adult, epithelial-derived kidney cancers dedifferentiate towards a fetal state. Importantly, when we found transcriptional evidence of dedifferentiation in adult tumors, it conferred a dismal prognosis. Furthermore, amongst childhood tumors we found examples of cell signals representing differentiation trajectories, such as the neural crest to mesenchyme conversion in MRT. By contrast, the different types of adult tumors resembled specific renal tubular cells.

A central question that our findings raise is whether mRNA signals point to the cell of origin of tumors. When the similarity between mRNA signals and specific cell types was high, as found in most tumor types, this may be a plausible proposition. For example, in CMN, which typically occurs within the first weeks of life, our analysis identified an early mesenchymal progenitor cell population, characterized by the disease-defining oncogenes of CMN, as the likely cell of origin of CMN. In some tumors, transformation may entirely distort and obliterate gene expression profiles of the cell of origin. We found CCSK transcriptomes to represent such an extreme modification of the transcriptome of the developing kidney A further finding of our study was that within each category, the majority of tumors exhibited remarkably uniform cellular signals. This indicates that there are overarching transcriptional features, beyond individual gene markers, that unite tumor entities despite underlying intra- and inter-tumor genetic heterogeneity. Therefore, cellular signals of renal tumors may lend themselves as diagnostic adjuncts, as illustrated here by our ability to resolve the identify of a histologically unclassifiable childhood tumor. Moreover, the cellular transcriptome itself may represent a therapeutic target that transcends individual patients, if we had tools available to manipulate transcription in a predictable manner. This may be a particularly attractive approach for targeting transcriptional states of fetal cells retained in childhood cancer that are absent from normal post-natal tissues.

Overall our findings attach specific cell labels to human renal tumors that are underpinned by quantitative molecular data obtained from single cell mRNA sequences, independent of the interpretation of marker genes. As reference data from single cell transcriptomes expand through efforts such as the *Human Cell Atlas*, it will be feasible to annotate existing large repositories of tumor bulk transcriptomes, to derive a cellular transcriptional definition of human cancer.

## Acknowledgments

We acknowledge funding from Wellcome (Fellowships to S.B., K.S., J.C.A., core funding to the Sanger Institute, strategic award 211276/Z/18/Z). Additional support was provided by the St Baldrick’s Foundation (S.B.), Great Ormond Street Hospital Children’s Charity (J.C.A.), Great Ormond Street Hospital Biomedical Research Centre, Olivia Hodson Cancer Fund (K.S.), CRUK (Fellowship to T.J.M.), National Institute for Health Research (J.C.A., R.A.B.). The views expressed are those of the authors and not necessarily those of the National Health Service, National Institute for Health Research, or Department of Health. The CUH adult renal cancer sampling had infrastructure support from the Cancer Research UK Cambridge Cancer Centre Urological Malignancy Program and NIHR Biomedical Research Centre. J.D. acknowledges funding from the European Research Council (ERC) starting grant 850571, Dutch Cancer Society (KWF/Alpe d’HuZes Bas Mulder Award; KWF/Alpe d’HuZes, #10218), Foundation Children Cancer Free (KiKa #338, L.C.), Oncode Institute. We thank Dr Amos Burke for his contribution. We are grateful to our patients, young and old, for participating in our study.

## Author Contributions

M.D.Y. and S.B. conceived of the experiment and wrote the manuscript. M.D.Y. performed analyses, aided by T.J.M., E.K., G.K., and T.H.H.C. I.D.V. and J.C.A. provided expertise on adrenal gland analysis. L.C. performed organoid experiments with F.A.V.B. T.R.W.O., N.S., D.R., N.C., L.H., R.K., A.W. provided pathological expertise. F.C., M.M.H.E., and A.S. provided clinical data. A.P., E.B.B., F.M., C.T., C.B., G.D.S., V.J.G., M.H., M.K., S.M.P., O.A.B., K.R., K.K, F.H., J.D., F.C.H., E.P., K.A., contributed to discussions and / or data. S.T., T.M., F.H., R.R.K, contributed fetal and tumor single cell data, together with K.B.M., R.A.B., X.H., A.W.C, L.M. S.B. and M.D.Y. directed the study, in conjunction with K.S. (single cell cancer work) and J.D. (organoid work).

