## Supplementary Materials for "Single cell derived mRNA signals across human kidney tumors"

### Supplementary Material

#### Ethics statement

All human material was obtained through ethically approved studies with the following references: NHS National Research Ethics Service reference 03/018 (DIAMOND study; adult kidney tissues); NHS National Research Ethics Service reference 16/EE/0394 (pediatric tissues); NHS National Research Ethics Service reference 96/085 (fetal tissues). Organoids were generated from human tissue as approved by the medical ethics committee of the Erasmus Medical Center (Rotterdam, the Netherlands). Additional fetal tissue was provided by the Joint MRC / Wellcome Trust-funded (grant # 099175/Z/12/Z) Human Developmental BiologyResource (HBDR, http://www.hdbr.org; (10)), with appropriate maternal written consent and approval from the Newcastle and North Tyneside NHS Health Authority Joint Ethics Committee. HDBR is regulated by the UK Human Tissue Authority (HTA;www.hta.gov.uk) and operates in accordance with the relevant HTA Codes of Practice.

#### Tissue processing and data generation

##### 10X single cell sequencing of fresh tissue and bulk sequencing of DNA/RNA

Fresh tissues were processed to generate single suspensions for processing on the Chromium 10X controller (V2/3 3' chemistry), as previously described[1]. The MRT and normal kidney tissues organoids were derived and maintained as previously described[2]. Libraries were produced according to the manufacturer's instructions and sequenced on an Illumina HiSeq4000 device. Sequencing of bulk RNA and DNA was performed as previously described[1].

##### Cell-Seq2 experiments

Following resection, a random piece was selected from viable tumour tissue, minced, and viably frozen. On the day of the sorting, the sample was thawed and dissociated into a single-cell suspension in AdDF+++ (Advanced DMEM/F12 containing 1× Glutamax, 10 mM HEPES and antibiotics) containing Collagenase 1a (1mg/mL, Sigma, C9407) and DNase (0.25µg/mL, Stemcell), supplemented with Rho-kinase inhibitor Y-27632 (10 µM, Abmole). The samples were digested on an orbital shaker for 30 min at 37 °C. The suspension was washed first with AdDF+++ and next with MACS buffer (PBS pH 7.2+2mM EDTA +0.5% Bovine Serum Albumine), followed each time by centrifugation at 300×g. Viable single cells were sorted based on forward/side scatter properties and DAPI/DRAQ5 staining using FACS (MoFlo Astrios EQ, Beckman Coulter) into 384-well plates (Biorad) containing 10 µl mineral oil (Sigma) and 50 nl of RT primers.

##### 10X single nuclei sequencing

Single nuclei were isolated from frozen tissue using a glass dounce homogeniser. Samples were homogenised in buffer A (Sucrose 0.25 M, BSA 10 mg/ml, $MgCl_2$0.005 M, protease inhibitors and RNAse inhibitors RNAseIn – 0.12 U/ul and Superasin 0.06 U/ul), using ~25 strokes with the “loose” pestle and ~20 strokes with the “tight” pestle. Nuclei were cleaned up using a 30% Percol gradient and resuspended in buffer B (Sucrose 0.32 M, BSA 10 mg/ml, CaCl2 3 mM, MgAc2 2 mM, EDTA 0.1 mM, Tris-HCl 10 mM, DTT 1mM in the presence of protease and RNAse inhibitors as in buffer A).

Nuclei were mixed 1:1 with Tryphan blue and counted using a disposable haemocytometer, then diluted to the appropriate concentration. Nuclei were loaded on to the 10X Chromium controller as per the Chromium Single Cell 3′ Reagent Kits v3 User Guide, targeting to recover 5000 nuclei. Post GEM-RT cleanup, cDNA amplification and 3’ gene expression library construction were carried out according to the user guide. The resulting libraries were sequenced on the Novaseq platform.

##### Immunohistochemistry of MRT tissue

Immunohistochemistry was performed on 3–4 µm sections of tissue fixed in 4% paraformaldehyde, dehydrated and embedded in paraffin according to standard protocols. Sections were subjected to H&E and immunohistochemical staining using antibodies for INI-1 (BD Transduction Laboratories, 612111, 1:400) or TWIST (Abcam, ab50581, 1:500). Counterstaining was performed using Mayer’s Hematoxylin (1:3 dilution). The Leica DMi8 microscope was used for imaging.

##### RNAscope smFISH and immunohistochemistry

FFPE tissue sections of 5 µm thickness were processed using a Leica BOND RX to automate staining with the RNAscope Multiplex Fluorescent Reagent Kit v2 Assay and RNAscope 4-plex Ancillary Kit for Multiplex Fluorescent Reagent Kit v2 (Advanced Cell Diagnostics, Bio-Techne) in combination with immunohistochemistry (IHC) for PECAM1[3]. Owing to intense tissue autofluorescence, samples were treated with a photobleaching procedure based upon that of Lin et al.[4]. It was observed that photobleaching prior to RNAscope probe hybridisation adversely affected smFISH staining, presumably due to loss of RNA integrity in the alkaline solution. Therefore, photobleaching was conducted following RNAscope probe and tree amplification reagents (AMP1/2/3) but before channel-specific HRP reagents and fluorophores. Initial automated processing included baking at 60°C for 30 minutes and dewaxing, as well as heat-induced epitope retrieval at 95°C for 15 minutes in buffer ER2 and digestion with Protease III for 15 minutes. Following RNAscope probe and AMP hybridisation according to the manufacturer’s instructions, slides were briefly rinsed in PBS and then subjected to photobleaching. Slides were incubated horizontally in 4.5% hydrogen peroxide, 24 mM sodium hydroxide in PBS in a Nunc Square BioAssay Dish atop a white light box for 30 minutes. Slides were thoroughly rinsed with sterile deionised water and Leica BOND wash before RNAscope staining was resumed with the sequential development of three probe channels using tyramide signal amplification with Opal 570, Opal 650 (both Akoya Biosciences) and TSA-biotin (TSA Plus Biotin Kit, Perkin Elmer) and streptavidin-conjugated Atto 425 (Sigma Aldrich). Finally, IHC was carried out, beginning with a blocking step of 1 hour in Primary Antibody Diluent (Leica), followed by rabbit anti-PECAM1 (Abcam ab28364) at 1:600 at room temperature for 2 hours, and then HRP goat anti-rabbit IgG (Thermo G21234) at 1:1000 at room temperature for 1 hour. Both antibodies were diluted in Primary Antibody Diluent. IHC signal was developed using Opal 520 (Akoya).

Stained sections were imaged with a Perkin Elmer Opera Phenix High-Content Screening System, in confocal mode with 1 µm z-step size, using a 20× water-immersion objective (NA 0.16, 0.299 µm/pixel). Channels: DAPI (excitation 375 nm, emission 435-480 nm), Atto 425 (ex. 425 nm, em. 463-501 nm), Opal 520 (ex. 488 nm, em. 500- 550 nm), Opal 570 (ex. 561 nm, em. 570-630 nm), Opal 650 (ex. 640 nm, em. 650-760 nm).

#### Basic data processing and quality control

##### Mapping of DNA reads

DNA sequencing reads were aligned to the GRCh 37d5 reference genome using the Burrows-Wheeler transform (BWA-MEM)[5]. Sequencing depth at each base was assessed using Bedtools coverage v2.24.0.

##### Substitution calling

Single base somatic substitutions were called using an in-house version of CaVEMan v1.11.2 (Cancer Variants through Expectation Maximization)[6]. CaVEMan compares sequencing reads from tumor and matched normal samples and uses a naïve Bayesian model and expectation-maximization approach to calculate the probability of a somatic variant at each base (https://github.com/cancerit/CaVEMan). Small insertions and deletions (indels) were called using an in-house version of Pindel (v2.2.2; github.com/cancerit/cgpPindel). Post-processing filters required that the following criteria were met to call a somatic substitution:

1. At least a third of the reads calling the variant had a base quality of 25 or higher.
2. If coverage of the mutant allele was less than 8, at least one mutant allele was detected in the first 2/3 of the read.
3. Less than 5% of the mutant alleles with base quality ≥ 15 were found in the matched normal.
4. Bidirectional reads reporting the mutant allele.
5. Not all mutant alleles reported in the second half of the read.
6. Mean mapping quality of the mutant allele reads was ≥ 21.
7. Mutation does not fall in a simple repeat or centromeric region.
8. Position does not fall within a germline insertion or deletion.
9. Variant is not reported by ≥ 3 reads in more than one percent of samples in a panel of approximately 400 unmatched normal samples.
10. A minimum 2 reads in each direction reporting the mutant allele.
11. At least 10-fold coverage at the mutant allele locus.
12. Minimum variant allele fraction 5%.
13. No insertion or deletion called within a read length (150bp) of the putative substitution.
14. No soft-clipped reads reporting the mutant allele.
15. Median BWA alignment score of the reads reporting the mutant allele ≥ 140.

The following variants were flagged for additional inspection for potential artefacts, germline contamination or index-jumping event:

16. Any mutant allele reported within 150bp of another variant.
17. Mutant allele reported in >1% of the matched normal reads.

#### Copy number detection in bulk DNA

The ascatNGS algorithm (v4.0.1)[7] was used to estimate tumor purity and ploidy and to construct copy number profiles prior to running the Battenberg algorithm (v2.2.5) (github.com/cancerit/cgpBattenberg) to allow for tumor subclonality.

#### Bulk RNA mapping and quantification

Where possible, we have processed all bulk RNA-seq data using the exact same pipeline as the recount2 project[8]. That is, we used RAIL-RNA to produce counts of bases aligned to each gene in each sample[9]. Counts were then converted to fragments aligned to genes by dividing counts by the average fragment length for the sample. This approach allowed us to combine our in-house data with any dataset processed by the recount2 project, in particular the TCGA and GTEX projects.

The length for each gene was calculated as the sum of unique exonic bases for all transcripts associated with each gene. We used the gencode v25 GTF annotation[10] and GRCh38 human reference genome.

In order to run the recount2 pipeline, we required access to the sequencer output (BAM files or fastq). In some cases, we only had access to processed data, either in the form of raw fragment counts, or transcripts per million (TPM). TPMs were converted to fragment counts by multiplying by 1 million, rounding to an integer and assigning each gene an effective length of 1.

Where TPM values were needed for direct comparison of gene expression, we calculated TPM values from fragment counts by dividing by gene length, then normalizing the counts/bp by forcing them to sum to 1,000,000 across all genes in a sample.

#### Single cell RNA mapping, quantification, quality control and normalization

Single cell RNA-seq data were quantified using the 10X software package cellranger (version 2.0.2 for V2 chemistry, 3.0.2 for V3 chemistry) to map sequencing data to version 2.1.0 of the build of the GRCh38 reference genome supplied by 10X.

Data were normalized for sequencing depth by dividing by the total number of UMIs in each cell and then transformed to a log scale for each cell using the Seurat version 3.1.4[11] NormalizeData function. That is, the transformed data, y, is given by:

$$y_{gc} = \log\left(1 + F\frac{x_{gc}}{\sum_g x_{gc}}\right)$$

where x is the UMI count matrix with g indexing gene and c indexing the cell. F is the Seurat "scale.factor" parameter (which we left at the default value of 10,000).

Doublets were determined using scrublet[12] and ambient RNA contamination was removed with SoupX[13]. To filter lower quality cells, we performed high resolution clustering (Seurat graph-based clustering with resolution = 10) and filtered any cell which:

1) had greater than 5% expression originating from mitochondrial genes
2) was marked as a doublet
3) expressed fewer than 500 distinct transcripts
4) or belonged to a cluster where greater than 50% of cells failed one of 1-3.

The rationale behind this approach was to conservatively remove cells with a very similar transcriptome to cells which have failed QC.

To prevent similarity to reference maps (e.g. fetal kidney) being driven by cell cycle state, we also removed any cell with evidence of being in S or G2M phase. We determined the cell cycle phase by scoring each cell based on panel of genes specific to each phase using the Seurat CellCycleScoring function. We also removed all leucocytes from each tissues reference map.

### Analysis of processed data

#### Derivation of color scheme

In deriving a color scheme to represent the different types of cellular signal used in this paper, we started by designating a series of hue ranges to represent each tissue type. These hue ranges were then further sub-divided to represent more specific cell types. To separate fetal and mature versions of the same cell type, we used different values of the "value" parameter in hue, saturation, value color space to represent fetal (0.9) and mature (0.7) cell signals. Finally, we set the saturation value to 0.6 by default and allowed this to vary as necessary to emphasize differences between cell types with otherwise similar colors. This color scheme is summarized in **Fig. S16**.

We also constructed a color scheme for each sample type in this study. We used light/pastel colors to represent non-tumor or control samples and solid colors for tumors. We used the same color to represent Neuroblastoma and ChRCC tumors as they were never referenced in the same plot. This color scheme is summarized in **Fig. 17**.

#### Dimension reduction and cluster generation of single cell RNA data

Following normalization, we identified genes with high variability using the Seurat FindVariableGenes function. This function calculates the mean expression and dispersion for each gene, then groups genes into bins (of size 20) by their mean expression and identifies any gene for which the z-score calculated from the dispersion exceeds some cut-off. We used the default cut-off of z=1 and mean expression in the range 0.1 to 8.

The normalized data was scaled to have mean 0 and standard deviation 1 and principle component analysis was performed using the variable genes identified together with any gene that we identified as being potentially biologically interesting (regardless of its variability in the data).

We determined the optimal number of principle components (PCs) using molecular cross validation (https://github.com/constantAmateur/MCVR)[14]. We used these to construct a two-dimensional representation of the data using either tSNE[15,16] or UMAP[17]. This representation was then used only to visualize the data.

Clusters were identified using the community identification algorithm as implemented in the Seurat "FindClusters" algorithm. We used the number of PCs determined above as input to this method and set the resolution parameter to 1. We chose this value of the resolution parameter as it produced a number of clusters that was large enough to capture most of the important biological variability but not so large as to make detailed manual scrutiny of each cluster impractical. All other parameters were set to the function defaults.

#### Annotation of fetal kidney single cell data

To create the fetal kidney reference, we combined the raw 10X output from previous studies[1,18] together with data from 4 additional fetal kidneys. The combined data were quantified and clustered as described above, with the exception that the clustering resolution parameter was set to 2 to obtain a more granular annotation.

To annotate these clusters, we used a previously published detailed annotation of the fetal kidney as a reference[18]. That is, we first trained a logistic regression model on just the “PloS” data[18]. In training this model we used the elastic net regularization procedure with alpha=0.99 to produce strong regularization but prevent strongly co-linear genes being excluded. This model was fit using the “glmnet” R package[19].

To obtain regression coefficients specific to each cluster in our training data we fit a series of N binomial logistic regression models, where N is the number of clusters in the training data (i.e., one-versus-rest binomial logistic regression). To prevent the observed frequencies of cells (which we do not expect to accurately reflect the true abundances in situ) from biasing the regression coefficients we use an offset for each model given by,

$$\log\left(\frac{f}{1-f}\right)$$

where f is the fraction of cells in the cluster being trained.

In each case, we performed 10-fold cross validation and selected the regularization co-efficient, lambda, to be as large as possible (i.e., as few non-zero coefficients as possible) such that the cross validated accuracy was within 1 standard deviation of the minimum.

These models were then used to calculate a predicted similarity for each cell in the combined fetal kidney data set. In calculating the predicted values, an offset of 0 was used. Softmax normalization was not used to allow for the possibility that cell types were present in the combined reference not present in the “PLoS” map. Clusters with a similarity of less than 1 (logit scale) to any of the reference data were labelled as “undecided”.

Following the application of the logistic regression model, we elected to merge categories in the reference data that were commonly found in the same clusters. Specifically, we combined:

- NPCa, NPCb, and NPCc categories into CapMes.

- RVCSBa, RVCSBb into RVCSB.
- ErPrT and SSBpr into ErPrT

We removed Leu, Prolif, PTA and Mes as no cluster contained a majority of these cells. Each cluster was then annotated with whichever of the reference categories had the highest similarity score averaged across all cells in the cluster. This procedure left one cluster as "Undecided" (that is, most cells in these clusters could not be allocated unambiguously to one of the reference populations). Closer inspection of this cluster revealed in to be an early mesenchymal population, which we labeled as MPC for mesenchymal progenitor cells as discussed in the manuscript.

#### Additional reference signal sets

In addition to the above annotated single cell data sets, cellular reference signals were also taken from additional data sets:

1) A mature kidney single cell reference map[1].
2) The 10x demonstration PBMC data set[20], annotated as described here (https://satijalab.org/seurat/v3.0/pbmc3k_tutorial.html). This data set was used to define a set of leucocyte signals that were added to all other reference maps.
3) Fetal adrenal reference map[21]
4) Whole embryo mouse data[22]
5) A pan-tissue human reference from the human cell landscape publication[23]

#### Annotation of congenital mesoblastic nephroma single cell data

Single cell transcriptomes derived from a congenital mesoblastic nephroma were processed into clusters as described above. Following clustering, we assigned a cell type to each cluster using marker genes identified as in previous work[1,13].

To confirm that the cluster without expression of other known markers represented CMN tumor cells, we investigated the expression of ETV6, NTRK3 and EGFR in this cluster. CMN is known to be driven by an activating rearrangement between ETV6 and NTRK3, which results in a fusion product with the 5' end of ETV6 fused to the 3' end of NTRK3. As the 10X assay measures expression using a 3' enrichment strategy, CMN tumor cells should show a high level of NTRK3 expression. This is indeed what we find, with the cluster marked tumor expressing NTRK3 more than 10 times more strongly than the next closest cluster. By contrast, ETV6 was not highly expressed in this cluster, as would be expected. Finally, EGFR is known to be highly expressed by tumor cells and was also found to be highly expressed in the cluster designated as tumor.

#### Annotation of congenital malignant rhabdoid tumor cell data

For MRT single cell data, clusters were determined and marker genes identified as described above for CMN. Common non-tumor populations were annotated based on well known markers (**Fig. S9**). Tumor cells were identified by loss of the *SMARCB1*, except for MRT2 for which the molecular diagnostic workup did not identify mutation of *SMARCB1*. For this sample, tumor cells were identified based on the loss expression of other members of the SWI/SNF *SMARCA2* and *SMARCA4*.

##### Cell similarity inference using single cell data

To measure the similarity of a target single cell transcriptome to a reference single cell data-set we used the methodology based on logistic regression outlined in detail[1]. Briefly, we train a logistic regression model with elastic net regularization (alpha=0.99) on the reference training set. We then use this trained model to infer a similarity score for each cell in the query data set for each cell type in the reference data.

Softmax normalization was not used to allow for the possibility that some cells in the query data set do not resemble any of the cell types in the reference data set. Predicted logits were averaged within each cluster in the query dataset. This approach was implemented using the "glmnet" package in R[19].

###### Similarity to mouse data

In assessing the closest match to the organoid MRT data, we considered a comparison to a mouse reference dataset[22]. We performed the similarity analysis as described above, with the only difference being that we limit the analysis to orthologous genes as determined by ENSEMBL biomart.

##### Sensitivity and specificity of samples to particular cellular signals

To perform the sensitivity and specificity analysis (**Fig. 7A-B**) we first constructed a set of cellular signals that were indicative of a particular tumor type. These were MPCs and CMN, Intercalated cells and ChRCC, nephrogenic cells (i.e., cap mesenchyme, primitive vesicle, and ureteric bud) and Wilms tumor, PT1 and ccRCC/pRCC, and mature vasculature and ccRCC. We calculated all of these scores for every sample in our data set and then evaluated the sensitivity and specificity at different cut-offs for each score to construct the sensitivity/specificity curves in **Fig. 7A** (ROC curve).

##### Quantification of smFISH images

The tiled images exported from the Phenix were illumination corrected andstitched together into large, multi-channel fluorescence images by a specialized tool supplied by Perkin-Elmer. These images were then analysed in the Qupath[24] Bioimage Analysis program. First, all nuclei were segmented using Qupath's cell detection algorithm, then each nucleus was expanded by 3 microns to estimate the area covered by the entire cell. The subcellular spot detection option was then used to detect all fluorescent spots in all the RNA-SCOPE channels for the size range of 1-8 square microns. Each spot was automatically assigned to a detected cell. The detection data for each image was then exported as a .csv file for further analysis.

#### Method to quantify single cell derived signals in bulk transcriptomes

##### Data preparation

###### Bulk RNA-seq data

Each bulk RNA-seq sample required two pieces of information: fragment counts per gene and the effective length of each gene. As described above, gene counts for this paper were mostly generated using the Rail-RNA pipeline[9] and effective gene lengths as the sum of unique exonic bases per gene. However, fragment counts and effective gene lengths can be calculated in any way.

#### Single cell reference data

To calculate reference single cell signals, single cell data must first be clustered and annotated. Cells are then grouped together by annotation and raw counts summed across all cells within a group. Summed counts are then normalized to sum to 1 across all genes so that a reference signal is defined as a vector, S,

$$\mathbf{S} = \{s_1, s_2, \ldots, s_m\} \text{s.t.} \sum_{g=1}^{m} s_g = 1$$

where there are m genes.

Reference signals can be constructed in any way (e.g., including batch correction for combined data sets), so long as the final signal can be normalized such that $\sum_g s_g = 1$.

#### Model fit

The aim of the signal assignment method is to infer how much of each of the different reference signals best explains the supplied bulk transcriptome. That is, we are aiming to solve for the values of beta in,

$$y_{gp} = \beta_{0p} + \beta_{1p} s_{g1} + \beta_{2p} s_{g2} + \ldots + \beta_{np} s_{gn}$$

where $y_{gp}$ is the fragment counts for gene g in sample p, $s_{gc}$ is the reference signal c for gene g and $\beta_{cp}$ is the contribution of signal c to sample p. Note that the term $\beta_{0p}$ represents the contribution the intercept term for sample p, which is equivalent to the inclusion of an additional flat signature for which $s_g = constant \ \forall g$. The inclusion of this intercept term provides a measure of the extent to which the reference signal set is inappropriate for the sample given.

In order to efficiently calculate the contributions of the same set of reference signals simultaneously, we formulated the following model,

$$\beta_{cp} = e^{z_{cp}}$$

where c represents the signal and p the sample as above. We solve for $z_{cp}$ instead of $\beta_{cp}$ directly as this formulation ensures that the contributions of each signal are always strictly positive. Next, we calculate the expression for gene g and sample p implied by these values of $\beta_{cp}$

$$\lambda_{gp} = S_{gc} \beta_{cp}$$

where g represents the gene, and then calculate

$$\lambda'_{gp} = \lambda_{gp} l_{gp}$$

where $l_{gp}$ represents the effective length of gene g in sample p. This modification by length is necessary as the fragment counts by gene created by bulk RNA-seq are proportional to the length of a gene. Finally, a joint negative log likelihood is calculated as

$$-\log \mathcal{L} = \sum_p \sum_g w_g (\lambda'_{gp} - y_{gp} \log \lambda'_{gp})$$

where $w_g$ is an optional gene penalty applied to genes deemed to be biologically less important. In this paper we set $w_g$ to 1 for all genes except a set of housekeeping genes that are set to 0.5 and metabolic genes set to 0. It is this equation that is minimzed with respect to $z_{cp}$ in order to solve for the values of $\beta_{cp}$. This log-likelihood is simply the Poisson log-likelihood for a Poisson distribution with mean $\lambda'_{gp}$ (see section below for a discussion of this choice of distribution).

This model is directly specified using the tensorflow framework[25], which allows the efficient minimization of this equation utilizing graphics processors and multi-core machines. The optimization is performed using stochastic gradient decent using Adam[26]. We require two termination conditions to be met before the optimization is terminated:

1. the fractional decrease in the log-likelihood must be less than some tolerance parameter.
2. the fractional change in Q must be less than the same tolerance parameter.

Q is defined as the sum of the sigmoid transformation of $z_{cp}$ and roughly measures the number of signals with non-zero contributions to the fit. Without this second termination condition, optimization would terminate with the coefficients of many signals given small but non-zero values as these non-zero values barely shift the total log-likelihood.

#### Post processing

Having obtained optimized values for $\beta_{cp}$, we next normalize these values by first modifying the intercept term to

$$\beta'_{0p} = \beta_{0p}/m$$

which make the modified intercept term equivalent to fitting an additional signal with a completely flat profile. Following this modification, we then normalize the values of $\beta_{cp}$ to sum to 1 for each sample. These normalized values represent the relative contribution of each signal to each sample and are the values reported throughout this manuscript. The final normalization step essentially controls for differences in bulk RNA-seq library size and makes the coefficients comparable across samples.
As an additional measure of the goodness we re-fit the above model with only the intercept term and then calculate,

$$pR^2 = 1 - \frac{\log \mathcal{L}_{\text{full}}}{\log \mathcal{L}_{\text{int}}}$$

which is, a McFadden's pseudo R-squared value[27] $pR^2$ given by 1 minus the ratio of the log-likelihood of the full model fit over the model fit with only the intercept term.

In order to aid with interpretation of the fit, the contribution from similar cellular signals is often aggregated before being presented in the Figures and Supplementary Figures. For example, there are multiple endothelial signals in the mature kidney, but for simplicity and readability we have combined them throughout this study.

#### Benchmarking

We compared our method to two other methods: MuSiC[28] and BSeq-SC[29]. In both cases we used the default settings recommended by each method. As MuSiC requires a reference containing multiple cells derived from multiple samples, we were unable to create include leucocytes as part of our reference panel for **Fig. 1D** for MuSiC. BSeq-SC required marker genes for each population in the reference. To generate markers for each reference population we identified genes significantly enriched in the target population with a binomial test, using the quickMarkers function in the SoupX R package[13]. We further refined this set of markers by requiring that markers be expressed in at least 40% of cells within the cluster they mark and less than 5% of all other clusters. With these requirements, some clusters, notably the PT1 cluster, did not have any marker genes and so received a value of 0 in the BSeq-SC fit.

#### Calibration of intercept term

To assess the range of contributions from the intercept term (i.e., the "unexplained signal") when no appropriate reference is provided we constructed a series of inappropriate fits. In each case we selected a cell signal reference set that we knew was inappropriate to the set of samples being considered. The range of intercept values in this fit then gives a quantitative range that is indicative of how much weight is given to the intercept when no appropriate reference is present.

#### Choice of Poisson distribution

The choice of the Poisson distribution as the likelihood model at first glance seem a curious one, given that the Negative Binomial distribution (of which the Poisson distribution is a specific case) is widely used to model both bulk RNA-seq and single cell RNA-seq data. However, some reflection reveals that this choice is actually well justified.

We wish to evaluate the probability of observing a particular number of fragment counts in a bulk RNA-seq experiment, given that this experiment is composed of the addition of signals from a collection of single cell derived transcriptomic signals. To do this, we need to know how likely a particular set of fragment counts is, given a fixed contribution from each of the single cell signals.

Let us assume that the number of counts for a gene g in cell type c in a single cell RNA-seq experiment (with a fixed number of reads) can be well modelled by a negative binomial distribution with mean $\mu$ and over-dispersion $\varphi$. That is, the variance of this distribution is given by,

$$\sigma_{gc}^2 = \mu_{gc} + \mu_{gc}^2 \phi_{gc}$$

The distribution we are interested in, is then the distribution resulting from the sum of $\{N_0, N_1, N_2, \ldots, N_k\}$ random samples from the set of negative binomial distributions representing cell types {1,2,…,k}. That is, the distribution we are interested in is given by the sum of negative binomial distributions.

The moment generating function for the compound distribution is then given by,

$$M_{\text{comp}}(t) = \prod_{c \in C} \left(1 + \mu_c \phi_c (1 - e^t)\right)^{-\frac{1}{\phi_c}}$$

where C is the set of signals summed to form the compound distribution. This moment generating function completely specifies the compound distribution. However, we can use the method of moments to approximate this compound distribution with another Negative Binomial distribution with mean μ and over-dispersion φ. To do this, observe that the first and second moment of the compound distribution are,

$$\begin{aligned} E(X) &= \sum_{c \in C} \mu_c \\ E(X^2) &= \sum_{c \in C} \mu_c + \left(\sum_{c \in C} \mu_c\right)^2 + \sum_{c \in C} \mu_c^2 \phi_c \end{aligned}$$

while the first and second moments of a negative binomial distribution with mean μ and over-dispersion φ are,

$$\begin{aligned} E(X) &= \mu \\ E(X^2) &= \mu + \mu^2 + \mu^2 \phi \end{aligned}$$

Using the method of moments, this implies that the negative binomial approximation to the compound distribution has,

$$\begin{aligned} \mu &= \sum_{c \in C} \mu_c \\ \phi &= \sum_{c \in C} \left(\frac{\mu_c}{\mu}\right)^2 \phi_c \end{aligned}$$

That is, the mean of the compound distribution equals the sum of the means of each component distribution (as expected). The over-dispersion of the compound distribution is equal to the weighted sum of the component distributions. Closer consideration of the equation for the compound over-dispersion reveals that the over-dispersion of the compound distribution is almost always considerably **less** than the average over-dispersion of its component distributions.

For example, consider the case where all distributions have approximately the same mean and rewrite the compound over-dispersion as,

$$\phi = \sum_{c \in C} \left( \frac{\mu_c}{< \mu_c >} \right)^2 \frac{\phi_c}{N^2}$$

where angle brackets denote an average and N is the number of elements in C. Assuming the ratio in brackets is close to 1 gives,

$$\phi = \frac{< \phi_c >}{N}$$

So, in the case of distributions with similar means, the over-dispersion of the compound distribution is always N times less than the mean over-dispersion of the individual distributions. Consequently, as the number of distributions being summed over increases, the over-dispersion goes to zero and the compound Negative Binomial distribution approaches a Poisson distribution. The more general case where the means of the component distributions are not all similar is more complex, but in the limit of many distributions, the compound over-dispersion still approaches 0.

This result justifies the use of a Poisson distribution as the likelihood model in our fitting procedure. Although the individual signals from which the fit is derived are negative binomially distributed, the distribution of their sum is Poisson distributed. It may be that an extension of the Poisson model used here may prove useful, to model effects such as uncertainty in the effective length of genes for example, but it is not required to accurately represent the compound distribution on which our model depends.

#### Data availability

Raw sequencing data have been deposited in the EGA. Raw tables of counts for bulk RNA seq data are available in **Data S1** and are described in **Table S1**. Raw tables of counts for single cell data are available in **Data S2** and are described in **Table S2**. The data used to generate all figures in this manuscript are made available in **Data S3**.

#### Code availability

As an annex to the supplementary methods, we have provided all source code used in generating the results, figures, and tables used in this study as **Data S4**. The purpose of these code files is to provide additional details as to how we implemented the analyses described in the **Methods** section.

### Supplementary Figures

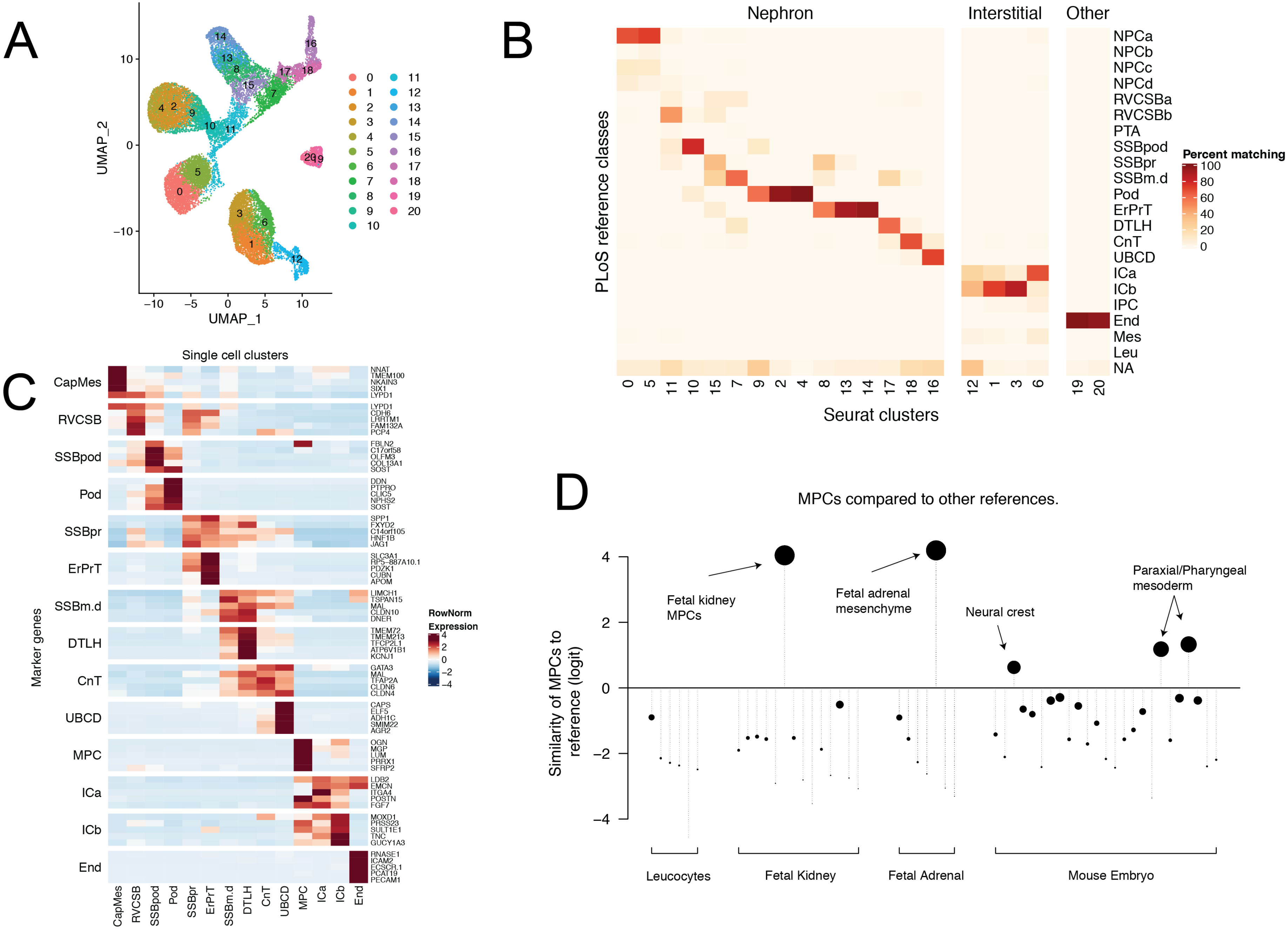
A
UMAP_1
UMAP_2
0 1 2 3 4 5 6 7 8 9 10 11 12 13 14 15 16 17 18 19 20
B
Nephron
Interstitial
Other
PLoS reference classes
NPCa
NPCb
NPCc
NPCd
RVCSBa
RVCSBb
PTA
SSBpod
SSBpr
SSBm.d
Pod
ErPrT
DTLH
CnT
UBCD
ICa
ICb
IPC
End
Mes
Leu
NA
Percent matching
Seurat clusters
C
Single cell clusters
Marker genes
CapMes
RVCSB
SSBpod
Pod
SSBpr
ErPrT
SSBm.d
DTLH
CnT
UBCD
MPC
ICa
ICb
End
RowNorm Expression
NNAT TMEM100 NKAIN3 SIX1 LYPD1
LYPD1 CDH6 LRRTM1 FAM132A PCP4
FBLN2 C17orf58 OLFM3 COL13A1 SOST
DDN PTPRO CLIC5 NPHS2 SOST
SPP1 FXYD2 C14orf105 HNF1B JAG1
SLC3A1 RP5−887A10.1 PDZK1 CUBN APOM
LIMCH1 TSPAN15 MAL CLDN10 DNER
TMEM72 TMEM213 TFCP2L1 ATP6V1B1 KCNJ1
GATA3 MAL TFAP2A CLDN6 CLDN4
CAPS ELF5 ADH1C SMIM22 AGR2
OGN MGP LUM PRRX1 SFRP2
LDB2 EMCN ITGA4 POSTN FGF7
MOXD1 PRSS23 SULT1E1 TNC GUCY1A3
RNASE1 ICAM2 ECSCR.1 PCAT19 PECAM1
D
MPCs compared to other references.
Similarity of MPCs to reference (logit)
Fetal kidney MPCs
Fetal adrenal mesenchyme
Neural crest
Paraxial/Pharyngeal mesoderm
Leucocytes
Fetal Kidney
Fetal Adrenal
Mouse Embryo

**Supplementary Figure1 – Fetal kidney reference map**

A. **UMAP of fetal kidney clusters –** Reduced dimension representation (UMAP) of the transcriptomes of 21,834 fetal kidney cells where each point represents a cell and nearby cells have similar transcriptomes. Each cell is labelled by a color corresponding to the cluster to which it belongs, as indicated by the legend on the right. Additionally, a label for each cluster is placed at the mean position of cells belonging to each cluster. Clusters are generated as discussed in the methods section.
B. **Comparison of clusters to PLoS annotation –** Each cell is assigned an annotation from [18] based on logistic regression, or NA if the similarity score is less than 1 (logit) or multiple populations have similarity greater than 1. The x-axis shows the clusters from **A,** the y-axis the reference populations and the color scale the fraction of cells assigned to each reference class. Note the high proportion of NAs in cluster 12.
C. **Marker genes defining cell types** – For each annotated cell population in **Fig. 1B**, the top 5 algorithmically determined marker genes are shown. The color scheme indicates the average normalized expression of the cells in the cluster indicated on the x-axis, z-scaled across all cell types to have mean 0 and standard deviation 1.
D. **Similarity of MPCs to other tissues** – Similarity score (logits) calculated by logistic regression trained on the fetal kidney reference to MPCs. Note that the extremely high similarity to the equivalent population in the fetal adrenal.

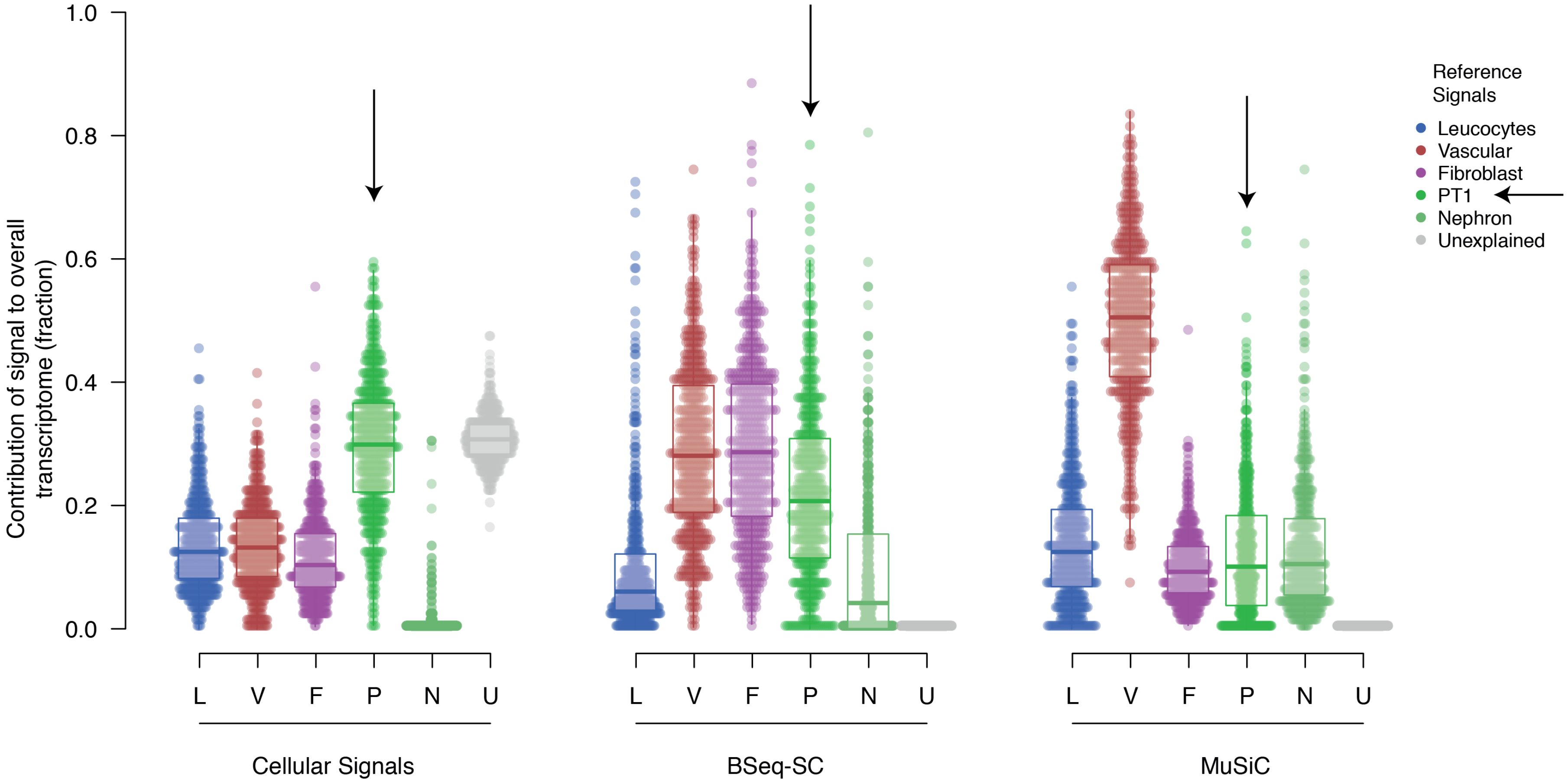

1.0
0.8
0.6
0.4
0.2
0.0
Contribution of signal to overall transcriptome (fraction)
L
V
F
P
N
U
Cellular Signals
BSeq-SC
MuSiC
Reference Signals
Leucocytes
Vascular
Fibroblast
PT1
Nephron
Unexplained

**Supplementary Figure 2 – Ability of different methods to recover known RCC signal**

We applied our cellular signal method, BSEQ-sc and MuSiC to a population of clear cell and papillary cell renal cell carcinomas, using reference signals derived from the mature normal kidney. This population of tumors is known to resemble the PT1 population of proximal tubular cells and have a strong vascular component. The relative contribution of each signal to each bulk RNA-seq sample is shown by the y-axis. The results for the three methods are split into blocks, with cellular signals on the left, BSEQ-sc in the middle, and MuSiC on the right. Each signal type is labelled with an abbreviation and colored as shown by the legend on the right. Signals are marked with a square for fetal kidney and circle for mature kidney. For our method, there is an additional signal which represents the unexplained signal for each sample (see Methods). Contributions due to vascular signals, fibroblast signals, leucocyte signals, and all non-proximal tubular nephron signals are aggregated together. Each signal/sample combination is represented by a single point and the distribution of relative signal contributions to the bulk transcriptomes are summarized with boxplots. Arrows mark the PT1 population which these tumors are known to resemble.

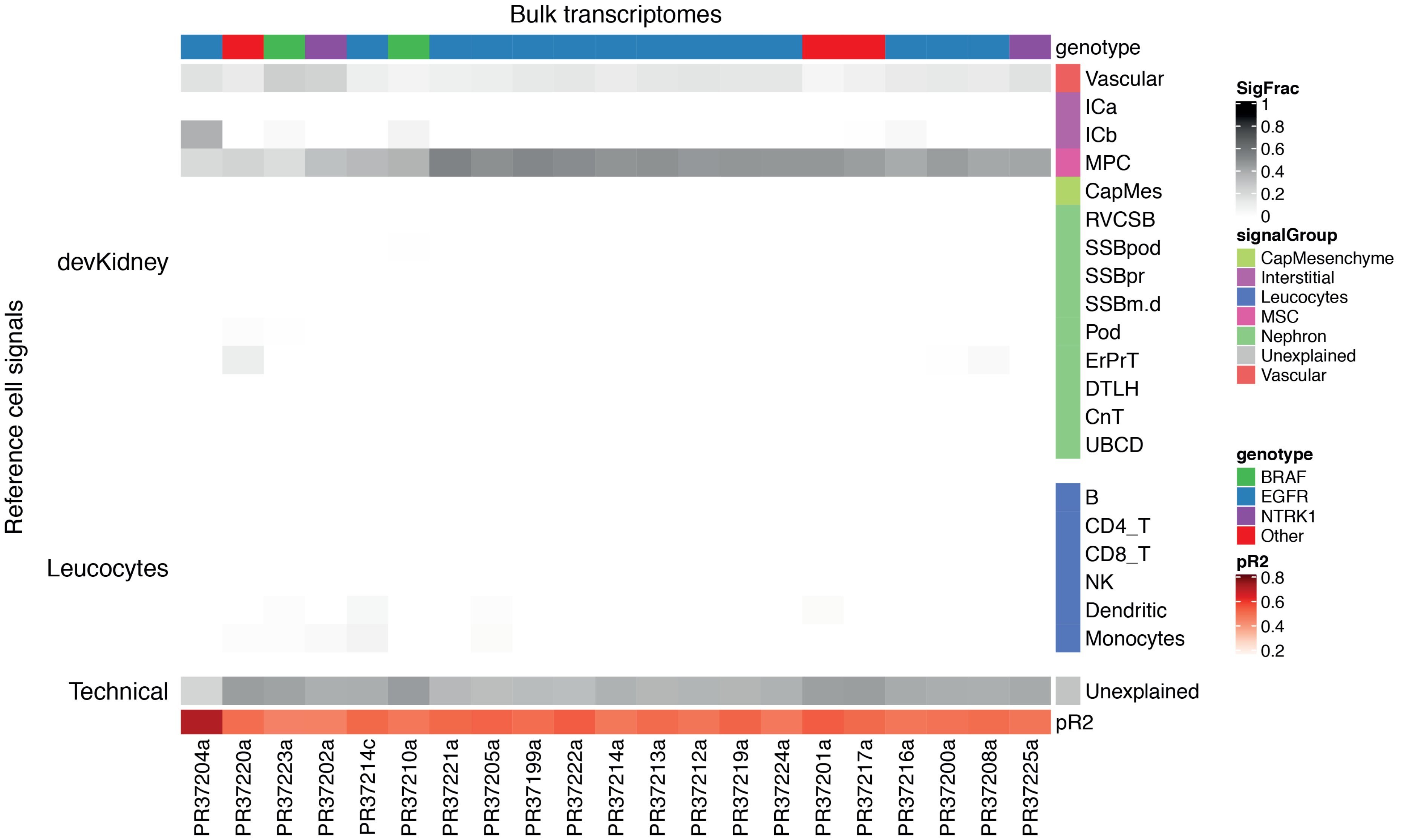

Bulk transcriptomes
genotype
Vascular
ICa
ICb
MPC
CapMes
RVCSB
SSBpod
SSBpr
SSBm.d
Pod
ErPrT
DTLH
CnT
UBCD
B
CD4_T
CD8_T
NK
Dendritic
Monocytes
Unexplained
pR2
Reference cell signals
devKidney
Leucocytes
Technical
PR37204a
PR37220a
PR37223a
PR37202a
PR37214c
PR37210a
PR37221a
PR37205a
PR37199a
PR37222a
PR37214a
PR37213a
PR37212a
PR37219a
PR37224a
PR37201a
PR37217a
PR37216a
PR37200a
PR37208a
PR37225a
SigFrac
1
0.8
0.6
0.4
0.2
0
signalGroup
CapMesenchyme
Interstitial
Leucocytes
MSC
Nephron
Unexplained
Vascular
genotype
BRAF
EGFR
NTRK1
Other
pR2
0.8
0.6
0.4
0.2

**Supplementary Figure 3 – CMNs resembles interstitial cells**

The same data as in **Fig. 3A**, but presented in heatmap form. Each column represents a sample and each row a collection of reference signal. The shading in each cell represents the relative contribution of that collection of reference signals to explaining the bulk transcriptome of that sample. The CMN samples are then further sub-divided by genotype. Sample IDs are printed below each column. The pR2 column represents a pseudo-R squared value for each sample, calculated as 1 minus the ratio of the log likelihoods of the full model to a model consisting of only the intercept term.

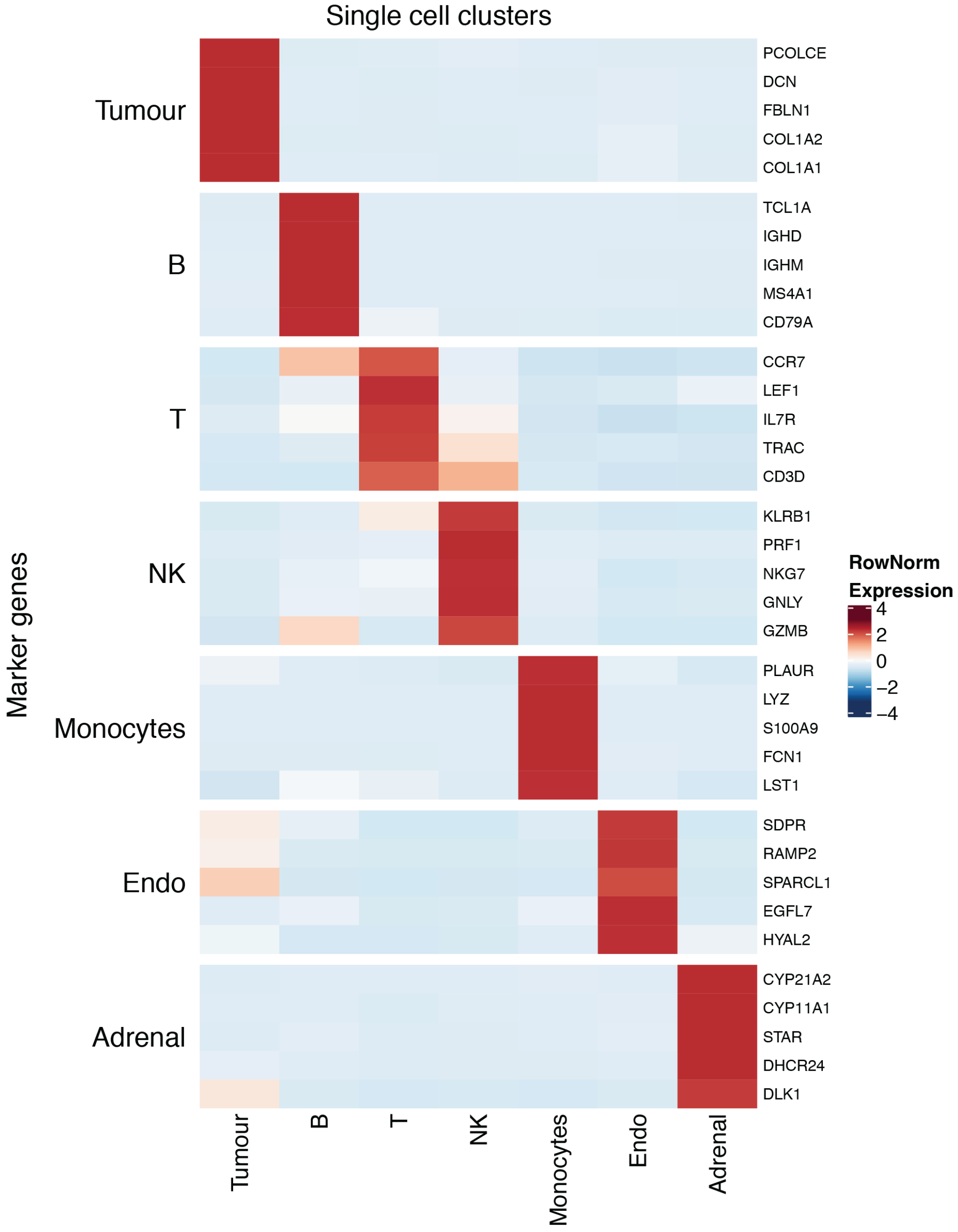


**Supplementary Figure 4 – Key markers of cell types of CMN** For each annotated cell population in **Fig. 3D**, the top 5 algorithmically determined marker genes are shown. The color scheme indicates the average normalized expression of the cells in the cluster indicated on the x-axis, z-scaled across all cell types to have mean 0 and standard deviation 1.

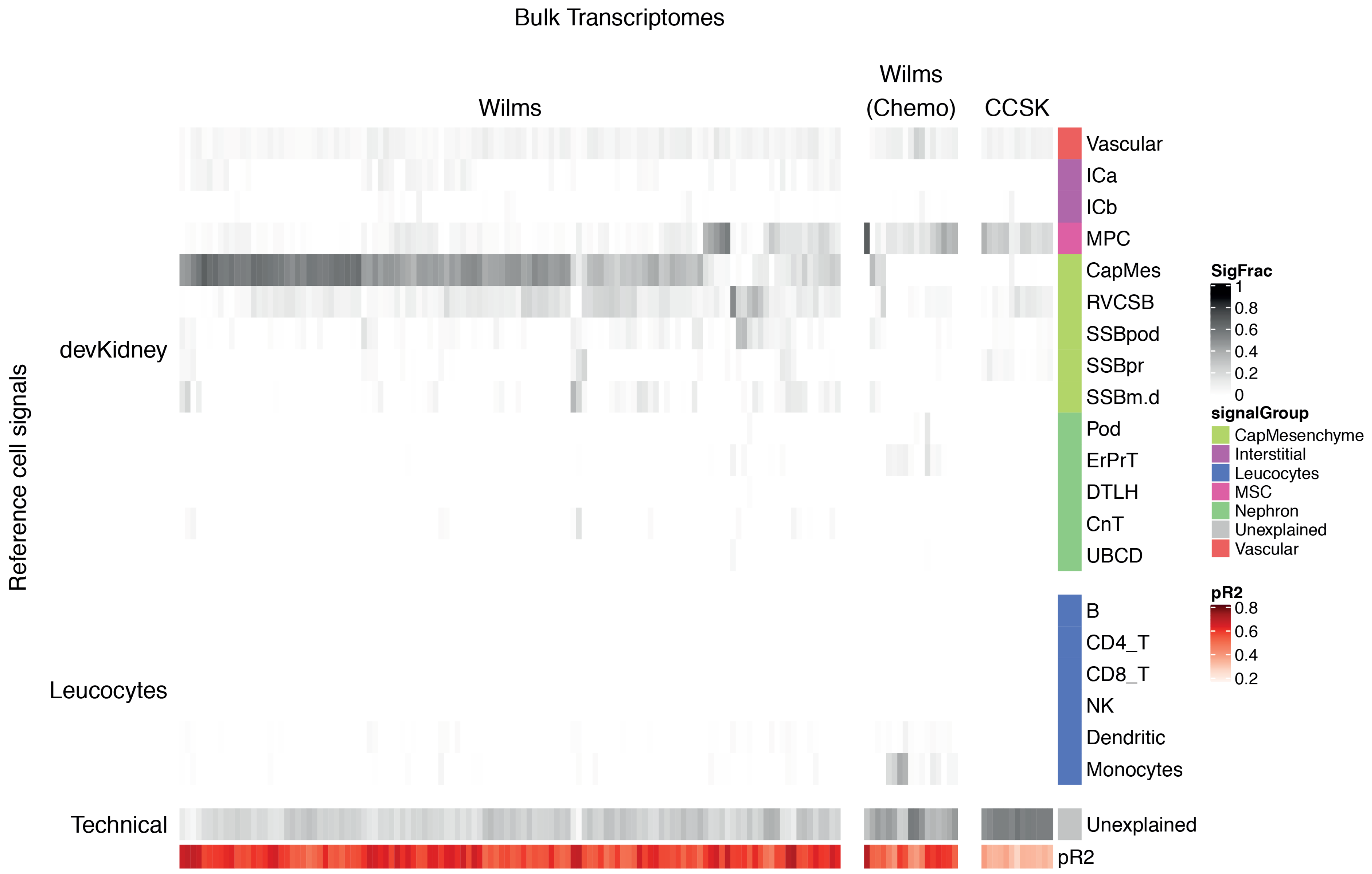

Bulk Transcriptomes
Wilms
Wilms (Chemo)
CCSK
Reference cell signals
devKidney
Leucocytes
Technical
Vascular
ICa
ICb
MPC
CapMes
RVCSB
SSBpod
SSBpr
SSBm.d
Pod
ErPrT
DTLH
CnT
UBCD
B
CD4_T
CD8_T
NK
Dendritic
Monocytes
Unexplained
pR2
SigFrac
1
0.8
0.6
0.4
0.2
0
signalGroup
CapMesenchyme
Interstitial
Leucocytes
MSC
Nephron
Unexplained
Vascular
pR2
0.8
0.6
0.4
0.2

**Supplementary Figure 5 – Wilms/Nephroblastoma and CCSK fit using fetal kidney signals**
The same data as in **Fig. 4A**, but presented in heatmap form. Each column represents a sample and each row a collection of reference signal. The shading in each cell represents the relative contribution of that collection of reference signals to explaining the bulk transcriptome of that sample. The pR2 column represents a pseudo-R squared value for each sample, calculated as 1 minus the ratio of the log likelihoods of the full model to a model consisting of only the intercept term.

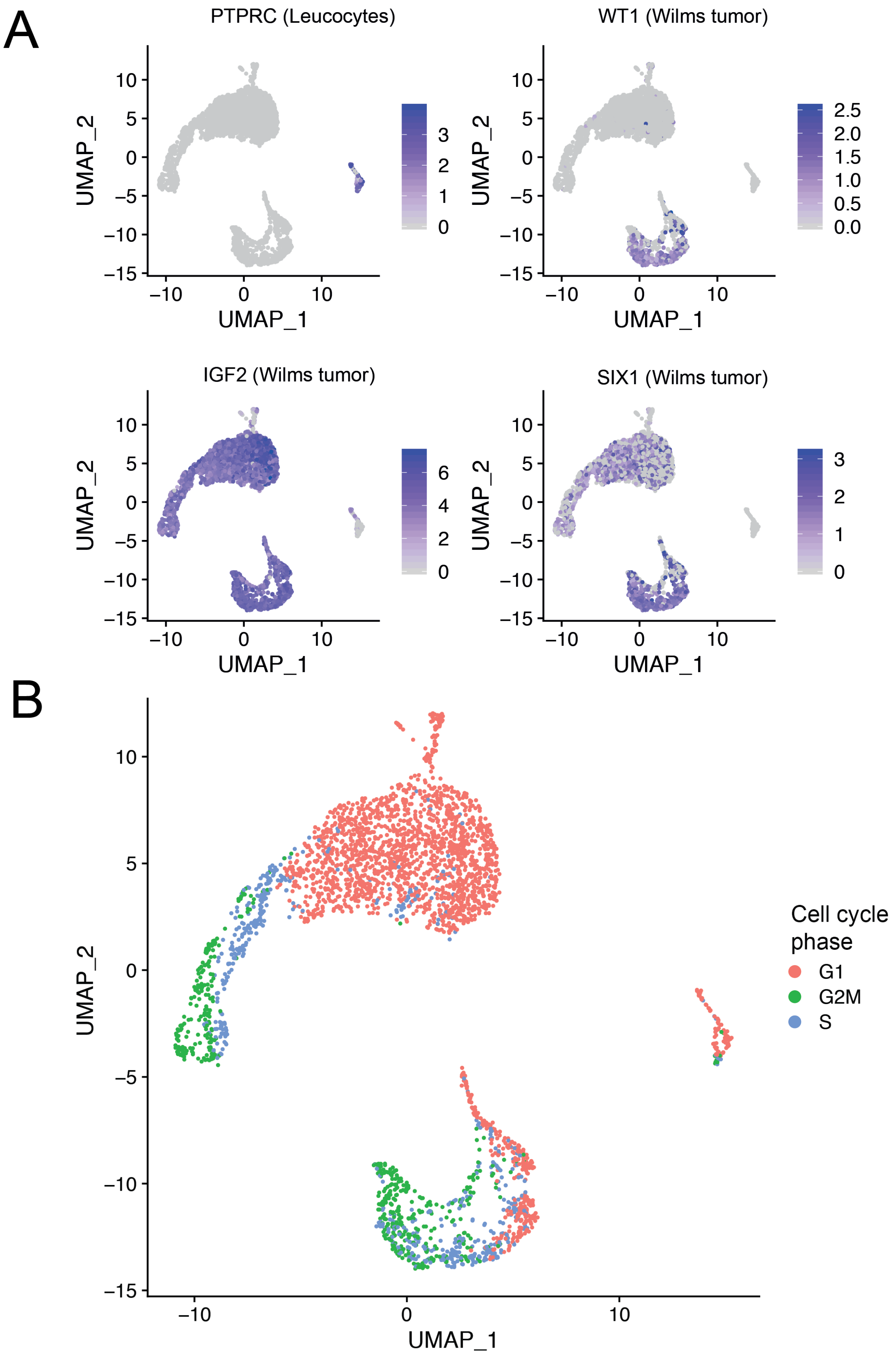

A
PTPRC (Leucocytes)
WT1 (Wilms tumor)
IGF2 (Wilms tumor)
SIX1 (Wilms tumor)
UMAP_1
UMAP_2
B
Cell cycle phase
G1
G2M
S
UMAP_1
UMAP_2

**Supplementary Figure 6 – Annotation of single cell Wilms data**

A. **Expression of key markers –** Reduced dimension representation (UMAP) of the transcriptomes of Wilms cells, colored by log-normalized expression of the gene in the panel title. The cell type that a gene is a marker of is shown in brackets.
B. **Phase of cell cycle –** The same data as panel **A**, but colored by inferred cell cycle phase as indicated by the legend on the right.

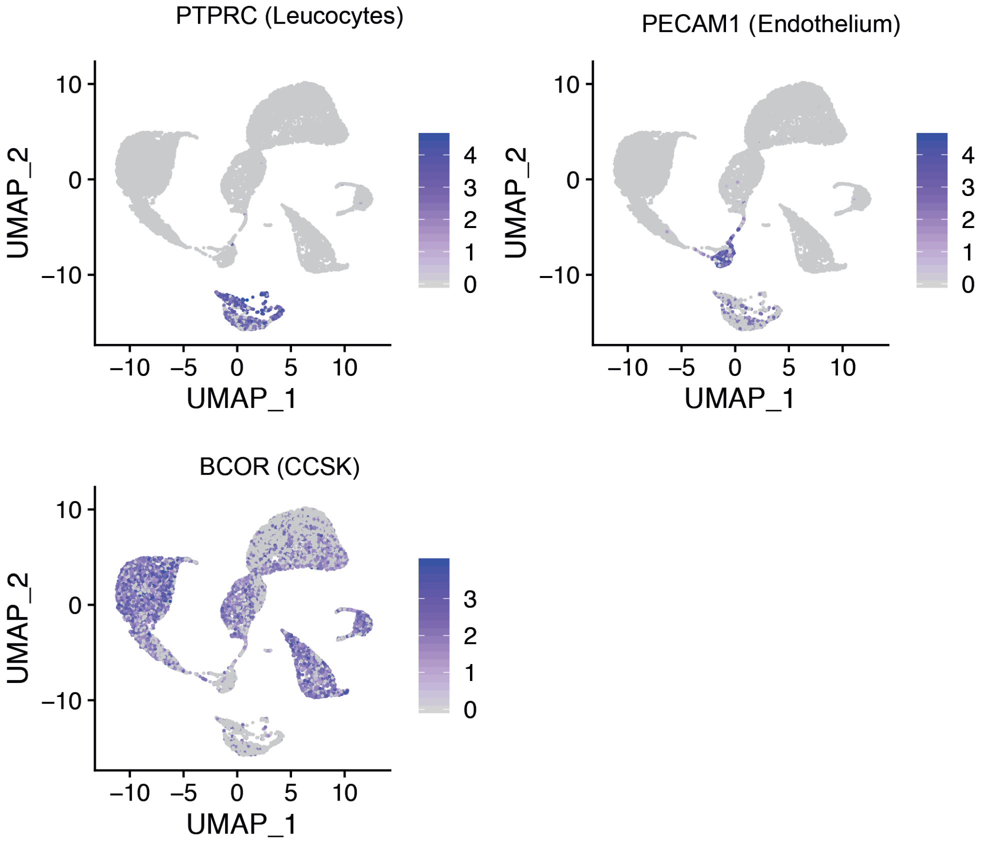


**Supplementary Figure 7 – Annotation of single cell CCSK data**

Reduced dimension representation (UMAP) of the transcriptomes of CCSK cells, colored by log-normalized expression of the gene in the panel title. The cell type that a gene is a marker of is shown in brackets.

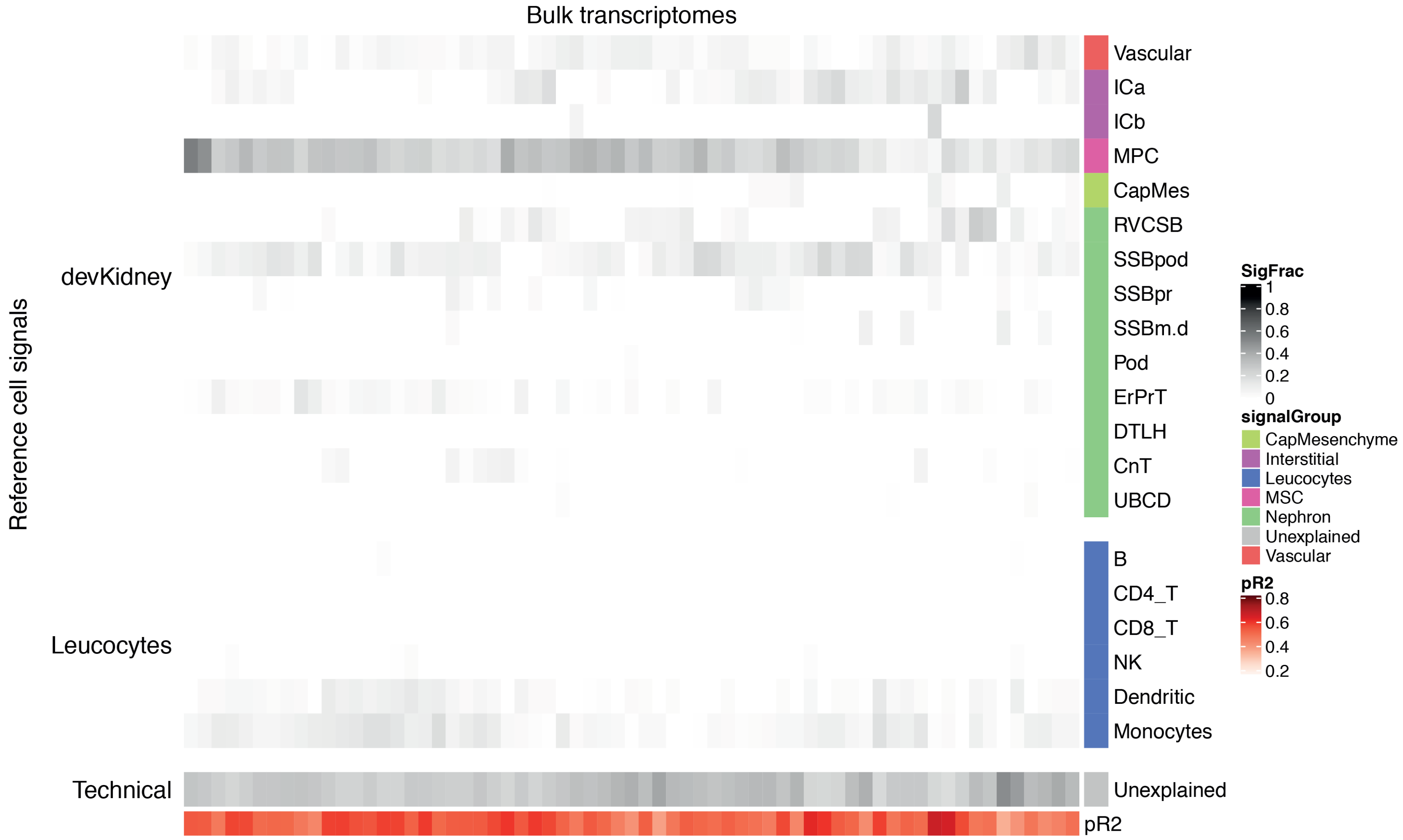
Bulk transcriptomes
Reference cell signals
devKidney
Leucocytes
Technical
Vascular
ICa
ICb
MPC
CapMes
RVCSB
SSBpod
SSBpr
SSBm.d
Pod
ErPrT
DTLH
CnT
UBCD
B
CD4_T
CD8_T
NK
Dendritic
Monocytes
Unexplained
pR2
SigFrac
1
0.8
0.6
0.4
0.2
0
signalGroup
CapMesenchyme
Interstitial
Leucocytes
MSC
Nephron
Unexplained
Vascular
pR2
0.8
0.6
0.4
0.2

**Supplementary Figure 8 – MRTs fit using fetal kidney signals**
The same data as in **Fig. 5A**, but presented in heatmap form. Each column represents a sample and each row a collection of reference signal. The shading in each cell represents the relative contribution of that collection of reference signals to explaining the bulk transcriptome of that sample. The pR2 column represents a pseudo-R squared value for each sample, calculated as 1 minus the ratio of the log likelihoods of the full model to a model consisting of only the intercept term.

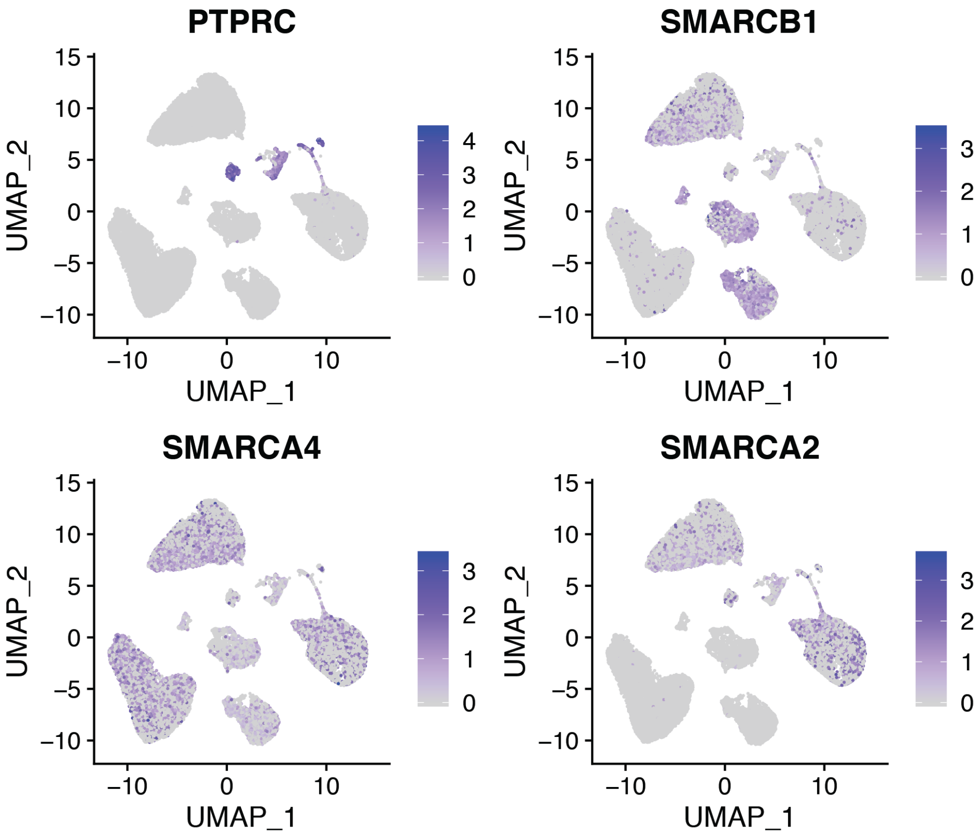


**Supplementary Figure 9 - Annotation of single cell MRT data**
Reduced dimension representation (UMAP) of the transcriptomes of MRT cells, colored by log-normalized expression of the gene in the panel title.

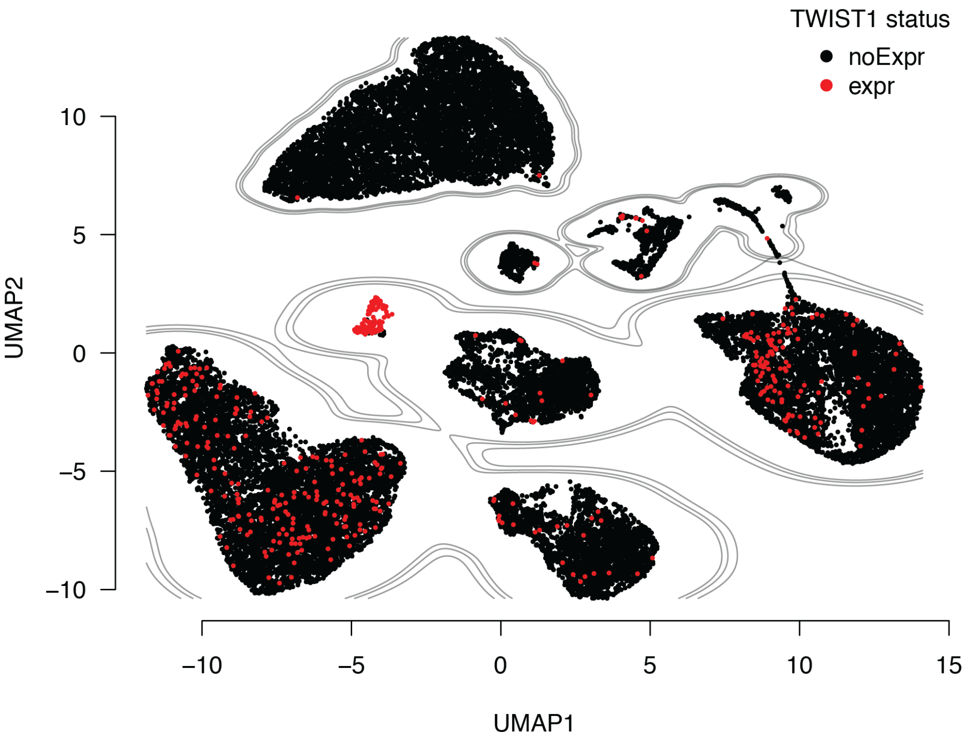


**Supplementary Figure 10 – TWIST1 positive cells in MRT**
Reduced dimension representation (UMAP) of the transcriptomes of MRT cells (as in **Fig. 5B**), with cells with any expression of *TWIST1* colored red and those without colored black.

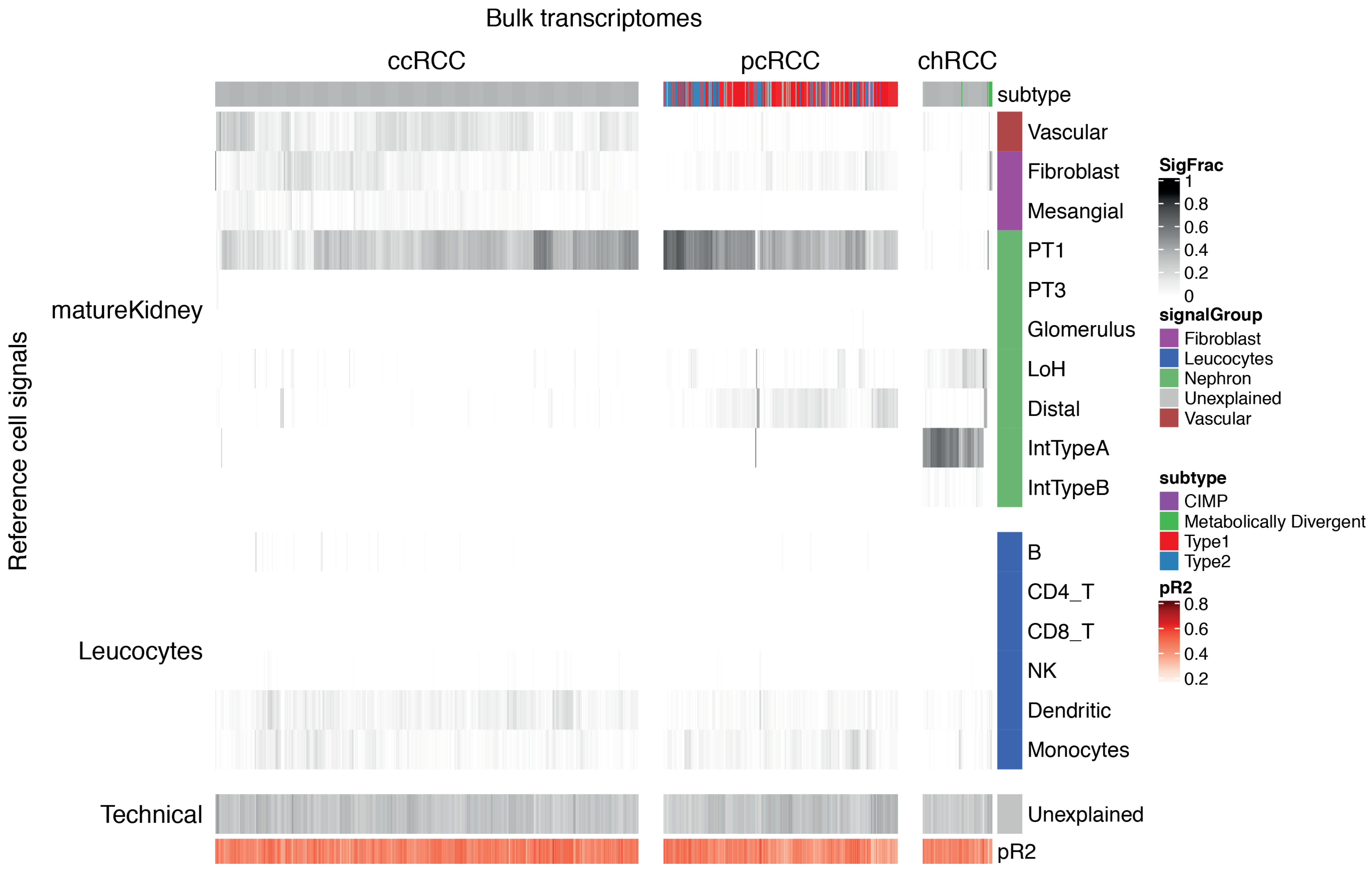
Bulk transcriptomes
ccRCC
pcRCC
chRCC
subtype
Vascular
Fibroblast
Mesangial
PT1
PT3
Glomerulus
LoH
Distal
IntTypeA
IntTypeB
B
CD4_T
CD8_T
NK
Dendritic
Monocytes
Unexplained
pR2
Reference cell signals
matureKidney
Leucocytes
Technical
SigFrac
1
0.8
0.6
0.4
0.2
0
signalGroup
Fibroblast
Leucocytes
Nephron
Unexplained
Vascular
subtype
CIMP
Metabolically Divergent
Type1
Type2
pR2
0.8
0.6
0.4
0.2

**Supplementary Figure 11 – All RCCs fit using mature kidney signals**

The same data as in **Fig. 6A**, but presented in heatmap form. Each column represents a sample and each row a collection of reference signal. Subtypes of tumor are indicated by the annotation row at the top of the figure. The shading in each cell represents the relative contribution of that collection of reference signals to explaining the bulk transcriptome of that sample. The pR2 column represents a pseudo-R squared value for each sample, calculated as 1 minus the ratio of the log likelihoods of the full model to a model consisting of only the intercept term.

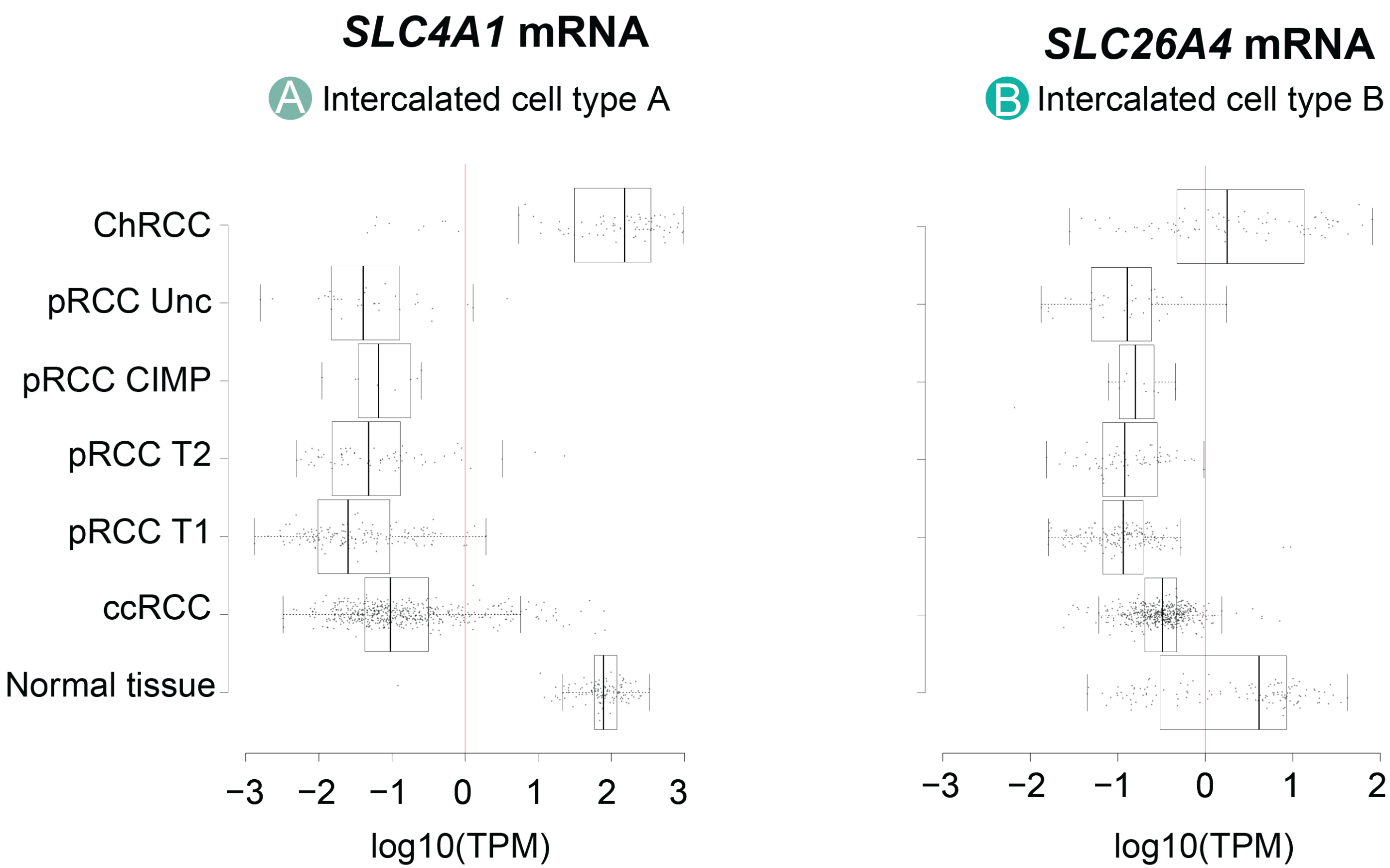


**Supplementary Figure 12 - Expression of markers of Type A and B intercalated cells in bulk transcriptomes**
Boxplots showing expression of canonical markers of Type A (left) and B (right) intercalated cells in different populations of bulk kidney tumor and normal transcriptomes. Expression values are normalized to transcripts per million (TPM) and log10 transformed and shown on the x-axis, with each point representing a sample and a boxplot showing the distribution of expression values for that sample type. A red line marks 0 on the log transformed expression scale.

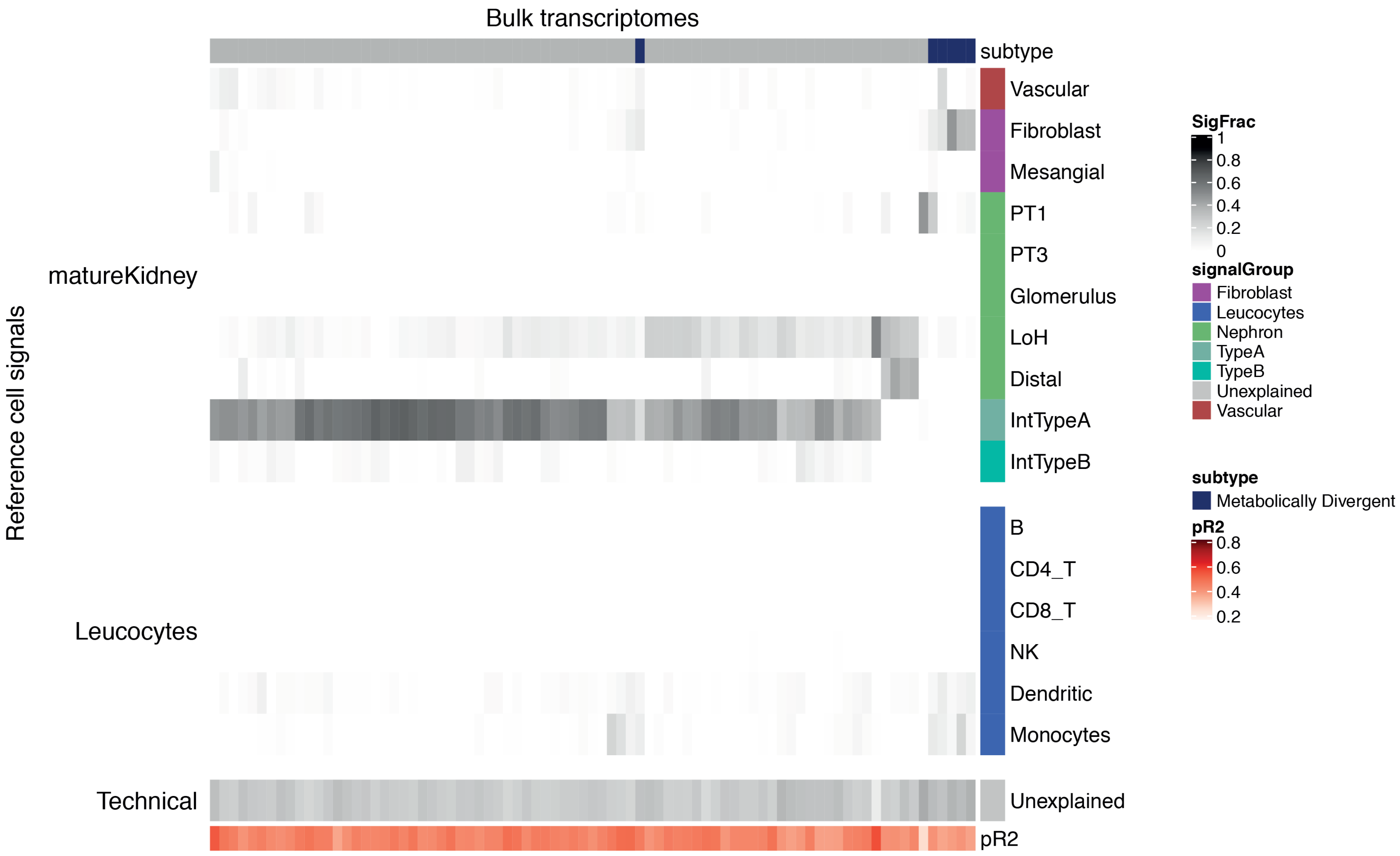
Bulk transcriptomes
subtype
Vascular
Fibroblast
Mesangial
PT1
PT3
Glomerulus
LoH
Distal
IntTypeA
IntTypeB
B
CD4_T
CD8_T
NK
Dendritic
Monocytes
Unexplained
pR2
matureKidney
Leucocytes
Technical
Reference cell signals
SigFrac
1
0.8
0.6
0.4
0.2
0
signalGroup
Fibroblast
Leucocytes
Nephron
TypeA
TypeB
Unexplained
Vascular
subtype
Metabolically Divergent
pR2
0.8
0.6
0.4
0.2

**Supplementary Figure 13 – ChRCC fit using mature kidney signals**
The same data as in **Fig. 6E**, but presented in heatmap form. Each column represents a sample and each row a collection of reference signal. Subtypes of tumor are indicated by the annotation row at the top of the figure. The shading in each cell represents the relative contribution of that collection of reference signals to explaining the bulk transcriptome of that sample. The pR2 column represents a pseudo-R squared value for each sample, calculated as 1 minus the ratio of the log likelihoods of the full model to a model consisting of only the intercept term.

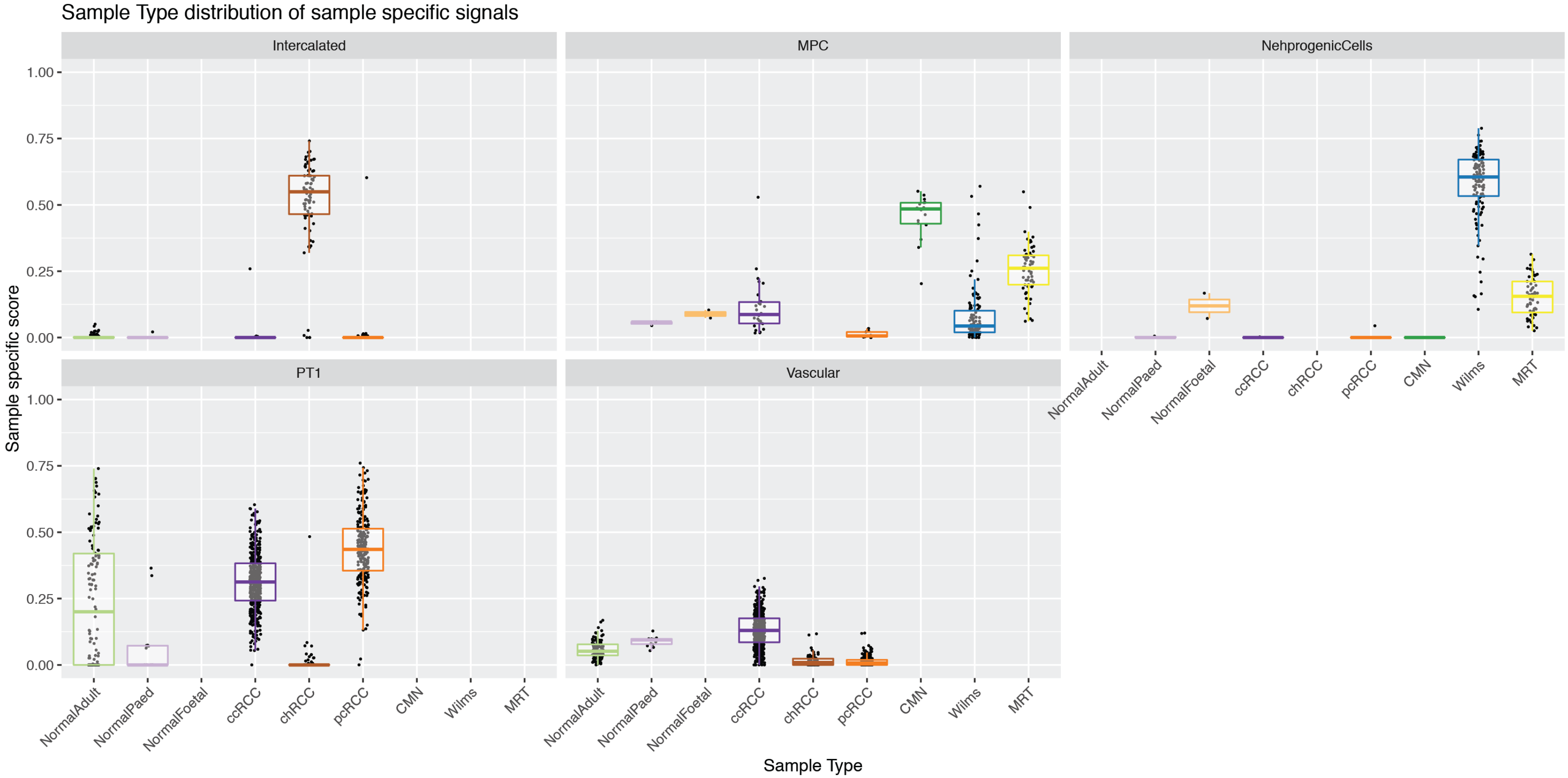


**Supplementary Figure 14 – Specificity of signal contributions to tumor types**
Each panel shows the distribution of the contribution of different "scores" calculated by aggregating the contribution from various signals, across a range of different tumor and normal samples. Each point represents a sample, with its y-axis value giving its score, with the score type given by the panel. Samples are broken down by sample/tumor type as shown on the x-axis, with each sample type given a different color. Box-plots show the distribution of the scores within each sample type. This is a more detailed version of the data shown in **Fig. 7B**.

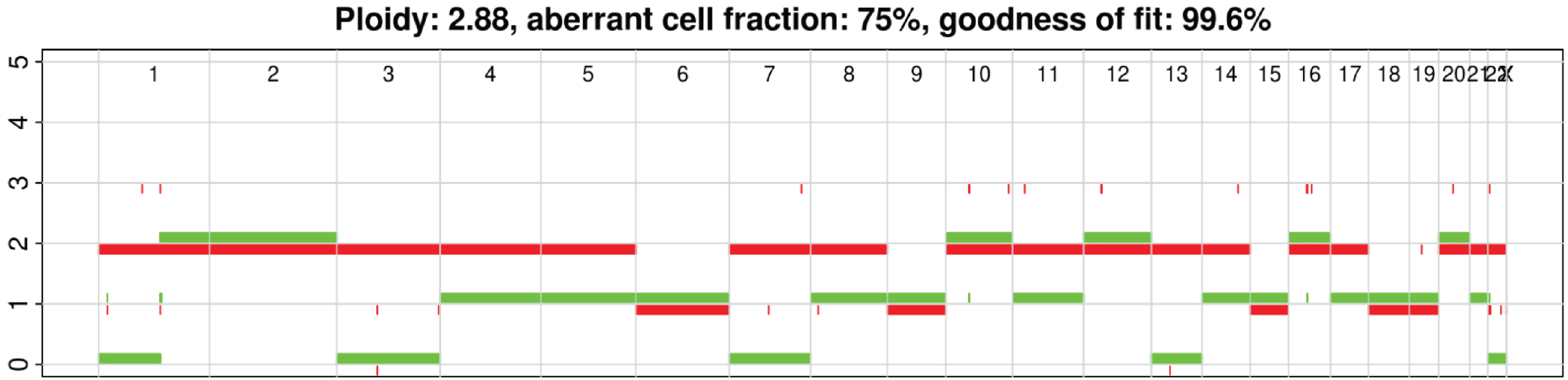


**Supplementary Figure 15 – CN profile of unknown childhood renal tumor**
Copy number profile of the unknown childhood renal tumor (SangerProject1700_PR40309a). The colored lines represent the major (red) and minor (green) alleles copy number (y-axis) along the chromosomes as shown at the top of the plot.

### Colour scheme for cellular signals

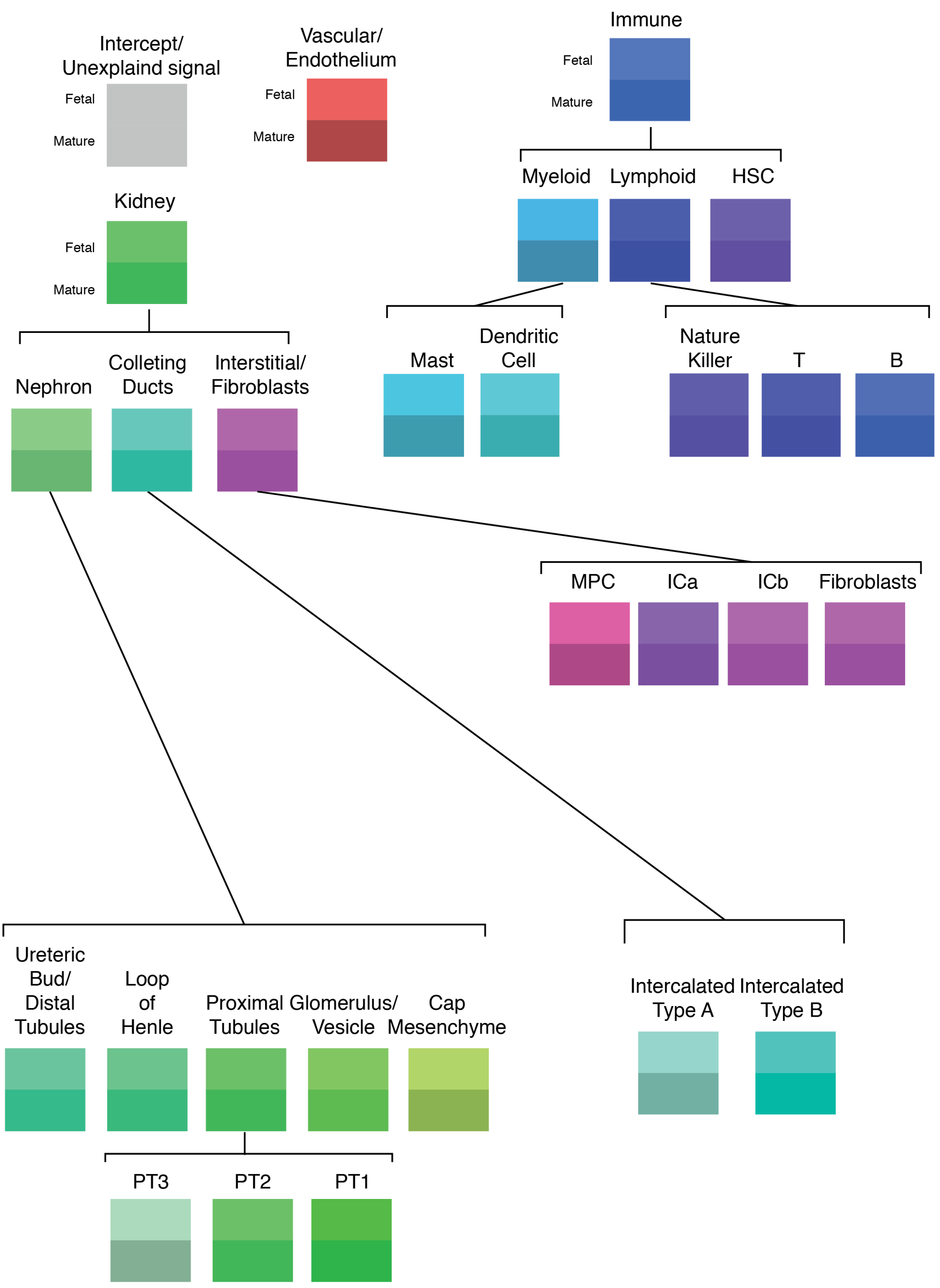

**Supplementary Figure 16 – Color scheme for cell types**
Overview of the color scheme used throughout the figures in this paper to represent different cell types. Cell types are organized hierarchically, with fetal/mature versions of the same cell type represented by colors of the same hue and saturation but different value. The exception to this rule is the unexplained signal color which is the same in both a fetal and mature context.

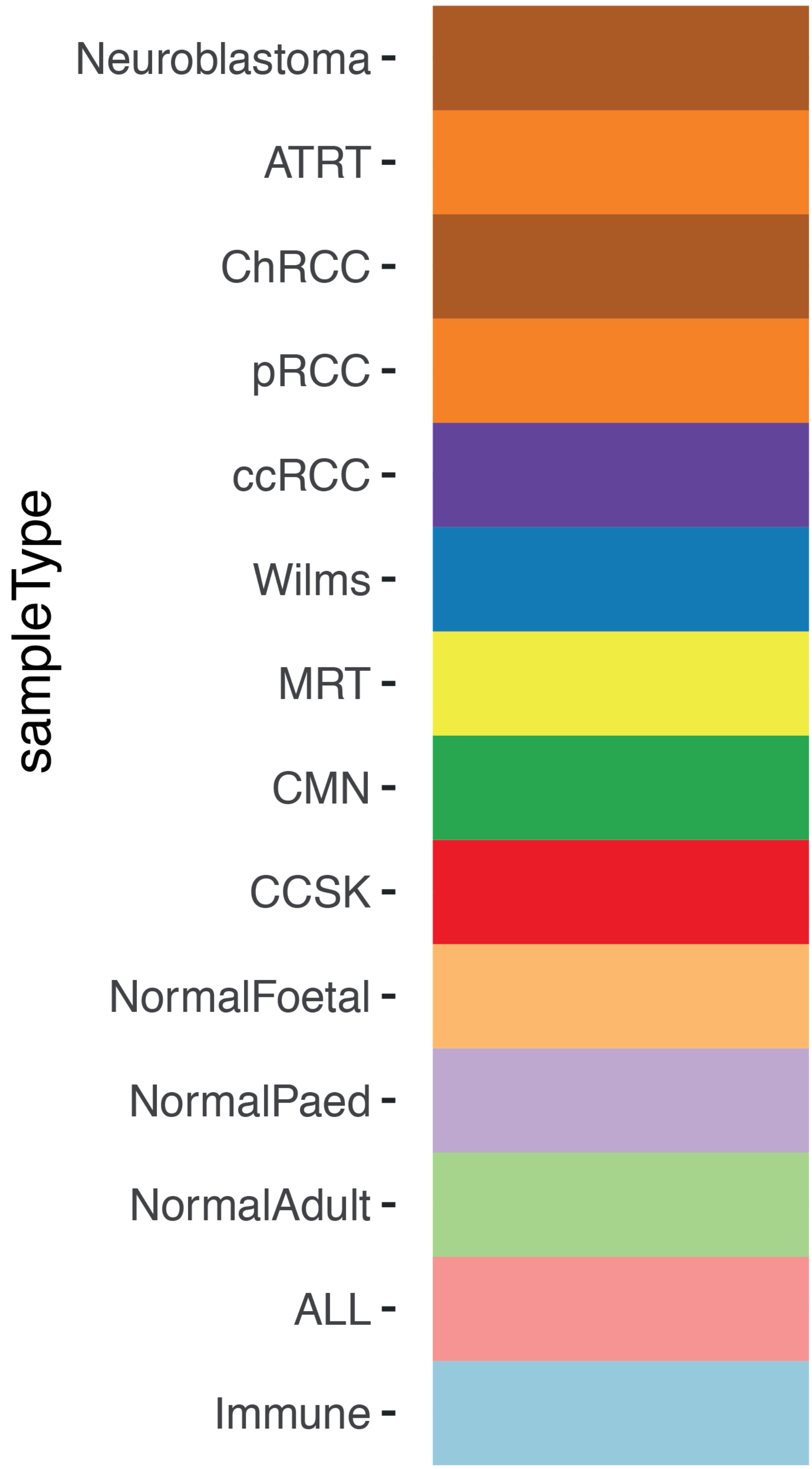


**Supplementary Figure 17 – Color scheme for sample types**
Overview of the color scheme used throughout the figures in this paper to represent different sample types. Where colors have been duplicated they either never occur in the same context, or it is obvious from the context which color refers to which sample type.

**Supplementary Table 1 – Bulk RNA manifest**
A table giving a description of each bulk RNA transcriptome used in this study as well as the path to the file containing the table of counts for that file (which can be extracted from **Data S1**).

**Supplementary Table 2 – Single cell manifest**
A table giving a description of each single cell experiment included in this study. For each row, the raw count data is available in **Data S2** in a folder with name given by the "channelLabel" column.

**Supplementary Table 3 – Linear model of immaturity by genotype**
Generated using "summary(fit)" in R, where fit is a linear model to predict the immaturity score of RCCs using sample genotype as covariates. Covariates are coded such that the intercept term corresponds to the wild type of non-tumor biopsy.

**Supplementary Table 4 – Linear model of immaturity by other features**
Generated using "summary(fit)" in R, where fit is a linear model to predict the immaturity score of ccRCCs using sample genotype as covariates. Covariates are coded such that the intercept term corresponds to unknown mRNA group, unknown miRNA group, grade G1, and stage I.

**Supplementary Table 5 – Mast cell fraction for RCCs**
Quantification of mast cell prevalence from smFISH of different tumors. In the column "tType" pRCC1/2 represents papillary cell renal cell carcinoma type 1/2.

**Supplementary Data 1 – Bulk transcriptomes**
Text files giving fragment counts and gene lengths for every sample in this study, with each sample's count table stored in a separate file.

**Supplementary Data 2 – Single cell transcriptomes**
Tables of counts and QC metrics for every single cell dataset generated in this study. The count tables are stored in h5 format, stored one folder per channel.

**Supplementary Data 3 – Data to generate figures**
The data files from which the figures and supplementary figures contained in this paper were generated. Each figure has a corresponding text file in this data object containing the matching data table.

**Supplementary Data 4 – Source code**
Collection of scripts used to perform the analysis and generate this manuscript. These scripts are provided to give additional details as to how we implemented the analyses described in the **Methods** section.
