## Supplementary material for "Single cell derived mRNA signals across human kidney tumors": Source Code: cellSignalAnalysis.py

```
'''
Takes a reference single cell data set and attempts to explain bulk samples as a linear combination of the population present in the single cell data.

*Installation*

This script relies on various packages and is written for python 3.  If you install tensorflow, pandas, numpy and scipy that should be enough.

*Documentation*

Read the documentation by running "python <this_file> -h".  Some specific notes:
    - The ideal way to provide bulk data is to have one sample per file.  This seems cumbersome but makes things much faster when you have a TCGA size data set and only want to load 10 samples.  If you just have 10 samples in one giant TCGA sized matrix, the whole file must be loaded into memory, then the samples you want selected.  This requires lots of memory and slows things done immensely.  Each individual file should look like this:
        GeneName    GeneLength  SampleID
        CD13    1210    3
        CD4 4320    0
        EPCAM1  399 2
        ...
    That is, it should be tab delimited, the first row must give column headers, the first column must give the names of genes and the second column must give the gene length.
    - If gene names differ between files (e.g. different bulk sample files are created using different annotations), only the global intersection of gene names is considered.  That is, any information pertaining to a gene not present in all samples and the reference is not considered and thrown out.  So take care not to use wildly discrepant annotations.
    - The output consists of:
        + *_usedBulkCounts.tsv - The exact table of fragment counts used to do the fit.
        + *_usedBulkGeneLengths.tsv - The exact table of per-sample gene lengths used to do the fit.
        + *_usedCellularSignals.tsv - The definition of the cellular signals used to perform the fit.
        + *_usedGeneWeights.tsv - The exact gene weights used to perform the fit.
        + *_fitExposures.tsv - The main results file.  Fitted values for the relative contribution of the cellular signals and intercept term, for the samples provided.  There are also some goodness of fit metrics included in this table.  pR2, which is a pseudo-R-squared value, obsCount, which is the total number of fragment counts observed for each sample, and fitCount, which is the total fitted fragment count for each sample.
        + *_fitCounts.tsv - The full fitted table of counts.
        + *_negativeLogLikelihoodFullFit.tsv - This table gives the contribution to the total negative log-likelihood for each gene in each sample.  This is most useful for assesing which genes are most poorly explained by the model fit.  This log-likelihood is for the full model fit consisting of all celullalar signals and an intercept term.
        + *_negativeLogLikelihoodNullFit.tsv - The same as above, but for the Null model where only an intercept term is used.


*Test run*
ipython -i -- ./cellSignalAnalysis.py \
        -b ./bulkData/SangerProjectTARGET_ALL_phase1.txt \
        -s ./scData/PBMCRef \
        -w ./geneWeights.tsv \
        ./output/TestRun

'''

#############
# Libraries #
#############
import os
import datetime
import argparse
#Suppress lots of memory warning
os.environ['TF_CPP_MIN_LOG_LEVEL'] = '3' 
from os.path import expanduser as eu
from os import path
from math import ceil
import tensorflow as tf
import pandas as pd
import numpy as np
from scipy.stats import poisson 
from scipy.special import loggamma
from scipy.io import mmread
import scipy.sparse as sparse
from pandas.core.algorithms import match
import warnings
tf.compat.v1.disable_eager_execution() 

##################################
# Define command line parameters #
##################################
parser = argparse.ArgumentParser()
parser.add_argument('output',type=str,help='Path where output is saved.  Files are created by adding a suffix to this argument for each output file.  For example, results/RCC would produce results/RCC_fit.tsv, results/RCC_sigs.tsv, etc.')
parser.add_argument('-b','--bulk',type=str,nargs='+',help='Where to get the bulk data from.  Bulk data must be provided as a tab separated text file with rows indicating genes and columns indicating samples.  The first column must give gene names, the second column the effective length of each gene and the first row sample names.  In the case of multiple samples, you can provide multiple arguments or a text file with the names of the files containing the samples you want, one per line.  If multiple samples are in the same file, the gene length is assumed to be the same across all samples.',required=True)
parser.add_argument('-s','--sigs',type=str,nargs='+',help='''Where to get the "cellular signals" that the bulk data will be decomposed into.  This must either be a single cell data-set with annotation indicating how to combine data, or a matrix giving the already summarised cellular signals.  
        Single cell data must be provided as a sparse matrix in Matrix Mart format.  In this case there are 3 files required, the actual counts, the row labels (gene-names), and the column labels (cell annotations).  These are identified by adding the suffixes .mtx, _rowNames.tsv, and _columnNames.tsv respectively to the provided argument.  Annotation is infered from the column labels by taking everything between the start of the string and the first colon as the cell annotation.  E.g., the cell with label "T:ACGTATTTACGA-1___STDY3411" would be annotated as "T".  The row and column name files should be one line per row/column with no headers giving the label in the matching matrix mart count file.
        If instead a summary file is given, the format must be the same as with bulk data.  The only difference is that the "geneLength" column is no longer needed and columns are now cellular signals representing populations of cells from single cell data and must be normalised to sum to 1 across each column.
        Regardless of wether single cell data or a summary file is provided, multiple sources can be provided, either through providing multiple arguments to this command, or providing the path to a file containing one source per line.''',required=True)
parser.add_argument('-rs','--refine-sigs',type=str,nargs='+',help='Either a file containing the IDs of cellular signals to include or each cellular signal to include passed as an argument.  The intersection of this list and all those cellular signals specified by -s are used.  If not specified, all signals given by -s are used.',default=None)
parser.add_argument('-w','--weights',type=str,help='Weights for each gene.  If omitted, all set to 1.  Must be a two column tab delimited file with no headers, the first column giving gene names, the second the weight to apply for that gene in the model, a value between 0 and 1.',default=None)
parser.add_argument('--no-collapse-ref',action='store_true',help='Usually signatures are created by collapsing cells within a cluster to create a cell specific signal.  This flag will instead use every cell as its own signal.')
parser.add_argument('--l1-lambda',type=float,help="Regularisation lambda parameter for L1 norm.  Controls how much regularisation to apply, with larger values providing a stronger penalty and more regularisation.  Default does no regularisation.  The scale of lambda is set such that when lambda=1 the negative log-likelihood of the NULL model is approximately equal to the penalty, assuming that the sum of the exposures equals the sum of the molecules in bulk (where number of molecules is estimated using the reads/length*insert_size) and that exposures are uniformly distributed.  So a lamda of 0.1 would apply essentially no constraint until the likelihood became comparable with that of the NULL model, with it becoming comparable in magnitude around the point where the likelihood of the model has improved over the NULL by 0.1.  Includes the intercept.",default=0)
parser.add_argument('--l2-lambda',type=float,help="Regularisation lambda parameter for L2 norm.  Everything applies as with L1 lambda, except this penalty does NOT include the intercept.",default=0)
parser.add_argument('-i','--insert-size',type=int,help="To convert between the cellular signals, which give the relative abundance of molecules, and the bulk counts, which give counts of fragments mapped to genes, a conversion factor is needed.  This is taken care of by the gene length supplied along with the bulk data, up to a multipliciative constant.  That is, if the model predicts N molecules of gene X, which has length L, the number of bulk fragment counts predicted is X*L/iSize where iSize is the insert-size of a fragment (the value specified here).  As this value is the same for all genes, getting it wrong will only change the scale of the raw fitted co-efficients.  As these values are basically useless anyway, setting this parameter 'inaccurately' changes nothing of importance.  So don't obsess about it being set 'correctly'.",default=500)
parser.add_argument('-dg','--drop-zero-sig-genes',action='store_true',help='Drop any gene that is not expressed in any cellular signals from the fit.  With a zero intercept, these genes would add an infinite contribution to the log-likelihood.  So the main way this changes is the fit is via the goodness-of-fit metric and possibly the size of the required intercept term if there are many such genes.')
parser.add_argument('-ie','--init-log-exposure',type=float,help='Initial value of log-exposure to use when training model.  A moderately negative value helps speed up the fit.  If set too far from 0 the fit will fail to find a sensible solution.',default=-10)
parser.add_argument('-r','--learn-rate',type=float,help='The learn rate to use.',default=0.01)
parser.add_argument('-p','--poll-interval',type=int,help='Poll fit after this many iterations',default=100)
parser.add_argument('--max-it',type=int,help='Maximum number of iterations',default=1e7)
parser.add_argument('-tl','--log-likelihood-tolerance',type=float,help='Termination condition.  Stop when improvement in the log-likelihood less than this.',default=1e-6)
parser.add_argument('-ts','--sparsity-tolerance',type=float,help='Termination condition.  Stop when improvement in the sparsity is less than this.',default=1e-4)
args = parser.parse_args()

###################
# Save parameters #
###################
print("Launching deconvolution with arguments:")
print(args)
tmp = vars(args)
args.output = eu(args.output)
with open(args.output+'_arguements.tsv','w') as f:
  x = f.write('Parameter\tValue\n')
  x = f.write('Launched\t'+str(datetime.datetime.now())+'\n')
  x = f.write('WorkDir\t'+os.getcwd()+'\n')
  for k in tmp.keys():
    x = f.write(k+'\t')
    x = f.write(str(tmp[k])+'\n')


##################
# Load bulk data #
##################
#Determine list of files to load
#Go through them one at a time and either parse the file list or store the path
bulkSrcs = []
bulkPointers = [eu(x) for x in args.bulk]
for bulkPointer in bulkPointers:
  #Peak inside file
  with open(bulkPointer,'r') as f:
    dat = f.readline().strip('\n').split('\t')
    #Is it a pointing file?
    if len(dat)==1:
      #This is a one-argument per line file.  Should point to other files.
      f.seek(0)
      #Paths in file are relative to the pointing file.
      bulkSrcs = bulkSrcs + [os.path.join(os.path.dirname(bulkPointer),eu(x.strip('\n'))) for x in f.readlines()]
    else:
      #It is the file itself
      bulkSrcs.append(bulkPointer)
#Now load the files containing data one at a time
toc = []
for bulkSrc in bulkSrcs:
  toc.append(pd.read_csv(bulkSrc,sep='\t',index_col=0))
#Do a rough check that the input files are sensible
if min([x.shape[1] for x in toc])<2:
  raise ValueError("Some source files have fewer than 3 columns.  Each source file must contain at least three columns: geneName,geneLength, and data for one or more samples.")
#Get the common gene names
bulkGenes = list(set.intersection(*[set(x.index.values) for x in toc]))
bulkGenes.sort()
nGenes = [x.shape[0] for x in toc]
if len(bulkGenes) < max(nGenes):
  warnings.warn("Bulk samples have between %d and %d genes, with only %d in common.  All non-shared genes will be dropped."%(min(nGenes),max(nGenes),len(bulkGenes)))
#Create a length matrix from this
tol = []
for x in toc:
  #Create expanded length matrix
  tmp = x.iloc[:,[0]*(x.shape[1]-1)]
  tmp.columns = x.columns[1:]
  tol.append(tmp)
tol = pd.concat([x.loc[bulkGenes,] for x in tol],axis=1)
#And one with the bulk data
toc = pd.concat([x.drop(columns=x.columns[0]).loc[bulkGenes,] for x in toc],axis=1)

#######################
# Load reference data #
#######################
#Get a list of sources
#Go through them one at a time and either parse the file list or store the path
refPointers = [eu(x) for x in args.sigs]
refSrcs=[]
for refPointer in refPointers:
  try:
    with open(refPointer,'r') as f:
      dat = f.readline().strip('\n').split('\t')
      #Is it a pointer?
      if len(dat)==1:
        #This is a one-argument per line file.
        f.seek(0)
        refSrcs = refSrcs + [os.path.join(os.path.dirname(refPointer),eu(x.strip('\n'))) for x in f.readlines()]
      else:
        refSrcs.append(refPointer)
  except:
    #If we're here, it's a single cell data-set so should be stored as is too
    refSrcs.append(refPointer)
#Now load the files one at a time
scSigs = []
for refSrc in refSrcs:
  #Try loading it as single cell data, if not assume it's a summary file.
  try:
    with open(refSrc+'.mtx','rb') as mm:
      with open(refSrc+'_rowNames.tsv','r') as rn:
        with open(refSrc+'_columnNames.tsv','r') as cn:
          print("Loading and processing single cell data from %s"%refSrc)
          #Load and create summary stats
          dat = mmread(mm).tocsc()
          #Get the row/col names
          rowNames = [x.strip('\n') for x in rn.readlines()]
          colNames = [x.strip('\n') for x in cn.readlines()]
          #Extract grouping annotation from labels
          if args.no_collapse_ref:
            cellSigs = np.array([x for x in colNames])
          else:
            cellSigs = np.array([x if x.find(':')==-1 else x[:x.find(':')] for x in colNames])
          #Create a summary object for them.
          tmp = []
          for cellSig in list(set(cellSigs)):
            idxs = np.where(cellSigs==cellSig)[0]
            tgts = dat[:,idxs]
            #Collapse across cells
            tgts = tgts.sum(axis=1)
            #Normalise to sum to 1 and store
            tmp.append(tgts/tgts.sum())
          #Now reformat and add the relevant labels
          tmp = pd.DataFrame([np.squeeze(np.asarray(x)) for x in tmp],index=list(set(cellSigs)),columns=rowNames).transpose()
          scSigs.append(tmp)
  except:
    #Assume it's already a summary file, just need to load it
    tmp = pd.read_csv(refSrc,sep='\t',index_col=0)
    scSigs.append(tmp)
#Now need to concatenate and get common genes
refGenes = list(set.intersection(*[set(x.index.values) for x in scSigs]))
refGenes.sort()
nGenes = [x.shape[0] for x in scSigs]
if len(refGenes) < max(nGenes):
  warnings.warn("Reference cellular signals have between %d and %d genes, with only %d in common.  All non-shared genes will be dropped."%(min(nGenes),max(nGenes),len(refGenes)))
#Merge into one matrix 
scSigs = [x.loc[refGenes] for x in scSigs]
scSigs = pd.concat(scSigs,axis=1)
#Drop unwanted signature is there are any
if args.refine_sigs is not None:
  #Load the signals we do want, assume it's a file first
  try:
    with open(eu(args.refine_sigs),'r') as f:
      tgts = [x.strip('\n') for x in f.readlines()]
  except:
    #If it's not a file, it's a raw list of ones to include
    tgts = args.refine_sigs
  commonSigs = list(set(scSigs.columns.values).intersection(tgts))
  if length(commonSigs)==0:
    raise ValueError("No common signals found.")
  scSigs = scSigs.loc[:,commonSigs]

################
# Load weights #
################
if args.weights is not None:
  geneWeights = pd.read_csv(eu(args.weights),sep='\t',index_col='geneID')
else:
  geneWeights = pd.DataFrame(np.ones((toc.shape[0],1)),index=toc.index,columns=['weight'])


##################
# Harmonise data #
##################
commonGenes = list(set(bulkGenes).intersection(refGenes))
if len(commonGenes) < max(len(bulkGenes),len(refGenes)):
  warnings.warn("Bulk data has %d genes, reference data %d, with only %d in common.  All information relating to non-shared genes will be dropped."%(len(bulkGenes),len(refGenes),len(commonGenes)))
#Do the more stringent thing of dropping anything zero in sigs
if args.drop_zero_sig_genes:
  print("Dropping %d genes with zero expression in any of the supplied cellular signals. %d genes remain"%(sum(scSigs.loc[commonGenes].sum(axis=1)==0),sum(scSigs.loc[commonGenes].sum(axis=1)>0)))
  commonGenes = list(np.array(commonGenes)[(scSigs.loc[commonGenes].sum(axis=1))>0])
#Drop the things we're not going to use
toc = toc.loc[commonGenes]
tol = tol.loc[commonGenes]
scSigs = scSigs.loc[commonGenes]
#Finalise weights
geneWeights = geneWeights.reindex(commonGenes)
geneWeights = geneWeights.fillna(1.0)
#Re-normalise signals?
#scSigs = scSigs/scSigs.sum(axis=0)
#Some useful messages
print("Fitting %d bulk samples to %d cellular signals using %d genes"%(toc.shape[1],scSigs.shape[1],toc.shape[0]))
print("Signals named:")
tmp = scSigs.columns.values
print(str(tmp))
print("Regularising with lambda %g/%g for L1/L2 "%(args.l1_lambda,args.l2_lambda))

####################
# Define the model #
####################
p,n = toc.shape
s = scSigs.shape[1]
#########
# Inputs
S = tf.constant(scSigs.values.astype('float32'),name='fixed_signals')
#Note the insert-size conversion factor to convert length of gene to expected number of fragments per molecule of mRNA from gene
C = tf.constant((tol.values/args.insert_size).astype('float32'),name='mols_to_reads')
k = tf.constant(toc.values.astype('float32'),name='obs_counts')
w = tf.constant(geneWeights.weight.values.astype('float32'),name='gene_weights')
#Create "Signal" that is really just the intercept term
Si = tf.constant((np.ones((p,1))/p).astype('float32'),name='intercept_signal')
Sp = tf.concat([S,Si],axis=1)
############
# Exposures
#These are the things we actually train
#Initialise to -10000, which converts to 0.  This value will not be changed when fitting null
z = tf.Variable(tf.zeros([s,n])+args.init_log_exposure,name='exposures')
#Define dynamic intercept
int0 = tf.Variable(tf.zeros([1,n]),name='intercept')
#Merge
zz = tf.concat([z,int0],name='exposures_with_int',axis=0)
#Positive exposures, including intercept
E = tf.exp(zz,name='positive_exposures')
###################
# Predicted Counts
#Predicted number of molecules
q = tf.matmul(Sp,E,name='pred_mols')
#Convert to number of reads
y = tf.multiply(q,C,name='pred_reads')
##########################
# Poisson log-likelihood
#Variable part of Poisson log-likelihood
Dij = k*tf.math.log(y)- y
#Constant part
D0 = tf.math.lgamma(k+1)
#Add gene weights and sum to get negative log-likelihood
LL = tf.transpose(a=D0-Dij)*w
tLL = tf.reduce_sum(input_tensor=LL,name='NLL')
#############################
# Define objective function
#Count of non-zeroish co-efficients
cCnt = tf.reduce_sum(input_tensor=tf.sigmoid(zz))
# The final penalised, adjust NLL
O = tLL
############
# Optimiser
opt_E = tf.compat.v1.train.AdamOptimizer(args.learn_rate)


############
# Fit NULL #
############
print('''
######################
# Fitting null Model #
######################
''')
#Fit null first so we can inform the regularisation
update_op = z.assign(np.zeros([s,n])-1000)
learners = opt_E.minimize(O,var_list=[int0],name='learn_exposures')
#Initialise
nullSess = tf.compat.v1.Session()
init = tf.compat.v1.global_variables_initializer()
nullSess.run(init)
_ = nullSess.run(update_op)
#Record initial values
last = nullSess.run(O)
lastNonZero = nullSess.run(cCnt)
#Record the movements
null_nll = np.zeros(int(ceil(args.max_it/args.poll_interval)))
null_nsigs = np.zeros(int(ceil(args.max_it/args.poll_interval)))
i=0
while True:
  #Take the exposure step
  nullSess.run([learners])
  #Every now and then, record our progress and check if we've converged
  if i%args.poll_interval == 0:
    ii = i//args.poll_interval
    #Record object function and number of non-zero exposures
    null_nll[ii] = nullSess.run(O)
    null_nsigs[ii] = nullSess.run(cCnt)
    #Record how much we've changed since we last checked
    diff = (last-null_nll[ii])
    last = null_nll[ii]
    diffCnts = (lastNonZero - null_nsigs[ii])
    lastNonZero = null_nsigs[ii]
    #Calculate per-element summaries
    sigsPerSample = lastNonZero/n
    llPerEntry = null_nll[ii]/n/p
    #And the average intercept
    avgInt = np.mean(np.exp(nullSess.run(int0)))
    #The average coverage relative to the observed
    avgCov = np.mean(nullSess.run(y).sum(axis=0)/nullSess.run(k).sum(axis=0))
    print("[%s] step %d, training O=%g, cnt=%g,dO=%g,dCnt=%g,nSigsAvg=%g/%d,avgNLL=%g,avgCov=%g" %(datetime.datetime.now(),i,last,lastNonZero,diff,diffCnts,sigsPerSample,s+1,llPerEntry,avgCov))
    #Test if we should terminate
    if diff<=0 and diff/last < args.log_likelihood_tolerance  and diffCnts/lastNonZero < args.sparsity_tolerance:
      break
  i = i+1
  if i>args.max_it:
    break

################
# Update model #
################
#Get the NULL likelihoods per samples
sampL = nullSess.run(LL).sum(axis=1)
#And the penalty under the assumption of uniform exposures, that total to the number of reads
#This is the average number of molecules per sample
mols_per_samp = (toc/tol).sum(axis=0)*args.insert_size
#NOTE: This sets the scale for lambda so that for lambda=1 the sample -LL under the NULL model is equal to the imposed penalty if exposures are uniformly distributed.
#The L1 Penalty
#Construct the total penalty under the uniformity and avg mol assumptions.  The denominator is really s*(mols_per_samp/s)
avg_pen = mols_per_samp
avg_pen = sampL/avg_pen
#And the penalty itself
penL1 = args.l1_lambda*tf.reduce_sum(avg_pen.values*E)
#The L2 penalty, does not include lambda (we don't want to penalise large exposures in intercept)
#Under the uniformity assumption and sum to avg mol assumption, this is the total penalty
#NOTE: The s+1 instead of s is to account for there being the unpenalized intercept in the model
avg_pen = s*((mols_per_samp/(s+1))**2)
avg_pen = sampL/avg_pen
#Don't want to include the intercept so have to re-exponetiate
penL2 = tf.exp(z)**2
penL2 = args.l2_lambda*tf.reduce_sum(avg_pen.values*penL2)
#And the penalised likelihood
O_pen = O+penL1+penL2


############
# Main fit #
############
print('''
######################
# Fitting main Model #
######################
''')
#Define optimisers
toLearn = [z,int0]
learners = opt_E.minimize(O_pen,var_list=toLearn,name='learn_exposures')
#Initialise
sess = tf.compat.v1.Session()
init = tf.compat.v1.global_variables_initializer()
sess.run(init)
#Record initial values
last = sess.run(O_pen)
lastNonZero = sess.run(cCnt)
#Record the movements
nll = np.zeros(int(ceil(args.max_it/args.poll_interval)))
nsigs = np.zeros(int(ceil(args.max_it/args.poll_interval)))
i=0
while True:
  #Take the exposure step
  sess.run([learners])
  #Every now and then, record our progress and check if we've converged
  if i%args.poll_interval == 0:
    ii = i//args.poll_interval
    #Record object function and number of non-zero exposures
    nll[ii] = sess.run(O_pen)
    nsigs[ii] = sess.run(cCnt)
    #Record how much we've changed since we last checked
    diff = (last-nll[ii])
    last = nll[ii]
    diffCnts = (lastNonZero - nsigs[ii])
    lastNonZero = nsigs[ii]
    #Calculate per-element summaries
    sigsPerSample = lastNonZero/n
    llPerEntry = nll[ii]/n/p
    #And the average intercept
    avgInt = np.mean(np.exp(sess.run(int0)))
    #The average coverage relative to the observed
    avgCov = np.mean(sess.run(y).sum(axis=0)/sess.run(k).sum(axis=0))
    print("[%s] step %d, training O=%g, cnt=%g,dO=%g,dCnt=%g,nSigsAvg=%g/%d,avgNLL=%g,avgCov=%g" %(datetime.datetime.now(),i,last,lastNonZero,diff,diffCnts,sigsPerSample,s+1,llPerEntry,avgCov))
    #Test if we should terminate
    if diff<0 and diff/last < args.log_likelihood_tolerance  and diffCnts/lastNonZero < args.sparsity_tolerance:
      break
  i = i+1
  if i>args.max_it:
    break


###################
# Post processing #
###################
#First save the things used to fit.
#The bulk counts
toc.to_csv(args.output + '_usedBulkCounts.tsv',sep='\t',index_label=False)
#And their lengths
tol.to_csv(args.output + '_usedBulkGeneLengths.tsv',sep='\t',index_label=False)
#The cellular Signals
scSigs.to_csv(args.output + '_usedCellularSignals.tsv',sep='\t',index_label=False)
#And gene weights
geneWeights.to_csv(args.output + '_usedGeneWeights.tsv',sep='\t',index_label=False)
#Now the things we infered
#The exposures, plus goodness of fit metrics
pred_E = pd.DataFrame(sess.run(E),index=list(scSigs.columns)+['Intercept'],columns=toc.columns)
pR2 = 1-sess.run(LL).sum(axis=1)/nullSess.run(LL).sum(axis=1)
pred_E.loc['pR2'] = pR2
pred_E.loc['fitCount'] = sess.run(y).sum(axis=0)
pred_E.loc['obsCount'] = sess.run(k).sum(axis=0)
pred_E.to_csv(args.output + '_fitExposures.tsv',sep='\t',index_label=False)
#Full predicted table of counts
pred_y = pd.DataFrame(sess.run(y),index=toc.index,columns=toc.columns)
pred_y.to_csv(args.output + '_fitCounts.tsv',sep='\t',index_label=False)
#Full negative log-likelihood table
pred_LL = pd.DataFrame(sess.run(LL),index=toc.columns,columns=toc.index)
pred_LL.to_csv(args.output + '_negativeLogLikelihoodFullFit.tsv',sep='\t',index_label=False)
#Full negative log-likelihood table under null
pred_nLL = pd.DataFrame(nullSess.run(LL),index=toc.columns,columns=toc.index)
pred_nLL.to_csv(args.output + '_negativeLogLikelihoodNullFit.tsv',sep='\t',index_label=False)
#Stats about fitting itself
#Main fit
df = pd.DataFrame(np.c_[nll,nsigs],columns=['NLL','numExposures'])
df = df.iloc[np.logical_and(nll!=0,nsigs!=0),]
df.loc[:,'step'] = df.index*args.poll_interval
df.loc[:,'fitType']='FullFit'
fitStats = df
#Null Fit
df = pd.DataFrame(np.c_[null_nll,null_nsigs],columns=['NLL','numExposures'])
df = df.iloc[np.logical_and(null_nll!=0,null_nsigs!=0),]
df.loc[:,'step'] = df.index*args.poll_interval
df.loc[:,'fitType']='NullFit'
fitStats = pd.concat([fitStats,df])
fitStats.to_csv(args.output + '_fitStats.tsv',sep='\t',index=False)
```
