## Supplementary material for "Single cell derived mRNA signals across human kidney tumors": Source Code: cellSignalAnalysis.R

```
#' A script to visualise the results generated by running cellSignalAnalysis.py
#'
#' Note: The process of writing out and reading in sample names can cause issues.  Particularly if those sample IDs are foolish enough to contain dashes, which R will convert to '.' when it reads them in.

##############
# Libraries  #
##############
library(ComplexHeatmap)

#' Load the fitted exposures and normalise them
#'
#' Load's the exposure table, separates out the goodness-of-fit metrics and exposures and normalises the exposures to sum to 1 across each sample.
#'
#' @param tgt The relevant *_fitExposures.tsv file that results should be loaded from.  Can either be the exact file, or the base part to which _fitExposures.tsv will be added.
#' @return A list containing the normalised exposure table, raw exposure table, and a goodness-of-fit table
normaliseExposures = function(tgt){
  #Is this just the base?
  if(file.exists(paste0(tgt,'_fitExposures.tsv')))
    tgt = paste0(tgt,'_fitExposures.tsv')
  fit = read.table(tgt,sep='\t',header=TRUE)
  #Extract the goodness of fit rows
  gofNoms = c('pR2','fitCount','obsCount')
  gof = fit[gofNoms,]
  gof['log2(countRatio)',] = log2(unlist(gof['fitCount',]/gof['obsCount',]))
  #And the exposure table
  exposures = fit[-match(gofNoms,rownames(fit)),]
  #Normalise
  exposures = t(t(exposures)/colSums(exposures))
  #Done
  return(list(exposures=exposures,gof=gof,raw=fit[-match(gofNoms,rownames(fit)),]))
}


#' Plot normalised exposures
#' 
#' Takes the normalised exposure object produced by \code{normaliseExposures} and creates a basic heatmap to visualise the results.
#'
#' @param fit The normalised exposure object returned by \code{normaliseExposures}
#' @param exposureScale Range of exposures to show.
#' @param cluster_rows Should we cluster the rows.
#' @param ... Passed to Heatmap.
plotExposures = function(fit,exposureScale=c(0,0.5),cluster_rows=FALSE,...){
  #Colours for exposures
  hmCols = c('#ffffff','#f0f0f0','#d9d9d9','#bdbdbd','#969696','#737373','#525252','#252525','#000000')
  #Colours for pR2 metric
  pR2Cols  = c('#fff5eb','#fee6ce','#fdd0a2','#fdae6b','#fd8d3c','#f16913','#d94801','#a63603','#7f2704')
  #Colours for log library sizes
  libCols = c('#fcfbfd','#efedf5','#dadaeb','#bcbddc','#9e9ac8','#807dba','#6a51a3','#54278f','#3f007d')
  #And library size ratio
  libRatCols = c('#8c510a','#bf812d','#dfc27d','#f6e8c3','#f5f5f5','#c7eae5','#80cdc1','#35978f','#01665e')
  #Create the bottom annotation
  gof = fit$gof
  gof['fitCount',] = log10(gof['fitCount',])
  gof['obsCount',] = log10(gof['obsCount',])
  rownames(gof) = gsub('(.*Count)','log10(\\1)',rownames(gof))
  #Decide on range for library size
  libRange = range(gof[grep('Count',rownames(gof)),])
  #Convert colours to ramps
  bCols = circlize::colorRamp2(seq(0,1,length.out=length(pR2Cols)),pR2Cols)
  lCols = circlize::colorRamp2(seq(libRange[1],libRange[2],length.out=length(libCols)),libCols)
  hmColObj = circlize::colorRamp2(seq(exposureScale[1],exposureScale[2],length.out=length(hmCols)),hmCols)
  lrCols = circlize::colorRamp2(seq(-1,1,length.out=length(libRatCols)),libRatCols)
  botAnno = HeatmapAnnotation(df = t(gof),
                              annotation_name_side = 'left',
                              col = list(pR2 =bCols,
                                         `log10(fitCount)` = lCols,
                                         `log10(obsCount)` = lCols,
                                         `log2(countRatio)` = lrCols)
                              )
  hm = Heatmap((fit$exposures),
               col=hmColObj,
               name='Exposures',
               bottom_annotation=botAnno,
               cluster_rows = cluster_rows,
               ...
               )
  return(hm)
}
```
