## Supplementary material for "Single cell derived mRNA signals across human kidney tumors": Source Code: geneSets.R

```
#Non pseudo-autosomal genes on chrY.  Based on wikipedia definitions of pseudo autosomal regions and the following code
#gtf='~/trueHome/scratch/Common/refdata-cellranger-GRCh38-1.2.0/genes/genes.gtf'
#ann = rtracklayer::import(gtf)
#parY = GRanges('Y',IRanges(c(10001,56887903),c(2781479,57217415)))
#chrYnonPar = subsetByOverlaps(ann[seqnames(ann)=='Y',],parY,invert=TRUE)
#chrYnonPar = sort(unique(chrYnonPar$gene_name))
chrYnonPar = c("AC006328.4", "AC006386.1", "AC007359.6", "AC008175.1", "AC009491.1", 
"AC009491.2", "AC009977.1", "AC010084.1", "AC010723.1", "AC010891.2", 
"AC012005.3", "AC012005.4", "AC022486.1", "AC064829.1", "AMELY", 
"BPY2", "BPY2B", "BPY2C", "CDY1", "CDY1B", "CDY2A", "CDY2B", 
"CSPG4P1Y", "DAZ1", "DAZ2", "DAZ3", "DAZ4", "DDX3Y", "EIF1AY", 
"FAM197Y5P", "FAM224A", "FAM224B", "FAM41AY1", "FAM41AY2", "HSFY1", 
"HSFY2", "KDM5D", "LINC00266-4P", "LINC00278", "LINC00279", "LINC00280", 
"NLGN4Y", "NLGN4Y-AS1", "PCDH11Y", "PRORY", "PRY", "PRY2", "RBMY1A1", 
"RBMY1B", "RBMY1D", "RBMY1E", "RBMY1F", "RBMY1J", "RP11-122L9.1", 
"RP11-414C23.1", "RP11-424G14.1", "RP11-65G9.1", "RPS4Y1", "RPS4Y2", 
"SRY", "TBL1Y", "TGIF2LY", "TMSB4Y", "TSPY1", "TSPY10", "TSPY2", 
"TSPY3", "TSPY4", "TSPY8", "TSPY9P", "TTTY1", "TTTY10", "TTTY11", 
"TTTY12", "TTTY13", "TTTY14", "TTTY15", "TTTY16", "TTTY17A", 
"TTTY17B", "TTTY17C", "TTTY18", "TTTY19", "TTTY1B", "TTTY2", 
"TTTY20", "TTTY21", "TTTY21B", "TTTY22", "TTTY23", "TTTY23B", 
"TTTY2B", "TTTY3", "TTTY3B", "TTTY4", "TTTY4B", "TTTY4C", "TTTY5", 
"TTTY6", "TTTY6B", "TTTY7", "TTTY7B", "TTTY8", "TTTY8B", "TTTY9B", 
"USP9Y", "UTY", "VCY", "VCY1B", "ZFY", "ZFY-AS1")
#Housekeeping genes
#hkGenes = read.table('http://www.tau.ac.il/~elieis/HKG/HK_genes.txt',sep='\t')
hkGenes = c("AAAS", "AAGAB", "AAMP", "AAR2", "AARS", "AARS2", "AARSD1", 
"AASDHPPT", "AATF", "ABCB10", "ABCB7", "ABCD3", "ABCE1", "ABCF1", 
"ABCF2", "ABCF3", "ABHD10", "ABHD12", "ABHD13", "ABHD14A", "ABHD16A", 
"ABHD4", "ABHD8", "ABI1", "ABT1", "ACAD9", "ACADVL", "ACAP3", 
"ACBD3", "ACBD5", "ACBD6", "ACIN1", "ACLY", "ACOT13", "ACOT8", 
"ACOT9", "ACOX1", "ACOX3", "ACP1", "ACSF3", "ACSL3", "ACSS2", 
"ACTR10", "ACTR1A", "ACTR1B", "ACTR5", "ACTR8", "ACVR1", "ACVR1B", 
"ADCK2", "ADCK4", "ADH5", "ADI1", "ADIPOR1", "ADIPOR2", "ADK", 
"ADNP", "ADO", "ADPRH", "ADPRHL2", "ADPRM", "ADSL", "AES", "AFF4", 
"AFTPH", "AGFG1", "AGGF1", "AGPAT1", "AGPAT3", "AGPAT6", "AGPS", 
"AHCY", "AHSA1", "AIMP1", "AIP", "AK2", "AK3", "AKAP8", "AKAP9", 
"AKIP1", "AKIRIN1", "AKIRIN2", "AKR1A1", "AKR7A2", "AKT1", "AKT1S1", 
"AKTIP", "ALAD", "ALDH3A2", "ALDH9A1", "ALG11", "ALG5", "ALG8", 
"ALG9", "ALKBH1", "ALKBH2", "ALKBH3", "ALKBH5", "ALS2", "ALYREF", 
"AMBRA1", "AMD1", "ANAPC10", "ANAPC11", "ANAPC13", "ANAPC15", 
"ANAPC16", "ANAPC2", "ANAPC5", "ANAPC7", "ANKFY1", "ANKH", "ANKHD1", 
"ANKHD1-EIF4EBP3", "ANKRD10", "ANKRD17", "ANKRD28", "ANKRD39", 
"ANKRD46", "ANO6", "ANP32A", "ANP32B", "ANP32C", "ANP32E", "ANXA6", 
"ANXA7", "AP1B1", "AP1G1", "AP1M1", "AP2A1", "AP2A2", "AP2M1", 
"AP2S1", "AP3B1", "AP3D1", "AP3M1", "AP3S1", "AP3S2", "AP4B1", 
"AP5M1", "APEH", "APEX1", "APEX2", "APH1A", "API5", "APIP", "APOA1BP", 
"APOL2", "APOOL", "APOPT1", "APPL2", "APTX", "ARAF", "ARCN1", 
"ARF1", "ARF5", "ARF6", "ARFGAP2", "ARFGAP3", "ARFGEF2", "ARFIP1", 
"ARFIP2", "ARFRP1", "ARHGAP35", "ARHGAP5", "ARHGDIA", "ARHGEF10L", 
"ARHGEF11", "ARHGEF40", "ARIH1", "ARIH2", "ARIH2OS", "ARL1", 
"ARL14EP", "ARL5A", "ARL6IP4", "ARL8A", "ARL8B", "ARMC1", "ARMC10", 
"ARMC5", "ARMC6", "ARMC7", "ARMC8", "ARMCX3", "ARMCX5", "ARNT", 
"ARPC1A", "ARPC2", "ARPC5L", "ARV1", "ASB1", "ASB6", "ASB7", 
"ASB8", "ASCC1", "ASCC3", "ASF1A", "ASH2L", "ASNA1", "ASNSD1", 
"ASPSCR1", "ASUN", "ASXL1", "ATAD1", "ATAD3A", "ATE1", "ATF1", 
"ATF2", "ATF4", "ATF6", "ATF7", "ATF7IP", "ATG12", "ATG13", "ATG16L1", 
"ATG2A", "ATG2B", "ATG3", "ATG4B", "ATG4D", "ATG5", "ATG7", "ATIC", 
"ATL2", "ATMIN", "ATOX1", "ATP2C1", "ATP5A1", "ATP5B", "ATP5C1", 
"ATP5D", "ATP5F1", "ATP5G2", "ATP5G3", "ATP5H", "ATP5J", "ATP5J2", 
"ATP5J2-PTCD1", "ATP5L", "ATP5O", "ATP5S", "ATP5SL", "ATP6AP1", 
"ATP6V0A2", "ATP6V0B", "ATP6V0C", "ATP6V0D1", "ATP6V0E1", "ATP6V1C1", 
"ATP6V1D", "ATP6V1E1", "ATP6V1F", "ATP6V1G1", "ATP6V1H", "ATPAF2", 
"ATPIF1", "ATRAID", "ATRN", "ATXN10", "ATXN1L", "ATXN2", "ATXN2L", 
"ATXN7L3", "ATXN7L3B", "AUH", "AUP1", "AURKAIP1", "AXIN1", "AZI2", 
"AZIN1", "B3GALT6", "B4GALT3", "B4GALT5", "B4GALT7", "BABAM1", 
"BAD", "BAG1", "BAG4", "BAG6", "BAHD1", "BANF1", "BAP1", "BAZ1B", 
"BBS4", "BCAP29", "BCAP31", "BCAS2", "BCAT2", "BCCIP", "BCKDHA", 
"BCKDK", "BCL2L1", "BCL2L13", "BCL2L2-PABPN1", "BCL7B", "BCLAF1", 
"BCS1L", "BECN1", "BFAR", "BIRC2", "BIVM-ERCC5", "BLMH", "BLOC1S1", 
"BLOC1S2", "BLOC1S3", "BLOC1S4", "BLOC1S6", "BLZF1", "BMI1", 
"BMS1", "BNIP1", "BNIP2", "BOD1", "BOLA1", "BOLA3", "BPGM", "BPNT1", 
"BPTF", "BRAT1", "BRD2", "BRD4", "BRD7", "BRD9", "BRE", "BRF1", 
"BRF2", "BRIX1", "BRK1", "BRMS1", "BRPF1", "BRPF3", "BSDC1", 
"BSG", "BTBD2", "BTD", "BTF3", "BUB3", "BZW1", "C10orf12", "C10orf2", 
"C10orf76", "C10orf88", "C11orf1", "C11orf24", "C11orf31", "C11orf57", 
"C11orf58", "C11orf73", "C11orf83", "C12orf10", "C12orf23", "C12orf29", 
"C12orf44", "C12orf45", "C12orf5", "C12orf52", "C12orf57", "C12orf65", 
"C12orf66", "C14orf1", "C14orf119", "C14orf142", "C14orf166", 
"C14orf2", "C14orf28", "C15orf38-AP3S2", "C15orf57", "C16orf13", 
"C16orf62", "C16orf72", "C16orf91", "C17orf49", "C17orf51", "C17orf58", 
"C17orf59", "C17orf70", "C17orf85", "C18orf21", "C18orf25", "C18orf32", 
"C18orf8", "C19orf43", "C19orf53", "C19orf60", "C19orf70", "C1GALT1", 
"C1QBP", "C1orf109", "C1orf122", "C1orf123", "C1orf174", "C1orf43", 
"C1orf50", "C1orf52", "C20orf111", "C20orf24", "C21orf2", "C21orf33", 
"C21orf59", "C22orf28", "C22orf29", "C22orf32", "C2orf47", "C2orf49", 
"C2orf69", "C2orf74", "C2orf76", "C3orf17", "C3orf37", "C3orf38", 
"C3orf58", "C4orf27", "C4orf3", "C4orf52", "C5orf15", "C5orf24", 
"C6orf1", "C6orf106", "C6orf120", "C6orf136", "C6orf226", "C6orf47", 
"C6orf57", "C6orf62", "C6orf89", "C7orf25", "C7orf26", "C7orf49", 
"C7orf50", "C7orf55", "C7orf55-LUC7L2", "C7orf73", "C8orf33", 
"C8orf40", "C8orf59", "C8orf76", "C8orf82", "C9orf123", "C9orf16", 
"C9orf37", "C9orf64", "C9orf69", "C9orf78", "C9orf89", "CAB39", 
"CALCOCO2", "CALM1", "CALR", "CALU", "CAMTA1", "CAMTA2", "CANT1", 
"CANX", "CAPN1", "CAPN7", "CAPNS1", "CAPRIN1", "CAPZA2", "CAPZB", 
"CARKD", "CARS", "CARS2", "CASC3", "CASC4", "CASP3", "CASP7", 
"CASP9", "CBR4", "CBX3", "CBX5", "CC2D1A", "CC2D1B", "CCAR1", 
"CCBL1", "CCDC12", "CCDC124", "CCDC127", "CCDC130", "CCDC137", 
"CCDC149", "CCDC174", "CCDC22", "CCDC23", "CCDC25", "CCDC47", 
"CCDC50", "CCDC51", "CCDC59", "CCDC71", "CCDC86", "CCDC90A", 
"CCDC92", "CCDC94", "CCM2", "CCNB1IP1", "CCNDBP1", "CCNG1", "CCNH", 
"CCNK", "CCNL1", "CCNL2", "CCNY", "CCPG1", "CCT3", "CCT4", "CCT5", 
"CCT6A", "CCT7", "CCT8", "CD164", "CD320", "CD46", "CD63", "CD81", 
"CD82", "CD99L2", "CDC123", "CDC16", "CDC23", "CDC27", "CDC37", 
"CDC37L1", "CDC40", "CDC42", "CDC5L", "CDIP1", "CDIPT", "CDK12", 
"CDK13", "CDK16", "CDK2AP1", "CDK4", "CDK5RAP1", "CDK8", "CDK9", 
"CDS2", "CDV3", "CDYL", "CEBPG", "CEBPZ", "CECR5", "CELF1", "CENPB", 
"CENPT", "CEP104", "CEP57", "CEP63", "CERK", "CERS2", "CGGBP1", 
"CHAMP1", "CHCHD1", "CHCHD2", "CHCHD3", "CHCHD4", "CHCHD5", "CHCHD7", 
"CHD1L", "CHD4", "CHD8", "CHERP", "CHID1", "CHKB", "CHMP1A", 
"CHMP2A", "CHMP2B", "CHMP4A", "CHMP4B", "CHMP5", "CHMP6", "CHP1", 
"CHPT1", "CHRAC1", "CHST12", "CHST7", "CHTOP", "CHUK", "CHURC1", 
"CHURC1-FNTB", "CIAO1", "CIB1", "CIC", "CINP", "CIR1", "CIRH1A", 
"CISD1", "CISD2", "CISD3", "CKAP4", "CLCC1", "CLCN3", "CLCN7", 
"CLINT1", "CLK3", "CLNS1A", "CLOCK", "CLP1", "CLPP", "CLPTM1", 
"CLPTM1L", "CLPX", "CLTA", "CLTB", "CLTC", "CMAS", "CMC1", "CMC2", 
"CMC4", "CMPK1", "CNBP", "CNIH", "CNIH4", "CNNM2", "CNNM3", "CNOT1", 
"CNOT11", "CNOT2", "CNOT3", "CNOT4", "CNOT7", "CNST", "COA1", 
"COA3", "COA4", "COA5", "COA6", "COASY", "COG1", "COG2", "COG3", 
"COG4", "COG7", "COG8", "COMMD1", "COMMD10", "COMMD3", "COMMD3-BMI1", 
"COMMD5", "COMMD6", "COMMD7", "COMMD9", "COMT", "COPA", "COPB1", 
"COPB2", "COPE", "COPG1", "COPS2", "COPS3", "COPS4", "COPS5", 
"COPS6", "COPS7A", "COPS7B", "COPS8", "COPZ1", "COQ10B", "COQ2", 
"COQ4", "COQ5", "COQ6", "CORO1C", "COX11", "COX14", "COX15", 
"COX16", "COX19", "COX20", "COX4I1", "COX5B", "COX6B1", "COX6C", 
"COX7A2", "COX7A2L", "COX7C", "COX8A", "CPD", "CPNE1", "CPNE2", 
"CPNE3", "CPOX", "CPSF2", "CPSF3L", "CPSF4", "CPSF6", "CPSF7", 
"CRADD", "CRBN", "CRCP", "CREB3", "CREBZF", "CREG1", "CRELD1", 
"CRIPAK", "CRIPT", "CRK", "CRKL", "CRLS1", "CRNKL1", "CRTC2", 
"CRY2", "CSGALNACT2", "CSNK1A1", "CSNK1A1L", "CSNK1D", "CSNK1G3", 
"CSNK2A3", "CSNK2B", "CSRP2BP", "CST3", "CSTB", "CSTF1", "CSTF2T", 
"CTAGE5", "CTBP1", "CTCF", "CTDSP2", "CTNNA1", "CTNNB1", "CTNNBIP1", 
"CTNNBL1", "CTNND1", "CTSA", "CTSD", "CTTN", "CTU2", "CUEDC2", 
"CUL1", "CUL2", "CUL4A", "CUL4B", "CUL5", "CUTA", "CUX1", "CWC15", 
"CWC22", "CWC25", "CXXC1", "CXXC5", "CXorf40A", "CXorf40B", "CXorf56", 
"CYB5B", "CYB5D2", "CYB5R3", "CYC1", "CYFIP1", "CYHR1", "CYP2U1", 
"D2HGDH", "DAD1", "DAG1", "DAGLB", "DALRD3", "DAP3", "DARS", 
"DARS2", "DAXX", "DAZAP1", "DBT", "DCAF10", "DCAF11", "DCAF12", 
"DCAF13", "DCAF5", "DCAF7", "DCAF8", "DCAKD", "DCTD", "DCTN2", 
"DCTN3", "DCTN4", "DCTN5", "DCTN6", "DCTPP1", "DCUN1D3", "DCUN1D4", 
"DCUN1D5", "DDA1", "DDB1", "DDB2", "DDOST", "DDRGK1", "DDX1", 
"DDX10", "DDX17", "DDX18", "DDX19A", "DDX19B", "DDX21", "DDX23", 
"DDX24", "DDX27", "DDX39B", "DDX3X", "DDX41", "DDX42", "DDX46", 
"DDX47", "DDX49", "DDX54", "DDX56", "DDX59", "DEDD", "DEF8", 
"DEGS1", "DEK", "DENND1A", "DENND4A", "DENR", "DERA", "DERL1", 
"DERL2", "DESI1", "DEXI", "DFFA", "DGCR14", "DGCR2", "DGCR6L", 
"DHPS", "DHRS12", "DHRS7B", "DHX15", "DHX16", "DHX29", "DHX30", 
"DHX32", "DHX33", "DHX36", "DHX38", "DHX8", "DHX9", "DIABLO", 
"DIDO1", "DIEXF", "DIMT1", "DIRC2", "DIS3", "DIS3L2", "DKC1", 
"DLD", "DLG1", "DLGAP4", "DLST", "DMAP1", "DNAAF2", "DNAJA2", 
"DNAJA3", "DNAJB11", "DNAJB12", "DNAJB9", "DNAJC10", "DNAJC11", 
"DNAJC14", "DNAJC17", "DNAJC19", "DNAJC2", "DNAJC21", "DNAJC3", 
"DNAJC4", "DNAJC5", "DNAJC7", "DNAJC8", "DNAJC9", "DNASE2", "DNLZ", 
"DNM1L", "DNM2", "DNTTIP1", "DNTTIP2", "DOHH", "DOLK", "DPAGT1", 
"DPH1", "DPH2", "DPH3", "DPH5", "DPM1", "DPP7", "DPY30", "DR1", 
"DRAM2", "DRAP1", "DRG2", "DROSHA", "DSCR3", "DTWD1", "DUSP11", 
"DUSP14", "DUSP16", "DUSP22", "DUT", "DVL3", "DYM", "DYNC1LI1", 
"DYNLL2", "DYNLRB1", "DYNLT1", "E2F4", "E4F1", "EAF1", "EAPP", 
"EARS2", "EBAG9", "EBNA1BP2", "ECD", "ECH1", "ECHDC1", "ECHS1", 
"ECI1", "ECI2", "ECSIT", "EDC3", "EDC4", "EDEM3", "EDF1", "EED", 
"EEF1B2", "EEF1E1", "EEF2", "EEFSEC", "EFCAB14", "EFHA1", "EFR3A", 
"EFTUD1", "EFTUD2", "EGLN2", "EHMT1", "EI24", "EID2", "EIF1", 
"EIF1AD", "EIF1B", "EIF2A", "EIF2AK1", "EIF2AK3", "EIF2AK4", 
"EIF2B2", "EIF2B3", "EIF2B4", "EIF2B5", "EIF2D", "EIF2S1", "EIF2S2", 
"EIF3A", "EIF3B", "EIF3D", "EIF3E", "EIF3G", "EIF3H", "EIF3I", 
"EIF3J", "EIF3K", "EIF3L", "EIF3M", "EIF4A1", "EIF4A3", "EIF4E2", 
"EIF4G1", "EIF4G2", "EIF4G3", "EIF4H", "EIF5", "EIF5A", "EIF5AL1", 
"EIF5B", "EIF6", "ELAC2", "ELAVL1", "ELF2", "ELK1", "ELK4", "ELL2", 
"ELMOD3", "ELOVL1", "ELP2", "ELP3", "ELP4", "ELP6", "EMC1", "EMC10", 
"EMC2", "EMC3", "EMC4", "EMC6", "EMC7", "EMC8", "EMC9", "EMD", 
"EMG1", "ENDOG", "ENOPH1", "ENSA", "ENTPD4", "ENTPD6", "ENY2", 
"EPC1", "EPM2AIP1", "EPN1", "EPRS", "ERAL1", "ERAP1", "ERCC1", 
"ERCC2", "ERCC3", "ERCC5", "ERGIC2", "ERGIC3", "ERH", "ERI3", 
"ERICH1", "ERLEC1", "ERO1L", "ERP44", "ESD", "ESF1", "ETF1", 
"ETFA", "ETFB", "ETV6", "EWSR1", "EXD2", "EXOC1", "EXOC2", "EXOC3", 
"EXOC4", "EXOC7", "EXOC8", "EXOSC1", "EXOSC10", "EXOSC2", "EXOSC4", 
"EXOSC7", "EXOSC8", "EXT2", "EXTL3", "FADD", "FAF1", "FAF2", 
"FAHD1", "FAM104B", "FAM108A1", "FAM108B1", "FAM114A2", "FAM118B", 
"FAM120A", "FAM120AOS", "FAM120B", "FAM122A", "FAM127B", "FAM134A", 
"FAM134C", "FAM136A", "FAM149B1", "FAM160A2", "FAM160B1", "FAM160B2", 
"FAM162A", "FAM168B", "FAM173A", "FAM173B", "FAM174A", "FAM175B", 
"FAM177A1", "FAM178A", "FAM192A", "FAM199X", "FAM200A", "FAM204A", 
"FAM206A", "FAM208B", "FAM20B", "FAM210B", "FAM32A", "FAM35A", 
"FAM3A", "FAM50A", "FAM50B", "FAM58A", "FAM63A", "FAM73B", "FAM8A1", 
"FAM96A", "FAM96B", "FAM98A", "FARS2", "FARSA", "FARSB", "FASTK", 
"FASTKD2", "FASTKD5", "FBRSL1", "FBXL15", "FBXL17", "FBXL3", 
"FBXL4", "FBXL5", "FBXL6", "FBXO11", "FBXO18", "FBXO22", "FBXO28", 
"FBXO3", "FBXO38", "FBXO42", "FBXO45", "FBXO6", "FBXO7", "FBXW11", 
"FBXW2", "FBXW4", "FBXW5", "FBXW7", "FCF1", "FDFT1", "FDPS", 
"FDX1", "FECH", "FEM1C", "FEN1", "FEZ2", "FGFR1OP2", "FH", "FIBP", 
"FICD", "FIP1L1", "FIS1", "FIZ1", "FKBP3", "FKBP8", "FKBPL", 
"FKRP", "FLAD1", "FLCN", "FLOT1", "FLOT2", "FNDC3A", "FNTA", 
"FNTB", "FOPNL", "FOXK2", "FOXP4", "FOXRED1", "FPGS", "FPGT", 
"FRA10AC1", "FTO", "FTSJ1", "FTSJ2", "FTSJ3", "FTSJD1", "FTSJD2", 
"FUBP1", "FUK", "FUNDC2", "FXN", "FYTTD1", "FZR1", "G3BP1", "GAA", 
"GABARAP", "GABARAPL2", "GABPB1", "GADD45GIP1", "GALK2", "GALNS", 
"GALNT1", "GALNT2", "GALT", "GANAB", "GAPVD1", "GARS", "GART", 
"GATAD2A", "GATAD2B", "GATC", "GBA", "GBA2", "GBF1", "GCC1", 
"GCDH", "GCLC", "GCLM", "GDE1", "GDI2", "GDPGP1", "GEMIN7", "GEMIN8", 
"GET4", "GFER", "GFM1", "GFOD2", "GGCT", "GGNBP2", "GGT7", "GHDC", 
"GHITM", "GID8", "GINM1", "GIPC1", "GLCE", "GLE1", "GLG1", "GLI4", 
"GLO1", "GLRX2", "GLRX3", "GLRX5", "GLT8D1", "GLTP", "GLTPD1", 
"GLYR1", "GMPPA", "GMPR2", "GNB1", "GNB2", "GNE", "GNL2", "GNL3", 
"GNPAT", "GNPDA1", "GNPNAT1", "GNPTG", "GNS", "GOLGA1", "GOLGA2", 
"GOLGA3", "GOLGA5", "GOLGA7", "GOLGB1", "GOLPH3", "GOLT1B", "GOPC", 
"GORASP1", "GORASP2", "GOSR1", "GOSR2", "GPAA1", "GPANK1", "GPATCH4", 
"GPBP1", "GPBP1L1", "GPHN", "GPI", "GPKOW", "GPN1", "GPN2", "GPN3", 
"GPR107", "GPR108", "GPS1", "GPS2", "GPX4", "GRAMD4", "GRHPR", 
"GRINA", "GRIPAP1", "GRPEL1", "GRSF1", "GRWD1", "GSK3A", "GSK3B", 
"GSPT1", "GSPT2", "GSR", "GSS", "GSTK1", "GSTM4", "GSTO1", "GTDC2", 
"GTF2A1", "GTF2B", "GTF2F1", "GTF2F2", "GTF2H1", "GTF2H4", "GTF2H5", 
"GTF2I", "GTF3A", "GTF3C1", "GTF3C2", "GTF3C3", "GTF3C5", "GTF3C6", 
"GTPBP10", "GTPBP4", "GTPBP5", "GTPBP8", "GUK1", "GZF1", "H1FX", 
"H2AFV", "H2AFX", "H2AFY", "H2AFZ", "HADH", "HADHA", "HAGH", 
"HARS", "HARS2", "HAT1", "HAUS3", "HAUS4", "HAUS7", "HAX1", "HBP1", 
"HBS1L", "HCCS", "HCFC1", "HDAC2", "HDAC3", "HDAC6", "HDAC8", 
"HDDC3", "HDGF", "HDHD3", "HDLBP", "HEATR2", "HEATR5A", "HEBP1", 
"HECTD3", "HELZ", "HEMK1", "HERC4", "HERPUD1", "HERPUD2", "HEXA", 
"HEXDC", "HEXIM1", "HGS", "HIAT1", "HIATL1", "HIBADH", "HIGD1A", 
"HIGD2A", "HINFP", "HINT1", "HINT2", "HIST1H2BC", "HIVEP1", "HMBS", 
"HMG20A", "HMG20B", "HMGB1", "HMGN3", "HMGXB3", "HMGXB4", "HMOX2", 
"HN1L", "HNRNPA0", "HNRNPA2B1", "HNRNPAB", "HNRNPC", "HNRNPD", 
"HNRNPF", "HNRNPH1", "HNRNPH2", "HNRNPK", "HNRNPL", "HNRNPM", 
"HNRNPR", "HNRNPU", "HNRNPUL1", "HNRNPUL2", "HNRPDL", "HNRPLL", 
"HP1BP3", "HPS1", "HPS6", "HS1BP3", "HS2ST1", "HS6ST1", "HSBP1", 
"HSCB", "HSD17B10", "HSD17B12", "HSD17B4", "HSPA14", "HSPA4", 
"HSPA5", "HSPA8", "HSPA9", "HSPBP1", "HSPE1-MOB4", "HTATIP2", 
"HTRA2", "HTT", "HUS1", "HUWE1", "HYOU1", "HYPK", "IAH1", "IARS", 
"IARS2", "IBA57", "IBTK", "ICK", "ICMT", "ICT1", "IDE", "IDH3A", 
"IDH3B", "IDH3G", "IDI1", "IER3IP1", "IFNAR1", "IFNGR1", "IFRD1", 
"IFT27", "IKZF5", "IL13RA1", "IL6ST", "ILF2", "ILKAP", "ILVBL", 
"IMMT", "IMP3", "IMP4", "IMPAD1", "INF2", "ING1", "INO80B", "INO80E", 
"INPP5A", "INPP5K", "INSIG2", "INTS1", "INTS10", "INTS12", "INTS3", 
"INTS4", "INVS", "IP6K1", "IP6K2", "IPO7", "IPO8", "IPO9", "IRAK1", 
"IREB2", "IRF2BP1", "IRF2BP2", "IRF2BPL", "IRGQ", "ISCU", "ISOC2", 
"IST1", "ISY1", "ISY1-RAB43", "ITCH", "ITFG1", "ITFG3", "ITGB1", 
"ITGB1BP1", "ITM2B", "ITPA", "ITPK1", "ITPKC", "ITPRIPL2", "IVNS1ABP", 
"IWS1", "JAGN1", "JAK1", "JKAMP", "JMJD4", "JMJD6", "JMJD7", 
"JMJD8", "JOSD2", "JTB", "JUND", "KANSL2", "KANSL3", "KARS", 
"KAT2B", "KAT5", "KAT8", "KBTBD2", "KBTBD4", "KBTBD7", "KCMF1", 
"KCTD20", "KCTD21", "KCTD6", "KDM2A", "KDM4A", "KDM5C", "KDSR", 
"KHDRBS1", "KHNYN", "KHSRP", "KIAA0100", "KIAA0141", "KIAA0195", 
"KIAA0196", "KIAA0232", "KIAA0319L", "KIAA0391", "KIAA0754", 
"KIAA0947", "KIAA1143", "KIAA1191", "KIAA1429", "KIAA1430", "KIAA1586", 
"KIAA1704", "KIAA1715", "KIAA1919", "KIAA1967", "KIAA2013", "KLC4", 
"KLF3", "KLF9", "KLHDC2", "KLHDC3", "KLHL20", "KLHL25", "KLHL36", 
"KLHL5", "KLHL8", "KPNA1", "KPNB1", "KRCC1", "KRR1", "KTI12", 
"KTN1", "KXD1", "L3MBTL2", "LACTB", "LAGE3", "LAMP1", "LAMP2", 
"LAMTOR1", "LAMTOR2", "LAMTOR3", "LAMTOR4", "LAMTOR5", "LAP3", 
"LAPTM4A", "LARP1", "LARP4", "LARP7", "LARS2", "LCOR", "LDHA", 
"LEMD2", "LENG1", "LEPROT", "LETM1", "LETMD1", "LGALSL", "LHPP", 
"LIAS", "LIG3", "LIG4", "LIN37", "LIN54", "LIN7C", "LINS", "LIPT1", 
"LMAN1", "LMBRD1", "LMF2", "LMO4", "LNX2", "LOC100129361", "LOC100289561", 
"LOC441155", "LOC729020", "LONP1", "LONP2", "LPCAT3", "LPIN1", 
"LPPR2", "LRFN3", "LRPAP1", "LRPPRC", "LRRC14", "LRRC24", "LRRC28", 
"LRRC40", "LRRC41", "LRRC42", "LRRC47", "LRRC57", "LRRC59", "LRRC8A", 
"LRRFIP2", "LRSAM1", "LSG1", "LSM1", "LSM10", "LSM14A", "LSM14B", 
"LSM2", "LSM3", "LSM4", "LSM5", "LSM6", "LSM7", "LSMD1", "LSS", 
"LTV1", "LUC7L2", "LUC7L3", "LUZP6", "LYRM1", "LYRM4", "LYRM5", 
"LYSMD1", "LYSMD3", "LYSMD4", "LZTR1", "M6PR", "MAD2L1BP", "MAD2L2", 
"MAEA", "MAGED1", "MAGEF1", "MAGOH", "MAGT1", "MAK16", "MALSU1", 
"MAN1A2", "MAN1B1", "MAN2A2", "MAN2B2", "MAN2C1", "MAP1LC3B2", 
"MAP2K1", "MAP2K2", "MAP2K5", "MAP3K7", "MAP4K4", "MAPK1", "MAPK1IP1L", 
"MAPK6", "MAPK8", "MAPK9", "MAPKAP1", "MAPKAPK2", "MAPKAPK5", 
"MAPRE2", "MARCH2", "MARCH5", "MARCH6", "MARCH7", "MARK3", "MARK4", 
"MARS", "MARS2", "MAT2B", "MAVS", "MAX", "MAZ", "MBD1", "MBD2", 
"MBD3", "MBD4", "MBLAC1", "MBNL2", "MBTPS1", "MBTPS2", "MCAT", 
"MCCC1", "MCEE", "MCFD2", "MCM3AP", "MCM7", "MCMBP", "MCOLN1", 
"MCPH1", "MCRS1", "MCTS1", "MCU", "MDC1", "MDP1", "ME2", "MEAF6", 
"MECP2", "MED10", "MED11", "MED13", "MED14", "MED16", "MED19", 
"MED20", "MED21", "MED24", "MED29", "MED31", "MED4", "MED6", 
"MED7", "MED8", "MEF2A", "MEF2BNB", "MEMO1", "MEN1", "MEPCE", 
"METAP1", "METAP2", "METRN", "METTL13", "METTL14", "METTL16", 
"METTL17", "METTL18", "METTL20", "METTL21A", "METTL23", "METTL2A", 
"METTL2B", "METTL3", "METTL5", "MFAP1", "MFAP3", "MFF", "MFN1", 
"MFSD11", "MFSD12", "MFSD3", "MFSD5", "MGAT2", "MGAT4B", "MGME1", 
"MGMT", "MGRN1", "MGST3", "MIA3", "MIB1", "MICALL1", "MICU1", 
"MID1IP1", "MIDN", "MIEN1", "MIER1", "MIF", "MIF4GD", "MIIP", 
"MINOS1", "MIS12", "MITD1", "MKI67IP", "MKKS", "MKLN1", "MKNK1", 
"MKRN2", "MLEC", "MLF2", "MLH1", "MLLT1", "MLLT10", "MLST8", 
"MLX", "MMAA", "MMADHC", "MMS19", "MNAT1", "MNF1", "MOB4", "MOGS", 
"MON1A", "MON2", "MORC2", "MORF4L2", "MOSPD1", "MPC2", "MPDU1", 
"MPG", "MPHOSPH10", "MPI", "MPLKIP", "MPND", "MPPE1", "MPV17L2", 
"MRFAP1", "MRFAP1L1", "MRI1", "MRM1", "MRP63", "MRPL1", "MRPL10", 
"MRPL11", "MRPL12", "MRPL13", "MRPL14", "MRPL15", "MRPL16", "MRPL17", 
"MRPL18", "MRPL19", "MRPL2", "MRPL20", "MRPL21", "MRPL22", "MRPL23", 
"MRPL24", "MRPL27", "MRPL28", "MRPL3", "MRPL30", "MRPL32", "MRPL33", 
"MRPL35", "MRPL36", "MRPL37", "MRPL38", "MRPL4", "MRPL40", "MRPL41", 
"MRPL42", "MRPL43", "MRPL44", "MRPL45", "MRPL46", "MRPL47", "MRPL48", 
"MRPL49", "MRPL50", "MRPL51", "MRPL52", "MRPL53", "MRPL54", "MRPL55", 
"MRPL9", "MRPS10", "MRPS11", "MRPS12", "MRPS14", "MRPS15", "MRPS16", 
"MRPS17", "MRPS18A", "MRPS18B", "MRPS18C", "MRPS2", "MRPS21", 
"MRPS22", "MRPS23", "MRPS24", "MRPS25", "MRPS26", "MRPS27", "MRPS28", 
"MRPS30", "MRPS31", "MRPS33", "MRPS34", "MRPS35", "MRPS5", "MRPS6", 
"MRPS7", "MRPS9", "MRRF", "MRS2", "MRTO4", "MSANTD3", "MSH3", 
"MSH6", "MSL3", "MSMP", "MSRA", "MSRB2", "MTA2", "MTCH1", "MTCH2", 
"MTDH", "MTERFD1", "MTERFD2", "MTERFD3", "MTFMT", "MTFR1", "MTFR1L", 
"MTIF3", "MTM1", "MTMR1", "MTMR3", "MTMR6", "MTO1", "MTPAP", 
"MTRR", "MTSS1", "MTX2", "MUL1", "MUS81", "MUT", "MVD", "MXD4", 
"MXI1", "MYBBP1A", "MYEOV2", "MYL12B", "MYNN", "MYO1E", "MYPOP", 
"MZF1", "MZT2A", "MZT2B", "N4BP1", "N4BP2L2", "NAA10", "NAA15", 
"NAA20", "NAA38", "NAA50", "NAA60", "NABP2", "NACA", "NACA2", 
"NACC1", "NACC2", "NAE1", "NAMPT", "NANS", "NAP1L4", "NAPA", 
"NARF", "NARFL", "NARG2", "NARS", "NARS2", "NAT10", "NBN", "NBR1", 
"NCAPH2", "NCBP2", "NCK1", "NCKIPSD", "NCL", "NCLN", "NCOA1", 
"NCOA6", "NCOR1", "NCSTN", "NDEL1", "NDFIP1", "NDNL2", "NDST1", 
"NDUFA10", "NDUFA11", "NDUFA12", "NDUFA13", "NDUFA2", "NDUFA3", 
"NDUFA4", "NDUFA5", "NDUFA6", "NDUFA7", "NDUFA8", "NDUFA9", "NDUFAF2", 
"NDUFAF3", "NDUFAF4", "NDUFB10", "NDUFB11", "NDUFB2", "NDUFB3", 
"NDUFB4", "NDUFB5", "NDUFB6", "NDUFB7", "NDUFB8", "NDUFB9", "NDUFC1", 
"NDUFC2", "NDUFC2-KCTD14", "NDUFS2", "NDUFS3", "NDUFS4", "NDUFS5", 
"NDUFS6", "NDUFS7", "NDUFS8", "NDUFV1", "NDUFV2", "NECAP1", "NEDD8", 
"NEDD8-MDP1", "NEIL2", "NEK4", "NEK9", "NELFB", "NELFCD", "NELFE", 
"NENF", "NEU1", "NF2", "NFATC2IP", "NFE2L2", "NFIL3", "NFKBIB", 
"NFKBIL1", "NFU1", "NFX1", "NFYB", "NFYC", "NGDN", "NGLY1", "NGRN", 
"NHP2", "NHP2L1", "NIF3L1", "NINJ1", "NIP7", "NIPA2", "NIPBL", 
"NISCH", "NIT1", "NIT2", "NKAP", "NKIRAS2", "NMD3", "NME1-NME2", 
"NME2", "NME3", "NME6", "NMRK1", "NMT1", "NOA1", "NOB1", "NOC2L", 
"NOL10", "NOL11", "NOL12", "NOL6", "NOL7", "NOL8", "NOLC1", "NOM1", 
"NONO", "NOP10", "NOP14", "NOP16", "NOP2", "NOP56", "NOP58", 
"NOP9", "NPC1", "NPC2", "NPLOC4", "NPRL2", "NPRL3", "NQO2", "NR1H2", 
"NR2C1", "NR2C2AP", "NR3C2", "NRBP1", "NRDE2", "NRIP1", "NSA2", 
"NSD1", "NSDHL", "NSFL1C", "NSMCE1", "NSMCE2", "NSMCE4A", "NSRP1", 
"NSUN2", "NSUN5", "NSUN6", "NT5C", "NT5C3", "NT5DC1", "NTAN1", 
"NTMT1", "NTPCR", "NUB1", "NUBP1", "NUBP2", "NUCB1", "NUCKS1", 
"NUDC", "NUDCD1", "NUDCD2", "NUDT14", "NUDT15", "NUDT2", "NUDT21", 
"NUDT22", "NUDT3", "NUDT9", "NUFIP2", "NUP107", "NUP133", "NUP153", 
"NUP54", "NUP62", "NUP85", "NUPL2", "NUTF2", "NXF1", "NXT1", 
"OAT", "OAZ1", "OAZ2", "OBFC1", "OCEL1", "OCIAD1", "ODC1", "OGFOD1", 
"OGFOD3", "OGFR", "OGG1", "OGT", "OLA1", "OPA1", "OPA3", "ORC4", 
"ORMDL1", "ORMDL2", "ORMDL3", "OS9", "OSBP", "OSBPL2", "OSBPL9", 
"OSGEP", "OSGIN2", "OSTM1", "OTUB1", "OTUD5", "OVCA2", "OXA1L", 
"OXNAD1", "P4HTM", "PA2G4", "PABPN1", "PACSIN2", "PAF1", "PAFAH1B1", 
"PAGR1", "PAICS", "PAIP1", "PAIP2", "PAK1IP1", "PAK2", "PAM16", 
"PANK2", "PANK3", "PANK4", "PANX1", "PAPD4", "PAPD7", "PAPOLA", 
"PARK7", "PARL", "PARN", "PARP1", "PARP3", "PARP9", "PATL1", 
"PATZ1", "PAXBP1", "PBDC1", "PBX2", "PCBP1", "PCBP2", "PCDHGB5", 
"PCF11", "PCGF1", "PCGF5", "PCID2", "PCIF1", "PCM1", "PCMT1", 
"PCNA", "PCNX", "PCNXL4", "PCSK7", "PCYOX1", "PCYT1A", "PDAP1", 
"PDCD2", "PDCD5", "PDCD6", "PDCD6IP", "PDE12", "PDE6D", "PDGFC", 
"PDHB", "PDHX", "PDK2", "PDLIM5", "PDP2", "PDS5A", "PDZD11", 
"PDZD8", "PEBP1", "PEF1", "PELO", "PELP1", "PEPD", "PES1", "PET100", 
"PET117", "PEX1", "PEX11A", "PEX11B", "PEX12", "PEX13", "PEX14", 
"PEX16", "PEX19", "PEX2", "PEX26", "PEX5", "PEX6", "PFDN2", "PFDN4", 
"PFDN5", "PFDN6", "PFN1", "PGAM5", "PGBD3", "PGK1", "PGLS", "PGP", 
"PGPEP1", "PGRMC2", "PHACTR4", "PHAX", "PHB", "PHB2", "PHC2", 
"PHF10", "PHF12", "PHF20L1", "PHF23", "PHF5A", "PHKB", "PHPT1", 
"PHRF1", "PI4K2A", "PI4KA", "PI4KB", "PIAS1", "PICALM", "PICK1", 
"PIGC", "PIGF", "PIGG", "PIGH", "PIGK", "PIGP", "PIGS", "PIGT", 
"PIGU", "PIGW", "PIGX", "PIGY", "PIH1D1", "PIK3C3", "PIK3CB", 
"PIK3R1", "PIK3R4", "PIN1", "PINK1", "PINX1", "PIP5K1A", "PITHD1", 
"PITPNA", "PITPNB", "PITRM1", "PLA2G12A", "PLAA", "PLBD2", "PLD3", 
"PLEKHA1", "PLEKHJ1", "PLEKHM1", "PLGRKT", "PLIN3", "PLOD1", 
"PLOD3", "PLRG1", "PMF1", "PMF1-BGLAP", "PMPCA", "PMPCB", "PMS1", 
"PMVK", "PNISR", "PNKD", "PNKP", "PNN", "PNO1", "PNPLA6", "PNPLA8", 
"PNPO", "PNPT1", "PNRC2", "POFUT1", "POLD2", "POLDIP2", "POLDIP3", 
"POLE3", "POLE4", "POLG", "POLH", "POLK", "POLL", "POLM", "POLR1C", 
"POLR1D", "POLR1E", "POLR2A", "POLR2B", "POLR2C", "POLR2D", "POLR2E", 
"POLR2F", "POLR2G", "POLR2H", "POLR2I", "POLR2J", "POLR2K", "POLR2L", 
"POLR3C", "POLR3E", "POLR3GL", "POLR3K", "POM121", "POM121C", 
"POMGNT1", "POMP", "POMT1", "POP4", "POP5", "POP7", "PPA1", "PPA2", 
"PPAN", "PPAN-P2RY11", "PPARA", "PPARD", "PPCS", "PPFIA1", "PPHLN1", 
"PPID", "PPIE", "PPIF", "PPIG", "PPIH", "PPIL4", "PPM1A", "PPM1B", 
"PPP1CA", "PPP1CC", "PPP1R10", "PPP1R11", "PPP1R15B", "PPP1R37", 
"PPP1R7", "PPP1R8", "PPP2CA", "PPP2CB", "PPP2R1A", "PPP2R2A", 
"PPP2R2D", "PPP2R3C", "PPP2R4", "PPP2R5A", "PPP2R5B", "PPP2R5C", 
"PPP2R5D", "PPP2R5E", "PPP4C", "PPP4R1", "PPP4R2", "PPP5C", "PPP6C", 
"PPP6R2", "PPP6R3", "PPWD1", "PQBP1", "PQLC1", "PQLC2", "PRADC1", 
"PRCC", "PRDM4", "PRDX1", "PRDX2", "PRDX3", "PRDX5", "PRDX6", 
"PREB", "PREP", "PRKAA1", "PRKAB1", "PRKACA", "PRKAG1", "PRKAR1A", 
"PRKRIP1", "PRMT1", "PRMT5", "PRMT7", "PROSC", "PRPF18", "PRPF19", 
"PRPF3", "PRPF31", "PRPF4", "PRPF40A", "PRPF4B", "PRPF6", "PRPF8", 
"PRPS1", "PRPSAP1", "PRR14", "PRRC1", "PRRC2A", "PRRC2B", "PRUNE", 
"PSEN1", "PSEN2", "PSENEN", "PSKH1", "PSMA1", "PSMA2", "PSMA3", 
"PSMA4", "PSMA5", "PSMA6", "PSMA7", "PSMB1", "PSMB2", "PSMB3", 
"PSMB4", "PSMB5", "PSMB6", "PSMB7", "PSMC2", "PSMC3", "PSMC4", 
"PSMC5", "PSMC6", "PSMD1", "PSMD10", "PSMD11", "PSMD12", "PSMD13", 
"PSMD14", "PSMD2", "PSMD3", "PSMD4", "PSMD5", "PSMD6", "PSMD7", 
"PSMD8", "PSMD9", "PSME1", "PSME3", "PSMF1", "PSMG2", "PSMG3", 
"PSMG4", "PSPC1", "PTCD1", "PTCD3", "PTDSS1", "PTEN", "PTGES2", 
"PTGES3", "PTOV1", "PTP4A2", "PTPMT1", "PTPN1", "PTPN11", "PTPN23", 
"PTRH1", "PTRH2", "PTRHD1", "PUF60", "PUM1", "PUM2", "PURA", 
"PURB", "PUS3", "PUS7", "PUSL1", "PWP1", "PWP2", "PWWP2A", "PXMP4", 
"PYCR2", "PYGO2", "PYURF", "QARS", "QRICH1", "QRSL1", "QSOX1", 
"QTRT1", "R3HCC1", "R3HDM2", "RAB10", "RAB11A", "RAB11B", "RAB14", 
"RAB18", "RAB1A", "RAB1B", "RAB21", "RAB22A", "RAB2A", "RAB2B", 
"RAB3GAP1", "RAB3GAP2", "RAB40C", "RAB4A", "RAB5A", "RAB5B", 
"RAB5C", "RAB6A", "RAB7A", "RAB9A", "RABEP1", "RABEPK", "RABGEF1", 
"RABGGTA", "RABGGTB", "RAD1", "RAD17", "RAD23B", "RAD50", "RAD51C", 
"RAF1", "RALA", "RALBP1", "RALY", "RAN", "RANBP1", "RANBP2", 
"RANBP3", "RANBP6", "RANGAP1", "RANGRF", "RAP1A", "RAPGEF1", 
"RAPGEF2", "RARS", "RARS2", "RB1CC1", "RBAK", "RBBP4", "RBBP7", 
"RBCK1", "RBFA", "RBM10", "RBM12", "RBM12B", "RBM14", "RBM14-RBM4", 
"RBM15", "RBM15B", "RBM17", "RBM18", "RBM19", "RBM23", "RBM27", 
"RBM28", "RBM33", "RBM34", "RBM39", "RBM4", "RBM41", "RBM42", 
"RBM5", "RBM6", "RBM7", "RBM8A", "RBMX", "RBMXL1", "RBX1", "RC3H2", 
"RCAN1", "RCHY1", "RCN2", "RDH14", "RDX", "REEP3", "REEP5", "RELA", 
"REPIN1", "REPS1", "RER1", "REST", "REXO1", "RFC1", "RFC2", "RFC5", 
"RFK", "RFNG", "RFT1", "RFWD2", "RFXANK", "RGP1", "RHBDD1", "RHBDD3", 
"RHOA", "RHOB", "RHOT1", "RHOT2", "RIC8A", "RIN2", "RING1", "RINT1", 
"RIOK1", "RIOK2", "RIOK3", "RIPK1", "RMDN1", "RMDN3", "RMI1", 
"RMND1", "RMND5A", "RMND5B", "RNASEH1", "RNASEH2C", "RNASEK", 
"RNF10", "RNF103", "RNF11", "RNF111", "RNF113A", "RNF115", "RNF121", 
"RNF126", "RNF13", "RNF14", "RNF141", "RNF146", "RNF167", "RNF181", 
"RNF185", "RNF187", "RNF216", "RNF220", "RNF25", "RNF26", "RNF31", 
"RNF34", "RNF4", "RNF40", "RNF5", "RNF6", "RNF7", "RNH1", "RNMTL1", 
"RNPEP", "ROMO1", "RP9", "RPA2", "RPA3", "RPAIN", "RPAP3", "RPF1", 
"RPF2", "RPL10A", "RPL11", "RPL14", "RPL26L1", "RPL27", "RPL30", 
"RPL31", "RPL32", "RPL34", "RPL35", "RPL35A", "RPL36AL", "RPL4", 
"RPL6", "RPL7L1", "RPL8", "RPN1", "RPN2", "RPP14", "RPP25L", 
"RPP30", "RPP38", "RPRD1B", "RPS13", "RPS19BP1", "RPS23", "RPS24", 
"RPS27L", "RPS5", "RPS6", "RPS6KA3", "RPS6KB1", "RPS6KB2", "RPUSD3", 
"RQCD1", "RRAGA", "RRM1", "RRN3", "RRNAD1", "RRP1", "RRP36", 
"RRP7A", "RRP8", "RRS1", "RSAD1", "RSBN1L", "RSC1A1", "RSL1D1", 
"RSPRY1", "RSRC1", "RSRC2", "RTCA", "RTFDC1", "RTN4", "RUFY1", 
"RUVBL1", "RWDD1", "RWDD3", "RXRA", "RXRB", "SAE1", "SAMD1", 
"SAMD4B", "SAMD8", "SAMM50", "SAP18", "SAP30", "SAP30BP", "SAP30L", 
"SAR1A", "SARNP", "SARS", "SART1", "SART3", "SAT2", "SAV1", "SBDS", 
"SCAF1", "SCAF11", "SCAF4", "SCAF8", "SCAMP2", "SCAMP3", "SCAND1", 
"SCAP", "SCARB2", "SCFD1", "SCFD2", "SCNM1", "SCO1", "SCO2", 
"SCOC", "SCP2", "SCRIB", "SCRN3", "SCYL1", "SCYL2", "SCYL3", 
"SDAD1", "SDCBP", "SDCCAG3", "SDCCAG8", "SDE2", "SDF2", "SDF4", 
"SDHA", "SDHAF2", "SDHB", "SDHC", "SDHD", "SDR39U1", "SEC11A", 
"SEC13", "SEC16A", "SEC22B", "SEC22C", "SEC23A", "SEC23IP", "SEC24A", 
"SEC24B", "SEC24C", "SEC31A", "SEC61A1", "SEC61B", "SEC61G", 
"SEC62", "SEC63", "SECISBP2", "SEH1L", "SEL1L", "SELK", "SELO", 
"SELRC1", "SELT", "SENP2", "SENP3", "SENP5", "SENP6", "SEPHS1", 
"SERBP1", "SERF2", "SERGEF", "SERINC1", "SERINC3", "SERPINB6", 
"SERTAD2", "SET", "SETD2", "SETD3", "SETD5", "SETD6", "SETD7", 
"SETD8", "SETDB1", "SF1", "SF3A1", "SF3A3", "SF3B1", "SF3B14", 
"SF3B2", "SF3B3", "SF3B4", "SF3B5", "SFSWAP", "SGK196", "SGMS1", 
"SGPL1", "SGSM3", "SGTA", "SH3BP5L", "SH3GLB1", "SHARPIN", "SHOC2", 
"SIAH1", "SIAH2", "SIGMAR1", "SIKE1", "SIL1", "SIRT2", "SIRT3", 
"SIRT5", "SIRT6", "SIVA1", "SKIL", "SKIV2L", "SKIV2L2", "SKP1", 
"SLC15A4", "SLC20A1", "SLC25A11", "SLC25A26", "SLC25A28", "SLC25A3", 
"SLC25A32", "SLC25A38", "SLC25A39", "SLC25A44", "SLC25A46", "SLC25A5", 
"SLC27A4", "SLC30A1", "SLC30A5", "SLC30A9", "SLC35A2", "SLC35A4", 
"SLC35B1", "SLC35B2", "SLC35C2", "SLC35E1", "SLC35E3", "SLC35F5", 
"SLC38A2", "SLC39A1", "SLC39A3", "SLC39A7", "SLC41A3", "SLC46A3", 
"SLC48A1", "SLIRP", "SLMO2", "SLTM", "SMAD2", "SMAD4", "SMAD5", 
"SMAP1", "SMARCA2", "SMARCA4", "SMARCAL1", "SMARCB1", "SMARCE1", 
"SMC1A", "SMC5", "SMCR7L", "SMEK1", "SMEK2", "SMG5", "SMG7", 
"SMG8", "SMIM11", "SMIM12", "SMIM8", "SMNDC1", "SMPD1", "SMPD4", 
"SMU1", "SMUG1", "SNAP23", "SNAP29", "SNAP47", "SNAPC3", "SNAPC5", 
"SNAPIN", "SND1", "SNF8", "SNRNP200", "SNRNP25", "SNRNP27", "SNRNP35", 
"SNRNP40", "SNRNP48", "SNRNP70", "SNRPA", "SNRPB", "SNRPB2", 
"SNRPC", "SNRPD1", "SNRPD2", "SNRPD3", "SNRPG", "SNUPN", "SNW1", 
"SNX12", "SNX13", "SNX17", "SNX18", "SNX19", "SNX2", "SNX25", 
"SNX3", "SNX4", "SNX5", "SNX6", "SNX9", "SOCS4", "SOCS6", "SOD1", 
"SON", "SPAG7", "SPAG9", "SPATA2", "SPATA5L1", "SPCS1", "SPCS3", 
"SPECC1L", "SPEN", "SPG11", "SPG21", "SPG7", "SPHAR", "SPNS1", 
"SPOP", "SPPL2B", "SPPL3", "SPRYD3", "SPRYD7", "SPSB3", "SPTSSA", 
"SPTY2D1", "SRA1", "SRD5A3", "SREBF2", "SREK1IP1", "SRM", "SRP14", 
"SRP19", "SRP54", "SRP68", "SRP72", "SRP9", "SRPR", "SRPRB", 
"SRR", "SRRD", "SRRM1", "SRSF1", "SRSF10", "SRSF11", "SRSF2", 
"SRSF3", "SRSF4", "SRSF7", "SRSF8", "SS18L2", "SSB", "SSBP1", 
"SSNA1", "SSR1", "SSR2", "SSR3", "SSRP1", "SSSCA1", "SSU72", 
"ST3GAL2", "ST6GALNAC6", "ST7", "STAM", "STAM2", "STAMBP", "STARD3", 
"STARD7", "STAT3", "STAU1", "STAU2", "STIM1", "STIP1", "STK11", 
"STK16", "STOM", "STOML1", "STOML2", "STRAP", "STRIP1", "STRN3", 
"STT3A", "STT3B", "STUB1", "STX10", "STX17", "STX4", "STX5", 
"STX8", "STXBP3", "STYXL1", "SUB1", "SUCLA2", "SUCLG1", "SUCLG2", 
"SUGP1", "SUGT1", "SUMO1", "SUMO3", "SUN2", "SUPT4H1", "SUPT5H", 
"SUPT6H", "SUPT7L", "SUPV3L1", "SURF1", "SURF4", "SURF6", "SUV420H1", 
"SUZ12", "SYAP1", "SYF2", "SYMPK", "SYNCRIP", "SYNJ2BP", "SYNJ2BP-COX16", 
"SYPL1", "SYS1", "SYVN1", "SZRD1", "TAB1", "TAB2", "TACO1", "TADA1", 
"TADA3", "TAF10", "TAF11", "TAF12", "TAF13", "TAF15", "TAF1D", 
"TAF4", "TAF5L", "TAF8", "TAF9", "TALDO1", "TAMM41", "TANGO2", 
"TANGO6", "TANK", "TAOK2", "TAPBP", "TAPT1", "TARDBP", "TARS", 
"TATDN1", "TATDN2", "TAX1BP1", "TAZ", "TBC1D1", "TBC1D14", "TBC1D15", 
"TBC1D20", "TBC1D22A", "TBC1D23", "TBC1D7", "TBC1D9B", "TBCA", 
"TBCB", "TBCC", "TBCCD1", "TBCD", "TBCE", "TBK1", "TBRG1", "TBRG4", 
"TCAIM", "TCEANC2", "TCEB1", "TCEB2", "TCEB3", "TCERG1", "TCF12", 
"TCF20", "TCF25", "TCP1", "TCTN3", "TDP2", "TDRD3", "TECR", "TEF", 
"TEFM", "TELO2", "TERF2", "TERF2IP", "TEX2", "TEX261", "TEX264", 
"TFAM", "TFB1M", "TFB2M", "TFCP2", "TFDP1", "TFE3", "TFG", "TFIP11", 
"TFPT", "TGIF2-C20orf24", "TGOLN2", "THADA", "THAP3", "THAP4", 
"THAP5", "THAP7", "THOC5", "THOC7", "THOP1", "THRAP3", "THTPA", 
"THUMPD3", "THYN1", "TIA1", "TIAL1", "TICAM1", "TIGD5", "TIGD6", 
"TIMM10", "TIMM10B", "TIMM13", "TIMM17A", "TIMM17B", "TIMM21", 
"TIMM22", "TIMM44", "TIMM50", "TIMM8B", "TIMM9", "TIMMDC1", "TINF2", 
"TIPRL", "TJAP1", "TLE1", "TLK1", "TM2D1", "TM2D2", "TM2D3", 
"TM9SF1", "TM9SF2", "TM9SF3", "TM9SF4", "TMBIM1", "TMBIM4", "TMBIM6", 
"TMCC1", "TMCO1", "TMCO3", "TMED1", "TMED10", "TMED2", "TMED4", 
"TMED5", "TMED7", "TMED7-TICAM2", "TMED9", "TMEM101", "TMEM106B", 
"TMEM106C", "TMEM115", "TMEM120A", "TMEM126A", "TMEM127", "TMEM128", 
"TMEM129", "TMEM131", "TMEM134", "TMEM141", "TMEM147", "TMEM14B", 
"TMEM14C", "TMEM161A", "TMEM167B", "TMEM168", "TMEM177", "TMEM179B", 
"TMEM18", "TMEM184C", "TMEM185B", "TMEM186", "TMEM187", "TMEM189", 
"TMEM189-UBE2V1", "TMEM19", "TMEM192", "TMEM199", "TMEM203", 
"TMEM205", "TMEM214", "TMEM219", "TMEM222", "TMEM223", "TMEM230", 
"TMEM242", "TMEM248", "TMEM251", "TMEM256", "TMEM258", "TMEM259", 
"TMEM30A", "TMEM33", "TMEM39A", "TMEM41A", "TMEM41B", "TMEM42", 
"TMEM5", "TMEM50A", "TMEM50B", "TMEM55B", "TMEM57", "TMEM59", 
"TMEM60", "TMEM62", "TMEM63B", "TMEM64", "TMEM66", "TMEM69", 
"TMEM70", "TMEM81", "TMEM87A", "TMEM9", "TMEM9B", "TMF1", "TMLHE", 
"TMPO", "TMUB1", "TMUB2", "TMX1", "TMX2", "TMX4", "TNFAIP1", 
"TNFAIP8L2-SCNM1", "TNIP1", "TNKS2", "TNPO1", "TNPO3", "TNRC6A", 
"TOB1", "TOLLIP", "TOMM20", "TOMM22", "TOMM40", "TOMM5", "TOMM6", 
"TOMM7", "TOMM70A", "TOP1", "TOP2B", "TOPORS", "TOR1A", "TOR1AIP2", 
"TOR1B", "TOR3A", "TOX4", "TP53RK", "TPCN1", "TPD52L2", "TPGS1", 
"TPI1", "TPP2", "TPRA1", "TPRG1L", "TPRKB", "TPRN", "TPST2", 
"TRA2A", "TRA2B", "TRAF6", "TRAF7", "TRAP1", "TRAPPC1", "TRAPPC10", 
"TRAPPC11", "TRAPPC12", "TRAPPC13", "TRAPPC2L", "TRAPPC3", "TRAPPC4", 
"TRAPPC5", "TRAPPC6B", "TRAPPC8", "TRAPPC9", "TRIAP1", "TRIM26", 
"TRIM27", "TRIM28", "TRIM3", "TRIM39", "TRIM39-RPP21", "TRIM41", 
"TRIM44", "TRIM56", "TRIM65", "TRIM8", "TRIP12", "TRIP4", "TRMT1", 
"TRMT10C", "TRMT112", "TRMT12", "TRMT1L", "TRMT2A", "TRNAU1AP", 
"TRNT1", "TRPC4AP", "TRPT1", "TRUB2", "TSC2", "TSEN15", "TSEN34", 
"TSFM", "TSG101", "TSN", "TSNAX", "TSPAN17", "TSPAN31", "TSPYL1", 
"TSR1", "TSR2", "TSR3", "TSSC4", "TSTA3", "TSTD2", "TTC1", "TTC17", 
"TTC19", "TTC32", "TTC33", "TTC37", "TTC4", "TTC7B", "TTC9C", 
"TTI1", "TTI2", "TUBA1B", "TUBA1C", "TUBB", "TUBD1", "TUBGCP2", 
"TUBGCP4", "TUFM", "TUSC2", "TUT1", "TVP23B", "TXLNA", "TXLNG", 
"TXN2", "TXNDC11", "TXNDC12", "TXNDC15", "TXNDC17", "TXNDC9", 
"TXNL1", "TXNL4A", "TXNL4B", "TXNRD1", "TYK2", "TYW1", "U2AF1", 
"U2AF1L4", "U2AF2", "UAP1", "UBA1", "UBA2", "UBA3", "UBA5", "UBA52", 
"UBAC2", "UBALD1", "UBAP1", "UBAP2L", "UBB", "UBC", "UBE2A", 
"UBE2B", "UBE2D2", "UBE2D3", "UBE2D4", "UBE2E1", "UBE2E2", "UBE2E3", 
"UBE2F", "UBE2G2", "UBE2H", "UBE2I", "UBE2J1", "UBE2J2", "UBE2K", 
"UBE2L3", "UBE2M", "UBE2N", "UBE2NL", "UBE2Q1", "UBE2R2", "UBE2V1", 
"UBE2V2", "UBE2W", "UBE2Z", "UBE3A", "UBE3B", "UBE3C", "UBE4A", 
"UBE4B", "UBFD1", "UBIAD1", "UBL3", "UBL4A", "UBL5", "UBL7", 
"UBOX5", "UBP1", "UBQLN1", "UBQLN2", "UBQLN4", "UBR2", "UBR7", 
"UBTD1", "UBTF", "UBXN2A", "UBXN4", "UBXN6", "UCHL3", "UCHL5", 
"UCK1", "UCK2", "UCKL1", "UEVLD", "UFC1", "UFD1L", "UFL1", "UFSP2", 
"UGP2", "UHRF1BP1L", "ULK1", "ULK3", "UNC50", "UNG", "UPF1", 
"UPF2", "UPF3B", "UPRT", "UQCC", "UQCR10", "UQCR11", "UQCRB", 
"UQCRC1", "UQCRC2", "UQCRHL", "UQCRQ", "URGCP", "URI1", "URM1", 
"UROD", "UROS", "USB1", "USE1", "USF1", "USF2", "USP10", "USP14", 
"USP16", "USP19", "USP22", "USP25", "USP27X", "USP33", "USP38", 
"USP39", "USP4", "USP47", "USP5", "USP7", "USP8", "USP9X", "UTP11L", 
"UTP14A", "UTP14C", "UTP15", "UTP23", "UTP3", "UTP6", "UXS1", 
"UXT", "VAC14", "VAMP3", "VAMP5", "VAPA", "VAPB", "VARS2", "VBP1", 
"VCP", "VDAC3", "VEZT", "VIMP", "VMA21", "VPS16", "VPS18", "VPS25", 
"VPS26A", "VPS26B", "VPS28", "VPS29", "VPS33A", "VPS36", "VPS37A", 
"VPS4A", "VPS51", "VPS52", "VPS53", "VPS72", "VRK2", "VRK3", 
"VTA1", "VTI1A", "VTI1B", "WAC", "WAPAL", "WARS2", "WBP11", "WBP1L", 
"WBP2", "WBP4", "WBSCR22", "WDR1", "WDR12", "WDR13", "WDR18", 
"WDR20", "WDR24", "WDR25", "WDR26", "WDR3", "WDR33", "WDR36", 
"WDR41", "WDR43", "WDR44", "WDR45", "WDR45B", "WDR46", "WDR55", 
"WDR59", "WDR5B", "WDR6", "WDR61", "WDR70", "WDR73", "WDR74", 
"WDR75", "WDR77", "WDR81", "WDR83OS", "WDR85", "WDR89", "WDTC1", 
"WIBG", "WIPI2", "WIZ", "WRAP53", "WRB", "WRNIP1", "WSB2", "WTAP", 
"WTH3DI", "WWP1", "WWP2", "XIAP", "XPA", "XPC", "XPNPEP1", "XPO1", 
"XPO7", "XPOT", "XRCC5", "XRCC6", "XYLT2", "YAF2", "YARS", "YARS2", 
"YIF1A", "YIF1B", "YIPF1", "YIPF3", "YIPF4", "YIPF5", "YIPF6", 
"YKT6", "YME1L1", "YPEL2", "YRDC", "YTHDC1", "YTHDF1", "YTHDF2", 
"YTHDF3", "YWHAB", "YWHAE", "YY1", "YY1AP1", "ZADH2", "ZBED4", 
"ZBED6", "ZBTB1", "ZBTB10", "ZBTB11", "ZBTB14", "ZBTB17", "ZBTB18", 
"ZBTB21", "ZBTB25", "ZBTB33", "ZBTB39", "ZBTB44", "ZBTB45", "ZBTB5", 
"ZBTB6", "ZBTB7A", "ZBTB8OS", "ZC3H10", "ZC3H11A", "ZC3H13", 
"ZC3H15", "ZC3H18", "ZC3H3", "ZC3H7A", "ZC3H7B", "ZCCHC10", "ZCCHC11", 
"ZCCHC3", "ZCCHC7", "ZCCHC9", "ZCRB1", "ZDHHC14", "ZDHHC16", 
"ZDHHC2", "ZDHHC3", "ZDHHC4", "ZDHHC5", "ZDHHC8", "ZFAND1", "ZFAND2B", 
"ZFAND3", "ZFAND5", "ZFAND6", "ZFP91", "ZFPL1", "ZFR", "ZFYVE1", 
"ZFYVE19", "ZFYVE27", "ZGPAT", "ZHX1", "ZHX1-C8ORF76", "ZHX2", 
"ZHX3", "ZKSCAN1", "ZMAT2", "ZMAT3", "ZMAT5", "ZMPSTE24", "ZMYM2", 
"ZMYND11", "ZNF121", "ZNF131", "ZNF134", "ZNF138", "ZNF142", 
"ZNF143", "ZNF146", "ZNF174", "ZNF181", "ZNF189", "ZNF195", "ZNF197", 
"ZNF207", "ZNF22", "ZNF226", "ZNF232", "ZNF24", "ZNF259", "ZNF274", 
"ZNF277", "ZNF280D", "ZNF281", "ZNF3", "ZNF32", "ZNF322", "ZNF326", 
"ZNF330", "ZNF335", "ZNF33A", "ZNF343", "ZNF347", "ZNF37A", "ZNF384", 
"ZNF394", "ZNF397", "ZNF398", "ZNF408", "ZNF41", "ZNF410", "ZNF414", 
"ZNF419", "ZNF438", "ZNF444", "ZNF446", "ZNF48", "ZNF480", "ZNF491", 
"ZNF506", "ZNF507", "ZNF513", "ZNF518A", "ZNF526", "ZNF561", 
"ZNF574", "ZNF576", "ZNF579", "ZNF580", "ZNF592", "ZNF593", "ZNF598", 
"ZNF620", "ZNF622", "ZNF623", "ZNF638", "ZNF639", "ZNF641", "ZNF644", 
"ZNF649", "ZNF654", "ZNF655", "ZNF664", "ZNF668", "ZNF672", "ZNF687", 
"ZNF688", "ZNF691", "ZNF7", "ZNF706", "ZNF721", "ZNF740", "ZNF76", 
"ZNF764", "ZNF770", "ZNF777", "ZNF787", "ZNF805", "ZNF814", "ZNF830", 
"ZNF865", "ZNF91", "ZNHIT1", "ZNHIT3", "ZNRD1", "ZRANB1", "ZRANB2", 
"ZSCAN21", "ZSCAN29", "ZSCAN32", "ZSWIM1", "ZSWIM7", "ZSWIM8", 
"ZW10", "ZXDA", "ZXDB", "ZZZ3")
#Taken from here Table S2
#http://journals.plos.org/plosone/article?id=10.1371/journal.pone.0012862
nuclearMTGenes = c("ABCB10", "ABCD3", "ACADM", "ACP6", "AGMAT", "AK2", "AKR7A2",
"ALDH4A1", "ALDH9A1", "APOA1BP", "ATAD3A", "ATAD3B", "ATP5F1",
"ATPAF1", "ATPIF1", "AURKAIP1", "BOLA1", "BRP44", "C1orf151",
"C1orf31", "C1orf69", "CABC1", "CCBL2", "CLIC4", "CPT2", "CYB5R1",
"DAP3", "DARS2", "DBT", "DDAH1", "DNAJC11", "FAM36A", "FDPS",
"FH", "GLRX2", "GNG5", "HAO2", "HBXIP", "HMGCL", "HMGCS2", "HSD3B1",
"HSPB7", "IARS2", "KIF1B", "MECR", "MFN2", "MMACHC", "MOSC1",
"MOSC2", "MRPL20", "MRPL24", "MRPL37", "MRPL55", "MRPL9", "MRPS14",
"MRPS15", "MRPS21", "MTX1", "MUL1", "MUTYH", "NDUFS2", "NDUFS5",
"NIT1", "NRD1", "OMA1", "PARK7", "PARS2", "PEX11B", "PINK1",
"PPOX", "PRDX1", "PRDX6", "SARS", "SCCPDH", "SCP2", "SDHB", "SDHC",
"SLC16A1", "SLC25A24", "SLC25A33", "SLC25A34", "SLC25A44", "TARS2",
"TATDN3", "TDRKH", "TFB2M", "THEM4", "THEM5", "TIMM17A", "TOMM20",
"TOMM40L", "TRIT1", "UQCRH", "WARS2", "ABCB6", "ACADL", "ACYP2",
"AGPS", "AGXT", "ATIC", "ATP5G3", "BCS1L", "C2orf47", "C2orf56",
"CASP8", "CCT7", "COQ10B", "COX5B", "COX7A2L", "CPS1", "CYP27A1",
"DGUOK", "DNAJC27", "EFHD1", "FAM136A", "FASTKD2", "FSIP2", "GLS",
"GPD2", "HADHA", "HADHB", "HIBCH", "HK2", "HSPD1", "HSPE1", "HTRA2",
"IDH1", "IMMT", "KYNU", "LIPT1", "LRPPRC", "MAP1D", "MARS2",
"MCEE", "MDH1", "MPV17", "MRPL19", "MRPL30", "MRPL33", "MRPL35",
"MRPL44", "MRPS9", "MTHFD2", "MTIF2", "MTX2", "NDUFA10", "NDUFB3",
"NDUFS1", "NFU1", "NIF3L1", "OSGEPL1", "PDK1", "PNKD", "PNPT1",
"PTCD3", "RAB11FIP5", "RDH14", "RNASEH1", "SFXN5", "SLC25A12",
"SLC30A6", "SPR", "SUCLG1", "UXS1", "VAMP8", "ABHD10", "ACAA1",
"ACAD11", "ACAD9", "ALAS1", "ALDH1L1", "AMT", "BDH1", "C3orf1",
"C3orf23", "C3orf31", "C3orf60", "CCDC51", "CCDC58", "CHCHD4",
"CHDH", "CMC1", "COX17", "CPOX", "EEFSEC", "EHHADH", "EXOG",
"FAM162A", "GFM1", "GMPPB", "GPX1", "GTPBP8", "HEMK1", "HIGD1A",
"LARS2", "MCCC1", "MFN1", "MOBP", "MRPL3", "MRPL47", "MRPS22",
"MRPS25", "NDUFB4", "NDUFB5", "NME6", "OGG1", "OPA1", "OXNAD1",
"OXSM", "PARL", "PCCB", "PDE12", "PDHB", "RG9MTD1", "RHOA", "RPL35A",
"SLC25A20", "SLC25A26", "SLC25A36", "SUCLG2", "TOMM70A", "TRNT1",
"UQCRC1", "AADAT", "ACOX3", "ACSL1", "ATP10D", "ATP5I", "C4orf14",
"CBR4", "COQ2", "COX18", "ETFDH", "GK2", "GRPEL1", "GRSF1", "GUF1",
"HADH", "LAP3", "LETM1", "LIAS", "MRPL1", "MRPS18C", "MTHFD2L",
"NFXL1", "OCIAD1", "PDHA2", "PET112L", "PIGY", "PPA2", "PPM1K",
"QDPR", "RPL34", "SLC25A31", "SLC25A4", "UCP1", "AGXT2", "AGXT2L2",
"ALDH7A1", "AMACR", "CKMT2", "DHX29", "DMGDH", "FTMT", "GFM2",
"GLRX", "GRPEL2", "HARS2", "HIGD2A", "HK3", "HSD17B4", "HSPA9",
"ISCA1L", "KIAA0141", "LOC153328", "LOC727947", "LYRM7", "MAT2B",
"MCCC2", "MRPL22", "MRPL36", "MRPS27", "MRPS30", "MRPS36", "NDUFA2",
"NDUFAF2", "NDUFS4", "NDUFS6", "NLN", "NNT", "OXCT1", "PCBD2",
"PRELID1", "PRELID2", "PTCD2", "RAB24", "RARS", "RPS14", "SDHA",
"SFXN1", "SLC22A4", "SLC25A2", "SLC25A46", "THG1L", "UQCRQ",
"VDAC1", "AARS2", "ACAT2", "ALDH5A1", "BAK1", "BCKDHB", "BPHL",
"BRP44L", "C6orf125", "C6orf203", "C6orf57", "C6orf66", "CCDC90A",
"COQ3", "COX7A2", "DACT2", "FARS2", "HDDC2", "HSD17B8", "LACE1",
"LOC643233", "LYRM4", "ME1", "MRPL14", "MRPL18", "MRPL2", "MRPS10",
"MRPS18A", "MRPS18B", "MRS2", "MTCH1", "MTHFD1L", "MTO1", "MTRF1L",
"MUT", "PACRG", "PDSS2", "PECI", "PRICKLE4", "PRSS35", "QRSL1",
"RAB32", "RARS2", "RMND1", "RPL10A", "RTN4IP1", "SIRT5", "SLC25A27",
"SOD2", "TAP1", "TFB1M", "THEM2", "TMEM14C", "VARS2", "AASS",
"ABCB8", "ABCF2", "ABHD11", "ACN9", "ADCK2", "AGK", "AGR2", "ARMC10",
"ATP5J2", "C7orf10", "C7orf55", "CCDC136", "CHCHD2", "CHCHD3",
"CROT", "CYCS", "DLD", "DNAJC30", "ELN", "FASTK", "FIS1", "FTSJ2",
"GARS", "GBAS", "HIBADH", "IMMP2L", "IQCE", "MDH2", "MRPL32",
"MRPS17", "MRPS24", "MRPS33", "MTERF", "MYLC2PL", "NDUFA4", "NDUFA5",
"NDUFB2", "NT5C3", "OGDH", "PDK4", "PMPCB", "SLC25A13", "SLC25A40",
"SND1", "SSBP1", "TBRG4", "tcag7.1260", "TOMM7", "ADCK5", "ADHFE1",
"AGPAT5", "ARMC1", "C8orf38", "CHCHD7", "COX6C", "CYC1", "CYP11B1",
"CYP11B2", "DECR1", "DUSP26", "FAM82B", "GSR", "HRSP12", "LACTB2",
"MRPL13", "MRPL15", "MRPS28", "MSRA", "MTERFD1", "MTFR1", "NDUFB9",
"OXR1", "PPM2C", "SLC25A32", "SLC25A37", "STAR", "TMEM65", "TMEM70",
"TOP1MT", "TTC35", "UQCRB", "VDAC3", "ACO1", "AK3", "ALDH1B1",
"AUH", "C9orf46", "COQ4", "CRAT", "ENDOG", "FPGS", "FXN", "GLDC",
"HINT2", "HSDL2", "ISCA1", "MCART1", "MRPL41", "MRPL50", "MRPS2",
"MRRF", "NDUFA8", "NDUFB6", "NIPSNAP3A", "NIPSNAP3B", "NUDT2",
"PMPCA", "PTGES2", "PTRH1", "RFK", "SARDH", "SECISBP2", "SLC25A25",
"STOML2", "SURF1", "TXN", "ACADSB", "ACSL5", "ADO", "AIFM2",
"ALDH18A1", "ANKRD26", "ARMC4", "AS3MT", "ASAH2", "ATAD1", "ATP5C1",
"C10orf10", "C10orf2", "C10orf58", "C10orf65", "CBARA1", "CCDC109A",
"CHAT", "CHCHD1", "CISD1", "COMTD1", "COX15", "DHTKD1", "ECHDC3",
"ECHS1", "GHITM", "GLUD1", "GPAM", "HK1", "IDE", "IDI1", "MRPL43",
"MRPS16", "MSRB2", "MTG1", "MTPAP", "NCOA4", "NUDT13", "OAT",
"OGDHL", "PDSS1", "PHYH", "PHYHIPL", "PITRM1", "PPIF", "PRDX3",
"PSTK", "SFXN2", "SFXN3", "SFXN4", "SLC25A16", "SLC25A28", "SUPV3L1",
"TFAM", "VDAC2", "YME1L1", "ACAD8", "ACAT1", "ATP5L", "BAD",
"BBOX1", "C11orf79", "CCDC90B", "CLPB", "COX8A", "CPT1A", "CYB5R2",
"DLAT", "DNAJC4", "FDX1", "FLJ14213", "FOXRED1", "FTH1", "FXC1",
"GLYAT", "LDHA", "LOC387787", "ME3", "MRPL16", "MRPL17", "MRPL21",
"MRPL23", "MRPL48", "MRPL49", "MTCH2", "NARS2", "NDUFC2", "NDUFS3",
"NDUFS8", "NDUFV1", "NLRX1", "NUDT8", "PC", "PDHX", "PRDX5",
"PTS", "REXO2", "RPUSD4", "SDHD", "SIRT3", "TCIRG1", "TIMM10",
"TIMM8B", "TMEM126A", "TMEM223", "UCP2", "UCP3", "ZBED5", "ABCD2",
"ACACB", "ACAD10", "ACADS", "ACSM4", "ALDH1L2", "ALDH2", "ATP5G2",
"BCAT1", "BLOC1S1", "C12orf62", "COQ10A", "COQ5", "COX6A1", "CRY1",
"CS", "CYP27B1", "DIABLO", "GATC", "GLS2", "ISCU", "KRT5", "LDHB",
"LETMD1", "LYRM5", "MARS", "MMAB", "MRPL42", "MRPL51", "MRPS35",
"MTERFD3", "NDUFA12", "NDUFA9", "PHB2", "PPTC7", "RAB35", "SDSL",
"SHMT2", "SLC25A3", "SPRYD4", "SUOX", "TMTC1", "TRIAP1", "TSFM",
"TXNRD1", "UNG", "VAMP1", "YARS2", "CARKD", "CARS2", "CLYBL",
"DNAJC15", "EFHA1", "hCG_1790806", "MIPEP", "MRP63", "MRPS31",
"MTIF3", "MTRF1", "PCCA", "SLC25A15", "SLC25A30", "SUCLA2", "ACOT2",
"ADCK1", "ALDH6A1", "ARG2", "ATP5S", "C14orf153", "C14orf156",
"C14orf159", "C14orf68", "COQ6", "COX16", "COX8C", "DHRS1", "DHRS4",
"DLST", "GALC", "GLRX5", "ISCA2", "L2HGDH", "METT11D1", "MRPL52",
"MTHFD1", "NUBPL", "OXA1L", "PCK2", "PTGR2", "SLC25A21", "SLC25A29",
"SPTLC2", "TIMM9", "WDR22", "C15orf48", "CKMT1A", "CKMT1B", "CLPX",
"COX5A", "CYP11A1", "DUT", "ETFA", "FAM82A2", "GATM", "IDH2",
"IDH3A", "IREB2", "IVD", "LACTB", "LDHAL6B", "MRPL46", "MRPS11",
"MTFMT", "MTHFS", "NDUFAF1", "POLG", "RAB11A", "RAB8B", "SQRDL",
"ABAT", "ABCC12", "ACSF3", "ACSM1", "ACSM2A", "ACSM3", "ACSM5",
"BCKDK", "CA5A", "COQ7", "COQ9", "COX4I1", "CYB5B", "DDX28",
"DHODH", "DNAJA3", "DUS2L", "EARS2", "FAHD1", "GFER", "GOT2",
"GPT2", "HAGH", "HSDL1", "KARS", "LDHD", "LYRM1", "Magmas", "MLYCD",
"MRPL28", "MRPS34", "NDUFAB1", "NDUFB10", "NME3", "NME4", "NTHL1",
"PDP2", "PDPR", "PLA2G15", "RHOT2", "SPG7", "TK2", "TMEM186",
"TRAP1", "TUFM", "UQCRC2", "ABCA9", "ACACA", "ACLY", "ACSF2",
"AKAP1", "AKAP10", "ALDH3A2", "ATP5G1", "ATP5H", "ATPAF2", "C17orf90",
"C1QBP", "CCDC44", "COX10", "COX11", "DCAKD", "DRG2", "ELAC2",
"FASN", "FDXR", "GLOD4", "ICT1", "MRM1", "MRPL10", "MRPL12",
"MRPL27", "MRPL45", "MRPS23", "MRPS7", "NAGS", "NME1", "NME1-NME2",
"NME2", "NT5C", "NT5M", "PDK2", "PGS1", "PHB", "POLDIP2", "POLG2",
"PTRH2", "RHOT1", "RILP", "RNMTL1", "RSAD1", "SCO1", "SEPT4",
"SHMT1", "SLC25A10", "SLC25A11", "SLC25A19", "SLC25A35", "SLC25A39",
"TIMM22", "TMEM11", "TTC19", "ACAA2", "AFG3L2", "ATP5A1", "BCL2",
"C18orf19", "C18orf55", "FECH", "ME2", "NARS", "NDUFV2", "SLMO1",
"ZADH2", "ADCK4", "ATP5SL", "ATPBD3", "BAX", "BCAT2", "BCKDHA",
"CLPP", "COX6B1", "COX7A1", "CPT1C", "ECH1", "ECSIT", "ETFB",
"ETHE1", "FDX1L", "FKBP8", "FKSG24", "GCDH", "GPX4", "GTPBP3",
"LONP1", "MRPL34", "MRPL4", "MRPL54", "MRPS12", "NDUFA13", "NDUFA3",
"NDUFA7", "NDUFB7", "NDUFS7", "NUDT19", "OPA3", "POLRMT", "PRODH2",
"RAB11B", "RAB3D", "RAB4B", "RDH13", "SARS2", "SLC25A23", "SLC25A42",
"TIMM44", "TMEM143", "TOMM40", "TSHZ3", "UQCR", "UQCRFS1", "ACSS1",
"ATP5E", "C20orf142", "C20orf24", "C20orf72", "COX4I2", "CRLS1",
"CYP24A1", "FAM65C", "GTPBP5", "IDH3B", "MRPS26", "NFS1", "PAK7",
"PANK2", "ROMO1", "SLMO2", "SNPH", "TOMM34", "UQCC", "VISA",
"ATP5J", "ATP5O", "C21orf33", "C21orf57", "MRPL39", "MRPS6",
"NDUFV3", "SOD1", "ACO2", "ADSL", "AIFM3", "BCL2L13", "BID",
"C22orf32", "CECR5", "CERK", "CHCHD10", "COMT", "CPT1B", "GCAT",
"HSCB", "MCAT", "MPST", "MRPL40", "MTP18", "NDUFA6", "NIPSNAP1",
"PISD", "PRODH", "SAMM50", "SERHL2", "SLC25A1", "SLC25A18", "SMCR7L",
"TOMM22", "TRMU", "TSPO", "TST", "TXN2", "TXNRD2", "UCRC")
#Taken from HUGO "ribosomal protein" family
riboGenes = c('RPLP0','RPLP1','RPLP2','RPL3','RPL3L','RPL4','RPL5','RPL6','RPL7','RPL7A','RPL7L1','RPL8','RPL9','RPL10','RPL10A','RPL10L','RPL11','RPL12','RPL13A','RPL14','RPL15','RPL17','RPL18A','RPL19','RPL21','RPL22','RPL23','RPL23A','RPL24','RPL26','RPL26L','RPL27','RPL27A','RPL28','RPL29','RPL30','RPL31','RPL32','RPL34','RPL35','RPL35A','RPL36','RPL36A','RPL36A','RPL37','RPL37A','RPL38','RPL39','RPL39L','UBA52','RPL41','MRPL1','MRPL2','MRPL3','MRPL4','MRPL9','MRPL10','MRPL11','MRPL12','MRPL13','MRPL14','MRPL15','MRPL16','MRPL17','MRPL18','MRPL19','MRPL20','MRPL21','MRPL22','MRPL23','MRPL24','MRPL27','MRPL28','MRPL30','MRPL32','MRPL33','MRPL34','MRPL35','MRPL36','MRPL37','MRPL38','MRPL39','MRPL40','MRPL41','MRPL42','MRPL43','MRPL44','MRPL45','MRPL46','MRPL47','MRPL48','MRPL49','MRPL50','MRPL51','MRPL52','MRPL53','MRPL54','MRPL55','LACTB','MRPL57','MRPS2','MRPS5','MRPS6','MRPS7','MRPS9','MRPS10','MRPS11','MRPS12','MRPS14','MRPS15','MRPS16','MRPS17','MRPS18','MRPS18','MRPS18','MRPS21','MRPS22','MRPS23','MRPS24','MRPS25','MRPS26','MRPS27','MRPS28','DAP3','MRPS30','MRPS31','MRPS33','MRPS34','MRPS35','MRPS36','RPS2','RPS3','RPS3A','RPS4X','RPS4Y1','RPS4Y2','RPS5','RPS6','RPS7','RPS8','RPS9','RPS10','RPS11','RPS12','RPS13','RPS14','RPS15','RPS15A','RPS16','RPS17','RPS18','RPS19','RPS20','RPS21','RPS23','RPS24','RPS25','RPS26','RPS27','RPS27A','RPS27L','RPS28','RPS29','FAU')
#Taken from HUGO "ribosomal rnas"
riboRNAGenes = c('RNA5S1','RNA5S2','RNA5S3','RNA5S4','RNA5S5','RNA5S6','RNA5S7','RNA5S8','RNA5S9','RNA5S10','RNA5S11','RNA5S12','RNA5S13','RNA5S14','RNA5S15','RNA5S16','RNA5S17','RNA18S1','RNA18S2','RNA18S3','RNA18S4','RNA18S5','RNA28S1','RNA28S2','RNA28S3','RNA28S4','RNA28S5','RNA45S1','RNA45S2','RNA45S3','RNA45S4','RNA45S5','RNA5-8S1','RNA5-8S2','RNA5-8S3','RNA5-8S4','RNA5-8S5','MT-RNR1','MT-RNR2','RNR1RNA','RNR2RNA','RNR3RNA','RNR4RNA','RNR5RNA','RNA18SN1','RNA18SN2','RNA18SN3','RNA18SN4','RNA18SN5','RNA28SN1','RNA28SN2','RNA28SN3','RNA28SN4','RNA28SN5','RNA45SN1','RNA45SN2','RNA45SN3','RNA45SN4','RNA45SN5','RNA5-8SN1','RNA5-8SN2','RNA5-8SN3','RNA5-8SN4','RNA5-8SN5')
#Taken from HUGO "Heat shock genes"
hspGenes = c('HSPA1A','HSPA1B','HSPA1L','HSPA2','HSPA4','HSPA4L','HSPA5','HSPA6','HSPA7','HSPA8','HSPA9','HSPA12A','HSPA12B','HSPA13','HSPA14','HSPH1','HYOU1','DNAJA1','DNAJA2','DNAJA3','DNAJA4','DNAJB1','DNAJB2','DNAJB3','DNAJB4','DNAJB5','DNAJB6','DNAJB7','DNAJB8','DNAJB9','DNAJB11','DNAJB12','DNAJB13','DNAJB14','DNAJC1','DNAJC2','DNAJC3','DNAJC4','DNAJC5','DNAJC5B','DNAJC5G','DNAJC6','DNAJC7','DNAJC8','DNAJC9','DNAJC10','DNAJC11','DNAJC12','DNAJC13','DNAJC14','DNAJC15','DNAJC16','DNAJC17','DNAJC18','DNAJC19','HSCB','DNAJC21','DNAJC22','SEC63','DNAJC24','DNAJC25','GAK','DNAJC27','DNAJC28','SACS','DNAJC30','HSPB1','HSPB2','HSPB3','CRYAA','CRYAB','HSPB6','HSPB7','HSPB8','HSPB9','ODF1','HSPB11','HSP90AA1','HSP90AA3P','HSP90AB1','HSP90B1','TRAP1','BBS10','BBS12','TCP1','CCT2','CCT3','CCT4','CCT5','CCT6A','CCT6B','CCT7','CCT8',
'CLPB','HSPD1','HSPE1','MKKS')
#Usual 13 MT genes
mtGenes = c("MT-ND1", "MT-ND2", "MT-CO1", "MT-CO2", "MT-ATP8","MT-ATP6", "MT-CO3", "MT-ND3", "MT-ND4L", "MT-ND4", "MT-ND5","MT-ND6", "MT-CYB")
#MHC
mhcGenes = c('HLA-A','HLA-B','HLA-C','HLA-DMA','HLA-DMB','HLA-DOA','HLA-DOB','HLA-DPA1','HLA-DPA2','HLA-DPA3','HLA-DPB1','HLA-DPB2','HLA-DQA1','HLA-DQA2','HLA-DQB1','HLA-DQB2','HLA-DQB3','HLA-DRA','HLA-DRB1','HLA-DRB2','HLA-DRB3','HLA-DRB4','HLA-DRB5','HLA-DRB6','HLA-DRB7','HLA-DRB8','HLA-DRB9','HLA-E','HLA-F','HLA-G','HLA-H','HLA-J','HLA-K','HLA-L','HLA-N','HLA-P','HLA-S','HLA-T','HLA-U','HLA-V','HLA-W','HLA-X','HLA-Y','HLA-Z')
mhc1Genes = grep('HLA-[^D]',mhcGenes,value=TRUE)
mhc2Genes = grep('HLA-D',mhcGenes,value=TRUE)
#Haemoglobin genes from HUGO
hgGenes = c('HBA1','HBA2','HBB','HBD','HBE1','HBG1','HBG2','HBM','HBQ1','HBZ')
#Constant part of immunoglobulin from HUGO
igGenes = c('IGHA1','IGHA2','IGHG1','IGHG2','IGHG3','IGHG4','IGHD','IGHE','IGHM','IGLC1','IGLC2','IGLC3','IGLC4','IGLC5','IGLC6','IGLC7','IGKC')
#Antisense genes.  Defined by regex '-AS[0-9]+$'
antisenseGenes = c("TTLL10-AS1", "TP73-AS1", "ENO1-AS1", "PIK3CD-AS1", "PIK3CD-AS2", 
"MTOR-AS1", "NPPA-AS1", "TMEM51-AS1", "ARHGEF19-AS1", "UBXN10-AS1", 
"ZNF436-AS1", "TCEB3-AS1", "MATN1-AS1", "CSMD2-AS1", "ZMYM4-AS1", 
"NFYC-AS1", "SLFNL1-AS1", "SLC2A1-AS1", "SZT2-AS1", "HYI-AS1", 
"KDM4A-AS1", "MKNK1-AS1", "EFCAB14-AS1", "CYP4A22-AS1", "FOXD2-AS1", 
"TTC39A-AS1", "TXNDC12-AS1", "SSBP3-AS1", "DAB1-AS1", "NFIA-AS2", 
"FOXD3-AS1", "ROR1-AS1", "GNG12-AS1", "DEPDC1-AS1", "ZRANB2-AS1", 
"ERICH3-AS1", "NEXN-AS1", "PKN2-AS1", "DPYD-AS1", "DPYD-AS2", 
"RTCA-AS1", "VAV3-AS1", "KCNC4-AS1", "FAM212B-AS1", "KCND3-AS1", 
"SLC16A1-AS1", "AP4B1-AS1", "HIPK1-AS1", "SRGAP2-AS1", "FLG-AS1", 
"UBE2Q1-AS1", "RUSC1-AS1", "ASH1L-AS1", "CADM3-AS1", "GAS5-AS1", 
"RASAL2-AS1", "KIAA1614-AS1", "IPO9-AS1", "LEMD1-AS1", "SERTAD4-AS1", 
"FLVCR1-AS1", "PROX1-AS1", "SPATA17-AS1", "TGFB2-AS1", "LYPLAL1-AS1", 
"HLX-AS1", "TAF1A-AS1", "ITPKB-AS1", "SNAP47-AS1", "IBA57-AS1", 
"LYST-AS1", "LGALS8-AS1", "CHRM3-AS2", "CHRM3-AS1", "HNRNPU-AS1", 
"GCSAML-AS1", "MYT1L-AS1", "TRAPPC12-AS1", "RNASEH1-AS1", "ID2-AS1", 
"DNAJC27-AS1", "AGBL5-AS1", "GTF3C2-AS1", "BIRC6-AS1", "BIRC6-AS2", 
"RMDN2-AS1", "CYP1B1-AS1", "SLC8A1-AS1", "SIX3-AS1", "MEIS1-AS3", 
"MEIS1-AS2", "ATP6V1B1-AS1", "DGUOK-AS1", "BOLA3-AS1", "DCTN1-AS1", 
"LBX2-AS1", "ST3GAL5-AS1", "RGPD4-AS1", "GCC2-AS1", "LIMS1-AS1", 
"SH3RF3-AS1", "PGM5P4-AS1", "DPP10-AS3", "DPP10-AS2", "DPP10-AS1", 
"STEAP3-AS1", "NIFK-AS1", "CCNT2-AS1", "DARS-AS1", "ZEB2-AS1", 
"CCDC148-AS1", "XIRP2-AS1", "DLX2-AS1", "RAPGEF4-AS1", "MLK7-AS1", 
"HOXD-AS2", "TTN-AS1", "FSIP2-AS1", "OSGEPL1-AS1", "SATB2-AS1", 
"CFLAR-AS1", "IDH1-AS1", "LANCL1-AS1", "DIRC3-AS1", "CATIP-AS2", 
"SPATA3-AS1", "IQCA1-AS1", "CAPN10-AS1", "BOK-AS1", "CHL1-AS1", 
"CNTN4-AS2", "CNTN4-AS1", "ITPR1-AS1", "BHLHE40-AS1", "GRM7-AS3", 
"GRM7-AS2", "GRM7-AS1", "LMCD1-AS1", "SRGAP3-AS1", "SRGAP3-AS2", 
"SRGAP3-AS3", "SRGAP3-AS4", "THUMPD3-AS1", "PRRT3-AS1", "SLC6A1-AS1", 
"HDAC11-AS1", "FGD5-AS1", "SH3BP5-AS1", "EAF1-AS1", "PLCL2-AS1", 
"ZNF385D-AS1", "ZNF385D-AS2", "UBE2E2-AS1", "UBE2E1-AS1", "RBMS3-AS3", 
"RBMS3-AS2", "RBMS3-AS1", "OSBPL10-AS1", "CNOT10-AS1", "ARPP21-AS1", 
"ACVR2B-AS1", "ENTPD3-AS1", "VIPR1-AS1", "HHATL-AS1", "CCDC13-AS1", 
"KRBOX1-AS1", "SNRK-AS1", "ZNF197-AS1", "LARS2-AS1", "LIMD1-AS1", 
"LRRC2-AS1", "KIF9-AS1", "CELSR3-AS1", "PRKAR2A-AS1", "BSN-AS2", 
"BSN-AS1", "RBM5-AS1", "SEMA3F-AS1", "SEMA3B-AS1", "RASSF1-AS1", 
"ZMYND10-AS1", "IQCF5-AS1", "ITIH4-AS1", "CACNA2D3-AS1", "WNT5A-AS1", 
"ARHGEF3-AS1", "ARF4-AS1", "DENND6A-AS1", "FLNB-AS1", "FAM3D-AS1", 
"C3orf67-AS1", "SYNPR-AS1", "THOC7-AS1", "PSMD6-AS2", "PRICKLE2-AS1", 
"PRICKLE2-AS1", "PRICKLE2-AS2", "PRICKLE2-AS3", "ADAMTS9-AS1", 
"ADAMTS9-AS2", "MAGI1-AS1", "SUCLG2-AS1", "FOXP1-AS1", "PDZRN3-AS1", 
"CADM2-AS2", "CADM2-AS1", "ST3GAL6-AS1", "ZBTB11-AS1", "MORC1-AS1", 
"PVRL3-AS1", "PLCXD2-AS1", "CFAP44-AS1", "SIDT1-AS1", "ZBTB20-AS1", 
"ZBTB20-AS2", "ZBTB20-AS3", "ZBTB20-AS4", "LSAMP-AS1", "IGSF11-AS1", 
"B4GALT4-AS1", "ARHGAP31-AS1", "MYLK-AS1", "MYLK-AS2", "ITGB5-AS1", 
"ALDH1L1-AS1", "ALDH1L1-AS2", "CCDC37-AS1", "RUVBL1-AS1", "DNAJB8-AS1", 
"GATA2-AS1", "H1FX-AS1", "TMCC1-AS1", "NPHP3-AS1", "TMEM108-AS1", 
"BFSP2-AS1", "NCK1-AS1", "IL20RB-AS1", "CLSTN2-AS1", "ATP1B3-AS1", 
"PLS1-AS1", "PAQR9-AS1", "SLC9A9-AS1", "SLC9A9-AS2", "PLSCR5-AS1", 
"ZIC4-AS1", "HLTF-AS1", "TM4SF1-AS1", "WWTR1-AS1", "ERICH6-AS1", 
"SIAH2-AS1", "CLRN1-AS1", "AADACL2-AS1", "MBNL1-AS1", "MME-AS1", 
"PLCH1-AS1", "PLCH1-AS2", "KCNAB1-AS2", "KCNAB1-AS1", "TIPARP-AS1", 
"IQCJ-SCHIP1-AS1", "IL12A-AS1", "SEC62-AS1", "TMEM212-AS1", "NLGN1-AS1", 
"NAALADL2-AS2", "NAALADL2-AS1", "TBL1XR1-AS1", "KCNMB2-AS1", 
"PEX5L-AS1", "PEX5L-AS2", "CCDC39-AS1", "MCCC1-AS1", "KLHL6-AS1", 
"YEATS2-AS1", "ABCC5-AS1", "HTR3E-AS1", "EIF2B5-AS1", "EHHADH-AS1", 
"IGF2BP2-AS1", "ETV5-AS1", "ADIPOQ-AS1", "LPP-AS2", "LPP-AS1", 
"TPRG1-AS1", "TPRG1-AS2", "P3H2-AS1", "OSTN-AS1", "FGF12-AS1", 
"FGF12-AS2", "FGF12-AS3", "ATP13A5-AS1", "ATP13A4-AS1", "OPA1-AS1", 
"TMEM44-AS1", "XXYLT1-AS1", "XXYLT1-AS2", "TNK2-AS1", "TM4SF19-AS1", 
"UBXN7-AS1", "NCBP2-AS1", "NCBP2-AS2", "MFI2-AS1", "DLG1-AS1", 
"LMLN-AS1", "CTBP1-AS2", "NOP14-AS1", "STX18-AS1", "AFAP1-AS1", 
"CPEB2-AS1", "TAPT1-AS1", "SEPSECS-AS1", "UGDH-AS1", "UCHL1-AS1", 
"OCIAD1-AS1", "USP46-AS1", "LNX1-AS1", "IGFBP7-AS1", "ADGRL3-AS1", 
"EPHA5-AS1", "UBA6-AS1", "THAP9-AS1", "WDFY3-AS1", "FAM13A-AS1", 
"SNCA-AS1", "BMPR1B-AS1", "STPG2-AS1", "TET2-AS1", "RPL34-AS1", 
"SEC24B-AS1", "IL21-AS1", "SCOC-AS1", "SMARCA5-AS1", "HHIP-AS1", 
"SMAD1-AS2", "SMAD1-AS1", "HAND2-AS1", "WWC2-AS2", "WWC2-AS1", 
"F11-AS1", "EXOC3-AS1", "ROPN1L-AS1", "ANKRD33B-AS1", "NADK2-AS1", 
"GDNF-AS1", "EGFLAM-AS4", "EGFLAM-AS3", "EGFLAM-AS2", "EGFLAM-AS1", 
"LIFR-AS1", "OSMR-AS1", "NNT-AS1", "FGF10-AS1", "MAST4-AS1", 
"FOXD1-AS1", "ZBED3-AS1", "SCAMP1-AS1", "RASGRF2-AS1", "CKMT2-AS1", 
"ATG10-AS1", "VCAN-AS1", "TMEM161B-AS1", "MEF2C-AS1", "ARRDC3-AS1", 
"NR2F1-AS1", "RGMB-AS1", "STARD4-AS1", "NREP-AS1", "EPB41L4A-AS1", 
"EPB41L4A-AS2", "SEMA6A-AS1", "ADAMTS19-AS1", "P4HA2-AS1", "C5orf66-AS1", 
"C5orf66-AS2", "SMAD5-AS1", "TRPC7-AS1", "TRPC7-AS2", "ARHGAP26-AS1", 
"JAKMIP2-AS1", "GRPEL2-AS1", "SAP30L-AS1", "HMMR-AS1", "PRR7-AS1", 
"GMDS-AS1", "LY86-AS1", "TFAP2A-AS1", "ELOVL2-AS1", "GFOD1-AS1", 
"JARID2-AS1", "ZSCAN16-AS1", "ZNRD1-AS1", "TRIM31-AS1", "MDC1-AS1", 
"DDR1-AS1", "DDX39B-AS1", "C6orf47-AS1", "SAPCD1-AS1", "EHMT2-AS1", 
"C2-AS1", "C4A-AS1", "C4B-AS1", "HLA-DQB1-AS1", "PSMB8-AS1", 
"BTBD9-AS1", "FOXP4-AS1", "TRAM2-AS1", "MLIP-AS1", "KCNQ5-AS1", 
"OOEP-AS1", "HMGN3-AS1", "TBX18-AS1", "MANEA-AS1", "UFL1-AS1", 
"MCHR2-AS1", "BVES-AS1", "OSTM1-AS1", "ARMC2-AS1", "TRAF3IP2-AS1", 
"RNF217-AS1", "NCOA7-AS1", "MYB-AS1", "PHACTR2-AS1", "STXBP5-AS1", 
"UST-AS1", "RAET1E-AS1", "RAET1E-AS1", "SYNE1-AS1", "EZR-AS1", 
"PACRG-AS2", "PACRG-AS3", "PACRG-AS1", "RPS6KA2-AS1", "MLLT4-AS1", 
"KIF25-AS1", "PSMG3-AS1", "ELFN1-AS1", "MEOX2-AS1", "ISPD-AS1", 
"MACC1-AS1", "KLHL7-AS1", "HOXA-AS2", "HOXA-AS3", "JAZF1-AS1", 
"SEPT7-AS1", "ELMO1-AS1", "TRG-AS1", "POU6F2-AS2", "POU6F2-AS1", 
"INHBA-AS1", "DDC-AS1", "EGFR-AS1", "ABHD11-AS1", "MAGI2-AS1", 
"MAGI2-AS2", "CYP51A1-AS1", "DLX6-AS1", "PCOLCE-AS1", "KMT2E-AS1", 
"SLC26A4-AS1", "DOCK4-AS1", "ST7-AS1", "ST7-AS2", "FEZF1-AS1", 
"POT1-AS1", "JHDM1D-AS1", "NDUFB2-AS1", "WEE2-AS1", "EPHA1-AS1", 
"OR2A1-AS1", "ATP6V0E2-AS1", "PRKAG2-AS1", "PAXIP1-AS2", "PAXIP1-AS1", 
"HTR5A-AS1", "MNX1-AS2", "MNX1-AS1", "ASMTL-AS1", "GYG2-AS1", 
"ARSD-AS1", "PRKX-AS1", "WWC3-AS1", "FRMPD4-AS1", "TLR8-AS1", 
"NHS-AS1", "PPEF1-AS1", "PHKA2-AS1", "EIF1AX-AS1", "PHEX-AS1", 
"ZFX-AS1", "PCYT1B-AS1", "GK-AS1", "TAB3-AS1", "TAB3-AS2", "DMD-AS3", 
"MID1IP1-AS1", "CASK-AS1", "NDP-AS1", "ZNF674-AS1", "UXT-AS1", 
"ZNF630-AS1", "SYP-AS1", "USP27X-AS1", "SPIN4-AS1", "IGBP1-AS2", 
"IGBP1-AS1", "DLG3-AS1", "PHKA1-AS1", "PABPC1L2B-AS1", "PABPC1L2B-AS1", 
"PABPC5-AS1", "DIAPH2-AS1", "ARMCX3-AS1", "TCEAL3-AS1", "MORF4L2-AS1", 
"FRMPD3-AS1", "ALG13-AS1", "PLS3-AS1", "RHOXF1-AS1", "XIAP-AS1", 
"RAP2C-AS1", "HS6ST2-AS1", "DDX26B-AS1", "FGF13-AS1", "MAGEA8-AS1", 
"GPR50-AS1", "HCFC1-AS1", "TMLHE-AS1", "ERICH1-AS1", "DLGAP2-AS1", 
"MCPH1-AS1", "LZTS1-AS1", "EXTL3-AS1", "RBPMS-AS1", "YTHDF3-AS1", 
"RRS1-AS1", "C8orf34-AS1", "LACTB2-AS1", "MSC-AS1", "RDH10-AS1", 
"STAU2-AS1", "ZFHX4-AS1", "PKIA-AS1", "CA3-AS1", "OTUD6B-AS1", 
"C8orf37-AS1", "UBR5-AS1", "AZIN1-AS1", "BAALC-AS2", "BAALC-AS1", 
"RAD21-AS1", "SAMD12-AS1", "HAS2-AS1", "FAM83A-AS1", "FER1L6-AS1", 
"FER1L6-AS2", "RNF139-AS1", "KIAA0196-AS1", "ZFAT-AS1", "RHPN1-AS1", 
"MAFA-AS1", "FAM83H-AS1", "TONSL-AS1", "ZNF252P-AS1", "PGM5P3-AS1", 
"VLDLR-AS1", "RFX3-AS1", "GLIS3-AS1", "CDC37L1-AS1", "PTPRD-AS1", 
"PTPRD-AS2", "LURAP1L-AS1", "FOCAD-AS1", "CDKN2A-AS1", "CDKN2B-AS1", 
"IFT74-AS1", "TOPORS-AS1", "B4GALT1-AS1", "UBE2R2-AS1", "CNTFR-AS1", 
"DNAJB5-AS1", "PGM5-AS1", "PGM5-AS1", "C9orf135-AS1", "MAMDC2-AS1", 
"SMC5-AS1", "RORB-AS1", "C9orf41-AS1", "VPS13A-AS1", "GNA14-AS1", 
"BARX1-AS1", "STX17-AS1", "TMEM246-AS1", "SMC2-AS1", "PAPPA-AS2", 
"PAPPA-AS1", "ASTN2-AS1", "PSMD5-AS1", "GSN-AS1", "SLC25A25-AS1", 
"PTGES2-AS1", "ODF2-AS1", "PRRX2-AS1", "DBH-AS1", "COL5A1-AS1", 
"PPP1R26-AS1", "CCDC183-AS1", "MAN1B1-AS1", "C9orf173-AS1", "ARRDC1-AS1", 
"TOLLIP-AS1", "KRTAP5-AS1", "MRPL23-AS1", "CD81-AS1", "KCNQ1-AS1", 
"CARS-AS1", "MRGPRG-AS1", "RRM1-AS1", "TMEM9B-AS1", "SBF2-AS1", 
"MRVI1-AS1", "ZBED5-AS1", "SPTY2D1-AS1", "NAV2-AS5", "NAV2-AS4", 
"NAV2-AS3", "NAV2-AS2", "NAV2-AS1", "BBOX1-AS1", "CSTF3-AS1", 
"FBXO3-AS1", "ALKBH3-AS1", "LRP4-AS1", "PTPRJ-AS1", "OVOL1-AS1", 
"MRGPRF-AS1", "ANO1-AS2", "ANO1-AS1", "SHANK2-AS1", "SHANK2-AS2", 
"SHANK2-AS3", "ARAP1-AS1", "ARAP1-AS2", "KCTD21-AS1", "RAB30-AS1", 
"PCF11-AS1", "GRM5-AS1", "JRKL-AS1", "NCAM1-AS1", "TMPRSS4-AS1", 
"USP2-AS1", "PKNOX2-AS1", "STT3A-AS1", "ST3GAL4-AS1", "KIRREL3-AS1", 
"KIRREL3-AS2", "KIRREL3-AS3", "NTM-AS1", "IDI2-AS1", "ADARB2-AS1", 
"PITRM1-AS1", "CALML3-AS1", "GATA3-AS1", "CELF2-AS2", "CELF2-AS1", 
"PROSER2-AS1", "VIM-AS1", "ST8SIA6-AS1", "STAM-AS1", "SLC39A12-AS1", 
"NEBL-AS1", "GPR158-AS1", "MKX-AS1", "WAC-AS1", "SVIL-AS1", "ZEB1-AS1", 
"PARD3-AS1", "ZNF32-AS3", "ZNF32-AS1", "ZNF32-AS2", "TMEM72-AS1", 
"FAM170B-AS1", "C10orf71-AS1", "SGMS1-AS1", "PRKG1-AS1", "TMEM26-AS1", 
"JMJD1C-AS1", "UNC5B-AS1", "CDH23-AS1", "DNAJC9-AS1", "ZSWIM8-AS1", 
"ZNF503-AS1", "ZNF503-AS2", "KCNMA1-AS1", "KCNMA1-AS2", "KCNMA1-AS3", 
"ZMIZ1-AS1", "NUTM2B-AS1", "SFTPD-AS1", "TMEM254-AS1", "GRID1-AS1", 
"NUTM2A-AS1", "ACTA2-AS1", "SLC16A12-AS1", "TNKS2-AS1", "PLCE1-AS2", 
"PLCE1-AS1", "ENTPD1-AS1", "SLIT1-AS1", "DNMBP-AS1", "KCNIP2-AS1", 
"CYP17A1-AS1", "NEURL1-AS1", "SH3PXD2A-AS1", "CFAP58-AS1", "SORCS3-AS1", 
"PDCD4-AS1", "WDR11-AS1", "ATE1-AS1", "FAM53B-AS1", "TEX36-AS1", 
"EDRF1-AS1", "FANK1-AS1", "TCERG1L-AS1", "ADGRA1-AS1", "CACNA1C-AS4", 
"CACNA1C-AS3", "CACNA1C-AS2", "CACNA1C-AS1", "CCND2-AS1", "CCND2-AS2", 
"C1RL-AS1", "A2ML1-AS1", "A2ML1-AS2", "A2M-AS1", "PLBD1-AS1", 
"RERG-AS1", "RASSF8-AS1", "ARNTL2-AS1", "OVCH1-AS1", "DDX11-AS1", 
"DENND5B-AS1", "BCDIN3D-AS1", "KRT73-AS1", "HOXC-AS1", "AGAP2-AS1", 
"TMEM5-AS1", "LLPH-AS1", "IFNG-AS1", "CEP83-AS1", "TMPO-AS1", 
"USP30-AS1", "FAM222A-AS1", "MAPKAPK5-AS1", "LHX5-AS1", "TBX5-AS1", 
"TESC-AS1", "PXN-AS1", "CLIP1-AS1", "LATS2-AS1", "SACS-AS1", 
"C1QTNF9B-AS1", "SPATA13-AS1", "C1QTNF9-AS1", "C1QTNF9B-AS1", 
"WASF3-AS1", "USP12-AS1", "USP12-AS2", "PDX1-AS1", "PAN3-AS1", 
"MTUS2-AS2", "MTUS2-AS1", "TEX26-AS1", "FRY-AS1", "SPG20-AS1", 
"FREM2-AS1", "VWA8-AS1", "ENOX1-AS2", "ENOX1-AS1", "SMIM2-AS1", 
"TSC22D1-AS1", "TPT1-AS1", "SLC25A30-AS1", "CPB2-AS1", "HTR2A-AS1", 
"SUCLA2-AS1", "MED4-AS1", "DLEU1-AS1", "DLEU7-AS1", "RNASEH2B-AS1", 
"INTS6-AS1", "DIAPH3-AS1", "DIAPH3-AS2", "PCDH9-AS1", "PCDH9-AS2", 
"PCDH9-AS3", "PCDH9-AS4", "LMO7-AS1", "MYCBP2-AS1", "SCEL-AS1", 
"EDNRB-AS1", "RNF219-AS1", "RBM26-AS1", "NDFIP2-AS1", "GPC5-AS2", 
"GPC5-AS1", "GPC6-AS2", "GPC6-AS1", "SOX21-AS1", "CLDN10-AS1", 
"DNAJC3-AS1", "FARP1-AS1", "STK24-AS1", "DOCK9-AS1", "DOCK9-AS2", 
"UBAC2-AS1", "CLYBL-AS2", "CLYBL-AS1", "PCCA-AS1", "NALCN-AS1", 
"FGF14-AS1", "FGF14-AS2", "DAOA-AS1", "MYO16-AS2", "MYO16-AS1", 
"COL4A2-AS2", "COL4A2-AS1", "ARHGEF7-AS2", "ARHGEF7-AS1", "ATP11A-AS1", 
"MCF2L-AS1", "F10-AS1", "GRTP1-AS1", "GAS6-AS1", "GAS6-AS2", 
"PRMT5-AS1", "DHRS4-AS1", "NOVA1-AS1", "FOXG1-AS1", "ARHGAP5-AS1", 
"EGLN3-AS1", "NKX2-1-AS1", "SLC25A21-AS1", "FRMD6-AS2", "FRMD6-AS1", 
"KTN1-AS1", "OTX2-AS1", "PSMA3-AS1", "HIF1A-AS1", "HIF1A-AS2", 
"FUT8-AS1", "ACTN1-AS1", "DIO2-AS1", "FOXN3-AS1", "FOXN3-AS2", 
"ITPK1-AS1", "FAM181A-AS1", "DICER1-AS1", "MKRN3-AS1", "GABRG3-AS1", 
"DPH6-AS1", "SRP14-AS1", "PLCB2-AS1", "EHD4-AS1", "PLA2G4E-AS1", 
"EIF3J-AS1", "GATM-AS1", "GABPB1-AS1", "RORA-AS1", "RORA-AS2", 
"USP3-AS1", "IQCH-AS1", "THSD4-AS1", "HEXA-AS1", "ADPGK-AS1", 
"C15orf59-AS1", "LOXL1-AS1", "UBL7-AS1", "LINGO1-AS1", "LINGO1-AS2", 
"ANKRD34C-AS1", "ST20-AS1", "TMC3-AS1", "CPEB1-AS1", "AGBL1-AS1", 
"NTRK3-AS1", "CRTC3-AS1", "PRC1-AS1", "NR2F2-AS1", "SPATA8-AS1", 
"CERS3-AS1", "PCSK6-AS1", "LMF1-AS1", "SRRM2-AS1", "MMP25-AS1", 
"ZNF213-AS1", "GLIS2-AS1", "CRYM-AS1", "LCMT1-AS1", "LCMT1-AS2", 
"IL21R-AS1", "ATP2A1-AS1", "CDIPT-AS1", "FBXL19-AS1", "PYCARD-AS1", 
"ITFG1-AS1", "BEAN1-AS1", "VAC14-AS1", "TAT-AS1", "FOXC2-AS1", 
"SNAI3-AS1", "VPS9D1-AS1", "GAS8-AS1", "PITPNA-AS1", "ALOX12-AS1", 
"TMEM220-AS1", "MPRIP-AS1", "RAI1-AS1", "EPN2-AS1", "CCDC144NL-AS1", 
"ABHD15-AS1", "RARA-AS1", "RAMP2-AS1", "ASB16-AS1", "RUNDC3A-AS1", 
"MAP3K14-AS1", "MAPT-AS1", "KANSL1-AS1", "SP2-AS1", "HOXB-AS1", 
"HOXB-AS3", "HOXB-AS2", "HOXB-AS4", "TMEM92-AS1", "CACNA1G-AS1", 
"BZRAP1-AS1", "SEPT4-AS1", "TBX2-AS1", "PRR29-AS1", "PRKCA-AS1", 
"ABCA9-AS1", "KCNJ2-AS1", "HID1-AS1", "C1QTNF1-AS1", "BAIAP2-AS1", 
"AATK-AS1", "MAFG-AS1", "DLGAP1-AS1", "DLGAP1-AS2", "DLGAP1-AS3", 
"DLGAP1-AS4", "DLGAP1-AS5", "L3MBTL4-AS1", "NDUFV2-AS1", "PPP4R1-AS1", 
"LDLRAD4-AS1", "GATA6-AS1", "TTC39C-AS1", "AQP4-AS1", "DSG1-AS1", 
"DSG2-AS1", "SLC14A2-AS1", "TCF4-AS1", "TCF4-AS2", "PARD6G-AS1", 
"NRSN2-AS1", "SDCBP2-AS1", "SIRPG-AS1", "UBOX5-AS1", "MCM8-AS1", 
"LAMP5-AS1", "SNAP25-AS1", "ISM1-AS1", "MACROD2-AS1", "KIZ-AS1", 
"NKX2-2-AS1", "ZNF337-AS1", "HM13-AS1", "ZNF341-AS1", "RALY-AS1", 
"ITCH-AS1", "MMP24-AS1", "FAM83C-AS1", "DLGAP4-AS1", "PLCG1-AS1", 
"OSER1-AS1", "HNF4A-AS1", "KCNK15-AS1", "STK4-AS1", "CSE1L-AS1", 
"CEBPB-AS1", "ADNP-AS1", "BMP7-AS1", "GNAS-AS1", "LAMA5-AS1", 
"C20orf166-AS1", "SLCO4A1-AS1", "OGFR-AS1", "ZBTB46-AS1", "UCKL1-AS1", 
"CIRBP-AS1", "CSNK1G2-AS1", "CACTIN-AS1", "DPP9-AS1", "RAB11B-AS1", 
"ILF3-AS1", "FARSA-AS1", "CEBPA-AS1", "ZNF30-AS1", "HPN-AS1", 
"TMEM147-AS1", "UPK1A-AS1", "ZNF529-AS1", "ZNF790-AS1", "ZNF793-AS1", 
"ZNF571-AS1", "LIPE-AS1", "DACT3-AS1", "NAPA-AS1", "GLTSCR1-AS1", 
"GLTSCR2-AS1", "PLA2G4C-AS1", "CARD8-AS1", "NUCB1-AS1", "PTOV1-AS1", 
"PTOV1-AS2", "ZNF649-AS1", "ZNF350-AS1", "LENG8-AS1", "ZNF582-AS1", 
"ZNF667-AS1", "ZIM2-AS1", "A1BG-AS1", "MZF1-AS1", "ZFY-AS1", 
"NLGN4Y-AS1", "POTEH-AS1", "CECR5-AS1", "THAP7-AS1", "MIF-AS1", 
"ADORA2A-AS1", "ZNRF3-AS1", "HORMAD2-AS1", "MORC2-AS1", "RNF185-AS1", 
"PIK3IP1-AS1", "ISX-AS1", "APOBEC3B-AS1", "MGAT3-AS1", "EP300-AS1", 
"EFCAB6-AS1", "NUP50-AS1", "PRR34-AS1", "GTSE1-AS1", "CHKB-AS1", 
"SAMSN1-AS1", "CHODL-AS1", "CYYR1-AS1", "BACH1-AS1", "GRIK1-AS1", 
"HUNK-AS1", "MIS18A-AS1", "URB1-AS1", "PAXBP1-AS1", "C21orf62-AS1", 
"IL10RB-AS1", "TTC3-AS1", "BRWD1-AS2", "BRWD1-AS1", "B3GALT5-AS1", 
"DSCAM-AS1", "ZNF295-AS1", "LRRC3-AS1", "TSPEAR-AS1", "TSPEAR-AS2", 
"ITGB2-AS1", "COL18A1-AS2", "COL18A1-AS1", "FTCD-AS1", "MCM3AP-AS1"
)
```
