## Supplementary material for "Single cell derived mRNA signals across human kidney tumors": Source Code: logisticRegression.R

```
#' Code to do the cell matching from Young et al., Science, 2018.
#'
#' Train a regularised logistic regression model on some training single cell data, usually reference normal.  Then use this trained model to infer the similarity of every cell in some target data set to the reference.  A basic plotting function to visualise the results.
#'
#' Performance notes:
#'
#' Because this is such a slow process, these functions have been rather aggressively parallelised.  However, parallelism in R is pretty shit, so you need to be careful and pass workers=NULL to disable parallelism if you have issues.  The two functions that use this are trainModel and predictSimilarity.  The most common mode of failure is for the parallel threads to consume all the memory available for the machine.  As long as your data set is small enough that data size * num CPUs < available Mem this should not be an issue.
#'
#' General notes:
#'
#' The first thing you need to do is load this code by running source('<Path_to_this_file>').
#'
#' The basic data you need for this to be useful are two single cell data-sets (I've assumed 10X, but in theory anything should work).  One of those data sets must be your 'reference' or 'training' data-set that you want to match the second data set (the 'test' or 'target') to.  The training data-set must have an annotation; that is each cell must have a class associated with it.  This doesn't have to be anything special, it could be the cluster number if you want to do something quickliy.
#'
#' The idea is that you first train the logistic regression model using the 'trainModel' function on your training data-set.  This will produce a fit object, or more accurately a list of fit objects, one per class in your annotation.  The main thing to think about when performing this step is how to label each cell into classes.  The way the model fit works, each class is considered one at a time and genes are identified that can be used to tell it apart from all other cells in the data set.  So if you have multiple classes that really represent the same thing (e.g., you've over clustered your data), then the model will be confused as it will see things in the "out" group that really should be in the "in" group.
#'
#' Sometimes the fit will fail for a particular class.  This is usually because there are too few cells in a class to get a meaningful fit.  In this case you can try passing the trainModel nfold=x, where x is some number less than 10 (which is the default).  However, I wouldn't recommend it, as any fit will be of dubious biological value.  When a fit fails for one class, the whole function will still proceed, it will just infer NA values for that class when you apply it later on.
#'
#' Fitting the model can take a loooooong time on even a moderate amount of data (>10,000 cells).  My current approach to this is to intelligently sub-sample the data in such cases, which will happen automatically.  But you could also choose to be very patient or implement something more clever.
#'
#' Having trained the model, you then pass the resulting object and your second single cell data set (the 'test' or 'target' data) to the predictSimilarity function.  This will return logit similarity scores for each cell by default.  If you have some annotation or clustering of the target data set, you can supply it to the predictSimilarity function via the "classes" parameter and it will instead give you the predicted similarity on a per-class level.  This is what you will want most of the time.
#'
#' There is a simple little plotting function which can aid in visualising the results.  This is called similarityHeatmap.
#'
#' You can interogate what genes define each model by running 'getMarkers' on the output of trainModel.

#############
# Libraries #
#############
library(glmnet)
if(!suppressWarnings(require(ComplexHeatmap)))
  message("ComplexHeatmap not found.  This library is needed for plotting functions.  It's the best heatmap library, so install it anyway :)")
#Define dummy functions to have parallel calls behave in a serial way
serialParallelCalls = list(MulticoreParam=function(workers) {invisible()},
                           bplapply = function(X,FUN,BPPARAM,...) {lapply(X,FUN,...)},
                           clusterEvalQ = function(cl,expr) {return(list(eval(expr)))},
                           clusterExport = function(cl,varlist,envir='a') {invisible()},
                           makeCluster = function(n) {},
                           stopCluster = function(cl) {invisible()},
                           multicoreWorkers = function() {1}
                           )
if(!suppressWarnings(require(BiocParallel))){
  message("BiocParallel not found. Parallel processing will not work without this library.")
  #Make dummy functions that will allow fall-back to serial.
  list2env(serialParallelCalls,environment())
}

 

#############
# Functions #
#############

#' Smart subset
#'
#' Sub sample classes so that no more than maxCells are selected, but ensuring that populations with fewer than minCells don't get downsampled and that no population falls below this threshold.
#' 
#' @param classes Vector of labels.
#' @param maxSize Downsample the reference data intelligently if the dataset contains more than this many entries total.
#' @param minSize When downsampling, do not let number of entries in a class be reduced below this number due to downsampling.
#' @param verbose Be chatty?
#' @return Vector of indices to use.
smartSubset = function(classes,maxSize=2000,minSize=100,verbose=TRUE){
  #No need to subset
  nEntries = length(classes)
  if(nEntries<=maxSize)
    return(seq_along(classes))
  cnts = table(classes)
  #Sub-sampling ensures those below minSize are untouched and no group goes below minSize after sampling.  
  ssRate = minSize/cnts
  ssRate[ssRate>1] = 1
  #Really inefficient root finder, but should work reliablyish
  optFun = function(e) abs(sum(cnts*((1-ssRate)*e+ssRate))-maxSize)
  x = optimize(optFun,c(0,1))$minimum
  ssRate = (1-ssRate)*x+ssRate
  w = split(seq_along(classes),classes)
  w = unlist(lapply(names(ssRate),function(e) sample(w[[e]],ceiling(length(w[[e]])*ssRate[e]))))
  w = sort(w)
  if(verbose)
    message(sprintf("Sub-sampled from %d entries to %d",nEntries,length(w)))
  return(w)
}
 

#' Normalise data
#'
#' Normalise raw count data.  This is applied to both test and training data.
#'
#' @param refMat A spares matrix, with genes as rows, cells as columns on which the model is to be trained.  Data should be in raw format (i.e., counts).
#' @param lengths If refMat is bulk RNA-seq, use this length vector (or matrix) to normalise the genes.  Equivalent to converting to TPMs.  Defaults does nothing.
#' @return Normalised data matrix.
normData = function(refMat,lengths){
  #Help out sparse functions, so it doesn't make this dense
  if(is(refMat,'sparseMatrix')){
    #If length is a matrix, just divide
    if(!is.null(dim(lengths))){
      dat = refMat/lengths
    }else{
      #Otherwise, save Matrix from itself.
      dat = refMat
      dat@x = dat@x/lengths[dat@i+1]
    }
    dat = t(dat)
    normFacs =Matrix::rowSums(dat)/1e4
    dat@x = dat@x/normFacs[dat@i+1]
    dat@x = log(dat@x+1)
  }else{
    #Easy dense matrix logic
    dat = refMat/lengths
    dat = t(dat)
    dat = dat/rowSums(dat)*1e4
    dat = log(dat+1)
  }
  return(dat)
}

#' Train the model against a reference
#' 
#' Fit the logistic regression model with regularisation using a One versus Rest approach for the single cell data given in the matrix ref, with cluster labels given.  As the model fit procedure is slow, the number of cells is downsampled if it exceeds some threshold.  To preserve the signal from smaller cell clusters, this downsampling is done in such a way as smaller clusters are not downsampled below a certain threshold.
#'
#' @param refMat A spares matrix, with genes as rows, cells as columns on which the model is to be trained.  Data should be in raw format (i.e., counts).
#' @param classes Vector whose length must match the number of columns in \code{ref}.  Each entry indicates which cluster a particular cell belongs to.
#' @param lengths If refMat is bulk RNA-seq, use this length vector (or matrix) to normalise the genes.  Equivalent to converting to TPMs.  Defaults does nothing.
#' @param maxCells Downsample the reference data intelligently if the dataset contains more than this many cells.
#' @param minCells When downsampling, do not let number of cells in a cluster be reduced below this number due to downsampling.
#' @param workers How many parallel threads to use.  Upon failure due to memory exhaustion, workers will be halved until success is reached.
#' @param ... Extra parameters passed to fitting function.
#' @return A model fit, to be used to predict similarities on the target dataset.
trainModel = function(refMat,classes,lengths=rep(1,nrow(refMat)),maxCells=2000,minCells=100,workers=multicoreWorkers(),...){
  ##############
  # Sub-sample #
  ##############
  w = smartSubset(classes,maxCells,minCells)
  ###################
  # Train the model #
  ###################
  #Prepare the data
  dat = normData(refMat[,w],lengths)
  #Train the model
  #Aggressive parallelisation
  while(TRUE){
    fit = tryCatch(multinomialFitCV(dat,classes[w],workers=workers,...),
                   error = function(e) NULL)
    if(!is.null(fit))
      break
    if(is.null(workers))
      stop("Fitting failed.")
    workers = ceiling(workers/2)
    if(workers==1)
      workers=NULL
  }
  #####################
  # Return the result #
  #####################
  return(fit)
}

#' Use trained model to predict similarity on target data
#'
#' Given the output of \code{trainModel} and a target single cell data-set, calculate the similarity score for each cluster.  Optionally can merge estimates per-cluster.  Classes that could not be trained in the reference data are given NA similarity.
#'
#' @param fit Model fit on reference data, from \code{trainModel}
#' @param tgtData Sparse single cell reference data matrix.  Columns are cells, rows are genes.  Gene names must match gene names use when training reference data.  Data should be in raw format (i.e., counts).
#' @param classes If not NULL, merges similarities to give a per-class/cluster similarity score.
#' @param lengths If tgtData is bulk RNA-seq, use this length vector (or matrix) to normalise the genes.  Equivalent to converting to TPMs.  Defaults does nothing.
#' @param minGeneMatch If the fraction of matching genes is less than this, do not proceed.
#' @param logits If FALSE, convert logits into probabilities.
#' @return A similarity matrix giving similarity scores for cells or clusters (columns) against each population in the reference data set(rows).
predictSimilarity = function(fit,tgtData,classes=NULL,lengths=rep(1,nrow(tgtData)),minGeneMatch=0.99,logits=TRUE){
  ###################
  # Reguralise data #
  ###################
  tgtGenes = rownames(fit[[1]]$glmnet.fit$beta)
  if(!all(tgtGenes %in% rownames(tgtData))){
    warning(sprintf("Of %d genes used in training data, only %d found in current data set.",length(tgtGenes),sum(tgtGenes %in% rownames(tgtData))))
    if(sum(tgtGenes %in% rownames(tgtData))/length(tgtGenes) < minGeneMatch){
      stop("Could not match enough genes between training data and tgtData")
    }
    tgtGenes = tgtGenes[tgtGenes %in% rownames(tgtData)]
  }
  #Order them as in reference data
  m = match(tgtGenes,rownames(tgtData))
  if(is.null(dim(lengths))){
    lengths = lengths[m] 
  }else{
    lengths = lengths[m,,drop=FALSE] 
  }
  tgtData = tgtData[m,,drop=FALSE]
  #Process in same way
  dat = normData(tgtData,lengths)
  ######################
  # Infer similarities #
  ######################
  #I've been over-complicating this.  Just want to do a matrix multiplication and need to construct the right B matrix.
  B = lapply(names(fit),function(mark){
               m = match(fit[[mark]]$lambda.1se,fit[[mark]]$glmnet.fit$lambda)
               if(is.null(fit[[mark]]) || m==1){
                 return(NULL)
               }else{
                 tmp = fit[[mark]]$glmnet.fit$beta[tgtGenes,m,drop=FALSE]
               }
               #Name the column
               colnames(tmp) = mark
               return(tmp)
                           })
  #Drop the ones that are NA and merge.
  nullMarks = sapply(B,is.null)
  B = do.call(cbind,B[!nullMarks])
  #Get the results.  No point in keeping sparse past here.
  preds = as.matrix(dat %*% B)
  #Fill in the NAs
  preds = cbind(preds,matrix(NA,nrow=nrow(preds),ncol=sum(nullMarks),dimnames=list(rownames(preds),names(fit)[nullMarks])))
  #Order
  pp = preds[,names(fit),drop=FALSE]
  ####The superior multicore way
  ####Set up cluster
  ###cl = makeCluster(1)
  ####Make the dots normal and bring functions we need into local env
  ###theDots = list(...)
  ###getPopulationOffset=getPopulationOffset
  ###clusterExport(cl,c('fit','dat','tgtGenes','verbose','workers','theDots'),envir=environment())
  ###clusterEvalQ(cl,{library(BiocParallel);library(Matrix)})
  ####Run the thing
  ###tst = lapply(names(fit),function(mark){
  ###               m = match(fit[[mark]]$lambda.1se,fit[[mark]]$glmnet.fit$lambda)
  ###               if(is.null(fit[[mark]]) | m==1){
  ###                 tmp = data.frame(mark = rep(NA,nrow(dat)))
  ###               }else{
  ###                 tmp = fit[[mark]]$glmnet.fit$beta[tgtGenes,m,drop=FALSE]
  ###               }
  ###               #Name the column
  ###               colnames(tmp) = mark
  ###               return(tmp)
  ###                         })
  ###preds = clusterEvalQ(cl,
  ###                     do.call(bplapply,
  ###                             c(list(X=names(fit),
  ###                                    BPPARAM=MulticoreParam(workers),
  ###                                    FUN = function(mark,...) {
  ###                                      if(verbose)
  ###                                        message(sprintf("Doing prediction for label %s",mark))
  ###                                      #Do the prediction manually so we can see what is happening
  ###                                      #Get the lambda value to use
  ###                                      m = match(fit[[mark]]$lambda.1se,fit[[mark]]$glmnet.fit$lambda)
  ###                                      #Is there no good match?  If so, set NAs
  ###                                      if(is.null(fit[[mark]]) | m==1){
  ###                                        tmp = data.frame(mark = rep(NA,nrow(dat)))
  ###                                      }else{
  ###                                        tmp = data.frame(mark = colSums(t(dat) * fit[[mark]]$glmnet.fit$beta[tgtGenes,m]))
  ###                                      }
  ###                                      #Name the column
  ###                                      colnames(tmp) = mark
  ###                                      return(tmp)
  ###                                    }),
  ###                               theDots))
  ###                     )[[1]]
  ###stopCluster(cl)
  ###preds = list()
  ###for(mark in names(fit)){
  ###  message(sprintf("Predicting similarities for class %s",mark))
  ###  #Do the prediction manually so we can see what is happening
  ###  #Get the lambda value to use
  ###  m = match(fit[[mark]]$lambda.1se,fit[[mark]]$glmnet.fit$lambda)
  ###  #Is there no good match?  If so, set NAs
  ###  if(is.null(fit[[mark]]) | m==1){
  ###    preds[[mark]] = data.frame(mark = rep(NA,nrow(dat)))
  ###  }else{
  ###    preds[[mark]] = data.frame(mark = Matrix::colSums(t(dat) * fit[[mark]]$glmnet.fit$beta[tgtGenes,m]))
  ###  }
  ###  #Name the column
  ###  colnames(preds[[mark]]) = mark
  ###  #preds[[mark]] = predict(fit[[mark]],newx=dat[,rownames(fit[[mark]]$glmnet.fit$beta)],s='lambda.1se',newoffset=rep(0,nrow(dat)))
  ###}
  ###pp = do.call(cbind,preds)
  ################
  # Post-process #
  ################
  #Return cluster level summary
  if(!is.null(classes))
    pp = summariseToClass(pp,classes,logit=TRUE)
  #Now decide how to present the results
  if(!logits)
    pp = (1+exp(-pp))**-1
  return(pp)
}

#' Summarise to class
#'
#' Averages probabilities to the class level.  If logits supplied, the averaging is done in probability space and the reconverted to logits.
#'
#' @param mtx The similarity score matrix, with rows being the test data and columns training.
#' @param classes Classes to summarise to.
#' @param logit Are the data in logit form?
summariseToClass = function(mtx,classes,logit=any(mtx<0 | mtx>1)){
  if(length(classes)!=nrow(mtx))
    stop("Classes and matrix size mismatch")
  #If averaging across cluster, this **MUST** be done on probabilities not logits (logits are unbounded and so vulnerable to skewing the answer)
  if(logit)
    mtx = (1+exp(-mtx))**-1
  pp  = do.call(rbind,lapply(split(seq_len(nrow(mtx)),classes),function(e) colMeans(mtx[e,,drop=FALSE])))
  #If logits are then required, convert probability back to logits after the averaging using probabilities
  if(logit)
    pp = log(pp)-log(1-pp)
  return(pp)
} 


#' Results heatmap
#'
#' Create a heatmap showing the results.  Works best when \code{classes} is supplied to \code{predictSimilarity).
#'
#' @param sims Similarity scores as calculated by \code{predictSimilarity}.
#' @param ... Extra parameters to pass to ComplexHeatmaps
#' @return Heatmap object as constructed by Heatmap
similarityHeatmap = function(sims,...){
  #Convert to matrix
  sims = as.matrix(sims)
  #Set colour scheme differently for logits and probabilities
  isLogit = any(sims<0) | any(sims>1)
  probCols = c('#ffffff','#f0f0f0','#d9d9d9','#bdbdbd','#969696','#737373','#525252','#252525','#000000')
  logitCols = c('#8c510a','#d8b365','#f6e8c3','#c7eae5','#5ab4ac','#01665e')
  if(isLogit){
    cols = circlize::colorRamp2(seq(-5,5,length.out=length(logitCols)),logitCols)
  }else{
    cols = circlize::colorRamp2(seq(0,1,length.out=length(probCols)),probCols)
  }
  #Do we have too many target "entities" to show them individually?
  showTgts = nrow(sims)>50
  theDots = list(...)
  params = list('matrix'=sims,
               name=paste0('Predicted\nSimilarity\n',ifelse(isLogit,'(Logit)','(Prob)')),
               col = cols,
               show_row_names = !showTgts,
               show_row_dend = FALSE,
               cluster_rows = showTgts,
               show_column_names = TRUE,
               show_column_dend = FALSE,
               cluster_columns = FALSE,
               column_title = 'Reference classes'
               )
  #The dots take precendence
  w = names(theDots) %in% names(params)
  for(nom in names(theDots)[w]){
    params[[nom]]=theDots[[nom]]
  }
  params = c(params,theDots[!w])
  hm = do.call(Heatmap,params)
  return(hm)
}

#' Get marker genes
#'
#' From a logistic regression model (i.e., the output of trainModel), extract the vector of non-zero coefficients.
#'
#' @param fit The model fit.  Can be either a single glmnet fit object or a named list of them.
#' @param lambda1se Use 1 s.e. Lambda instead of minimum. 
#' @param gns A data.frame to convert the gene given to symbols. The first column must match the rownames of the fit and the second the symbol you want to use.
getMarkers = function(fit,lambda1se=TRUE,gns=NULL){
  if(class(fit)=='cv.glmnet'){
    fit = list(fit=fit)
  }
  out=list()
  for(nom in names(fit)){
    tgt = ifelse(lambda1se,fit[[nom]]$lambda.1se,fit[[nom]]$lambda.min)
    w = which.min(abs(fit[[nom]]$lambda-tgt))
    x = fit[[nom]]$glmnet.fit$beta[,w]
    x = x[x!=0]
    out[[nom]] = data.frame(gene = names(x),
                            coef = x,
                            class = rep(nom,length(x)))
  }
  out = do.call(rbind,out)
  out = out[order(out$class,abs(out$coef)),]
  rownames(out) = NULL
  if(!is.null(gns))
    out$symb = gns[match(out$gene,gns[,1]),2]
  return(out)
}


#' Fit the model
#'
#' Do the OvR fit for every variable.  This just does a simple cross-validation selection of regularisation amount.  Trys to handle common failure modes gracefully.
#'
#' @param x Data matrix
#' @param y Class definitions
#' @param workers Number of cores to use.  NULL kills all parallel machinary. 
#' @param ... Passed to cv.glmnet
#' @return A list containing a model fit object for each unique class in \code{y} or NULL if no model could be fit.
multinomialFitCV = function(x,y,workers=multicoreWorkers(),verbose=is.null(workers) || workers==1,...){
  #Force pure serial execution
  if(is.null(workers))
    list2env(serialParallelCalls,environment())
  #Do them in order of size
  marks = names(sort(table(as.character(y))))
  #Set up cluster
  cl = makeCluster(1)
  #Make the dots normal and bring functions we need into local env
  theDots = list(...)
  getPopulationOffset=getPopulationOffset
  clusterExport(cl,c('x','y','workers','verbose','theDots','marks','getPopulationOffset'),envir=environment())
  clusterEvalQ(cl,{library(BiocParallel);library(glmnet)})
  #Do the multtcore thing.
  fits = clusterEvalQ(cl,
                    do.call(bplapply,
                            c(
                              list(X=marks,
                                   BPPARAM=MulticoreParam(workers),
                                   FUN = function(mark,...){
                                     if(verbose)
                                       message(sprintf("Fitting model for variable %s",mark))
                                     fac = factor(y==mark)
                                     #The two main modes of failure are too few positives and errors constructing lambda.  These should be handled semi-gracefully
                                     tmp = tryCatch(
                                                    cv.glmnet(x,fac,offset=getPopulationOffset(fac),family='binomial',intercept=FALSE,alpha=0.99,type.measure='class',...),
                                                    error = function(e) {return(NULL)}
                                                    )
                                     #A bad fit can either fail entirely, or select a fit with no coefficients
                                     m = match(tmp$lambda.1se,tmp$glmnet.fit$lambda)
                                     #Deviance allows more fine-grained selections to be made when class error is too coarse.  Can be particularly useful when above model defaults to all-zero coefficients
                                     if(is.null(tmp) | m==1){
                                       if(m==1){
                                         warning(sprintf("%s:Default model set all coefficients to zero, refitting using deviance as CV-measure.",mark))
                                       }else{
                                         warning(sprintf("%s:No fit found, refitting using deviance as CV-measure.",mark))
                                       }
                                       tmp = tryCatch(
                                                      cv.glmnet(x,fac,offset=getPopulationOffset(fac),family='binomial',intercept=FALSE,alpha=0.99,type.measure='deviance',...),
                                                      error = function(e) {
                                                        warning(sprintf("%s:Could not fit model.",mark))
                                                        return(NULL)
                                                      })
                                     }
                                     return(tmp)
                                   }),
                              theDots))
                    )[[1]]
  stopCluster(cl)
  names(fits) = marks
  ###for(mark in marks){
  ###  message(sprintf("Fitting model for variable %s",mark))
  ###  fac = factor(y==mark)
  ###  #The two main modes of failure are too few positives and errors constructing lambda.  These should be handled semi-gracefully
  ###  fits[[mark]] = tryCatch(
  ###    cv.glmnet(x,fac,offset=getPopulationOffset(fac),family='binomial',intercept=FALSE,alpha=0.99,type.measure='class',parallel=nParallel>1,...),
  ###    error = function(e) {return(NULL)}
  ###    )
  ###  #A bad fit can either fail entirely, or select a fit with no coefficients
  ###  m = match(fits[[mark]]$lambda.1se,fits[[mark]]$glmnet.fit$lambda)
  ###  #Deviance allows more fine-grained selections to be made when class error is too coarse.  Can be particularly useful when above model defaults to all-zero coefficients
  ###  if(is.null(fits[[mark]]) | m==1){
  ###    if(m==1){
  ###      warning("Default model set all coefficients to zero, refitting using deviance as CV-measure.")
  ###    }else{
  ###      warning("No fit found, refitting using deviance as CV-measure.")
  ###    }
  ###    fits[[mark]] = tryCatch(
  ###                            cv.glmnet(x,fac,offset=getPopulationOffset(fac),family='binomial',intercept=FALSE,alpha=0.99,type.measure='deviance',parallel=nParallel>1,...),
  ###                            error = function(e) {
  ###                              warning(sprintf("Could not fit model for variable %s",mark))
  ###                              return(NULL)
  ###                            })
  ###  }
  ###  #If we still have an "m=1" error, just return the model, it can be handled at the user's discression at the prediction stage.
  ###}
  return(fits)
}

#' Get offset for each population/cluster
#'
#' Calculate appropriate intercept term to make the training insensitive to the observed frequency of different populations.
#'
#' @param y A factor or string with two-levels to calculate the appropriate offset.
#' @return A vector of offsets to use in the model fit.
getPopulationOffset = function(y){
  if(!is.factor(y))
    y=factor(y)
  if(length(levels(y))!=2)
    stop("y must be a two-level factor")
  off = sum(y==levels(y)[2])/length(y)
  off = log(off/(1-off))
  return(rep(off,length(y)))
}
```
