## Supplementary material for "Single cell derived mRNA signals across human kidney tumors": Source Code: plotParts.R

```
#' Add density contours for each group
#' 
#' Given x/y coordinates for data, split by group, creates 2D density contours for each and plots them.
#'
#' @param x x co-ord
#' @param y y co-ord
#' @param clust Grouping of points.
#' @param nGrid How many grid-points to use to evaluate density.
#' @param nSplits How many parts to split the density range into.
#' @param splitsToUse Which splits to use in drawing contours.  Must be in range 1 to nSplits.  Lower numbers are at edges of clusters.
#' @param col Colour to draw line.  Length must match number of levels in clust or be 1. 
#' @param lwd Line thickness.  Length must match number of levels in clust or be 1. 
#' @param shadeCol Shade within the outer contour.  No shading if NULL.  If not, must be 1 or number of clusters.
#' @param useKS Should we use the ks package?
#' @param ... Passed to contour.
addDensityContours = function(x,y,clust,nGrid=500,nSplits=20,splitsToUse=seq(2,min(4,nSplits)),col='black',lwd=1,shadeCol=NULL,useKS=FALSE,...){
  #plot(x,y,type='n')
  clust = factor(clust)
  if(length(col)==1)
    col = rep(col,nlevels(clust))
  if(length(lwd)==1)
    lwd = rep(lwd,nlevels(clust))
  if(!is.null(shadeCol) && length(shadeCol)==1)
    shadeCol = rep(shadeCol,nlevels(clust))
  ii=0
  for(cl in levels(clust)){
    ii=ii+1
    w = which(clust==cl)
    if(useKS){
      require(ks)
      #This approach is better in many ways, except in that the call it makes to contour internally does not set axes=FALSE or pass the dots.  This is what is needed to prevent redrawing the bounding box and axes over and over.
      kk = kde(cbind(x[w],y[w]),gridsize=nGrid,xmin=c(min(x),min(y)),xmax=c(max(x),max(y)))
      #First do the shading
      if(!is.null(shadeCol)){
        plot(kk,
             display='filled.contour2',
             cont=95,
             add=TRUE,
             drawLabels=FALSE,
             col=c(NA,shadeCol[ii]),
             lwd.fc=0,
             axes=FALSE,
             ...)
      }
      #Then the lines
      plot(kk,
           display='filled.contour2',
           cont=95,
           add=TRUE,
           drawLabels=FALSE,
           col=c(NA),
           lwd.fc=1,
           axes=FALSE,
           ...)
    }else{
      require(MASS)
      kk = kde2d(x[w],y[w],n=nGrid,lims=c(range(x),range(y)))
      contour(kk,
              levels=pretty(range(kk$z),nSplits)[splitsToUse],
              col = col[ii],
              lwd = lwd[ii],
              drawlabels=FALSE,
              add=TRUE,
              ...)
      #Add shading?
      cl = contourLines(kk,levels=pretty(range(kk$z),nSplits)[splitsToUse])
      if(!is.null(shadeCol)){
        polygon(cl[[1]]$x,cl[[1]]$y,col=shadeCol[ii])
      }
    }
  }
  #points(x,y,cex=0.1,pch=19)
}

#' Adds transparency to colour
#'
#' @param cols Vector of colours.
#' @param alphas Single value or vector of alphas
#' @param ... Passed to rgb
#' @return rgb colours with transparency set.
colAlpha = function(cols,alphas,...) {
  if(length(alphas)==1)
    alphas = rep(alphas,length(cols))
  tmp = col2rgb(cols)
  sapply(seq_len(ncol(tmp)),function(e) rgb(tmp[1,e],tmp[2,e],tmp[3,e],alphas[e]*255,maxColorValue=255,...))
}

#' Adds a colorbar legend to a plot
#'
#' Specify x,y as in legend.  Then draws colorbar as a series of boxes with some ticks and labels for scale.
#'
#' @param x Left edge of colorbar.  Or a character like 'left' as in legend.
#' @param y Top edge of colorbar.
#' @param col The vector of colours that make up the bar.
#' @param breaks The values corresponding to those colours.
#' @param barFrac Fraction of y-axis plot area taken up with the bar.
#' @param ticks Number of ticks, or their values.  Must lie in range(breaks). If NULL, guess automatically.
#' @param tickLabs Labels to draw at ticks.  If NULL, use numbers.
#' @param nBoxes How many little boxes to use to build the bar.
#' @param cex cex value
#' @param xpd Passed where it's needed.
addColorBar = function(x,y=NULL,col,breaks,title=NULL,barFrac=0.2,ticks=NULL,tickLabs=NULL,nBoxes=50,cex=1,xpd=NA){
  if(length(breaks)!=length(col) & length(breaks)==2)
    breaks = seq(breaks[1],breaks[2],length.out=length(col))
  #How many colours?
  colMod = circlize::colorRamp2(breaks,col)
  colVals = seq(min(breaks),max(breaks),length.out=nBoxes)
  cols = colMod(colVals)
  #Currently defined plot area
  usr = par('usr')
  xLims = usr[1:2]
  yLims = usr[3:4]
  #Make the cex local
  Cex = cex * par('cex')
  #How much space does one character take up in this plot-space?
  charSize = xyinch(par('cin'),warn.log=FALSE)
  charSize = charSize * Cex
  #How big should we make the bar?
  barSize = c(charSize[1],(yLims[2]-yLims[1])*barFrac)
  #Get tick values
  if(is.null(ticks)){
    ticks = floor(barSize[2]/strheight(min(breaks),units='user',cex=cex))
    #Make sure it's at least 2
    if(ticks<2)
      ticks=2
    #And if it's bigger than 2, make sure it's odd
    if(ticks>2 & ticks%%2==0)
      ticks=ticks-1
  }
  if(length(ticks)==1)
    ticks = seq(min(breaks),max(breaks),length.out=ticks)
  if(!all(ticks<=max(breaks) & ticks>=min(breaks)))
    stop("Ticks must be within range of breaks.")
  if(is.null(tickLabs))
    tickLabs = sprintf('%.02g',ticks)
  ##How big should we make the bar?
  #barSize = c(1,min(5,length(ticks)))*charSize
  ##How big should we make a box
  #boxSize = c(barSize[1],barSize[2]/nBoxes)
  #The +1 is for some wiggle room
  legHeight = barSize[2] + (!is.null(title))*charSize[2]
  #Assume 4 figures for numbers, 1 for box, 1 for wiggle
  legWidth = barSize[1] + charSize[1] + max(strwidth(tickLabs,units='user',cex=cex))
  if(!is.character(x)){
    left = x
    top = y
  }else{
    insetx = 0
    insety = 0
    left = switch(x,
                  bottomright = , topright =, right = xLims[2] - insetx - legWidth,
                  bottomleft = ,left = , topleft = xLims[1] + insetx+legWidth,
                  bottom =, top= ,center = mean(xLims)-legWidth/2)
    top = switch(x,
                 bottomright = , bottom = , bottomleft = yLims[1]+legHeight + insety,
                 topleft = , top = , topright = yLims[2] - insety,
                 left = , right = , center = mean(yLims) +legHeight/2)
    if(is.null(top))
      stop(sprintf("%s is an invalid position string.  See ?legend",x))
  }
  #Now make the boxes with the colours
  tranFun = function(dim,val){
    isLog = ifelse(dim==1,par('xlog'),par('ylog'))
    if(isLog){
      10**val
    }else{
      val
    }
  }
  tmp = seq(top,top-barSize[2],length.out=nBoxes+1)
  rect(xleft = tranFun(1,rep(left,nBoxes)),
       xright = tranFun(1,rep(left+barSize[1],nBoxes)),
       ytop = tranFun(2,tmp[-length(tmp)]),
       ybottom = tranFun(2,tmp[-1]),
       col=cols,
       border=NA,
       xpd=xpd
  )
  #Make the border of the whole thing
  rect(xleft=tranFun(1,left),
       xright=tranFun(1,left+barSize[1]),
       ytop=tranFun(2,top),
       ybottom=tranFun(2,top-barSize[2]),
       col=NA,
       xpd=xpd,
       border=grey(0.2))
  #Add tick marks
  tickLocs = (ticks-breaks[1])/(breaks[length(breaks)]-breaks[1])
  for(i in seq_along(ticks)){
    y = tranFun(2,top-tickLocs[i]*barSize[2])
    lines(tranFun(1,c(left,left+barSize[1]*0.1)),
          c(y,y),
          col=grey(0.2),
          xpd=xpd)
    lines(tranFun(1,c(left+barSize[1]*0.9,left+barSize[1])),
          c(y,y),
          col=grey(0.2),
          xpd=xpd)
  }
  #And the labels
  text(x=tranFun(1,left+barSize[1]*1.1),y=tranFun(2,top-tickLocs*barSize[2]),labels = tickLabs,adj=c(0,0.5),cex=cex,xpd=xpd)
  #And the title
  if(!is.null(title)){
    text(x=tranFun(1,left+barSize[1]*0.5),y=tranFun(2,top+charSize[2]*0.5),labels=title,adj=c(0.5,0),cex=cex,xpd=xpd)
  }
}

#' Add Hexbin
#'
#' Basically a thin wrapper around hexbin that draws bins using polygon rather than the grids BS.
#'
#' @param ... Passed to hexbin
#' @param minCnts The lower limit on counts-per-bin for plotting.  Defaults to 0.
#' @param maxCnts The upper limit on counts-per-bin for plotting.  Defaults to maximum count.
#' @param colSet Colours to map to count range using colorRamp2.  From low to high.
#' @param alpha Transparency.
#' @param logCounts Should we log (base 10) the counts?
#' @return The bin range.
addHexbin = function(...,colSet=c(grey(.8),grey(.1)),alpha=1,logCounts=FALSE,minCnts=0,maxCnts=NULL){
  require(hexbin)
  dat = hexbin(...)
  cnts = dat@count
  xbins = dat@xbins
  shape = dat@shape
  tmp = hcell2xy(dat)
  xnew = tmp$x
  ynew = tmp$y
  sx = xbins/diff(dat@xbnds)
  sy = (xbins*shape)/diff(dat@ybnds)
  inner = 1/2
  outer = (2*inner)/sqrt(3)
  dx = inner/sx
  dy = outer/(2*sy)
  hexC = hexcoords(dx, dy, sep = FALSE)
  #Determine the top range
  if(is.null(maxCnts))
    maxCnts = max(cnts)
  #Decide on colours
  if(logCounts){
    cnts = log10(cnts)
    maxCnts = log10(maxCnts)
  }
  breaks = seq(0,maxCnts,length.out=length(colSet))
  col = circlize::colorRamp2(breaks,colSet,transparency=1-alpha)
  for(i in seq_along(xnew)){
    polygon(x = xnew[i]+hexC$x[seq(6)],
            y = ynew[i]+hexC$y[seq(6)],
            border = col(cnts[i]),
            col=col(cnts[i]))
  }
  return(list(breaks=rev(breaks),col=colSet))
}
```
