## Supplementary material for "Single cell derived mRNA signals across human kidney tumors": Source Code: processCCSK.R

```
setwd('~/trueHome/Projects/CellDeconvolution')

#############
# Libraries #
#############
library(Seurat)
library(Matrix)
import('Code/logisticRegression.R',as='lr')
import('Code/scQC.R')
import('Code/geneSets.R')


##############
# Parameters #
##############
outDir = 'Results/scData'
srcs = 'Data/scTumours'
srcs = file.path(srcs,list.files(srcs))
names(srcs) = c('MRTa','MRTb','CCSK1a','CCSK1b','CCSK2a','CCSK2b','CCSK3')
strained = setNames(file.path(outDir,paste0(names(srcs),'_strained')),names(srcs))
#Mask grouping things by expreiment
exMask = gsub('[ab]$','',names(srcs))
#MT cut-off
mtCut = 0.3
#nGene Cut
minNumGenes = 300
#nUMI cut
minNumUMI = 1000
#How many doublets in a cluster before you throw the entire cluster
maxScrubFrac = 0.5
#UMAP parameters to try in grid
minDists = c(0,.01,.1,.3,.5)
NNs = c(5,10,20,50,100)
#Final parameters
finalMinDist = 0.5
finalNN = 50
#Clustering resolution
clusterRes=1.0
#Populations of genes to exclude from the analysis
excludeGenes = unlist(as.list(geneSets)[c('mtGenes','hspGenes','riboGenes')],use.name=FALSE)
excludeGenes = sort(unique(excludeGenes))
#Number of PCs to use for quick analysis during QC
defNumPCs=75

#############
# Functions #
#############
#Copy/pasted from dropletutils.  A useful function, but the package has an absurd number of dependencies
write10xCounts = function (path, x, barcodes = colnames(x), gene.id = rownames(x),
    gene.symbol = gene.id, overwrite = FALSE)
{
    temp.path <- tempfile(tmpdir = dirname(path))
    dir.create(temp.path, showWarnings = FALSE)
    on.exit({
        if (file.exists(temp.path)) {
            unlink(temp.path, recursive = TRUE)
        }
    })
    writeMM(x, file = file.path(temp.path, "matrix.mtx"))
    if (ncol(x) != length(barcodes)) {
        stop("'barcodes' must of of the same length as 'ncol(x)'")
    }
    write(barcodes, file = file.path(temp.path, "barcodes.tsv"))
    if (length(gene.id) != length(gene.symbol) || length(gene.id) !=
        nrow(x)) {
        stop("lengths of 'gene.id' and 'gene.symbol' must be equal to 'nrow(x)'")
    }
    write.table(data.frame(gene.id, gene.symbol), file = file.path(temp.path,
        "genes.tsv"), row.names = FALSE, col.names = FALSE, quote = FALSE,
        sep = "\t")
    if (overwrite) {
        unlink(path, recursive = TRUE)
    }
    else if (file.exists(path)) {
        stop("specified 'path' already exists")
    }
    file.rename(temp.path, path)
    return(invisible(TRUE))
}
#' Fine clustering for SoupX and other things
#'
#' @param dat Raw filtered count matrix.
#' @param numPCs How many PCs to use.
#' @param clusteringRes Clustering parameter to use.
#' @return Seurat object.
quickCluster = function(dat,numPCs=75,clusteringRes=10.0){
  ss = CreateSeuratObject(dat)
  ss = NormalizeData(ss,verbose=FALSE)
  ss = FindVariableFeatures(ss,verbose=FALSE)
  ss = ScaleData(ss,verbose=FALSE)
  ss = RunPCA(ss,npcs=numPCs,approx=FALSE,verbose=FALSE)
  ss = FindNeighbors(ss,dims=seq(numPCs),verbose=FALSE)
  ss = FindClusters(ss,res=clusteringRes,verbose=FALSE)
  ss = RunUMAP(ss,dims=seq(numPCs),verbose=FALSE)
  return(ss)
}
#' Standard processing
stdProcess = function(dat,...){
  ss = CreateSeuratObject(dat)
  ss = NormalizeData(ss,verbose=FALSE)
  #ss = CellCycleScoring(ss, s.features=geneSets[['S']][geneSets[['S']] %in% rownames(dat)],g2m.features=geneSets[['G2M']][geneSets[['G2M']] %in% rownames(dat)],seed=sample(1e9,1))
  #ss = ScaleData(ss,vars.to.regress=c('S.Score','G2M.Score'),verbose=FALSE,...)
  ss = ScaleData(ss,verbose=FALSE,...)
  return(ss@assays$RNA@scale.data)
}
#' Automatically calculate the contamination fraction
#'
#' The idea of this method is that genes that are highly expressed in the soup and be marker genes for some population can be used to estimate the background.  Marker genes are identified using the tfidf method (see \code{\link{quickMarkers}}).  The contamination fraction is then calculated at the cluster level for each of these genes and clusters are then aggressively pruned to remove those that give implausible estimates.  A final estimate is based on the average across these estimates, under the assumption that biased estimates will not cluster, while true ones will (around the true value).
#'
#' Note that the interpretation of the tf-idf cut-off in this context is that a cut-off t implies that a marker gene has the property that geneFreqGlobal < exp(-t/geneFreqInClust).
#'
#' @param sc The SoupChannel object.
#' @param nMarks How many marker genes to use. These must also pass the tfidf cut-off and be highly expressed in the soup, so there may end up being fewer than this number.
#' @param tfidfMin Minimum value of tfidf to accept for a marker gene.
#' @param soupMin Minimum value of expression in the background to accept for marker.
#' @param rhoMax
#' @param FDR
#' @param clustPerGene
#' @param maxClustFrac
#' @param minCnts
autoCont = function(sc,nMarks = 50,tfidfMin=0.8,soupMin=1e-4,rhoMax=0.5,FDR=0.1,clustPerGene=1,maxClustFrac=0.5,minCnts=1){
  #First collapse by cluster
  s = split(rownames(sc$metaData),sc$metaData$clusters)
  tmp = do.call(cbind,lapply(s,function(e) rowSums(sc$toc[,e,drop=FALSE])))
  ssc = sc 
  ssc$toc = tmp
  ssc$metaData = data.frame(nUMIs = colSums(tmp),row.names=colnames(tmp))
  ###################
  # Get best markers
  #Get the top N soup Genes
  tgts = ssc$soupProfile[order(ssc$soupProfile$est,decreasing=TRUE),]
  tgts = rownames(tgts)[tgts$est>soupMin]
  #Refine this to the best markers we can manage
  mrks = quickMarkers(sc$toc,sc$metaData$clusters,N=Inf)
  #Keep only the ones that are high in soup
  mrks = mrks[mrks$gene %in% tgts,]
  #And only the most specific entry for each gene
  mrks = mrks[order(mrks$gene,-mrks$tfidf),]
  mrks = mrks[!duplicated(mrks$gene),]
  #Order by tfidif maxness
  mrks = mrks[order(-mrks$tfidf),]
  #Apply tf-idf cut-off
  mrks = mrks[mrks$tfidf > tfidfMin,]
  tgts = head(mrks$gene,nMarks)
  ############################
  # Get estimates in clusters
  #Get which ones we'd use and where with canonical method
  tmp = as.list(tgts)
  names(tmp) = tgts
  ute = estimateNonExpressingCells(sc,tmp,maximumContamination=rhoMax,FDR=FDR)
  m = rownames(sc$metaData)[match(rownames(ssc$metaData),sc$metaData$clusters)]
  ute = t(ute[m,])
  colnames(ute) = rownames(ssc$metaData)
  #Now calculate the observed and expected counts for each cluster for 
  expCnts = outer(ssc$soupProfile$est,ssc$metaData$nUMIs)
  rownames(expCnts) = rownames(ssc$soupProfile)
  colnames(expCnts) = rownames(ssc$metaData)
  expCnts = expCnts[tgts,]
  #And the observed ones
  obsCnts = ssc$toc[tgts,]
  #Filter out the shite
  #Get the p-value for this being less than 1
  pp = ppois(obsCnts,expCnts*rhoMax,lower.tail=TRUE)
  qq = p.adjust(pp,method='BH')
  qq = matrix(qq,nrow=nrow(pp),ncol=ncol(pp),dimnames=dimnames(pp))
  #Get the cluster level ratio
  rhos = obsCnts/expCnts
  #Index in range
  rhoIdx = t(apply(rhos,1,function(e) order(order(e))))
  #Make a data.frame with everything
  dd = data.frame(gene = rep(rownames(ute),ncol(ute)),
                  passNonExp = as.vector(ute),
                  rho = as.vector(rhos),
                  rhoIdx = as.vector(rhoIdx),
                  obsCnt = as.vector(obsCnts),
                  expCnt = as.vector(expCnts),
                  qVal = as.vector(qq)
                  )
  dd$pass = dd$obsCnt >= minCnts & 
    dd$qVal < FDR & 
    dd$rhoIdx <= min(clustPerGene,floor(ncol(rhoIdx)*maxClustFrac)) & 
    dd$passNonExp
  #I think the best way to do this is based on the density.
  tmp = density(dd$rho[dd$pass])
  dd$rhoEst = tmp$x[which.max(tmp$y)]
  return(dd)
}

#########
# SoupX #
#########
for(ex in names(srcs)){
  message(sprintf("Running SoupX on channel %d of %d in experiment %s",i,length(srcs),ex))
  sc = load10X(srcs[ex])
  tmp = autoCont(sc,rhoMax=1)
  sc = setContaminationFraction(sc,tmp$rhoEst[1])
  out = adjustCounts(sc,verbose=2,nCores=12)
  #Save it
  write10xCounts(strained[ex],out,overwrite=TRUE)
}


############
# Basic QC #
############
tgtFile = file.path(outDir,'baseSeurat.RDS')
if(file.exists(tgtFile)){
  srats = readRDS(tgtFile)
}else{
  #Create the basic Seurat object for each experiment
  srats = list()
  for(mask in unique(exMask)){
    noms = names(srcs)[exMask==mask]
    message(sprintf("QC filters on sample %s",mask))
    pdf(file.path(outDir,sprintf('QC_%s.pdf',mask)))
      srat = scQC$basicQC(dataDirs=strained[noms],
                          excludeGenes = excludeGenes,
                          geneSets = geneSets,
                          maxMT = mtCut,
                          minGenes = 300,
                          minUMIs = 300,
                          maxBadFrac = maxScrubFrac,
                          scrubScoreMax = 1.2,
                          numPCs = defNumPCs)
    dev.off()
    srats[[mask]] = srat
  }
  saveRDS(srats,tgtFile)
}

####################
# Final Processing #
####################
srat = ScaleData(srat)
srat = RunPCA(srat,npcs=nPCs,approx=FALSE)
#Run tSNE as well in case that turns out to be useful.
#srat = RunTSNE(srat)
#The old python based method seems to segfault every time...
srat = FindNeighbors(srat,dims=seq(nPCs))
srat = FindClusters(srat,res=clusterRes)
#Final UMAP
srat = RunUMAP(srat,dims=seq(nPCs),min.dist=finalMinDist,n.neighbors=finalNN)
mm = srat@meta.data$seurat_clusters
mm = ifelse(mm %in% c(),mm,
            ifelse(mm %in% c(),
                   'Top',
                   'Bottom'
                   )
            )
qq = quickMarkers(srat@assays$RNA@counts,mm)


############
# Train LR #
############
#Kidney
fKid = readMM('Data/scRef/fKidMerge.mtx')
rownames(fKid) = read.table('Data/scRef/fKidMergeRownames.tsv',sep='\t',header=TRUE)$geneID
colnames(fKid) = read.table('Data/scRef/fKidMergeColnames.tsv',sep='\t',header=TRUE)$cellID
fKidClasses = gsub(':.*','',colnames(fKid))
fKidFit = lr$trainModel(fKid,fKidClasses)
#Adrenal
fAd = readMM('Data/scRef/foetalAdrenalRef.mtx')
rownames(fAd) = read.table('Data/scRef/foetalAdrenalRefRownames.tsv',sep='\t',header=TRUE)$geneID
colnames(fAd) = read.table('Data/scRef/foetalAdrenalRefColnames.tsv',sep='\t',header=TRUE)$cellID
fAdClasses = gsub(':.*','',colnames(fAd))
fAdFit = lr$trainModel(fAd,fAdClasses)
gns = read.table('Data/genes.tsv',sep='\t')


#Process each separately
mask = gsub('[ab]$','',names(srcs))
srats = list()
for(nom in unique(masks)){
  tmp = srcs[mask==nom]
  srats[[nom]] = scQC$basicQC(dataDirs = setNames(file.path(srcs,'filtered_feature_bc_matrix'),names(srcs)),
                              excludeGenes = excludeGenes,

                              geneSets = geneSets,
                              maxMT = mtCut,
                              minGenes = minNumGenes,
                              minUMIs = minNumUMI,
                              maxBadFrac = maxScrubFrac,
                              numPCs = defNumPCs)
}
###sPhaseGenes = cc.genes.updated.2019$s.genes
###g2mPhaseGenes = cc.genes.updated.2019$g2m.genes
###clusteringRes=1.0
###numPCs=75
###srat = 
###srat = srcs[mask==nom]
###srat = Read10X(setNames(file.path(srat,'filtered_feature_bc_matrix'),names(srat)))
###srat = CreateSeuratObject(srat)
###srat = NormalizeData(srat,verbose=FALSE)
###srat = CellCycleScoring(srat, s.features=sPhaseGenes,g2m.features=g2mPhaseGenes,seed=sample(1e9,1))
###srat = FindVariableFeatures(srat,verbose=FALSE)
###srat = ScaleData(srat,verbose=FALSE)
###srat = RunPCA(srat,npcs=numPCs,approx=FALSE,verbose=FALSE)
###srat = FindNeighbors(srat,dims=seq(numPCs),verbose=FALSE)
###srat = FindClusters(srat,res=clusteringRes,verbose=FALSE)
###srat = RunUMAP(srat,dims=seq(numPCs),verbose=FALSE)
 
```
