## Supplementary material for "Single cell derived mRNA signals across human kidney tumors": Source Code: processRefData.R

```
#' Process the single cell data sets that we will use as references
#' 
#' Most references we re-process from scratch.  However, some samples (such as Marioni mouse data) are sufficiently processed and annotated that it makes no sense to be a wholesale re-processing and annotation.  These are handled at the end of the script.

#####################
# Project directory #
#####################
setwd('~/trueHome/Projects/CellDeconvolution')

#############
# Libraries #
#############
library(Seurat)
library(SoupX)
library(ggplot2)
library(DropletUtils)
library(biomaRt)
library(Matrix)
source('https://raw.githubusercontent.com/constantAmateur/MCVR/master/code.R')
import('Code/scQC.R',as='scQC')
import('Code/geneSets.R')
import('Code/logisticRegression.R',as='lr')

#############
# Functions #
#############

#' Standard processing
stdProcess = function(dat,...){
  ss = CreateSeuratObject(dat)
  ss = NormalizeData(ss,verbose=FALSE)
  #ss = CellCycleScoring(ss, s.features=geneSets[['S']][geneSets[['S']] %in% rownames(dat)],g2m.features=geneSets[['G2M']][geneSets[['G2M']] %in% rownames(dat)],seed=sample(1e9,1))
  #ss = ScaleData(ss,vars.to.regress=c('S.Score','G2M.Score'),verbose=FALSE,...)
  ss = ScaleData(ss,verbose=FALSE,...)
  return(ss@assays$RNA@scale.data)
}

#' Load a matrix
#'
#' @param src Where to load it
loadRefMtx = function(src){
  mtx = readMM(paste0(src,'.mtx'))
  rownames(mtx) = read.table(paste0(src,'Rownames.tsv'),sep='\t',header=TRUE)$geneID
  colnames(mtx) = read.table(paste0(src,'Colnames.tsv'),sep='\t',header=TRUE)$cellID
  return(mtx)
}


##############
# Parameters #
##############
#Output directory
outDir = 'Results/preProcess/scData'
srcs = list()
#Foetal kidney
tmp = list.files('Data/scData/fKidScience/',full.names=TRUE)
names(tmp) = basename(tmp)
srcs[['fKid']] = list(fKidScience=tmp,
                      fKidPlos='Data/scData/fKidPlos')
#Manifest for mKid
cMani = read.table('Data/TableS6.tsv',sep='\t',header=TRUE)
tmp = list.files('Data/scData/mKid',full.names=TRUE)
names(tmp) = cMani$Label[match(basename(tmp),cMani$SangerID)]
srcs[['mKid']] = list(mKidScience = tmp)
#Adult Bone Marrow Immune cells
srcs[['mBM']] = 'Data/scData/mBM/'
#Foetal adrenal
tmp = list.files('Data/scData/fAd',full.names=TRUE)
srcs[['fAd']] = lapply(tmp,function(e) {
                     tmp = grep('.RDS$',list.files(e,full.names=TRUE),invert=TRUE,value=TRUE)
                     setNames(tmp,basename(tmp))})
names(srcs[['fAd']]) = basename(tmp)
srcs$fAd$bilateralAdrenals = setNames(file.path(srcs$fAd$bilateralAdrenals,'outs'),names(srcs$fAd$bilateralAdrenals))
#Borring PBMCs
srcs$PBMC = list(PBMC_10k=c(PBMC_10k='Data/scData/PBMC'))
#Well annotated PBMCs
srcs$PBMC3k = list(PBMC_3k=c(PBMC_3k='Data/scData/PBMC3k'))
#MT cut-off
mtCut = 0.3
#nGene Cut
minNumGenes = 300
#nUMI cut
minNumUMI = 1000
#How many doublets in a cluster before you throw the entire cluster
maxScrubFrac = 0.5
#UMAP parameters to try in grid
minDists = c(0,.01,.1,.3,.5)
NNs = c(5,10,20,50,100)
#Final parameters
finalMinDist = 0.5
finalNN = 50
#Clustering resolution
clusterRes=1.0
#Populations of genes to exclude from the analysis
excludeGenes = unlist(as.list(geneSets)[c('mtGenes','hspGenes','riboGenes')],use.name=FALSE)
excludeGenes = sort(unique(excludeGenes))
#Number of PCs to use for quick analysis during QC
defNumPCs=75
#Gene map
gnsMap = read.table(file.path(srcs$fAd[[1]][1],'filtered_gene_bc_matrices','GRCh38','genes.tsv'),sep='\t',header=FALSE)
#Cycle genes
ccGns = cc.genes.updated.2019
ccGns = unlist(ccGns)
ccGns = data.frame(symbol = ccGns,
                   phase = gsub('\\..*','',names(ccGns)))
ccGns$geneID = gnsMap[match(ccGns$symbol,gnsMap[,2]),1]
ccGns = ccGns[!is.na(ccGns$geneID),]
#Use a cut-off on average expression to be a bit more conservative


#########
# SoupX #
#########
#Check if it's already been done
for(ex in names(srcs)){
  message(sprintf("Running decontamination for experiment %s",ex))
  #Can't do soup removal on bone marrow
  if(ex=='mBM'){
    next
    #But we should split it out into separate channels.
    #Read it in
    out = readMM(file.path(srcs[['mBM']],'ADULT_BM_RAW.mtx'))
    out = t(out)
    tmp = read.table(file.path(srcs[['mBM']],'ADULT_BM_cellNames.csv'))
    colnames(out) = tmp[,1]
    #Fix up labels
    colnames(out) = gsub('(HiSeq_[0-9]+)-','\\1_',colnames(out))
    colnames(out) = gsub('-BM_adult','',colnames(out))
    tmp = read.table(file.path(srcs[['mBM']],'ADULT_BM_genes.csv'))
    rownames(out) = tmp[,1]
    #Split it into channels for subsequent processing
    channels = gsub('_[ACGT]+.*','',colnames(out))
    for(channel in unique(channels)){
      tgt = file.path(outDir,'strained',ex,channel)
      message(sprintf("Splitting out channel %s",tgt))
      write10xCounts(tgt,out[,channels==channel],overwrite=TRUE)
    }
  }
  for(dSrc in names(srcs[[ex]])){
    message(sprintf("From source %s",dSrc))
    if(dSrc=='fKidPlos'){
      srcFiles = list.files(srcs[[ex]][[dSrc]],full.names=TRUE)
      for(wk in c('09','11','13','16','18')){
        lab=paste0('fKidPlosWk',wk)
        message(sprintf("Running week %s",wk))
        #Load matrix
        dat = readMM(grep(paste0('wk',wk,'_matrix.mtx'),srcFiles,value=TRUE))
        colnames(dat) = read.table(grep(paste0('wk',wk,'_barcodes.tsv'),srcFiles,value=TRUE),sep='\t',header=FALSE)[,1]
        rownames(dat) = read.table(grep(paste0('wk',wk,'_genes.tsv'),srcFiles,value=TRUE),sep='\t',header=FALSE)[,1]
        #And the real cells
        annTab = read.table(grep(paste0('w',wk,'_barcodes_celltypes'),srcFiles,value=TRUE),sep=',',header=TRUE)
        #Run SoupX if needed
        if(ncol(dat)<100000){
          out=dat[,annTab[,1]]
        }else{
          #Now do soup calculation
          #Have to cluster these things
          pdf(file.path(outDir,sprintf('%s_%s_SoupX_processing.pdf',ex,lab)),useDingbats=FALSE)
          toc = dat[,annTab[,1]]
          srat = quickCluster(toc,clusteringRes=0.8)
          sc = SoupChannel(dat,toc)
          sc = setClusters(sc,srat@active.ident)
          sc = autoEstCont(sc)
          out = adjustCounts(sc,verbose=2)
          dev.off()
        }
        #Save it
        tgt = file.path(outDir,'strained',ex,lab)
        dir.create(tgt,recursive=TRUE,showWarnings=FALSE)
        write10xCounts(tgt,out,overwrite=TRUE)
      }
    }else{
      for(lab in names(srcs[[ex]][[dSrc]])){
        tmp = sprintf("Processing channel %s",lab)
        tmp = paste0(paste(rep('#',nchar(tmp)+3),collapse=''),'\n',tmp)
        message(tmp)
        pdf(file.path(outDir,sprintf('%s_%s_SoupX_processing.pdf',ex,lab)),useDingbats=FALSE)
        sc = load10X(srcs[[ex]][[dSrc]][[lab]])
        if(!'clusters' %in% colnames(sc$metaData)){
          message("No clusters found, running Seurat.")
          srat = scQC$quickCluster(sc$toc,clusteringRes=0.8)
          sc = setClusters(sc,srat@meta.data$seurat_clusters[match(colnames(sc$toc),rownames(srat@meta.data))])
        }
        #For whatever reason, late Adrenals different (V3?) gives 0, manually set to 0.1 to err on the safe side.
        if(FALSE && dSrc=='lateAdrenals'){
          tmp = setContaminationFraction(sc,0.1)
        }else{
          tmp = tryCatch(autoEstCont(sc,forceAccept=TRUE,tfidfMin=def),
                         error = function(c) NULL,
                         warning = function(c) NULL)
          if(is.null(tmp)){
            message("Default failed, reducing tfidf to 0.8")
            tmp = tryCatch(autoEstCont(sc,tfidfMin=0.8,forceAccept=TRUE),
                           error = function(c) NULL,
                           warning = function(c) NULL)
            if(is.null(tmp)){
              message("Still failed, reducing tfidf to 0.6")
              tmp = tryCatch(autoEstCont(sc,tfidfMin=0.6,forceAccept=TRUE),
                                   error = function(c) NULL,
                                   warning = function(c) NULL
                                   )
              if(is.null(tmp)){
                message("Still failed, defaulting to 0.1")
                tmp = setContaminationFraction(sc,0.1)
              }
            }
          }
        }
        sc = tmp
        out = adjustCounts(sc,verbose=2)
        dev.off()
        #Save it
        tgt = file.path(outDir,'strained',ex,lab)
        dir.create(tgt,recursive=TRUE,showWarnings=FALSE)
        write10xCounts(tgt,out,overwrite=TRUE)
      }
    }
  }
}
strained = lapply(names(srcs),function(e) {
                    x=list.files(file.path(outDir,'strained',e),full.names=TRUE)
                    names(x) = basename(x)
                    x})
names(strained) = names(srcs)
#Drop samples too shit to bother with 
strained$mKid = strained$mKid[!names(strained$mKid) %in% c('Wilms3_Ure_N_ldc_1_1','Wilms2_Ure_N_ldc_1_1','RCC3_Ure_N_ldc_1_1','RCC1_Ure_N_ldc_1_1')]
#Duplicate mKid to make kidney immune profile
strained$mKidImm = strained$mKid

############
# Basic QC #
############
srats = list()
for(ex in names(strained)){
  tgtFile = file.path(outDir,sprintf('baseSeurat_%s.RDS',ex))
  if(file.exists(tgtFile)){
    srat = readRDS(tgtFile)
  }else{
    #Tweaked cut-offs for different data sets
    mtLoc = mtCut
    scrubLoc = 0.2
    umiLoc = minNumUMI
    if(ex=='fKid'){
      mtLoc = 0.2
      scrubLoc = 0.2
    }
    if(ex %in% c('mKidImm','mKid')){
      mtLoc = 0.4
      scrubLoc = 0.3
      umiLoc = 300
    }
    if(ex=='mBM'){
      mtLoc = 0.1
    }
    #Create the basic Seurat object for each experiment
    message(sprintf("QC filters on %s",ex))
    pdf(file.path(outDir,sprintf('%s_QC.pdf',ex)))
      srat = scQC$basicQC(dataDirs=strained[[ex]],
                          excludeGenes = excludeGenes,
                          geneSets = geneSets,
                          maxMT = mtLoc,
                          minGenes = minNumGenes,
                          minUMIs = umiLoc,
                          maxBadFrac = maxScrubFrac,
                          scrubScoreMax = scrubLoc,
                          numPCs = defNumPCs)
    dev.off()
    #####################################################
    # Special droping of things that should be excluded 
    if(ex=='mKidImm'){
      #Make a copy that is 
      #Extra things for mature kidney
      baseDir = 'Data/scienceKidneyAnalysis/'
      cMani = read.table('Data/TableS2.tsv',sep='\t',header=TRUE)
      comps = unique(cMani$Data_set)
      srat@meta.data$sciClust=NA
      srat@meta.data$sciAlias=NA
      for(comp in comps){
        mDat = readRDS(file.path(baseDir,comp,'batchCorrected','metadata.RDS'))$meta.data
        rownames(mDat) = paste0(mDat$Sanger_study_ID,'___',gsub('^[0-9]+_','',rownames(mDat)))
        #Pull cluster ID
        tmp = cMani[cMani$Data_set==comp,]
        m = match(mDat$res.1,gsub('[A-Z]*','',tmp$Cluster_ID))
        mDat$ClusterID = tmp$Cluster_ID[m]
        mDat$Alias = tmp$Alias[m]
        #Map these onto our data
        m = match(paste0(mDat$label,'_',gsub('.*___','',rownames(mDat))),rownames(srat@meta.data))
        w = !is.na(m)
        srat@meta.data$sciClust[m[w]] = mDat$ClusterID[w]
        srat@meta.data$sciAlias[m[w]] = mDat$Alias[w]
      }
      #Now use this to train model and allocate clusters
      refClass = srat@meta.data$sciAlias
      #Keep only the sensible, immune classes
      w = unique(refClass[grep('IN',srat@meta.data$sciClust)])
      w = refClass %in% w[w!='-']
      mDat = srat@meta.data[w,]
      misc = srat@misc
      srat = CreateSeuratObject(srat@assays$RNA@counts[,w])
      srat@meta.data = cbind(srat@meta.data,mDat[,!(colnames(mDat) %in% colnames(srat@meta.data)),drop=FALSE])
      srat@misc = misc
      srat = NormalizeData(srat,verbose=FALSE)
      srat = FindVariableFeatures(srat,verbose=FALSE)
    }
    if(ex=='mKid'){
      #Extra things for mature kidney
      baseDir = 'Data/scienceKidneyAnalysis/'
      cMani = read.table('Data/TableS2.tsv',sep='\t',header=TRUE)
      comps = unique(cMani$Data_set)
      srat@meta.data$sciClust=NA
      srat@meta.data$sciAlias=NA
      for(comp in comps){
        mDat = readRDS(file.path(baseDir,comp,'batchCorrected','metadata.RDS'))$meta.data
        rownames(mDat) = paste0(mDat$Sanger_study_ID,'___',gsub('^[0-9]+_','',rownames(mDat)))
        #Pull cluster ID
        tmp = cMani[cMani$Data_set==comp,]
        m = match(mDat$res.1,gsub('[A-Z]*','',tmp$Cluster_ID))
        mDat$ClusterID = tmp$Cluster_ID[m]
        mDat$Alias = tmp$Alias[m]
        #Map these onto our data
        m = match(paste0(mDat$label,'_',gsub('.*___','',rownames(mDat))),rownames(srat@meta.data))
        w = !is.na(m)
        srat@meta.data$sciClust[m[w]] = mDat$ClusterID[w]
        srat@meta.data$sciAlias[m[w]] = mDat$Alias[w]
      }
      #Now use this to train model and allocate clusters
      refClass = srat@meta.data$sciAlias
      #Keep only the sensible, non-immune classes
      w = refClass %in% c('AV1','AV2','C1','C2','D','DV','G','GE','H','M','PT1','PT3')
      mDat = srat@meta.data[w,]
      misc = srat@misc
      srat = CreateSeuratObject(srat@assays$RNA@counts[,w])
      srat@meta.data = cbind(srat@meta.data,mDat[,!(colnames(mDat) %in% colnames(srat@meta.data)),drop=FALSE])
      srat@misc = misc
      srat = NormalizeData(srat,verbose=FALSE)
      srat = FindVariableFeatures(srat,verbose=FALSE)
    }
    saveRDS(srat,tgtFile)
  }
  srats[[ex]]=srat
}

################
# Drop cycling #
################
for(ex in names(strained)){
  tgtFile = file.path(outDir,sprintf('seuratG0_%s.RDS',ex))
  if(file.exists(tgtFile)){
    srat = readRDS(tgtFile)
  }else{
    srat = srats[[ex]]
    if(ex %in% c('mKidImm','mKid','PBMC3k')){
      #The auto-estimating thing doesn't work well here, so manually specify which to drop
      tmp = c('MKI67','TOP2A','UBE2C','RRM2')
      w = which(colSums(srat@assays$RNA@counts[tmp,])==0)
    }else{
      #Aggressively filter out things
      w = which(srat@meta.data$S.Score<0 & srat@meta.data$G2M.Score<0)
    }
    mDat = srat@meta.data[w,]
    srat = CreateSeuratObject(srat@assays$RNA@counts[,w])
    srat@meta.data = cbind(srat@meta.data,mDat[,!(colnames(mDat) %in% colnames(srat@meta.data)),drop=FALSE])
    srat = NormalizeData(srat,verbose=FALSE)
    srat = FindVariableFeatures(srat,verbose=FALSE)
    saveRDS(srat,tgtFile)
  }
  srats[[ex]]=srat
}

###################
# Drop Leucocytes #
###################
#This is a bit tricky.  Want to drop CD45+ but also erythroblasts, megakaryoctes, which are CD45-.  Use HBB/HBA1 and ITGA2B/ITGB3 as markers for these.
for(ex in names(strained)){
  #Don't want to drop leucocytes from immune reference
  if(ex %in% c('PBMC','mBM','mKidImm','PBMC3k'))
    next
  tgtFile = file.path(outDir,sprintf('noLeucotyesG0_%s.RDS',ex))
  if(file.exists(tgtFile)){
    srat = readRDS(tgtFile)
  }else{
    srat = srats[[ex]]
    #Need to do this at the cluster level.
    tmp = quickCluster(srat@assays$RNA@counts,clusteringRes=0.8)
    #Get clusters
    clusts = tmp@active.ident
    leucFrac = sapply(split(tmp@assays$RNA@counts['PTPRC',]>=1,clusts),mean)
    #These two are sometimes CD45-
    bFrac = sapply(split(tmp@assays$RNA@counts['MS4A1',]>=1,clusts),mean)
    mgkFrac = sapply(split(tmp@assays$RNA@counts['ITGA2B',]>=1,clusts),mean)
    #Use geneset for RBC
    tmp@meta.data$hgGenes = srat@meta.data$hgGenes[match(rownames(tmp@meta.data),rownames(srat@meta.data))]
    rbcFrac = sapply(split(tmp@meta.data$hgGenes,clusts),mean)
    #Keep only the things that aren't erythroblasts, leucocytes, or megakaryctes
    toKeep = leucFrac<0.3 & rbcFrac<0.1 & mgkFrac<0.5 & bFrac<0.5
    if(ex=='fKid'){
      toKeep = leucFrac<0.15 & rbcFrac<0.1 & mgkFrac<0.5 & bFrac<0.5
    }
    toKeep = as.character(tmp@active.ident) %in% names(toKeep)[toKeep]
    #Also drop scattered ones with stupid hgFrac
    toKeep = toKeep & tmp@meta.data$hgGenes<0.1
    tmp@meta.data$toKeep=toKeep
    mDat = srat@meta.data[toKeep,]
    srat = CreateSeuratObject(srat@assays$RNA@counts[,toKeep])
    srat@meta.data = cbind(srat@meta.data,mDat[,!(colnames(mDat) %in% colnames(srat@meta.data)),drop=FALSE])
    srat = NormalizeData(srat,verbose=FALSE)
    srat = FindVariableFeatures(srat,verbose=FALSE)
    saveRDS(srat,tgtFile)
  }
  srats[[ex]]=srat
}

################
# Optimal nPCs #
################
#Work out what the best number of PCs is
mcvs = list()
nPCs = c()
for(ex in names(strained)){
  tgtFile=file.path(outDir,sprintf('MCV_%s.RDS',ex))
  srat = srats[[ex]]
  if(file.exists(tgtFile)){
    mcv = readRDS(tgtFile)
  }else{
    mcv = molecularCrossValidation(roundToInt(srat@assays$RNA@counts),VariableFeatures(srat),normalisation=stdProcess,features=VariableFeatures(srat),tFracs=c(1,0.9,0.8,0.5),nSplits=4)
    pdf(file.path(outDir,paste0(ex,'_mcv.pdf')))
      plotMCV(mcv)
    dev.off()
    saveRDS(mcv,tgtFile)
  }
  mcvs[[ex]] = mcv
  nPCs[ex] = max(mcv$mins1sd[[1]])
  #nPCs = sapply(mcvList,function(e) max(e$mins1sd[[1]]))
}


####################
# Final processing #
####################
for(ex in names(strained)){
  tgtFile = file.path(outDir,sprintf('processedSeurat_%s.RDS',ex))
  if(file.exists(tgtFile)){
    srat = readRDS(tgtFile)
  }else{
    srat = srats[[ex]]
    pdf(file.path(outDir,sprintf('standardPlots_%s.pdf',ex)),width=24,height=24)
      srat = ScaleData(srat)
      srat = RunPCA(srat,npcs=nPCs[ex],approx=FALSE)
      #Run tSNE as well in case that turns out to be useful.
      #srat = RunTSNE(srat)
      #The old python based method seems to segfault every time...
      srat = FindNeighbors(srat,dims=seq(nPCs[ex]))
      srat = FindClusters(srat,res=clusterRes)
      #Make a range of UMAP plots
      #umaps=list()
      #for(minDist in minDists){
      #  for(NN in NNs){
      #    message(sprintf("Calculating UMAP for minDist=%g, NN=%d",minDist,NN))
      #    srat = RunUMAP(srat,dims=seq(nPCs[ex]),min.dist=minDist,n.neighbors=NN,verbose=FALSE)
      #    umaps[[paste0(minDist,'_',NN)]] = DimPlot(srat,label=TRUE) +
      #      guides(colour=FALSE) +
      #      ggtitle(sprintf("UMAP with minDist=%g, NN=%d",minDist,NN))
      #  }
      #}
      #plot(plot_grid(plotlist=umaps,
      #          nrow=length(minDists),
      #          ncol=length(NNs)))
      #Final UMAP
      srat = RunUMAP(srat,dims=seq(nPCs[ex]),min.dist=finalMinDist,n.neighbors=finalNN)
      #Clustering
      plot(DimPlot(srat,label=TRUE))
      #My ones
      plot(FeaturePlot(srat,c('PTPRC','CD74','PDGFRB','EPCAM','PECAM1','HBB','ITGA2B')))
      #Source
      plot(DimPlot(srat,group.by='orig.ident'))
      #Cell cycle
      plot(DimPlot(srat,group.by='Phase'))
      #QC things
      plot(FeaturePlot(srat,c('scrubScore','nFeature_RNA','nCount_RNA',names(geneSets))))
      #Visualise cell cycle
      #plot(FeaturePlot(srat,c('PCNA','MKI67','MCM4','CDKN1C')))
      #Genes specific to Steroidogenisis
      #plot(FeaturePlot(srat,c('MC2R','CYP17A1','MRAP','STAR','CYP11B2','CYP11B1','INHA','GNRHR','HSD3B2')))
      #Medulla markers
      #plot(FeaturePlot(srat,c('PHOX2A','ISL1','CHRNA3','SOX2','SOX10','ERBB3','DBH','CHGB','TH')))
      #Others
      #plot(FeaturePlot(srat,c('NR5A1','NOV','HSD3B2','SULT2A1','FDX1')))
    dev.off()
    saveRDS(srat,tgtFile)
  }
  srats[[ex]] = srat
}

###################
# Post processing #
###################
for(ex in names(strained)){
  tgtFile = file.path(outDir,sprintf('finalisedSeurat_%s.RDS',ex))
  if(file.exists(tgtFile)){
    srat = readRDS(tgtFile)
  }else{
    srat = srats[[ex]]
    if(ex=='fKid'){
      #Extra things for foetal kidney
      srat@meta.data$channel = gsub('_[ACGT]+.?.?$','',rownames(srat@meta.data))
      nom = srat@meta.data$channel
      srat@meta.data$sort = ifelse(grepl('_neg_',nom) | grepl('45N',nom),
                                   'CD45-',
                                   ifelse(grepl('_pos_',nom) | grepl('45P',nom),
                                          'CD45+',
                                          'None'
                                          )
                                   )
      tmp = gsub('_.*','',nom)
      srat@meta.data$age = ifelse(grepl('fKidPlos',tmp),
                                  as.numeric(gsub('fKidPlosWk','',tmp)),
                                  c(F15=12,
                                    F16=8+1/7,
                                    F17=9+1/7,
                                    F35=7+6/7,
                                    F38=13+6/7,
                                    F41=16+6/7,
                                    F45=13+6/7)[tmp]
                                  )
      srat@meta.data$gest = cut(srat@meta.data$age,breaks=c(0,10,14,30))
      #Add in the annotation
      #Reload Plos data and fit model.
      srcDir = 'Data/scData/fKidPlos'
      tgts = grep('matrix.mtx',list.files(srcDir),value=TRUE)
      mtxs = list()
      for(tgt in tgts){
        #Load the matrix
        mtx = readMM(file.path(srcDir,tgt))
        colnames(mtx) = read.table(file.path(srcDir,gsub('_matrix.mtx','_barcodes.tsv',tgt)),sep='\t',header=FALSE)[,1]
        gns = read.table(file.path(srcDir,gsub('_matrix.mtx','_genes.tsv',tgt)),sep='\t',header=FALSE)
        rownames(mtx) = gns[,2]
        #Load the meta-data
        mDat = read.table(file.path(srcDir,gsub('.*_wk?([0-9]+)_.*','w\\1_barcodes_celltypes.csv',tgt)),sep=',',header=TRUE)
        #Keep only the informative ones
        mtx = mtx[,match(mDat[,1],colnames(mtx))]
        #Add everything to header and make things unique
        samp = gsub('.*_wk?([0-9]+)_.*','w\\1',tgt)
        print(samp)
        print(table(gsub('.*-','',colnames(mtx))))
        colnames(mtx) = paste0(gsub('/','.',mDat[,2]),':',samp,'___',colnames(mtx))
        mtxs[[tgt]]=mtx
      }
      mtx = do.call(cbind,mtxs)
      t2 = file.path(outDir,'lr_fKid.RDS')
      if(file.exists(t2)){
        lr_fKid = readRDS(t2)
      }else{
        lr_fKid = lr$trainModel(mtx,classes=gsub(':.*','',colnames(mtx)))
        saveRDS(lr_fKid,t2)
      }
      out = lr$predictSimilarity(lr_fKid,srat@assays$RNA@counts)
      #Get the most likely annotation
      annot = colnames(out)[apply(out,1,which.max)]
      #Set those with only moderate evidence to NA
      annot[apply(out,1,max)<1]=NA
      #And those with multiple annotations.
      annot[rowSums(out>3)>1] = NA
      #Finish up
      srat@meta.data$annot = annot
      #Full annotation.  Standardise on clusters
      srat@meta.data$fullAnnot = annot
      srat@meta.data$annot[grep('NPC[abcd]',srat@meta.data$annot)]='NPC'
      srat@meta.data$annot[grep('RVCSB[ab]',srat@meta.data$annot)]='RVCSB'
      srat@meta.data$annot[srat@meta.data$annot%in% c('Leu','IPC','Mes','PTA')]=NA
      tmp = table(srat@meta.data$annot,srat@meta.data$seurat_clusters)
      class(tmp) = 'matrix'
      tmp = setNames(rownames(tmp)[apply(tmp,2,which.max)],colnames(tmp))
      srat@meta.data$annot = tmp[srat@meta.data$seurat_clusters]
      srat@meta.data$annot[srat@meta.data$seurat_clusters=='12']='MSC'
    }
    if(ex=='mKid'){
      ####Extra things for mature kidney
      ###baseDir='~/trueHome/scratch/Plots/KidneySC/analysis20171204/'
      ###cMani = read.table('~/trueHome/scratch/Plots/KidneySC/Paper/Science_R1/TableS2.tsv',sep='\t',header=TRUE)
      ###comps = unique(cMani$Data_set)
      ###srat@meta.data$sciClust=NA
      ###srat@meta.data$sciAlias=NA
      ###for(comp in comps){
      ###  mDat = readRDS(file.path(baseDir,comp,'batchCorrected','metadata.RDS'))$meta.data
      ###  rownames(mDat) = paste0(mDat$Sanger_study_ID,'___',gsub('^[0-9]+_','',rownames(mDat)))
      ###  #Pull cluster ID
      ###  tmp = cMani[cMani$Data_set==comp,]
      ###  m = match(mDat$res.1,gsub('[A-Z]*','',tmp$Cluster_ID))
      ###  mDat$ClusterID = tmp$Cluster_ID[m]
      ###  mDat$Alias = tmp$Alias[m]
      ###  #Map these onto our data
      ###  m = match(paste0(mDat$label,'_',gsub('.*___','',rownames(mDat))),rownames(srat@meta.data))
      ###  w = !is.na(m)
      ###  srat@meta.data$sciClust[m[w]] = mDat$ClusterID[w]
      ###  srat@meta.data$sciAlias[m[w]] = mDat$Alias[w]
      ###}
      ####Now use this to train model and allocate clusters
      ###refClass = srat@meta.data$sciAlias
      ###w = (!refClass %in% c('-','NK2','F','U2','U1')) & !is.na(refClass)
      ####Drop things with fewer than 10
      ###w[w] = table(refClass[w])[refClass[w]]>10
      ###t2 = file.path(outDir,'lr_mKid.RDS')
      ###if(file.exists(t2)){
      ###  lrKid = readRDS(t2)
      ###}else{
      ###  lrKid = lr$trainModel(srat@assays$RNA@counts[,w],classes=refClass[w],maxCells=Inf)
      ###  saveRDS(lrKid,t2)
      ###}
      ###out = lr$predictSimilarity(lrKid,srat@assays$RNA@counts)
      ####Convert to probabilities for averaging
      ###tmp = 1/(1+exp(-out))
      ###tmp = do.call(rbind,lapply(split(names(srat@active.ident),srat@active.ident),function(e) -log(1/colMeans(tmp[e,,drop=FALSE])-1)))
      ###pdf(file.path(outDir,sprintf('finalisingPlots_%s.pdf',ex)))
      ###  hh = lr$similarityHeatmap(tmp)
      ###  draw(hh)
      ###dev.off()
      srat@meta.data$annot = srat@meta.data$sciAlias
      srat@meta.data$fullAnnot = srat@meta.data$annot
    }
    if(ex=='mBM'){
      #Extra things for mature Bone Marrow data
      annot = read.table('Data/scData/mBM/ADULT_BM_cellLabels.csv',sep=',')
      colnames(annot) = 'Label'
      annot$CellID = read.table('Data/scData/mBM/ADULT_BM_cellNames.csv',sep=',')[,1]
      annot$CellID = gsub('(HiSeq_[0-9]+)-','\\1_',gsub('-BM_adult','',annot$CellID))
      annot$CellID = gsub('-1$','',annot$CellID)
      m = match(sub('MantonBM[0-9]+_HiSeq_[0-9]+_','',rownames(srat@meta.data)),annot$CellID)
      srat@meta.data$annot = annot$Label[m]
    }
    if(ex=='fAd'){
      tmp = AverageExpression(srat)$RNA
      markers = list(SCPs=c('SOX10','MPZ','PLP1','ERBB3'),
                     Bridge = 'DLL3',
                     Sympathoblastic = c('GAP43','BCL2','NPY'),
                     Chromaffin = 'PNMT',
                     Mesenchyme = c('PDGFRB','TCF21'),
                     Endothelium = c('PLVAP','PECAM1','KDR','PTPRB'),
                     Cortex = c('STAR','MC2R'))
      annot = do.call(rbind,lapply(markers,function(e) colMeans(t(scale(t(tmp[e,,drop=FALSE]))))))
      t2 = apply(annot,2,max)
      t3 = apply(annot,2,which.max)
      annot = ifelse(t2>0,
                     rownames(annot)[t3],
                     'Cortex')
      srat@meta.data$annot = annot[srat@meta.data$seurat_clusters]
    }
    if(ex=='PBMC'){
      srat@meta.data$annot=NA
      srat@meta.data$annot[srat@meta.data$seurat_clusters=='0']='NK'
      srat@meta.data$annot[srat@meta.data$seurat_clusters=='12']='NKT'
      srat@meta.data$annot[srat@meta.data$seurat_clusters=='5']='B'
      srat@meta.data$annot[srat@meta.data$seurat_clusters=='2']='T_CD8'
      srat@meta.data$annot[srat@meta.data$seurat_clusters=='7']='T_CD4'
      srat@meta.data$annot[srat@meta.data$seurat_clusters=='16']='B_mature'
      srat@meta.data$annot[srat@meta.data$seurat_clusters%in% c(10,1,9)]='T'
      srat@meta.data$annot[srat@meta.data$seurat_clusters%in% c(10,1,9)]='T'
      srat@meta.data$annot[srat@meta.data$seurat_clusters%in% c(15,11,8,4,6,3,13,14)]='MNP'
    }
    if(ex=='mKidImm'){
      tmp = srat@meta.data$sciAlias
      tmp[grep('MNP',tmp)]='MNP'
      tmp[grep('NKT',tmp)]='NKT'
      tmp[grep('NK[0-9]',tmp)]='NK'
      tmp[tmp=='8T']='Tcyto'
      tmp[tmp %in% c('NP')]=NA
      srat@meta.data$annot = tmp
    }
    if(ex=='PBMC3k'){
      tmp = read.table('Data/PBMCRef_columnNames.tsv',header=FALSE)$V1
      m = match(gsub('.*_','',colnames(srat)),gsub('.*:','',tmp))
      srat@meta.data$annot = gsub(':.*','',tmp)[m]
      #Fix up the NAs
      tmp = table(srat@meta.data$seurat_clusters,srat@meta.data$annot)
      tmp = setNames(colnames(tmp)[apply(tmp,1,which.max)],rownames(tmp))
      w = is.na(srat@meta.data$annot) | srat@meta.data$annot=='PBMC_Megakaryocytes'
      srat@meta.data$annot[w] = tmp[as.character(srat@meta.data$seurat_clusters[w])]
      srat@meta.data$annot = gsub('PBMC_','',srat@meta.data$annot)
    }
    saveRDS(srat,tgtFile)
  }
  srats[[ex]]=srat
}

########################
# Non standard samples #
########################
##########
# sciRef #
##########
ex='sciKidRef'
tgtFile = file.path(outDir,sprintf('finalisedSeurat_%s.RDS',ex))
if(file.exists(tgtFile)){
  srat = readRDS(tgtFile)
}else{
  srat = loadRefMtx('~/trueHome/scratch/scSignatures/sciKidRef')
  srat = scQC$quickCluster(srat)
  srat = CellCycleScoring(srat, s.features=ccGns$geneID[ccGns$phase=='s'],g2m.features=ccGns$geneID[ccGns$phase=='g2m'],seed=sample(1e9,1))
  #Redo without cycing cells
  #Clean up colnames
  srat = srat@assays$RNA@counts[,srat@meta.data$Phase=='G1']
  annot = gsub('sciKidRef_(.*):.*','\\1',colnames(srat))
  colnames(srat) = gsub('.*:','',colnames(srat))
  srat = scQC$quickCluster(srat)
  srat@meta.data$annot = annot
  saveRDS(srat,tgtFile)
}
srats[[ex]]=srat
##########
# Immune #
##########
ex='PBMC'
tgtFile = file.path(outDir,sprintf('finalisedSeurat_%s.RDS',ex))
if(file.exists(tgtFile)){
  srat = readRDS(tgtFile)
}else{
  srat = loadRefMtx('~/trueHome/scratch/scSignatures/PBMCRef')
  srat = scQC$quickCluster(srat)
  srat = CellCycleScoring(srat, s.features=ccGns$geneID[ccGns$phase=='s'],g2m.features=ccGns$geneID[ccGns$phase=='g2m'],seed=sample(1e9,1))
  srat@meta.data$Phase = ifelse(srat@meta.data$G2M.Score>0.2,
                                'G2M',
                                ifelse(srat@meta.data$S.Score>0.2,
                                       'S',
                                       'G1'
                                       )
                                )
  #Redo without cycing cells
  #Clean up colnames
  srat = srat@assays$RNA@counts[,srat@meta.data$Phase=='G1']
  annot = gsub('PBMC_(.*):.*','\\1',colnames(srat))
  colnames(srat) = gsub('.*:','',colnames(srat))
  srat = scQC$quickCluster(srat)
  srat@meta.data$annot = annot
  saveRDS(srat,tgtFile)
}
srats[[ex]] = srat
########################
# Human Cell Landscape #
########################
ex='HCL'
tgtFile = file.path(outDir,sprintf('finalisedSeurat_%s.RDS',ex))
if(file.exists(tgtFile)){
  srat = readRDS(tgtFile)
}else{
  #######
  # Load
  hfile = hdf5r::h5file(filename = 'Data/scData/humanCellLandscape/HCL_final_USE.h5ad', mode = "r")
  #Need some complicated sequential readthrough to prevent memory exhaustion.  Data can be loaded with standard indexing.
  tgt = hfile[['raw.X']]
  chunkSize=10000
  out = list()
  totN = length(tgt[1,])
  totG = length(tgt[,1])
  i=0
  while(i*chunkSize<totN){
    print(i*chunkSize)
    x = tgt[,(i*chunkSize+1):min(totN,((i+1)*chunkSize))]
    w = which(x!=0,arr.ind=TRUE)
    out[[length(out)+1]] = data.frame(i = w[,1],
                                      j = (i*chunkSize)+w[,2],
                                      x=x[w])
    i=i+1
  }
  out = do.call(rbind,out)
  #Get the dimension names
  mDat = hfile[['obs']][1:totN]
  #Add the reduced dim coordinates
  mDat$tSNE1 = hfile[['obsm']][['X_tsne']][1,]
  mDat$tSNE2 = hfile[['obsm']][['X_tsne']][2,]
  mDat$UMAP1 = hfile[['obsm']][['X_umap']][1,]
  mDat$UMAP2 = hfile[['obsm']][['X_umap']][2,]
  mDat$annot = gsub('\\.[ACGT]+.*','',mDat$index)
  #Get gene information
  gDat = hfile[['raw.var']][1:totG]
  gns = read.table(file.path(srcs$fAd[[1]][1],'filtered_gene_bc_matrices','GRCh38','genes.tsv'),sep='\t',header=FALSE)
  m = match(gDat$index,gns$V2)
  gDat$ENS = gns$V1[m]
  out = sparseMatrix(i=out$i,
                     j=out$j,
                     x=expm1(out$x),
                     dims=c(totG,totN),
                     dimnames=list(gDat$index,mDat$index)
  )
  #Make Seurat
  srat = CreateSeuratObject(out)
  srat = NormalizeData(srat)
  srat@meta.data = cbind(srat@meta.data,mDat)
  srat@meta.data$devel = ifelse(grepl('Adult',srat@meta.data$annot),
                               'Adult',
                               ifelse(grepl('Fetal',srat@meta.data$annot),
                                      'Fetal',
                                      'Other'
                                      )
                               )
  srat@misc$geneMap = gDat
  #Force in dim reduction
  srat[['tsne']] = CreateDimReducObject(embeddings = as.matrix(srat@meta.data[,c('tSNE1','tSNE2')]),key='tsne',global=TRUE)
  srat[['umap']] = CreateDimReducObject(embeddings = as.matrix(srat@meta.data[,c('UMAP1','UMAP2')]),key='umap',global=TRUE)
  saveRDS(srat,tgtFile)
}
srats[[ex]]=srat
#################
# Marioni Mouse #
#################
ex='mouseDev'
tgtFile = file.path(outDir,sprintf('finalisedSeurat_%s.RDS',ex))
if(file.exists(tgtFile)){
  srat = readRDS(tgtFile)
}else{
  #######
  # Load
  dat = readMM('Data/scData/mouseDevMarioni/raw_counts.mtx')
  colnames(dat) = read.table('Data/scData/mouseDevMarioni/barcodes.tsv',sep='\t',header=FALSE)$V1
  gns = read.table('Data/scData/mouseDevMarioni/genes.tsv',sep='\t',header=FALSE)
  mDat = read.table('Data/scData/mouseDevMarioni/meta.tab',sep='\t',header=TRUE)
  #Get orthologs
  mart1 = useMart("ensembl", dataset="hsapiens_gene_ensembl")
  mart2 = useMart("ensembl", dataset="mmusculus_gene_ensembl")
  gnMap = getLDS(attributes=c("ensembl_gene_id"),
                 filters="ensembl_gene_id", 
                 values=gns$V1, 
                 mart=mart2,
                 attributesL=c("ensembl_gene_id"), 
                 martL=mart1)
  colnames(gnMap) = c('mouseID','humanID')
  gnMap$ambiguous = table(gnMap$humanID)[gnMap$humanID]>1 | table(gnMap$mouseID)[gnMap$mouseID]>1
  #Keep only the ones that we can map unambiguously and are in the Seurat annotation
  m = match(gns$V1,gnMap$mouseID)
  gns$humanID = ifelse(!is.na(m) & !gnMap$ambiguous[m], gnMap$humanID[m],NA)
  tmp = read.table('Data/scData/fAd/babyAdrenals/babyAdrenal1/filtered_gene_bc_matrices/GRCh38/genes.tsv',sep='\t',header=FALSE)
  gns$humanSym = tmp$V2[match(gns$humanID,tmp$V1)]
  #Keep and make seurat object
  cnts = dat[!is.na(gns$humanSym),]
  #Label by symbol
  rownames(cnts) = make.unique(gns$humanSym[!is.na(gns$humanSym)])
  #Make seurat object
  cellKeep = !is.na(mDat$celltype)
  cnts = cnts[,cellKeep]
  mDat = mDat[cellKeep,]
  srat = scQC$quickCluster(cnts,numPCs=100)
  srat@misc$geneMap = gns
  srat@misc$geneOrthologs = gnMap
  #Add meta-data
  srat@meta.data = cbind(srat@meta.data,mDat[,-match(c('cell'),colnames(mDat))])
  srat = SetIdent(srat,value='celltype')
  srat@meta.data$annot = srat@meta.data$celltype
  saveRDS(srat,tgtFile)
}
srats[[ex]]=srat
#####################
# Mouse Dev CellSeq #
#####################
ex='mouseDev_extra'
tgtFile = file.path(outDir,sprintf('finalisedSeurat_%s.RDS',ex))
if(file.exists(tgtFile)){
  srat = readRDS(tgtFile)
}else{
  #######
  # Load
  mDat = read.table('Data/Mouse_CCSK/metaData_Mm.txt',sep='\t',header=TRUE)
  dat = read.table('Data/Mouse_CCSK/normData_Mm.txt',sep='\t',header=TRUE)
  rownames(dat) = dat[,1]
  dat = dat[,-1]
  #Get orthologs
  gns = data.frame(mouseID = gsub('__.*','',rownames(dat)),
                   mouseSym = gsub('.*__','',rownames(dat)))
  mart1 = useMart("ensembl", dataset="hsapiens_gene_ensembl")
  mart2 = useMart("ensembl", dataset="mmusculus_gene_ensembl")
  gnMap = getLDS(attributes=c("ensembl_gene_id"),
                 filters="ensembl_gene_id", 
                 values=gns$mouseID, 
                 mart=mart2,
                 attributesL=c("ensembl_gene_id"), 
                 martL=mart1)
  colnames(gnMap) = c('mouseID','humanID')
  gnMap$ambiguous = table(gnMap$humanID)[gnMap$humanID]>1 | table(gnMap$mouseID)[gnMap$mouseID]>1
  #Keep only the ones that we can map unambiguously and are in the Seurat annotation
  m = match(gns$mouseID,gnMap$mouseID)
  gns$humanID = ifelse(!is.na(m) & !gnMap$ambiguous[m], gnMap$humanID[m],NA)
  tmp = read.table('Data/scData/fAd/babyAdrenals/babyAdrenal1/filtered_gene_bc_matrices/GRCh38/genes.tsv',sep='\t',header=FALSE)
  gns$humanSym = tmp$V2[match(gns$humanID,tmp$V1)]
  #Keep and make seurat object
  srat = dat[!is.na(gns$humanSym),]
  #Label by symbol
  rownames(srat) = make.unique(gns$humanSym[!is.na(gns$humanSym)])
  srat = CreateSeuratObject(srat)
  m = match(colnames(srat),mDat$id)
  srat@meta.data = cbind(srat@meta.data,mDat[m,])
  srat = NormalizeData(srat)
  srat = FindVariableFeatures(srat)
  srat = ScaleData(srat)
  srat = RunPCA(srat,npcs=75)
  srat = RunUMAP(srat,dims=seq(75))
  srat = FindNeighbors(srat,dims=seq(75))
  srat = FindClusters(srat,resolution=1)
  #####
  # QC
  avg = AverageExpression(srat)$RNA
  #Drop PTPRC
  w = which(!srat@active.ident %in% colnames(avg)[avg['PTPRC',]>1])
  #########
  # Finish
  srat = srat@assays$RNA@counts[,w]
  m = match(colnames(srat),mDat$id)
  srat = CreateSeuratObject(srat)
  srat@meta.data = cbind(srat@meta.data,mDat[m,])
  srat = NormalizeData(srat)
  srat = FindVariableFeatures(srat)
  srat = ScaleData(srat)
  srat = RunPCA(srat,npcs=50)
  srat = RunUMAP(srat,dims=seq(50))
  srat = FindNeighbors(srat,dims=seq(50))
  srat = FindClusters(srat,resolution=1)
  srat@meta.data$annot = srat@meta.data$ClusterNames_final
  srat@meta.data$annot[grep('stromal',srat@meta.data$annot)]='Stroma'
  srat@meta.data$annot[grep('endoth',srat@meta.data$annot)]='Endothelium'
  srat@meta.data$annot = gsub('^[ED]\\.','',srat@meta.data$annot)
  srat@meta.data$annot = gsub('_','',srat@meta.data$annot)
  srat = SetIdent(srat,value='annot')
  srat@misc$geneMap = gns
  srat@misc$geneOrthologs = gnMap
  saveRDS(srat,tgtFile)
}
srats[[ex]]=srat


##########################
# Save in desired format #
##########################
for(ex in names(srats)){
  message(sprintf("Saving raw tables for %s",ex))
  srat = srats[[ex]]
  tgtFile = file.path(outDir,ex)
  #The matrix
  mtx = srat@assays$RNA@counts
  #Row names
  #Convert to ENSEMBL and retain symbol
  if(ex=='HCL'){
    gns = read.table(file.path(srcs$fAd[[1]][1],'filtered_gene_bc_matrices','GRCh38','genes.tsv'),sep='\t',header=FALSE)
    m = match(rownames(mtx),gns$V2)
  }else{
    if(ex %in% c('mouseDev','mouseDev_extra')){
      w = !is.na(srat@misc$geneMap$humanSym)
      gns = data.frame(V1=srat@misc$geneMap$humanID[w],
                       V2=make.unique(srat@misc$geneMap$humanSym[w])
                       )
    }else if(ex %in% c('mKidImm','mBM','sciKidRef','PBMC')){
      gns = read.table(file.path(srcs$fAd[[1]][1],'filtered_gene_bc_matrices','GRCh38','genes.tsv'),sep='\t',header=FALSE)
    }else if(ex=='PBMC'){
      gns = read.table(file.path(srcs[[ex]][[1]][1],'filtered_feature_bc_matrix','features.tsv.gz'),sep='\t',header=FALSE)
    }else if(ex=='PBMC3k'){
      gns = read.table(file.path(srcs$PBMC3k[[1]][1],'filtered_gene_bc_matrices','hg19','genes.tsv'),sep='\t',header=FALSE)
    }else{
      gns = read.table(file.path(srcs[[ex]][[1]][1],'filtered_gene_bc_matrices','GRCh38','genes.tsv'),sep='\t',header=FALSE)
    }
    m = match(rownames(mtx),gns$V2)
    #Need to fix some stupid hyphen underscore conversion problem.
    w=which(is.na(m))
    m[w] = m[w-1]+1
    w=which(is.na(m))
    m[w] = m[w-1]+1
  }
  if(ex %in% c('PBMC','sciKidRef')){
    dd = data.frame(idx=seq(nrow(mtx)),
                    geneID = rownames(mtx),
                    geneSymbol = gns$V2[match(rownames(mtx),gns$V1)]
                    )
    m = seq(nrow(mtx))
  }else{
    dd = data.frame(idx=seq(nrow(mtx)),
                    geneID = gns$V1[m],
                    geneSymbol = gns$V2[m]
                    )
  }
  w = which(!is.na(m))
  write.table(dd[w,],paste0(tgtFile,'Rownames.tsv'),sep='\t',row.names=FALSE,col.names=TRUE,quote=FALSE)
  #Write the matrix
  writeMM(mtx[w,],paste0(tgtFile,'.mtx'))
  #Column names
  #Ensure the triple underscore
  if(ex=='HCL'){
    colnames(mtx) = gsub('\\.','___',colnames(mtx))
  }else{
    colnames(mtx) = gsub('^([^_]+)_+(.+)$','\\1___\\2',colnames(mtx))
  }
  dd = data.frame(idx=seq(ncol(mtx)),
                  cellID = paste0(ex,'_',srat@meta.data$annot,':',colnames(mtx))
                  )
  #De-stupidify the labels
  dd$cellID = gsub('\\+','pos',dd$cellID)
  dd$cellID = gsub('[ -]','_',dd$cellID)
  write.table(dd,paste0(tgtFile,'Colnames.tsv'),sep='\t',row.names=FALSE,col.names=TRUE,quote=FALSE)
}
 
```
