## Supplementary material for "Single cell derived mRNA signals across human kidney tumors": Source Code: processSingleCellData.R

```
setwd('~/trueHome/Projects/CellDeconvolution')
#Process single cell tumour data to be compared against

#############
# Libraries #
#############
library(DropletUtils)
library(Seurat)
library(SoupX)
library(ggplot2)
library(Matrix)
source('https://raw.githubusercontent.com/constantAmateur/MCVR/master/code.R')
import('Code/scQC.R',as='scQC')
import('Code/logisticRegression.R',as='lr')
import('Code/geneSets.R')

#############
# Functions #
#############


#' Standard processing
stdProcess = function(dat,...){
  ss = CreateSeuratObject(dat)
  ss = NormalizeData(ss,verbose=FALSE)
  #ss = CellCycleScoring(ss, s.features=geneSets[['S']][geneSets[['S']] %in% rownames(dat)],g2m.features=geneSets[['G2M']][geneSets[['G2M']] %in% rownames(dat)],seed=sample(1e9,1))
  #ss = ScaleData(ss,vars.to.regress=c('S.Score','G2M.Score'),verbose=FALSE,...)
  ss = ScaleData(ss,verbose=FALSE,...)
  return(ss@assays$RNA@scale.data)
}

##############
# Parameters #
##############
#Output directory
outDir = 'Results/preProcess/scData/Tumour/'
srcs = list()
#Wilms data
srcs[['Wilms']] = list(oldWilms = file.path('Data/scTumours/Wilms',sprintf('Wilms%d',1:3)),
                       newWilms = 'Data/scTumours/Wilms/IMP672')
srcs[['Wilms']] = lapply(srcs[['Wilms']],function(e) {tmp = list.files(e,full.names=TRUE);names(tmp) = paste0('Wilms___',basename(dirname(tmp)),'___',basename(tmp));tmp})
names(srcs$Wilms$newWilms) = gsub('(.*)___.*_CG_SB_(.*)_GRCh3.*','\\1___\\2',names(srcs$Wilms$newWilms))
#Just keep pre-treated wilms
srcs$Wilms = srcs$Wilms[names(srcs$Wilms)=='newWilms']
#CMN data
srcs$CMN=list()
tmp = list.files('Data/scTumours/CMN',full.names=TRUE)
for(i in seq_along(tmp)){
  srcs$CMN[[basename(tmp[i])]] = list.files(tmp[i],full.names=TRUE)
  if(length(srcs$CMN[[basename(tmp[i])]])==0){
    srcs$CMN[[basename(tmp[i])]]=NULL
    next
  }
  names(srcs$CMN[[basename(tmp[i])]]) = paste0('CMN___',basename(tmp[i]),'___',basename(srcs$CMN[[basename(tmp[i])]]))
}
#Drop lovely second CMN :(
#srcs$CMN = srcs$CMN[names(srcs$CMN)!='IMP628']
#Drop the normal channel
srcs$CMN$IMP628 = srcs$CMN$IMP628[grep('4602STDY7894259',srcs$CMN$IMP628,invert=TRUE)]
#And the frozen samples as they have no viable data
srcs$CMN = srcs$CMN[names(srcs$CMN)!='EGFR']
srcs$CMN = srcs$CMN[names(srcs$CMN)!='IMP67']
#MRT data
#Organoids
srcs$MRT_org = list.files('Data/scTumours/MRT/Organoids/60',full.names=TRUE)
srcs$MRT_org = grep('1143$',srcs$MRT_org,invert=TRUE,value=TRUE)
names(srcs$MRT_org) = paste0('MRT_org___60___',basename(srcs$MRT_org))
srcs$MRT_org = list('60' = srcs$MRT_org)
#Fresh
srcs$MRT_fresh = list.files('Data/scTumours/MRT/GOSH029/',full.names=TRUE)
names(srcs$MRT_fresh) = paste0('MRT_fresh___GOSH029___',gsub('.*CG_SB_(.*)_GRCh.*','\\1',basename(srcs$MRT_fresh)))
srcs$MRT_fresh = list(GOSH029 = srcs$MRT_fresh)
#Nuclear
srcs$MRT_nuc = list.files('Data/scTumours/MRT/GOSH024/',full.names=TRUE)
names(srcs$MRT_nuc) = paste0('MRT_nuc___GOSH024___',gsub('.*CG_SB_(.*)_GRCh.*','\\1',basename(srcs$MRT_nuc)))
srcs$MRT_nuc = list(GOSH024 = srcs$MRT_nuc)
#srcs$MRT=list()
#tmp = list.files('Data/scTumours/MRT',full.names=TRUE)
#for(i in seq_along(tmp)){
#  srcs$MRT[[basename(tmp[i])]] = list.files(tmp[i],full.names=TRUE)
#  names(srcs$MRT[[basename(tmp[i])]]) = paste0('MRT___',basename(tmp[i]),'___',
#                                               gsub('.*CG_SB_(.*)_GRCh.*','\\1',basename(srcs$MRT[[basename(tmp[i])]])))
#}
#ATRT data
srcs$ATRT=list()
tmp = list.files('Data/scTumours/ATRT',full.names=TRUE)
for(i in seq_along(tmp)){
  srcs$ATRT[[basename(tmp[i])]] = list.files(tmp[i],full.names=TRUE)
  names(srcs$ATRT[[basename(tmp[i])]]) = paste0('ATRT___',basename(tmp[i]),'___',
                                               gsub('.*CG_SB_(.*)_GRCh.*','\\1',basename(srcs$ATRT[[basename(tmp[i])]])))
}
#CCSK data
srcs$CCSK=list()
tmp = list.files('Data/scTumours/CCSK',full.names=TRUE)
for(i in seq_along(tmp)){
  srcs$CCSK[[basename(tmp[i])]] = list.files(tmp[i],full.names=TRUE)
  names(srcs$CCSK[[basename(tmp[i])]]) = paste0('CCSK___',basename(tmp[i]),'___',
                                               gsub('.*CG_SB_(.*)_GRCh.*','\\1',basename(srcs$CCSK[[basename(tmp[i])]])))
}
#RCC
srcs$RCC=list()
tmp = list.files('Data/scTumours/ccRCC',full.names=TRUE)
tmp = c(tmp,list.files('Data/scTumours/pcRCC',full.names=TRUE))
for(i in seq_along(tmp)){
  srcs$RCC[[basename(tmp[i])]] = list.files(tmp[i],full.names=TRUE)
  names(srcs$RCC[[basename(tmp[i])]]) = paste0('RCC___',basename(tmp[i]),'___',
                                               gsub('.*CG_SB_(.*)_GRCh.*','\\1',basename(srcs$RCC[[basename(tmp[i])]])))
}
#Drop RCC3, it's a complete failure
srcs$RCC = srcs$RCC[names(srcs$RCC)!='RCC3']
#And Umbrella, due to weird sequencing fuck-up
#srcs$RCC = srcs$RCC[names(srcs$RCC)!='UMB82691004']
#Add outs if needed
for(a in names(srcs)){
  for(b in names(srcs[[a]])){
    #Drop BAMs if they're there
    srcs[[a]][[b]] = srcs[[a]][[b]][grep('\\.bam(\\.bai)?$',names(srcs[[a]][[b]]),invert=TRUE)]
    for(c in names(srcs[[a]][[b]])){
      tmp = file.path(srcs[[a]][[b]][c],'strainedCounts')
      w=!file.exists(tmp)
      tmp[w] = file.path(srcs[[a]][[b]][c],'cleanCounts')
      srcs[[a]][[b]][c] = tmp
    }
  }
}
#MT cut-off
mtCut = 0.4
#nGene Cut
minNumGenes = 300
#nUMI cut
minNumUMI = 1000
#How many doublets in a cluster before you throw the entire cluster
maxScrubFrac = 0.5
#UMAP parameters to try in grid
minDists = c(0,.01,.1,.3,.5)
NNs = c(5,10,20,50,100)
#Final parameters
finalMinDist = 0.5
finalNN = 50
#Clustering resolution
clusterRes=1.0
#Populations of genes to exclude from the analysis
excludeGenes = unlist(as.list(geneSets)[c('mtGenes','hspGenes','riboGenes')],use.name=FALSE)
excludeGenes = sort(unique(excludeGenes))
#Number of PCs to use for quick analysis during QC
defNumPCs=75
#Default contamination to use if estimation fails
defRho = 0.1
#Minimum contamination fraction to accept
minRho = 0.01

############
# Basic QC #
############
srats = list()
for(ex in names(srcs)){
  message(ex)
  tgtFile = file.path(outDir,sprintf('baseSeurat_%s.RDS',ex))
  if(file.exists(tgtFile)){
    srat = readRDS(tgtFile)
  }else{
    #Tweaked cut-offs for different data sets
    mtLoc = mtCut
    scrubLoc = 0.4
    umiLoc = minNumUMI
    #Create the basic Seurat object for each experiment
    message(sprintf("QC filters on %s",ex))
    tgts = unlist(srcs[[ex]])
    names(tgts) = gsub('.*\\.','',names(tgts))
    pdf(file.path(outDir,sprintf('%s_QC.pdf',ex)))
      srat = scQC$basicQC(dataDirs=unlist(srcs[[ex]]),
                          excludeGenes = excludeGenes,
                          geneSets = geneSets,
                          maxMT = mtLoc,
                          minGenes = minNumGenes,
                          minUMIs = umiLoc,
                          maxBadFrac = maxScrubFrac,
                          scrubScoreMax = scrubLoc,
                          numPCs = defNumPCs)
    dev.off()
    #Load the non-SoupX version, provided it exists
    tgts = setNames(file.path(tgts,'..','cleanCounts'),names(tgts))
    if(all(file.exists(tgts))){
      pdf(file.path(outDir,sprintf('%s_QC_noSoupX.pdf',ex)))
        srat@misc$noSoupX = scQC$basicQC(dataDirs=tgts,
                                         excludeGenes = excludeGenes,
                                         geneSets = geneSets,
                                         maxMT = mtLoc,
                                         minGenes = minNumGenes,
                                         minUMIs = umiLoc,
                                         maxBadFrac = maxScrubFrac,
                                         scrubScoreMax = scrubLoc,
                                         numPCs = defNumPCs
        )
      dev.off()
      srat@misc$noSoupX@meta.data$orig.ident = gsub('.*___(.*)___.*','\\1',rownames(srat@misc$noSoupX@meta.data))
    }
    #Fix orig.ident
    srat@misc$preQC@meta.data$orig.ident = gsub('.*___(.*)___.*','\\1',rownames(srat@misc$preQC@meta.data))
    srat@meta.data$orig.ident = gsub('.*___(.*)___.*','\\1',rownames(srat@meta.data))
    #Add sample 
    srat@meta.data$samp = gsub('.*___(.*)_[ACGT]+(-[0-9]+)?$','\\1',rownames(srat@meta.data))
    ####Add source metadata
    ###m = match(rownames(srat@meta.data),paste0(passOut$chan,'_',rownames(passOut)))
    ###srat@meta.data$cellPass = ifelse(!is.na(passOut$FDR[m]) & passOut$FDR[m]<0.05,
    ###                                 ifelse(passOut$passCR[m],
    ###                                        'Both',
    ###                                        'ED'
    ###                                        ),
    ###                                 'CR'
    ###                                 )
    saveRDS(srat,tgtFile)
  }
  srats[[ex]]=srat
}


################
# Optimal nPCs #
################
#Work out what the best number of PCs is
mcvs = list()
nPCs = c()
for(ex in names(srcs)){
  message(sprintf('Finding optimal number of PCs for experiment %s',ex))
  tgtFile=file.path(outDir,sprintf('MCV_%s.RDS',ex))
  srat = srats[[ex]]
  if(file.exists(tgtFile)){
    mcv = readRDS(tgtFile)
  }else{
    mcv = molecularCrossValidation(roundToInt(srat@assays$RNA@counts),VariableFeatures(srat),normalisation=stdProcess,features=VariableFeatures(srat),tFracs=c(1,0.9,0.8,0.5),nSplits=4,BPPARAM=SerialParam(),maxPCs=200)
    pdf(file.path(outDir,paste0(ex,'_mcv.pdf')))
      plotMCV(mcv)
    dev.off()
    saveRDS(mcv,tgtFile)
  }
  mcvs[[ex]] = mcv
  nPCs[ex] = max(mcv$mins1sd[[1]])
  #nPCs = sapply(mcvList,function(e) max(e$mins1sd[[1]]))
}


####################
# Final processing #
####################
for(ex in names(srcs)){
  tgtFile = file.path(outDir,sprintf('processedSeurat_%s.RDS',ex))
  message(sprintf("Finalising fit for %s",ex))
  if(file.exists(tgtFile)){
    srat = readRDS(tgtFile)
  }else{
    srat = srats[[ex]]
    pdf(file.path(outDir,sprintf('standardPlots_%s.pdf',ex)),width=24,height=24)
      srat = ScaleData(srat)
      srat = RunPCA(srat,npcs=nPCs[ex],approx=FALSE)
      #Run tSNE as well in case that turns out to be useful.
      #srat = RunTSNE(srat)
      #The old python based method seems to segfault every time...
      srat = FindNeighbors(srat,dims=seq(nPCs[ex]))
      srat = FindClusters(srat,res=clusterRes)
      #Make a range of UMAP plots
      #umaps=list()
      #for(minDist in minDists){
      #  for(NN in NNs){
      #    message(sprintf("Calculating UMAP for minDist=%g, NN=%d",minDist,NN))
      #    srat = RunUMAP(srat,dims=seq(nPCs[ex]),min.dist=minDist,n.neighbors=NN,verbose=FALSE)
      #    umaps[[paste0(minDist,'_',NN)]] = DimPlot(srat,label=TRUE) +
      #      guides(colour=FALSE) +
      #      ggtitle(sprintf("UMAP with minDist=%g, NN=%d",minDist,NN))
      #  }
      #}
      #plot(plot_grid(plotlist=umaps,
      #          nrow=length(minDists),
      #          ncol=length(NNs)))
      #Final UMAP
      srat = RunUMAP(srat,dims=seq(nPCs[ex]),min.dist=finalMinDist,n.neighbors=finalNN)
      #Clustering
      plot(DimPlot(srat,label=TRUE))
      #My ones
      plot(FeaturePlot(srat,c('PTPRC','CD74','PDGFRB','EPCAM','PECAM1','HBB','ITGA2B')))
      #Source
      plot(DimPlot(srat,group.by='orig.ident'))
      #Cell cycle
      plot(DimPlot(srat,group.by='Phase'))
      #QC things
      plot(FeaturePlot(srat,c('scrubScore','nFeature_RNA','nCount_RNA',names(geneSets))))
      #Visualise cell cycle
      #plot(FeaturePlot(srat,c('PCNA','MKI67','MCM4','CDKN1C')))
      #Genes specific to Steroidogenisis
      #plot(FeaturePlot(srat,c('MC2R','CYP17A1','MRAP','STAR','CYP11B2','CYP11B1','INHA','GNRHR','HSD3B2')))
      #Medulla markers
      #plot(FeaturePlot(srat,c('PHOX2A','ISL1','CHRNA3','SOX2','SOX10','ERBB3','DBH','CHGB','TH')))
      #Others
      #plot(FeaturePlot(srat,c('NR5A1','NOV','HSD3B2','SULT2A1','FDX1')))
    dev.off()
    saveRDS(srat,tgtFile)
  }
  srats[[ex]] = srat
}

########################
# Non standard samples #
########################

##############
# Jarno CCSK #
##############
ex='CCSK_extra'
tgtFile = file.path(outDir,sprintf('processedSeurat_%s.RDS',ex))
if(file.exists(tgtFile)){
  message(sprintf("Finalising fit for %s",ex))
  ccsk = readRDS(tgtFile)
}else{
  #######
  # Load
  mDat = read.table('Data/Mouse_CCSK/metaData_CCSK.txt',sep='\t',header=TRUE)
  x = read.table('Data/Mouse_CCSK/TM211.transcripts.txt',sep='\t',header=TRUE)
  xx = x[,-c(1,2)]
  colnames(xx) = paste0('CSK_1015T1_pl1_TM211__',colnames(xx))
  y = read.table('Data/Mouse_CCSK/TM212.transcripts.txt',sep='\t',header=TRUE)
  yy = y[,-c(1,2)]
  colnames(yy) = paste0('CCSK_1015T1_pl2_TM212__',colnames(yy))
  gns = intersect(x$GENEID,y$GENEID)
  gns = grep('-cluster$',gns,value=TRUE,invert=TRUE)
  #Drop the duplicated symbols
  tmp = table(gsub('.*__','',gns))
  gns = gns[!gsub('.*__','',gns) %in% names(tmp[tmp>1])]
  dat = cbind(xx[match(gns,x$GENEID),],yy[match(gns,y$GENEID),])
  rownames(dat) = gns
  #Get meta-data stuff
  mm = lapply(geneSets,function(e) colSums(dat[match(e,gsub('.*__','',rownames(dat))),,drop=FALSE],na.rm=TRUE)/colSums(dat))
  mm = as.data.frame(do.call(cbind,mm))
  mm$ERCC =  colSums(dat[grep('^ERCC-',gns),])/colSums(dat)
  #Now drop the useless ones
  dat = dat[-c(grep('ERCC-',rownames(dat)),grep('MT-',rownames(dat))),]
  rownames(dat) = gsub('.*__','',rownames(dat))
  #####
  # QC
  ccsk = CreateSeuratObject(dat)
  ccsk@meta.data = cbind(ccsk@meta.data,mm)
  ccsk = NormalizeData(ccsk)
  ccsk = FindVariableFeatures(ccsk)
  ccsk = ScaleData(ccsk)
  ccsk = RunPCA(ccsk,npcs=75)
  ccsk = RunUMAP(ccsk,dims=seq(75))
  ccsk = FindNeighbors(ccsk,dims=seq(75))
  ccsk = FindClusters(ccsk,resolution=10)
  #Do the standard QC stuff.
  ccsk@meta.data$PASS_MT = !(ccsk@meta.data$mtGenes >= .2)
  ccsk@meta.data$PASS_ERCC = !(ccsk@meta.data$ERCC >= .2)
  ccsk@meta.data$PASS_nGenes = !(ccsk@meta.data$nFeature_RNA <= 300)
  ccsk@meta.data$PASS_nCounts = !(ccsk@meta.data$nCount_RNA <= 1000)
  tt = aggregate(!PASS_MT | !PASS_nGenes | !PASS_nCounts | !PASS_ERCC ~ seurat_clusters,FUN=mean,data=ccsk@meta.data)
  ccsk@meta.data$fracBadClust = tt[match(ccsk@meta.data$seurat_clusters,tt[,1]),2]
  ccsk@meta.data$PASS_cluster = !(ccsk@meta.data$fracBadClust > 0.5)
  ccsk@meta.data$PASS = with(ccsk@meta.data,PASS_MT & PASS_nGenes & PASS_nCounts & PASS_cluster & PASS_ERCC)
  #Now do the drop and proper process
  w = which(ccsk@meta.data$PASS)
  old = ccsk
  mDat = ccsk@meta.data[w,]
  #Drop old clustering
  mDat = mDat[,!colnames(mDat) %in% c('seurat_clusters',grep('^RNA_snn_res',colnames(mDat),value=TRUE))]
  #And drop genes that we don't care about
  genesToKeep = rownames(ccsk@assays$RNA@counts)
  genesToKeep = genesToKeep[!genesToKeep %in% excludeGenes]
  ccsk = CreateSeuratObject(ccsk@assays$RNA@counts[genesToKeep,w])
  #Merge in metadata
  ccsk@meta.data = cbind(ccsk@meta.data,mDat[,!(colnames(mDat) %in% colnames(ccsk@meta.data)),drop=FALSE])
  ccsk = NormalizeData(ccsk,verbose=FALSE)
  ccsk = FindVariableFeatures(ccsk,verbose=FALSE)
  ccsk@misc$preQC =old 
  #########
  # Finish
  mcv = molecularCrossValidation(roundToInt(ccsk@assays$RNA@counts),VariableFeatures(ccsk),normalisation=stdProcess,features=VariableFeatures(ccsk),tFracs=c(1,0.9,0.8,0.5),nSplits=4,BPPARAM=MulticoreParam(8),maxPCs=100)
  pdf(file.path(outDir,paste0(ex,'_mcv.pdf')))
    plotMCV(mcv)
  dev.off()
  nn = max(mcv$mins1sd[[1]])
  ccsk = NormalizeData(ccsk)
  ccsk = FindVariableFeatures(ccsk)
  ccsk = ScaleData(ccsk)
  ccsk = RunPCA(ccsk,npcs=nn)
  ccsk = RunUMAP(ccsk,dims=seq(nn))
  ccsk = FindNeighbors(ccsk,dims=seq(nn))
  ccsk = FindClusters(ccsk,resolution=1)
  saveRDS(ccsk,tgtFile)
}
srats[[ex]] = ccsk
  


 
 
```
