## Supplementary material for "Single cell derived mRNA signals across human kidney tumors": Source Code: revisionFigures.R

```
#' Generates main results for paper
#'
#' Creates for bulk and sc data against various references using LR and Signal analysis.

#################
# Materials map #
#################


#Supplementary Figures
#S1a-c - Justification for fetal kidney labels.
#S1d - MPCs in kidney strongly resemble the mesenchymal population in the adrenal gland.
#S1e - MPCs resemble primitive mesoderm.
#S2 - PT1 signal identified as major contributor to ccRCC/pRCC.
#S3 - Heatmap for CMN against fKid.
#S4 - Annotation for CMN.
#S5 - Heatmap for Wilms and CCSK against fKid.
#S6 - Annotation and proliferation status of Wilms.
#S7 - Annotation of CCSK.
#S8 - Heatmap for MRT against fKid.
#S9 - Annotation for MRT.
#S10 - Expression of TWIST1 in single cell MRT transcriptomes.
#S11 - Heatmap for RCCs against mKid.
#S12 - Type A/B gene expression in RCCs.
#S13 - Heatmap for chRCCs against mKid.
#S14 - Specificity of signal to different tumour types.
#S15 - Copy number profiles of Wilms.
#S16 - Colour map normal
#S17 - Colour map tumour


#Tables
#1 - List of all bulk transcriptomes
#2 - List of all single cell experiments.
#3 - Summary table for linear model associating immaturity score to genotype.
#4 - Summary table for linear model associating immaturity score to other groupings.
#5 - Mast and cell numbers from smFISH.

#Supplementary Data
#1 - All bulk RNA seq raw files
#2 - All single cell raw files
#3 - All data tables to generate figures
#4 - Supplementary code

###################
# Set Project Dir #
###################
setwd('~/trueHome/Projects/CellDeconvolution')

#############
# Libraries #
#############
#General
library(Matrix)
library(Matrix.utils)
library(Seurat)
library(SummarizedExperiment)
library(survival)
#Plotting libraries
library(ggplot2)
library(plotrix)
library(reshape2)
library(beeswarm)
library(RColorBrewer)
library(survminer)
import('Code/plotParts.R',as='pl')
#Alt fitting libraries
library(bseqsc)
library(MuSiC)
library(xbioc)
import('Code/logisticRegression.R',as='lr')
import('Code/scQC.R',as='scQC')
import('Code/geneSets.R')

####################
# Useful functions #
####################


#' Alternative visualisation of signals
#'
#' Plots the distribution as a boxplot style thing, but joins each point with faint line to get the connection between points.
#'
#' @param dat A matrix containing exposures, where columns are signals and rows are cells.
#' @param pR2 The goodness of fit metric.  One for each cell.
#' @param alpha Transparency value for ponits.
#' @param joinPoints Join ponits with a line.
altVis = function(dat,pR2,alpha=0.4,joinPoints=FALSE){
  dat = as.matrix(dat)
  if(is.null(rownames(dat)))
    rownames(dat) = paste0('Cell',seq(nrow(dat)))
  if(is.null(colnames(dat)))
    colnames(dat) = paste0('Sig',seq(ncol(dat)))
  names(pR2) = rownames(dat)
  #Define colour scheme for goodness of fit
  pR2cols = color.scale(pR2,extremes=c('red','black'),alpha=alpha)
  names(pR2cols) = names(pR2)
  #Work out exactly where everything will go
  dd = data.frame(value = as.vector(dat),
                  cellID = rep(rownames(dat),ncol(dat)),
                  sigID = rep(colnames(dat),each=nrow(dat))
                  )
  dd$xval = match(dd$sigID,colnames(dat))-1+rnorm(nrow(dd),0,0.1)
  #Plot area
  plot(0,
       type='n',
       xlim=c(-0.5,ncol(dat)-0.5),
       ylim=c(0,1),
       frame.plot=FALSE,
       xaxt='n',
       xlab='',
       ylab='Score'
       )
  axis(1,
       at=seq(ncol(dat))-1,
       lwd=0,
       lwd.ticks=1,
       las=2,
       label=colnames(dat))
  #Joining lines, one at a time
  if(joinPoints){
    for(cellID in rownames(dat)){
      lines(dd$xval[dd$cellID==cellID],dd$value[dd$cellID==cellID],
            col=pR2cols[cellID],
            lwd=0.1)
    }
  }
  points(dd$xval,dd$value,
         pch=19,
         col=pR2cols[dd$cellID],
         cex=0.5)
  #Make median plots
  for(sigID in colnames(dat)){
    i = match(sigID,colnames(dat))
    lines(x=i-1+c(-0.5,0.5)*2/3,
          y=rep(median(dd$value[dd$sigID==sigID]),2),
          lwd=3,
          col='grey'
    )
  }
  #Add the visualisation
  color.legend(ncol(dat)-0.8,0.4,ncol(dat)-0.5,1,
               legend=c(min(pR2),rep(NULL,length(pR2cols)-2),max(pR2)),
               rect.col=sort(pR2cols,decreasing=TRUE),
               align='rb',
               gradient='y'
               )
}

#' Colour to hue
#' 
#' Hue from colour
#' @param col Colour to Hue.
col2hsv = function(colour){
  tmp = col2rgb(colour)[,1]/255
  cmax = max(tmp)
  cmin = min(tmp)
  delta = cmax-cmin
  w = which.max(tmp)
  if(delta==0){
    hue = 0
  }else{
    if(names(w)=='red'){
      hue = (tmp['green']-tmp['blue'])/delta
    }else if(names(w)=='green'){
      hue = 2 + (tmp['blue']-tmp['red'])/delta
    }else{
      hue = 4 + (tmp['red']-tmp['green'])/delta
    }
  }
  hue = hue*60
  if(cmax==0){
    sat = 0
  }else{
    sat = delta/cmax
  }
  val = cmax
  return(as.numeric(c(hue/360,sat,val)))
}


#' Load a matrix
#'
#' @param src Where to load it
loadRefMtx = function(src){
  mtx = readMM(paste0(src,'.mtx'))
  rownames(mtx) = read.table(paste0(src,'Rownames.tsv'),sep='\t',header=TRUE)$geneID
  colnames(mtx) = read.table(paste0(src,'Colnames.tsv'),sep='\t',header=TRUE)$cellID
  return(mtx)
}

#' Save reference sparse matrix in format needed for python
#'
#' @param mtx Matrix to save.
#' @param tgtFile Where to save it.
saveRefMtx = function(mtx,tgtFile){
  #Matrix
  writeMM(mtx,paste0(tgtFile,'.mtx'))
  #Rownames
  write.table(rownames(mtx),paste0(tgtFile,'_rowNames.tsv'),sep='\t',row.names=FALSE,col.names=FALSE,quote=FALSE)
  #Column names
  write.table(colnames(mtx),paste0(tgtFile,'_columnNames.tsv'),sep='\t',row.names=FALSE,col.names=FALSE,quote=FALSE)
}

#' Save a matrix of counts and lengths in input format
#'
#' If all entries have the same length information (i.e. lengths is a vector), this is saved to one file.  Otherwise, saved to one file per sample in a new folder and a linking file is created.  Single file mode can be forced.
#'
#' @param mtx The count matrix to save.  Rows are genes, columns samples.
#' @param lengths A vector or matrix giving the lengths associated associated with each gene in each sample.
#' @param tgtFile File to save to.  If single file mode, ending is stripped and a folder created.
#' @param forceSingle Force function to write out one file per sample.
#' @return Nothing.  Saves files.
saveFitMtx = function(mtx,lengths,tgtFile,forceSingle=FALSE){
  #Which mode
  if(!is.null(dim(lengths)) | forceSingle){
    tgtDir = gsub('\\.[^\\.]+$','',tgtFile)
    dir.create(tgtDir,showWarnings=FALSE,recursive=TRUE)
    for(i in seq_len(ncol(mtx))){
      if(is.null(dim(lengths))){
        len = lengths
      }else{
        len = lengths[,i]
      }
      dd = cbind(geneName=rownames(mtx),geneLenths=lengths,XXX=mtx[,i])
      colnames(dd)[3] = colnames(mtx)[i]
      write.table(dd,file.path(tgtDir,paste0(colnames(mtx)[i],'.tsv')),sep='\t',row.names=FALSE,col.names=TRUE,quote=FALSE)
    }
    #Finally, write index file
    write.table(file.path('.',basename(tgtDir),paste0(colnames(mtx),'.tsv')),tgtFile,row.names=FALSE,col.names=FALSE,quote=FALSE)
  }else{
    dd = cbind(geneName=rownames(mtx),geneLengths=lengths,as.data.frame(as.matrix(mtx)))
    write.table(dd,tgtFile,sep='\t',row.names=FALSE,col.names=TRUE,quote=FALSE)
  }
}

#' Runs code for a set of samples and signatures and loads the results
#'
#' @param samples Samples to run.  Passed to -b
#' @param references Reference to fit against.  Passed to -s 
#' @param tgt Name for output
#' @param weights Weights for genes.  Passed to -w
#' @param collapseClasses Collapse reference data into classes instead of using each cell as a reference.
#' @param skipFit Don't run, just load.
#' @param debug Make ipython interactive.
#' @param returnCommand Just return the command and quit.
#' @param tol Termination tollerance.
#' @return A list with fit exposures, normalised exposures and extra info.
runAndLoad = function(samples,references,tgt,weights='Data/geneWeights.tsv',collapseClasses=TRUE,skipFit=TRUE,debug=FALSE,returnCommand=FALSE,tol=0.01){
  #create weights if it's not a file that exists
  if(!is.character(weights)){
    dd = data.frame(geneID = names(weights),
                    weight = weights)
    weights = tempfile() 
    write.table(dd,weights,row.names=FALSE,col.names=TRUE,sep='\t',quote=FALSE)
  }
  #Fix paths as python abhors non-absolute paths
  samples = normalizePath(samples)
  references = normalizePath(references)
  weights = normalizePath(weights)
  #################
  # Construct call
  code = paste0('ipython ',
                ifelse(debug,'-i',''),
                ' -- Code/cellSignalAnalysis.py ',
                sprintf(' -ts %g ',tol), #No need to waste time once likelihood converged.
                paste0(' -b ',samples),
                paste0(' -s ',references),
                paste0(' -w ',weights),
                ifelse(collapseClasses,' ',' --no-collapse-ref '),
                tgt)
  if(returnCommand)
    return(code)
  ###########
  # Run code
  if(!skipFit)
    system(code)
  ###################
  # Load the results
  fNom = paste0(tgt,'_fitExposures.tsv')
  fit = read.table(fNom,sep='\t',header=TRUE)
  extras = fit[match(c('Intercept','fitCount','obsCount','pR2'),rownames(fit)),,drop=FALSE]
  fit = fit[-match(c('Intercept','fitCount','obsCount','pR2'),rownames(fit)),,drop=FALSE]
  #Normalise, including the intercept
  norm = data.frame(t(t(fit)/(unlist(extras['Intercept',,drop=FALSE])+colSums(fit))))
  #Add intercept back in
  norm = rbind(norm,Intercept = pmax(0,1-colSums(norm)))
  out = list(fit=fit,extras=extras,norm=norm)
  return(out)
}

#' Load bulk data
#'
#' Load bulk data from a list of samples stored in the required format.
#'
#' @param tgts Files to load.
#' @return A list containing counts and lengths.
loadBulk = function(tgts){
  #Load them
  dat = lapply(tgts,read.table,sep='\t',header=TRUE)
  gns = Reduce(intersect,lapply(dat,function(e) e$geneName))
  cnts = do.call(cbind,lapply(dat,function(e) e[match(gns,e$geneName),-c(1,2),drop=FALSE])) 
  rownames(cnts) = gns
  lens = do.call(cbind,lapply(dat,function(e) {
                                tmp = e[match(gns,e$geneName),rep(match('geneLengths',colnames(e)),ncol(e)-2),drop=FALSE] 
                                colnames(tmp) = colnames(e)[-c(1,2)]
                                tmp}))
  rownames(lens) = gns
  return(list(cnts=cnts,lens=lens))
}

#' Load full matrix
#'
#' Loads the full matrix for a reference
#'
#' @param ref The reference object.
loadFullMtx = function(ref){
  mtx = lapply(ref$mtx,function(e) loadRefMtx(e))
  gns = Reduce(intersect,lapply(mtx,rownames))
  mtx = do.call(cbind,lapply(mtx,function(e) e[gns,]))
  #Train logistic regression model
  classes = gsub(':.*','',colnames(mtx))
  #Clean class labels 
  classes = gsub('/.*','',classes)
  #Drop anything with too few observations to train with
  w = which(gsub('^[^_]+_','',classes)!='NA' & table(classes)[classes]>20)
  return(mtx[,w])
}

#' Save raster and non-raster versions of plots
#'
#' Saves a pdf and png version of everything.
#'
#' @param baseName Name of file to save.  File name without extension.
#' @param plotFun Function to generate the plot.  Must take two arguments, noFrame and noPlot which control if the plot part / frame part are drawn.
#' @param width Width in inches.
#' @param heights Height in inches.
#' @param res Resolution in pixels per inch for rasterised image.
#' @param rawData If provided, save the raw data used to make the plot.
#' @param row.names Passed to write.table
#' @param col.names Passed to write.table
#' @param ... Passed to plotFun
saveFig = function(baseName,plotFun,width=4,height=3,res=300,rawData=NULL,row.names=FALSE,col.names=TRUE,...){
  #Save the base versions
  pdf(paste0(baseName,'.pdf'),width=width,height=height,useDingbats=FALSE)
  plotFun(noFrame=FALSE,noPlot=FALSE,...)
  dev.off()
  png(paste0(baseName,'.png'),width=width,height=height,res=300,units='in')
  plotFun(...)
  dev.off()
  #And the various bits of frame in vector format
  pdf(paste0(baseName,'_frame.pdf'),width=width,height=height,useDingbats=FALSE)
  plotFun(noPlot=TRUE,noFrame=FALSE,...)
  dev.off()
  #And the various bits of the actual plot in raster format
  png(paste0(baseName,'_plot.png'),width=width,height=height,res=300,units='in')
  plotFun(noPlot=FALSE,noFrame=TRUE,...)
  dev.off()
  #Save raw data if provided
  if(!is.null(rawData)){
    write.table(rawData,paste0(baseName,'_rawData.tsv'),
                quote=FALSE,
                sep='\t',
                row.names=row.names,
                col.names=col.names)
  }
}

#' Boxplots with points overlaid
#'
#' Groups data either first by tumour type, then signatures or the other way around.  Only boxplot is show for control samples and with decreased width.  Colours are used to indicate the second split.  Points are either shown using jitter or by binning and creating a dot plot.
#'
#' @param dat Matrix with rows representing signatures, columns samples named in the format <TumourType>___<SampleID> and entries the exposures.
#' @param isControl A named vector indicating which of the tumour types are being used as controls.
#' @param groupBySigs Main grouping is by tumour type by default, if set to TRUE, do main grouping by Signatures.
#' @param dotPlot Default behaviour is to jitter points.  This instead produces a dot plot.
#' @param greyScale Don't use colours, use greyscale.
#' @param controlWidth The width, relative to 1 for a normal group, devoted to each control.
#' @param colPal Colour palette to use.  Can also be named list of colours.
#' @param jitterSDs How many standard deviations to use to jitter x.  Higher number means less jitter
#' @param splitLims Region within [0,1] to use for actually plotting in the first split.
#' @param pointLims Region within [0,1] to use for plotting data within the second split.
#' @param nBins Number of vertical bins for dot plot.
#' @param xPtWid How much space per point in x direction.  If NULL set so that the biggest bin in the plot is filled.
#' @param yPtWid How much space per point in y direction.
#' @param fixNumX Have this many points per row, overrides xPtWid.
#' @param fixNumY Have this many points per col, overrides yPtWid.
#' @param scaleToBox Scale number of points so the fullest bin in each box fills that row.
#' @param packDense Err on the side of more points?
#' @param maxY y-limit for plot
#' @param ptAlpha Alpha value for points.  If a value greater than one, alpha set dynamically such that alpha*numPoints = ptAlpha
#' @param pch Shape of points.
#' @param cex Size of points.
#' @param labCex Size of labels.
#' @param boxSqueeze Scale boxplot width relative to dot plotting by this factor.
#' @param yLabel Y-axis label.
#' @param controlColChange Shift all the colour to the boxes.  Tweaks everything to make this work.
#' @param abb Abbreviation for split labels.  If NULL calculated automatically.
exposurePlot = function(dat,isControl=NULL,groupBySigs=FALSE,dotPlot=TRUE,greyScale=FALSE,controlWidth=0.2,colPal = brewer.pal(9,'Set1'),jitterSDs=5,splitLims=c(0.1,0.9),pointLims=c(0.05,0.95),nBins=100,xPtWid=0.01,yPtWid=0.01,fixNumX=1,fixNumY=1,scaleToBox=TRUE,packDense=TRUE,maxY=ceiling(max(dat)),ptAlpha=0.2,pch=19,cex=0.8,labCex=3.0,boxSqueeze=0.8,yLabel='Exposure',controlColChange=FALSE,abb=NULL){
  ##############
  # Preprocess #
  ##############
  #Stick the controls on the right
  dat = dat[,order(isControl[gsub('___.*','',colnames(dat))]),drop=FALSE]
  splitWidth = splitLims[2]-splitLims[1]
  df = melt(as.matrix(dat),as.is=TRUE,varnames=c('Signatures','Sample'))
  df$class = gsub('___.*','',df$Sample)
  if(is.null(isControl))
    isControl = setNames(rep(FALSE,length(unique(df$class))),unique(df$class))
  #Designate controls
  df$isControl = isControl[df$class]
  #Define order
  df$Signatures = factor(df$Signatures,levels=rownames(dat))
  df$class = factor(df$class,levels=unique(df$class[match(colnames(dat),df$Sample)]))
  #Check that controls are all together 
  if(length(which(isControl[levels(df$class)]))>1 & any(diff(which(isControl[levels(df$class)]))!=1))
    stop("Controls must form a continuous block.  Please re-order.")
  if(groupBySigs){
    tmp = colnames(df)
    tmp[match(c('Signatures','class'),tmp)]=c('class','Signatures')
    colnames(df) = tmp
    xBreaks = seq(nlevels(df$class)+1)-0.5
    defWidths = rep(splitWidth/nlevels(df$Signatures),nlevels(df$Signatures))
    defWidths = defWidths*ifelse(isControl[levels(df$Signatures)],controlWidth,1)
    defWidths = defWidths/sum(defWidths)*splitWidth
    x2Breaks = c(0.1,0.1+cumsum(defWidths))
  }else{
    #Boundaries of each class
    xBreaks = c(0.5,0.5+cumsum(ifelse(isControl[levels(df$class)],controlWidth,1)))
    #Boundaries of each signature, assuming class area has width 1
    x2Breaks = seq(splitLims[1],splitLims[2],length.out=nlevels(df$Signatures)+1)
  }
  #Boundaries of plot area within signature, assuming signature area has width 1
  x3Breaks = pointLims
  xWidths = diff(xBreaks)
  #y-breakpoints
  yBreaks = seq(floor(min(dat)),maxY,length.out=nBins+1)
  #############################
  # Calculate point locations #
  #############################
  xx = rep(NA,nrow(df))
  yy = rep(NA,nrow(df))
  #Coordinates for extra annotation
  boxVals = list()
  if(is.null(xPtWid)){
    #Determine the number of points in the largest bin that we will plot
    bigBinNo = df[!df$isControl,]
    bigBinNo = with(bigBinNo,split(value,interaction(Signatures,class)))
    bigBinNo = lapply(bigBinNo,cut,yBreaks,include.lowest=TRUE)
    bigBinNo = sapply(lapply(bigBinNo,table),max)
    bigBinNo = max(bigBinNo)
    #Set point spacing so the fullest bin would be just filled
    xPtWid = max(xWidths[!isControl[names(xWidths)]])*max(diff(x2Breaks))/bigBinNo
  }
  for(i in levels(df$class)){
    boxVals[[i]]=list()
    for(j in levels(df$Signatures)){
      #Get the data
      w = which(df$class==i & df$Signatures==j)
      #Define region
      xLeft = xBreaks[match(i,levels(df$class))] + x2Breaks[match(j,levels(df$Signatures))]*xWidths[match(i,levels(df$class))]
      xRight = xBreaks[match(i,levels(df$class))] + x2Breaks[match(j,levels(df$Signatures))+1]*xWidths[match(i,levels(df$class))]
      #Tweak by final level of breaks
      xMid = (xLeft+xRight)*0.5
      xLeft = xMid + (x3Breaks[1]-0.5)*(xRight-xLeft)
      xRight = xMid + (x3Breaks[2]-0.5)*(xRight-xLeft)
      #Get number of points in the x-direction
      if(is.null(fixNumX)){
        nX = (xRight-xLeft)/xPtWid
        if(packDense){
          nX = ceiling(nX)
        }else{
          nX = max(1,floor(nX))
        }
      }else{
        nX = fixNumX
      }
      #Get quantiles
      boxVals[[i]][[j]]=list(quant=quantile(df$value[w]),xLeft=xLeft,xRight=xRight,dat=df$value[w])
      #Give plotting order for each point in each bin
      if(dotPlot){
        #tmp = beeswarm(df$value[w],method='hex',breaks=seq(0,1,length.out=nBins+1),corral='wrap',do.plot=FALSE,cex=cex)
        #yy[w] = tmp$y
        ##Centre around 0
        #tmp = (tmp$x -1)
        ##Boundaries should be at -0.5/0.5, scale them so they're at xLeft/xRight
        #tmp = tmp*(xRight-xLeft)
        ##Shift centre
        #tmp = tmp+0.5*(xLeft+xRight)
        #xx[w] = tmp 
        #Split into bin
        cc = cut(df$value[w],yBreaks,include.lowest=TRUE)
        if(scaleToBox)
          nX = max(table(cc))
        for(k in levels(cc)){
          #Process each y-bin at a time
          ww = which(cc==k)
          o = seq_along(ww)-1
          #Get number of points in the y-direction
          ii = match(k,levels(cc))
          if(is.null(fixNumY)){
            nY = (yBreaks[ii+1]-yBreaks[ii])/yPtWid
            if(packDense){
              nY = ceiling(nY)
            }else{
              nY = max(1,floor(nY))
            }
          }else{
            nY = fixNumY
          }
          #Line stacker
          #nY = 1
          #nX = length(ww)
          #Convert plotting order to positions relative to middle of bin
          #Place in y-direction first
          a = o %% nY
          #Oscillate around middle
          a = ceiling(a/2)*ifelse(a%%2,1,-1)
          #Then in x-direction
          b = (o %/% nY) %% nX
          b = ceiling(b/2)*ifelse(b%%2,1,-1)
          #Define the x point positions for each
          xPos = seq(xLeft,xRight,length.out=nX+1)
          xPos = xPos[-length(xPos)]+diff(xPos)/2
          #And thy y point positions, which depends on the bin number
          yPos = seq(yBreaks[ii],yBreaks[ii+1],length.out=nY+1)
          yPos = yPos[-length(yPos)]+diff(yPos)/2
          #Finally construct the positions
          xPos = xPos[b+ceiling(nX/2)]
          yPos = yPos[a+ceiling(nY/2)]
          #Centre it?
          xPos = xPos - mean(xPos) + (xLeft+xRight)/2
          yPos = yPos - mean(yPos) + (yBreaks[ii]+yBreaks[ii+1])/2
          xx[w[ww]] = xPos
          yy[w[ww]] = yPos
        }
      }else{
        #Jitter instead
        xx[w] = rnorm(length(w),mean=0.5*(xLeft+xRight),sd = (xRight-xLeft)/jitterSDs) 
        yy[w] = df$value[w]
      }
    }
  }
  ###############
  # Do the plot #
  ###############
  #Define colours
  if(greyScale){
    cols = rep('grey',nlevels(df$Signatures))
  }else{
    if(is.function(colPal)){
      cols = colPal(nlevels(df$Signatures))
    }else{
      cols = colPal[levels(df$Signatures)]
    }
    #Repeat if needed
    if(any(is.na(cols))){
      nCols = max(which(!is.na(cols)))
      cols = cols[((seq_along(cols)-1) %% nCols)+1]
    }
  }
  names(cols) = levels(df$Signatures)
  par(mar=c(7.1, 4.1, 4.1, 9.1))
  #Define the empty plot area
  plot(0,0,type='n',xlab='',ylab=yLabel,ylim=c(floor(min(dat)),maxY),xlim=range(xx),xaxt='n',frame.plot=FALSE,las=1,cex.axis=labCex)
  #Dynamically determine alpha
  if(ptAlpha>1){
    dynAlpha = ptAlpha/table(df$Signatures,df$class)
    m = cbind(match(df$Signatures,rownames(dynAlpha)),match(df$class,colnames(dynAlpha)))
    alpha = pmin(1,dynAlpha[m])
  }else{
    alpha = ptAlpha
  }
  w = which(!controlColChange | !df$isControl)
  if(controlColChange){
    ptCols = pl$colAlpha('#606060',alpha)
  }else{
    ptCols = pl$colAlpha(cols[df$Signatures],alpha)[w] 
  }
  points(xx[w],yy[w],
       col=ptCols,
       pch=pch,
       cex=cex,
       xaxt='n',
       yaxt='n'
       )
  #Plot boxplots
  boxDat = unlist(lapply(boxVals,function(e) lapply(e,function(ee) ee$dat)),recursive=FALSE)
  boxPos = unlist(lapply(boxVals,function(e) lapply(e,function(ee) 0.5*(ee$xLeft+ee$xRight))))
  boxWidth = unlist(lapply(boxVals,function(e) lapply(e,function(ee) (ee$xRight-ee$xLeft))))
  #Boxplot
  if(!groupBySigs){
    sigNoms = gsub('^(.+)\\.(.+)$','\\1',names(boxDat))
    classNoms = gsub('^(.+)\\.(.+)$','\\2',names(boxDat))
  }else{
    #This is a bit weird and hacky, but is what is needed to make the colour selection work for boxplots
    sigNoms = gsub('^(.+)\\.(.+)$','\\2',names(boxDat))
    classNoms = gsub('^(.+)\\.(.+)$','\\2',names(boxDat))
  }
  boxplot(boxDat,
          at=boxPos,
          width=boxWidth*boxSqueeze,
          pars=list(boxwex=max(boxWidth)*boxSqueeze,#Make widths absolute, not relative
                    staplewex = 0, #Turn off things at top
                    whisklty=1), #Solid whiskers
          outline=FALSE,
          boxlwd = ifelse(controlColChange,3,1),
          whisklwd = ifelse(controlColChange,3,1),
          #border = ifelse(controlColChange & isControl[sigNoms],
          #                cols[classNoms],
          #                'black'
          #                ),
          border = cols[classNoms],
          xaxt='n',
          yaxt='n',
          col =pl$colAlpha('white',0.4),
          frame.plot=FALSE,
          add=TRUE
          )
  #Define an abbreviation for the inner split.  Increase string length until unique
  if(is.null(abb)){
    #Increase each name independently
    ll=rep(NA,nlevels(df$Signatures))
    for(i in seq_along(ll)){
      j=1
      while(j<=max(nchar(levels(df$Signatures)))){
        abb = substr(levels(df$Signatures),1,j)
        if(sum(abb==abb[i])==1)
          break
        j = j +1
      }
      ll[i]=j
    }
    abb = setNames(substr(levels(df$Signatures),1,ll),levels(df$Signatures))
  }
  #Create a series of axes labelling things
  for(i in seq_along(xBreaks[-1])){
    off = (x2Breaks[-length(x2Breaks)]+diff(x2Breaks)/2)*(xBreaks[i+1]-xBreaks[i])
    #Create the deeper split labels
    axis(1,at=xBreaks[i]+off,labels = abb,cex.axis=labCex)
    #Now add the larger axis
    axis(1,at=c(xBreaks[i]+off[1],0.5*(xBreaks[i]+xBreaks[i+1]),xBreaks[i]+off[length(off)]),labels = c(NA,levels(df$class)[i],NA),line=2.5,lwd.ticks=0,cex.axis=labCex)
  }
  #Finally add control label if needed
  if(!groupBySigs){
    tmp = isControl[levels(df$class)]
    #Do we need to draw it?
    if(any(tmp)){
      low = xBreaks[min(which(tmp))]
      high = xBreaks[max(which(tmp))+1]
      axis(1,at=c(low,0.5*(low+high),high),labels=c(NA,'Controls',NA),line=5,lwd.ticks=0,cex.axis=labCex)
    }
  }
  #Add axis label
  #title(xlab=ifelse(groupBySigs,'Signatures','TumourType'),line=5)
  #Add legends
  if(!groupBySigs){
    #Add legend
    legend(x=max(xx)+(max(xx)-min(xx))*0.05,
           y=max(yy),
           legend=levels(df$Signatures),
           col = cols,
           pch = pch,
           bty='n',
           title='Signatures',
           cex=cex,
           xpd=TRUE)
  }else{
    #Add extra heading to legend to show controls
    labs = levels(df$Signatures)
    cc = cols
    #Order so controls at the bottom
    o = order(isControl[labs])
    labs = labs[o]
    cc = cc[o]
    #Add in extra label
    cc = c(cc[seq(sum(!isControl))],NA,cc[-seq(sum(!isControl))])
    labs = c(labs[seq(sum(!isControl))],'Controls',labs[-seq(sum(!isControl))])
    #Add legend
    legend(x=max(xx)+(max(xx)-min(xx))*0.05,
           y=max(yy),
           legend=labs,
           col = cc,
           pch = pch,
           bty='n',
           title='TumourTypes',
           cex=cex,
           xpd=TRUE)
  }
}

#' Discrete colour set
#'
#' Given a set of discrete labels and a brewer pallette makes a named vector suitable for use with ComplexHeatmap
#'
#' @param labels Labels to use.
#' @param pal Brewer pallette.
#' @param na.rm Drop NAs
colSet = function(labels,pal,na.rm=TRUE){
  x = unique(labels)
  if(na.rm)
    x = x[!is.na(x)]
  setNames(brewer.pal(length(x),pal),x)
}

#' Cell signals heatmap
#'
#' Creates heatmap with pR2, and other meta-data
#'
#' @param norm Norm object.  Should
#' @param extras Extras object.
#' @param isControl Null sets Bench/ALL to controls.  If you want no controls set to FALSE.
#' @param colPal Colour scheme to use.  Range from 0 to 1.
csHeatmap = function(norm,extras,
                     isControl=NULL,
                     botAnno = list(),
                     topAnno = list(),
                     colPal = brewer.pal(9,'Greys'),
                     colPalPR2 = brewer.pal(9,'Reds'),
                     colPalIsControl = c('TRUE'='DarkRed','FALSE'='DarkGreen'),
                     limsNorm=c(0,1),
                     limsPR2=c(0.2,0.8),
                     ...
                     ){
  #Standard controls are Bench/ALL
  if(is.null(isControl))
    isControl = grepl('^SangerProjectBench',colnames(norm)) | grepl('^SangerProjectTARGET_ALL_phase1',colnames(norm))
  if(length(isControl)==1)
    isControl = rep(isControl,ncol(norm))
  #Construct the colour maps
  cMap = circlize::colorRamp2(seq(limsNorm[1],limsNorm[2],length.out=length(colPal)),colPal)
  cMapPR2 = circlize::colorRamp2(seq(limsPR2[1],limsPR2[2],length.out=length(colPalPR2)),colPalPR2)
  #Create meta-data track for pR2
  botAnno$pR2 = unlist(extras['pR2',])
  #Check if col exists, add it if not
  if(!'col' %in% names(botAnno))
    botAnno$col = list()
  botAnno$col$pR2 = cMapPR2
  bAnn = do.call(HeatmapAnnotation,botAnno)
  #Extra meta-data for top.  Suppress isControl track if all the same value
  if(length(unique(isControl))>1){
    topAnno$isControl = isControl
    if(!'col' %in% names(topAnno))
      topAnno$col = list()
    topAnno$col$isControl = colPalIsControl
  }
  if(length(topAnno)>0){
    tAnn = do.call(HeatmapAnnotation,topAnno)
  }else{
    tAnn=NULL
  }
  hh = list(matrix=as.matrix(norm),
            col=cMap,
            name='SigFrac',
            bottom_annotation = bAnn,
            top_annotation = tAnn,
            cluster_rows=FALSE,
            cluster_row_slices = FALSE,
            cluster_column_slices = FALSE,
            column_title = 'Bulk transcriptomes',
            row_gap = unit(5,'mm'),
            column_gap = unit(5,'mm'),
            show_column_names=FALSE,
            show_column_dend = FALSE,
            row_title_rot=0
            )
  #Update with values from the dots
  theDots = list(...)
  for(nom in names(theDots))
    hh[[nom]] = theDots[[nom]]
  do.call(Heatmap,hh)
}

#' Creates a heatmap showing top genes for each class
#'
#' Given a Seurat object and an annotation vector, gets the top markers and plots their expression.
#'
#' @param srat Seurat object to plot
#' @param annot Class vector to use.  If a factor, use levels to order.
annHeatmap = function(srat,annot=srat@meta.data$annot,...){
  if(!is.factor(annot))
    annot = factor(annot)
  #Get the top 5 to plot
  topMrks = SoupX::quickMarkers(srat@assays$RNA@counts,clusters = annot,expressCut = 0.9,N=5)
  #Get class averaged and re-normalised expression
  avgExpr = t(aggregate.Matrix(t(srat@assays$RNA@counts),groupings=annot))
  avgExpr = t(t(avgExpr)/colSums(avgExpr))
  avgExpr = log(1+1e4*avgExpr)
  avgExpr = avgExpr[,levels(annot)]
  mat = as.matrix(avgExpr[topMrks$gene,])
  #Row normalise
  mat = t(scale(t(mat)))
  hh = list(mat,
               name='RowNorm\nExpression',
               col = circlize::colorRamp2(seq(-3,3,length.out=length(divColsAlt)),divColsAlt),
               row_split = factor(topMrks$cluster,levels=levels(annot)),
               show_row_dend=FALSE,
               row_title_rot = 0,
               row_names_gp = gpar(fontsize = 8),
               row_gap = unit(2,'mm'),
               cluster_row_slices = FALSE,
               cluster_columns = FALSE,
               show_column_dend=FALSE,
               show_column_names = TRUE
               )
  theDots = list(...)
  for(nom in names(theDots)){
    hh[[nom]] = theDots[[nom]]
  }
  hh = do.call(Heatmap,hh)
  return(list(hh=hh,rawData=mat))
}


##############
# Parameters #
##############
outDir='Results/fits'
resDir='Results/virginFigs'
bulkAssayType = 'frag'
#Force recalculation istead of using previously saved data
forceRecalc=FALSE
#TARGET name codes
TGTtumourCodes = c('00'='Normal',
                   '52'='RT'
                   )
TGTtissueCodes = c('01'='Primary',
                   '02'='Secondary',
                   '03'='PrimaryBloodPB',
                   '04'='RecurrentBloodBM',
                   '06'='Metastatic',
                   '09'='PrimaryBloodBM',
                   '20'='CellLineControl',
                   '40'='RecurrentBloodPB',
                   '50'='CellLinePatient',
                   '11'='Normal'
                   )
#Cancer gene definitions
somaticGenes = list()
#From Treger, 2019 Onconephrology.  Listed in order of decreasing prevelance
somaticGenes$Wilms = c('WT1','CTNNB1','AMER1','IGF2','TP53','MYCN','DROSHA','DICER1','DGCR8','XPO5','DIS3L2','TARBP2','SIX1','SIX2','SMARCA4','MLLT1','BCORL1','COL6A3','NF1','BCOR','NONO','ARID1A','MAP3K4','MAX','ASXL1','BRD7','FGFR1','HDAC4','CHD4','ACTB')
#From Wegert, 2018 Nature Comms.
somaticGenes$CMN = c('EGFR','NTRK3','ETV6','NTRK1','NTRK2','BRAF','KIAA1549','TPR','LMNA')
#From Han, 2020, Pediatr Blood Cancer
somaticGenes$CCSK = c('BCOR','YWHAE','NUTM2B','NUTM2E','CCNB3')
#From Manley, 2017, Curr Opin Urol
somaticGenes$ccRCC = c('VHL', 'PBRM1', 'SETD2', 'BAP1', 'KDM5C', 'MTOR')
#MET is specific to Type 1, FH,CDKN2a Type2
somaticGenes$pcRCC = c('MET', 'SETD2', 'NF2', 'KDM6A', 'SMARCB1','FH','CDKN2A')
somaticGenes$chRCC = c('TP53','PTEN')
#MRT
somaticGenes$MRT = c('SMARCB1','SMARCA2','PBRM1','SMARCA4','ARID1A')
#Mouse signals to use, exclude external and pre-gast
mouseGold = c("mouseDev_Caudal_Mesoderm", "mouseDev_Caudal_neurectoderm", 
"mouseDev_Def_endoderm", "mouseDev_Intermediate_mesoderm", "mouseDev_Mixed_mesoderm", 
"mouseDev_Nascent_mesoderm", "mouseDev_Neural_crest", "mouseDev_NMP", 
"mouseDev_Paraxial_mesoderm", "mouseDev_Parietal_endoderm", "mouseDev_Pharyngeal_mesoderm", 
"mouseDev_Rostral_neurectoderm", "mouseDev_Somitic_mesoderm", 
"mouseDev_Surface_ectoderm", "mouseDev_Blood_progenitors_1", 
"mouseDev_Blood_progenitors_2", "mouseDev_Cardiomyocytes", "mouseDev_Endothelium", 
"mouseDev_Erythroid1", "mouseDev_Erythroid2", "mouseDev_Erythroid3", 
"mouseDev_Forebrain_Midbrain_Hindbrain", "mouseDev_Gut", "mouseDev_Haematoendothelial_progenitors", 
"mouseDev_Mesenchyme", "mouseDev_Notochord", "mouseDev_Spinal_cord", 
"mouseDev_Caudal_Mesoderm", "mouseDev_Def_endoderm", "mouseDev_Intermediate_mesoderm", 
"mouseDev_Mixed_mesoderm", "mouseDev_Nascent_mesoderm", "mouseDev_Neural_crest", 
"mouseDev_Paraxial_mesoderm", "mouseDev_Parietal_endoderm", "mouseDev_Pharyngeal_mesoderm", 
"mouseDev_Somitic_mesoderm", "mouseDev_Surface_ectoderm")
#Genes to exclude when performing fits
gns = read.table('Data/scData/fAd/babyAdrenals/babyAdrenal1/filtered_gene_bc_matrices/GRCh38/genes.tsv',sep='\t',header=FALSE)
gnsMap = gns
hkGeneREGEX='^(EIF[0-9]|RPL[0-9]|RPS[0-9]|RPN1|POLR[0-9]|SNX[0-9]|HSP[AB][0-9]|H1FX|H2AF[VXYZ]|PRKA|NDUF[ABCSV]|ATP[0-9]|PSM[ABCDEFG][0-9]|UBA[0-9]|UBE[0-9]|USP[0-9]|TXN)'
coreExcludeGenes = unique(c(grep('\\.[0-9]+_',gns[,2],value=TRUE), #Poorly characterised
                        grep('MALAT1',gns[,2],value=TRUE), #Contamination
                        grep('^HB[BGMQDAZE][12]?_',gns[,2],value=TRUE), #Contamination
                        grep('^MT-',gns[,2],value=TRUE), #Mitochondria
                        grep(hkGeneREGEX,gns[,2],value=TRUE) #Housekeeping genes
                        ))
coreExcludeGenes = unique(unlist(as.list(geneSets)[c('riboGenes','hspGenes','mtGenes')]))
gnsExclude = gns[gns$V2 %in% coreExcludeGenes,]
weightVec = setNames(rep(0,nrow(gnsExclude)),gnsExclude$V1)

##################
# Colour schemes #
##################
########################
# Random colour schemes
divColsAlt = c("#053061", "#2166ac", "#4393c3", "#92c5de", "#d1e5f0", "#f7f7f7","#fddbc7", "#f4a582", "#d6604d", "#b2182b", "#67001f")
###################
# Sample colouring
sampCols = c('#1b9e77','#d95f02','#7570b3')
sampCols = list(Normal = col2hsv(sampCols[1]),
                PaedTumour = col2hsv(sampCols[2]),
                AdultTumour = col2hsv(sampCols[3])
                )
sampColsName = list(Normal=c('NormalAdult','NormalPaed','NormalFoetal'),
                    PaedTumour = c('CCSK','CMN','MRT','Wilms'),
                    AdultTumour = c('ccRCC','pRCC','ChRCC')
                    )
#Create colours
sampCols = unlist(lapply(names(sampCols),function(e) setNames(hsv(sampCols[[e]][1],seq(0.5,1.0,length.out=length(sampColsName[[e]])),seq(0.5,1.0,length.out=length(sampColsName[[e]]))),sampColsName[[e]])))
sampCols = setNames(c('#a7cee2','#f69999','#b4d88a','#cab2d6','#fdbf6e','#e21f26','#34a048','#faeb30','#2078b4','#6b3e98','#f57f20','#b15a28','#f57f20','#b15a28'),
                    c('Immune','ALL','NormalAdult','NormalPaed','NormalFoetal','CCSK','CMN','MRT','Wilms','ccRCC','pRCC','ChRCC','ATRT','Neuroblastoma')
                    )
#Aliases
sampCols['chRCC'] = sampCols['ChRCC']
sampCols['pcRCC'] = sampCols['pRCC']
#################
# Signal colours
#First define broad groupings
sigHues = list(
            Vascular = c(0.0,0.0),#This is really a proxy for c(0.95,0.05) but the wrapping around is complex
            Adrenal = c(.05,.2),
            Kidney = c(.2,.5),
            Immune=c(.5,.75),
            Fibroblast = c(.75,.95)
            )
#Next level of split
sigHues[['Immune']] = list('.'=sigHues[['Immune']],
                           Myeloid = c(0.0,0.45),
                           Lymphoid= c (0.45,0.9),
                           HSC = c(0.9,1.0)
                           )
sigHues[['Kidney']] = list('.' = sigHues[['Kidney']],
                           Nephron = c(0,0.8),
                           CollectingDuct = c(0.8,1.0)
                           )
sigHues[['Adrenal']] = list('.' = sigHues[['Adrenal']],
                            Medulla = c(0,0.5),
                            Mesenchyme = c(0.5,0.8),
                            Cortex = c(0.8,1.0)
                            )
sigHues[['Fibroblast']] = list('.' = sigHues[['Fibroblast']],
                               ICa = c(0,0.35),
                               ICb = c(0.35,0.6),
                               MSC = c(0.6,1.0)
                               )
#Third split
sigHues[['Adrenal']][['Medulla']] = list('.' = sigHues[['Adrenal']][['Medulla']],
                                         SCPs = c(0,0.25),
                                         Bridge = c(0.25,0.5),
                                         Chromaffin = c(0.5,0.75),
                                         Sympathoblastic = c(0.75,1)
                                         )
sigHues[['Immune']][['Lymphoid']] = list('.' = sigHues[['Immune']][['Lymphoid']],
                                         B = c(0,0.4),
                                         'T' = c(0.4,0.8),
                                         NK = c(0.8,1)
                                         )
sigHues[['Immune']][['Myeloid']] = list('.' = sigHues[['Immune']][['Myeloid']],
                                        DC = c(0,0.2),
                                        Mast = c(0.2,0.4)
                                        )
sigHues[['Kidney']][['Nephron']] = list('.' = sigHues$Kidney$Nephron,
                                        CapMesenchyme = c(0,0.1),
                                        Glomerulus = c(0.3,0.4),
                                        ProximalTubules = c(0.4,0.8),
                                        LoopOfHenle = c(0.8,0.9),
                                        DistalTubules = c(0.9,1.0)
                                        )
sigHues[['Kidney']][['CollectingDuct']] = list('.' = sigHues$Kidney$CollectingDuct,
                                               Pelvis = c(0,0.2),
                                               TypeA = c(0.2,0.55),
                                               TypeB = c(0.55,0.9),
                                               Ureter = c(0.9,1.0)
                                               )
#Final split
sigHues$Kidney$Nephron$ProximalTubules = list('.' = sigHues$Kidney$Nephron$ProximalTubules,
                                              PT1 = c(0,0.33),
                                              PT2 = c(0.33,0.66),
                                              PT3 = c(0.66,1)
                                              )
#Evenly space fine grained cell types
#
#Translate relative hues into absolute
sigHues$.=c(0,1)
adjRange = function(absRange,relRange) absRange[1]+diff(absRange)*relRange
relToAbs = function(x){
  for(nom in names(x)){
    if(nom=='.')
      next
    #Is this a leaf?
    if(!is.list(x[[nom]])){
      x[[nom]] = adjRange(x$.,x[[nom]])
    }else{
      #Not a leaf, adjust top-level and descend
      x[[nom]]$. = adjRange(x$.,x[[nom]]$.)
      x[[nom]] = relToAbs(x[[nom]])
    }
  }
  return(x)
}
sigHues = relToAbs(sigHues)
#Build the colours
tst = melt(sigHues)
#Average entries
tmp = aggregate(value ~ apply(tst[,-1],1,paste,collapse='___'),tst,mean)
m = match(tmp[,1],apply(tst[,-1],1,paste,collapse='___'))
tst$value[m] = tmp$value
tst = tst[m,]
tst[tst=='.']=NA
tst = tst[!is.na(tst$L1),c(sort(grep('^L[0-9]+$',colnames(tst),value=TRUE)),'value')]
tst$name=NA
for(nom in sort(grep('^L[0-9]+$',colnames(tst),value=TRUE),decreasing=FALSE)){
  w = which(!is.na(tst[,nom]))
  tst$name[w] = tst[w,nom]
}
rownames(tst) = tst$name
colnames(tst) = gsub('value','hue',colnames(tst))
#Add some aliases
sigCols = tst
tmp = sigCols[c('DistalTubules','Glomerulus','Nephron','Nephron','Nephron','CapMesenchyme','Glomerulus','TypeA','TypeB','Immune','Fibroblast','Fibroblast'),]
tmp$name = c('UretericBud','PrimitiveVesicle','otherNephron','EarlyTubule','Tubules','NephronProgenitor','Vesicle','IntercalatedTypeA','IntercalatedTypeB','Leucocytes','Interstitial','InterstitialMesenchyme')
rownames(tmp) = tmp$name
sigCols = rbind(sigCols,tmp)
#Get depth of each entry
sigCols$level = factor(rowSums(!is.na(sigCols[,grep('^L[0-9]+$',colnames(sigCols),value=TRUE)])))
#Create colours - Hue specified, value separates mature from foetal, saturation is for fine tuning
sigCols$sats = 0.6
sigCols$sats[sigCols$hue==sigCols$hue[sigCols$name=='PT1']] = 1.0
sigCols$sats[sigCols$hue==sigCols$hue[sigCols$name=='PT2']] = 0.6
sigCols$sats[sigCols$hue==sigCols$hue[sigCols$name=='PT3']] = 0.2
sigCols$sats[sigCols$hue==sigCols$hue[sigCols$name=='Nephron']] = 0.4
sigCols$sats[sigCols$hue==sigCols$hue[sigCols$name=='TypeA']] = 0.3
sigCols$sats[sigCols$hue==sigCols$hue[sigCols$name=='TypeB']] = 0.9
sigCols$sats[sigCols$hue==sigCols$hue[sigCols$name=='SchwannPrecursor']] = 0.3
#Foetal version
tmp = sigCols
tmp$colour = hsv(tmp$hue,tmp$sats,0.9)
tmp$development = 'Foetal'
rownames(tmp) = paste0(rownames(tmp),'_Foetal')
#Mature version
sigCols$colour = hsv(sigCols$hue,sigCols$sats,0.7)
sigCols$development = 'Mature'
rownames(sigCols) = paste0(rownames(sigCols),'_Mature')
#Merge
sigCols = rbind(sigCols,tmp)
#Manually add colours
dummy = sigCols[1,]
dummy[1,]=NA
rownames(dummy)=NULL
tmp=dummy
extras = dummy[rep(1,1000),,drop=FALSE]
extras[1,c('L1','name','development','colour')]=c('Other','Intercept','Mature','#CCCCCC')
extras[2,c('L1','name','development','colour')]=c('Other','Intercept','Foetal','#CCCCCC')
extras = extras[!is.na(extras$colour),]
rownames(extras) = paste0(extras$name,'_',extras$development)
#Alias for intercept
extras2 = extras
extras2$name='Unexplained'
rownames(extras2) = gsub('Intercept','Unexplained',rownames(extras2))
sigCols = rbind(sigCols,extras,extras2)
#Order colours sensibly
o = do.call(order,c(
                    decreasing=TRUE,
                    list(sigCols$development),
                    list(sigCols$level),
                    lapply(grep('^L[0-9]+$',colnames(sigCols),value=TRUE),function(e) sigCols[,e]),
                    list(sigCols$name)
                    ))
sigCols = sigCols[o,]
#Visualise them sensibly
#First order appropriately
sigCols = sigCols[order(factor(sigCols$L1,levels=c('Immune','Adrenal','Kidney','Vascular','Fibroblast','Other')),
                        sigCols$L2,
                        sigCols$level,
                        sigCols$hue,
                        sigCols$name,
                        sigCols$development
                        ),]


###########################
# Define single cell data #
###########################
#Load Seurat objcets
rcc = readRDS('Results/preProcess/scData/Tumour/processedSeurat_RCC.RDS')
fKid = readRDS('Results/preProcess/scData/finalisedSeurat_fKid.RDS')
fAd = readRDS('Results/preProcess/scData/finalisedSeurat_fAd.RDS')
pbmc = readRDS('Results/preProcess/scData/finalisedSeurat_PBMC.RDS')
sciKidRef = readRDS('Results/preProcess/scData/finalisedSeurat_sciKidRef.RDS')
mrtFresh = readRDS('Results/preProcess/scData/Tumour//processedSeurat_MRT_fresh.RDS')
mrtNuc = readRDS('Results/preProcess/scData/Tumour//processedSeurat_MRT_nuc.RDS')
mrtOrg = readRDS('Results/preProcess/scData/Tumour//processedSeurat_MRT_org.RDS')
#################
# CCSK annotiion
ccsk = readRDS('Results/preProcess/scData/Tumour//processedSeurat_CCSK.RDS')
ccsk2 = readRDS('Results/preProcess/scData/Tumour//processedSeurat_CCSK_extra.RDS')
gns = intersect(rownames(ccsk),rownames(ccsk2))
ccsk = cbind(ccsk@assays$RNA@counts[gns,],ccsk2@assays$RNA@counts[gns,])
ccsk = scQC$quickCluster(ccsk)
annot = list(End=c(24,14),
             Leucocytes=c(10,15,20,23),
             TumourA = c(1,0,5,4,22),
             TumourB = c(19,8,9),
             TumourC = c(2,3,6,7,11),
             TumourD = c(16,17),
             TumourE = c(12,13,18,21)
             )
annot = setNames(rep(names(annot),lengths(annot)),unlist(annot))
ccsk@meta.data$annot = annot[as.character(ccsk@meta.data$seurat_clusters)]
###################
# Wilms annotation
wilms = readRDS('Results/preProcess/scData/Tumour//processedSeurat_Wilms.RDS')
annot = list('TumourA'=c(5,7,9,10,12,13),
             'TumourB'=c(0,1,2,3,6,8,11,16,15,4),
             Leucocytes=14
             )
annot = setNames(rep(names(annot),lengths(annot)),unlist(annot))
wilms@meta.data$annot = annot[as.character(wilms@meta.data$seurat_clusters)]
#################
# CMN annotation
cmn = readRDS('Results/preProcess/scData/Tumour/processedSeurat_CMN.RDS')
annot = list('Tumour'=c(11,16),
             Adrenal = c(9,10),
             End = c(7,12),
             'T' = c(1,2,4),
             B = c(6,15),
             NK = c(0,5,14),
             Monocytes = c(3,8,13)
             )
annot = setNames(rep(names(annot),lengths(annot)),unlist(annot))
cmn@meta.data$annot = annot[as.character(cmn@meta.data$seurat_clusters)]
###############
# Combined MRT
mrt = cbind(mrtFresh@assays$RNA@counts,mrtNuc@assays$RNA@counts,mrtOrg@assays$RNA@counts)
mrt = scQC$quickCluster(mrt)
tmp = list('T1'=c(0,1,4),
           'T2'=c(8,12,13),
           'T3'=c(7,19,20),
           'T4'=c(21),
           'T5'=c(6,9,10,11,14,15),
           'Stroma' = c(2,3,5,16),
           'L1' = c(18),
           'L2' = c(17),
           'L3' = c(22,23)
)
tmp = setNames(rep(names(tmp),times=lengths(tmp)),unlist(tmp))
mrt@meta.data$annot = tmp[as.character(mrt@meta.data$seurat_clusters)]
mrt@meta.data$type = ifelse(grepl('MRT_fresh',colnames(mrt)),
                            'Fresh',
                            ifelse(grepl('MRT_org',colnames(mrt)),
                                   'Organoid',
                                   'Nuclear'
                                   )
                            )

#####################
# Get nuclear genes #
#####################
dat = list()
dat$MRT = mrtNuc@assays$RNA@counts
dat$CCSK = ccsk@assays$RNA@counts[,grep('IM[0-9]+',ccsk@meta.data$orig.ident)]
gns1 = rownames(dat$MRT)[rowSums(dat$MRT>0)>10]
gns2 = rownames(dat$CCSK)[rowSums(dat$CCSK>0)>10]
gns = union(gns1,gns2)
#Convert to ENSEMBL
tmp = wilms@misc$geneMapCommon
nucGns = data.frame(ID = tmp[match(gns,tmp[,2]),1],sym = gns)


############################
# Define common references #
############################
#Define annots in sciKidRef for mature kidney
mKidAnnots = paste0('sciKidRef_',c('G', "PT1", "PT2","PT3",'H', "D",  "C1", "C2", "M",'F', "AV1", "AV2", "DV", "GE"))
#Define and process/load references.
refs = list(HCL = 'HCL',
            mouseDev = 'mouseDev',
            mouseDevPBMC = c('mouseDev','PBMC'),
            mouseDevExtra = 'mouseDev_extra',
            mouseDevExtraPBMC = c('mouseDev_extra','PBMC'),
            benchSet = c('PBMC','sciKidRef'),
            kidPBMCandMast = c('fKid','sciKidRef','PBMC'),
            mKidPBMCandMast = c('sciKidRef','PBMC'),
            allPBMC = c('fKid','sciKidRef','fAd','PBMC'),
            kidPBMC = c('fKid','sciKidRef','PBMC'),
            mKidPBMC = c('sciKidRef','PBMC'),
            fKidPBMCNuc = c('fKid','PBMC'),
            fKidPBMC = c('fKid','PBMC'),
            fAdPBMC = c('fAd','PBMC'), 
            fKidOnly = 'fKid',
            mKidOnly = 'sciKidRef',
            fAdOnly = 'fAd',
            PBMC = 'PBMC'
)
refs = rev(refs)
refs = lapply(refs,function(e) list(src=e))
for(nom in names(refs)){
  bigTgtFile = file.path(outDir,'refs',paste0('completeObject_',nom,'.RDS'))
  if(!forceRecalc && file.exists(bigTgtFile)){
    message(sprintf("Loading reference set %s",nom))
    refs[[nom]] = readRDS(bigTgtFile)
  }else{
    message(sprintf("Pre-processing reference set %s",nom))
    #Save exactly where to get things
    refs[[nom]]$mtx = file.path('Results/preProcess/scData',refs[[nom]]$src)
    mtx = lapply(refs[[nom]]$mtx,function(e) loadRefMtx(e))
    gns = Reduce(intersect,lapply(mtx,rownames))
    #And the set of genes
    refs[[nom]]$gns = gns
    mtx = do.call(cbind,lapply(mtx,function(e) e[gns,]))
    #Custom sets
    if(nom=='benchSet'){
      #Keep immune and add PT1
      mtx = mtx[,grepl('^PBMC_',colnames(mtx)) | grepl('sciKidRef_PT1',colnames(mtx))]
    }
    #Pull mature kidney out of sciKidRef
    if(grepl('^mKid',nom) | nom %in% c('kidPBMCandMast','allPBMC','kidPBMC')){
      #Drop those that aren't in the kidney
      mKidTgts = mKidAnnots
      #Add Mast cells if we need them
      if(grepl('Mast',nom)){
        mKidTgts = c(mKidTgts,'sciKidRef_MST','sciKidRef_tMST1','sciKidRef_tMST2')
      }
      #Drop anything from sciKidRef that isn't in the approved list
      mtx = mtx[,!grepl('^sciKidRef',colnames(mtx)) | gsub(':.*','',colnames(mtx)) %in% mKidTgts]
    }
    #Train logistic regression model
    classes = gsub(':.*','',colnames(mtx))
    #Clean class labels 
    classes = gsub('/.*','',classes)
    #Drop anything with too few observations to train with
    w = (gsub('^[^_]+_','',classes)!='NA' & table(classes)[classes]>20)
    if(nom %in% c('mouseDev','mouseDevPBMC'))
      w = w & (grepl('^PBMC',classes) | classes %in% mouseGold)
    w = which(w)
    nWorkers = NULL
    tgtFile = file.path(outDir,'refs',paste0('logisticRegression_',nom,'.RDS'))
    if(!forceRecalc && file.exists(tgtFile)){
      refs[[nom]]$lrFit = readRDS(tgtFile)
    }else{
      message('Fitting logistic regression')
      tmp = mtx[(!rownames(mtx) %in% gnsExclude$V1),w]
      if(grepl('Nuc$',nom))
        tmp = tmp[rownames(mtx) %in% nucGns$ID,]
      refs[[nom]]$lrFit= lr$trainModel(tmp,classes=classes[w],maxCells=5000)
      saveRDS(refs[[nom]]$lrFit,tgtFile)
    }
    #Make LR confusion matrix.  This is fast now, ne need to do the save/load loop.
    confMtx = lr$predictSimilarity(refs[[nom]]$lrFit,mtx,classes=classes)
    #tgtFile = file.path(outDir,'refs',paste0('confusionMatrix_',nom,'.RDS'))
    #if(!forceRecalc && file.exists(tgtFile)){
    #  confMtx = readRDS(tgtFile)
    #}else{
    #  message('Building confusion matrix')
    #  saveRDS(confMtx,tgtFile)
    #}
    confMtx = confMtx[,order(colnames(confMtx))]
    confMtx = confMtx[order(rownames(confMtx)),]
    refs[[nom]]$confMat = confMtx
    #And plot it.
    pdf(file.path(outDir,'refs',paste0('confusionMatrix_',nom,'.pdf')),width=14,height=14)
    hh = lr$similarityHeatmap(confMtx,
                              cluster_rows=FALSE,
                              show_row_names=TRUE,
                              column_split=gsub('_.*','',colnames(confMtx)),
                              row_split=gsub('_.*','',rownames(confMtx))
    )
    draw(hh)
    dev.off()
    #Make cluster level summaries for deconvolution
    clustSum = t(aggregate.Matrix(t(mtx[,w]),classes[w]))
    clustSum = t(t(clustSum)/colSums(clustSum))
    clustSum = as.data.frame(as.matrix(clustSum))
    clustSum = cbind(geneName=rownames(clustSum),clustSum)
    refs[[nom]]$clustSum = clustSum
    refs[[nom]]$clustSumFile = file.path(outDir,'refs',paste0('clustSummary_',nom,'.tsv'))
    #Write it out
    write.table(clustSum,refs[[nom]]$clustSumFile,row.names=FALSE,col.names=TRUE,sep='\t',quote=FALSE)
    #And define a random subset if we're going to do per cell fitting
    refs[[nom]]$w = w[lr$smartSubset(classes[w],minSize=200)]
    saveRDS(refs[[nom]],bigTgtFile)
  }
}

####################
# Define bulk data #
####################
#The output will have a bunch of source specific columns, but must have:
#source - Where did this sample come from.
#tissue - What is the tissue.
#disease - What is the disease status of the individual.
#biopsy - What type of biopsy is it (normal,tumour,etc).
#subtype - What subtype is it.
#outcome - Optional.  Outcome data if available.
#UniqueSampleID - A unique sample ID.
#tgtFile - Where to get the associated file.
#toKeep - Use this file?
cmnCols = c('dataset','source','tissue','disease','biopsy','subtype','outcome','UniqueSampleID','tgtFile','toKeep')
bulkMani = list()
########
# Wilms
#Work out which samples we're going to get.
maniWilms = read.table('Data/SangerWilmsManifest.txt',sep='\t',header=TRUE)
maniWilms$UniqueSampleID = paste0('SangerProject1700_',gsub('^PD','PR',maniWilms$Sample))
#Add the fixes
tmp = split(maniWilms$Tumour_type,maniWilms$Case)
tmp = lapply(tmp,function(e) unique(e[e!='-']))
tmp = sapply(tmp,function(e) ifelse(length(e)==1,e,'Normal'))
maniWilms$Tumour_type = tmp[maniWilms$Case]
yrs = as.numeric(gsub('(([0-9]+) years)?.*','\\2',maniWilms$Age))
mnths = as.numeric(gsub('(.*years (and )?)?(([0-9]+) months)?.*','\\4',maniWilms$Age))
maniWilms$AgeMonths = ifelse(is.na(yrs) & is.na(mnths),NA,
                                   ifelse(is.na(yrs),0,yrs*12) +
                                   ifelse(is.na(mnths),0,mnths)
                                   )
maniWilms$Disease = ifelse(maniWilms$Tumour_type=='Nephroblastoma','Wilms','ccRCC')
maniWilmsSanger = maniWilms
#Drop the ones we don't have
maniWilmsSanger = maniWilmsSanger[file.exists(file.path('Results/bulkRNAseq/SangerProject1700',bulkAssayType,paste0(maniWilmsSanger$UniqueSampleID,'.tsv'))),]
#Add in the extra data from Reem
tmp = read.table('Data/WilmsInhouseMeta.txt',sep='\t',header=TRUE)
m = match(maniWilmsSanger$Case,tmp$PD_ID)
maniWilmsSanger$Bilateral = tmp$Bilateral[m]
maniWilmsSanger$Presentation = ifelse(tmp$Localised.Metastatic[m]==1,
                                      'Localised',
                                      ifelse(tmp$Localised.Metastatic[m]==2,
                                             'CTmets',
                                             ifelse(tmp$Localised.Metastatic[m]==3,
                                                    'Metastatic',
                                                    tmp$Localised.Metastatic[m]
                                                    )
                                             )
                                      )
maniWilmsSanger$Relapsed = tmp$Relapse[m]
maniWilmsSanger$outcome = c('NOT RECIEVED'=NA,'1'='Alive','2'='Disease','3'='Dead')[tmp$A.D[m]]
#And add the target wilms
tgts = gsub('\\.tsv','',list.files(file.path('Results/bulkRNAseq/TARGET-WT',bulkAssayType)))
mDat = read.table('Data/TARGET_WT_Discovery_ClinicalData_20170525.txt',sep='\t',header=TRUE)
m = match(gsub('.*_(TARGET-[0-9]+-[A-Z]+)-.*','\\1',tgts),mDat$TARGET.USI)
mDat = mDat[m,]
rownames(mDat) = tgts
mDat$TissueCode = TGTtissueCodes[gsub('.*_TARGET-([0-9]+)-([A-Z]+)-([0-9]+)([A-Z])-([0-9]+[A-Z])','\\3',tgts)]
mDat$TissuePortion = gsub('.*_TARGET-([0-9]+)-([A-Z]+)-([0-9]+)([A-Z])-([0-9]+[A-Z])','\\4',tgts)
mDat$NucleicAcid = gsub('.*_TARGET-([0-9]+)-([A-Z]+)-([0-9]+)([A-Z])-([0-9]+[A-Z])','\\5',tgts)
#Legacy annotation
mDat1 = read.table('Data/TARGET_WT_SampleMatrix_Discovery_20190212.txt',sep='\t',header=TRUE)
mDat2 = read.table('Data/TARGET_WT_Percent_Tumor_Nuclei_and_Necrosis_20170525.txt',sep='\t',header=TRUE)
mDat3 = read.table('Data/TARGET_WT_Discovery_ClinicalData_20170525.txt',sep='\t',header=TRUE)
samps = mDat$TARGET.USI
#Now try and find a match for each of these
m1 = sapply(samps,function(e) grep(e,mDat1$Diagnostic.Tumor.RNA.Sample.ID))
m2 = sapply(samps,function(e) grep(e,mDat1$Matched.Normal.RNA.Sample.ID))
m3 = sapply(samps,function(e) grep(e,mDat1$Relapse.Tumor.RNA.Sample.ID))
m = rep(NA,length(samps))
m[lengths(m1)==1] = unlist(m1[lengths(m1)==1])
m[lengths(m2)==1] = unlist(m2[lengths(m2)==1])
m[lengths(m3)==1] = unlist(m3[lengths(m3)==1])
sampClass = ifelse(lengths(m1)==1,
                   'Tumour',
                   ifelse(lengths(m2)==1,
                          'Normal',
                          'Secondary'
                          )
                   )
#And add age where available
mm = match(mDat1$Case.USI[m],mDat2$TARGET.Case.USI)
mmm = match(mDat1$Case.USI[m],mDat3$TARGET.USI)
maniWilmsTARGET = cbind(mDat,mDat1[m,],mDat2[mm,],mDat3[mmm,])
maniWilmsTARGET$UniqueSampleID = rownames(maniWilmsTARGET)
#Mark the ones to keep
maniWilmsTARGET$toKeep = maniWilmsTARGET$TissueCode=='Primary'
tmp = split(maniWilmsSanger$UniqueSampleID,maniWilmsSanger$Case)
tmp = lapply(tmp,function(e) {
               #Special cases
               if(any(grepl('40718',e))){
                 return(grep('40718c',e,value=TRUE))
               }else if(any(grepl('40641',e))){
                 return(grep('40641m',e,value=TRUE))
               }else if(any(grepl('36159',e))){
                 return(grep('36159c',e,value=TRUE))
               }else if(any(grepl('24242',e))){
                 return(grep('24242c',e,value=TRUE))
               }else if(any(grepl('36165',e))){
                 return(grep('36165d',e,value=TRUE))
               }
               #Get a if it's there
               if(any(grepl('a$',e))){
                 return(grep('a$',e,value=TRUE))
               }else{
                 return(grep('b$',e,value=TRUE))
               }
                   })
maniWilmsSanger$toKeep = maniWilmsSanger$UniqueSampleID %in% unlist(tmp)
maniWilmsSanger$toKeep[maniWilmsSanger$Disease=='ccRCC']=FALSE
#Finally, get the samples that we want
maniWilmsSanger$tgtFile = file.path('Results/bulkRNAseq/SangerProject1700',bulkAssayType,paste0(maniWilmsSanger$UniqueSampleID,'.tsv'))
maniWilmsTARGET$tgtFile = file.path('Results/bulkRNAseq/TARGET-WT',bulkAssayType,paste0(maniWilmsTARGET$UniqueSampleID,'.tsv'))
#Do the join
maniWilmsSanger$source = 'GOSH'
maniWilmsSanger$tissue = 'Kidney'
colnames(maniWilmsSanger)= gsub('^Disease$','disease',colnames(maniWilmsSanger))
maniWilmsSanger$biopsy = maniWilmsSanger$Sample_type_1
maniWilmsSanger$subtype = maniWilmsSanger$Tumour_histology
maniWilmsSanger$dataset = 'WilmsSanger'
maniWilmsTARGET$source = 'TARGET'
maniWilmsTARGET$tissue = 'Kidney'
maniWilmsTARGET$disease = 'Wilms'
maniWilmsTARGET$biopsy = maniWilmsTARGET$TissueCode 
maniWilmsTARGET$subtype = maniWilmsTARGET$Histology.Classification.of.Primary.Tumor 
maniWilmsTARGET$outcome = maniWilmsTARGET$Vital.Status
maniWilmsTARGET$dataset = 'WilmsTARGET'
bulkMani[['WilmsSanger']] = maniWilmsSanger[,c(cmnCols,setdiff(colnames(maniWilmsSanger),cmnCols))]
bulkMani[['WilmsTARGET']] = maniWilmsTARGET[,c(cmnCols,setdiff(colnames(maniWilmsTARGET),cmnCols))]
#######
# CCSK
mDat = read.delim('Data/TARGET_CCSK_Discovery_ClinicalData_20170525.txt',sep='\t',header=TRUE)
mDat$tgtFile = file.path('Results/bulkRNAseq/SangerProjectCCSK',bulkAssayType,paste0('SangerProjectCCSK_',mDat$SRR_ID,'.tsv'))
mDat$source='TARGET'
mDat$tissue = 'Kidney'
mDat$disease = 'CCSK'
mDat$biopsy = 'Tumour'
mDat$subtype = NA
mDat$outcome = mDat$Reason.for.death
mDat$UniqueSampleID = paste0('SangerProjectCCSK_',mDat$SRR_ID)
mDat$toKeep=TRUE
mDat$dataset='CCSK'
bulkMani[['CCSK']] = mDat[,c(cmnCols,setdiff(colnames(mDat),cmnCols))]
######
# CMN
mDat = read.table('Data/SangerCMNManifest.txt',sep='\t',header=TRUE)
mDat$UniqueSampleID = paste0('SangerProject1716_',mDat$RNA_ID)
mDat = mDat[!mDat$RNA_ID %in% c('PR37581a','PR37582a','PR37218a'),]
colnames(mDat) = gsub('^Disease$','disease',colnames(mDat))
colnames(mDat) = gsub('^Subtype$','subtype',colnames(mDat))
mDat$source = 'Sanger'
mDat$tissue = 'Kidney'
mDat$biopsy = 'Tumour'
mDat$outcome = NA
mDat$toKeep=TRUE
mDat$tgtFile = file.path('Results/bulkRNAseq/SangerProject1716/',bulkAssayType,paste0(mDat$UniqueSampleID,'.tsv'))
mDat$dataset='CMN'
bulkMani[['CMN']] = mDat[,c(cmnCols,setdiff(colnames(mDat),cmnCols))]
######
# RCC
rc2 = readRDS('Data/bulkRNAseq/rse_gene_TCGA.RDS')
tcgaMani = read.table('Data/TCGA_RCC_update.txt',sep='\t',header=TRUE)
tcgaMani = tcgaMani[!is.na(tcgaMani[,1]),]
mDat = colData(rc2)
rownames(mDat) = paste0('TCGA_',rownames(mDat))
#Add in the updated RCC metadata where available
m = match(mDat$xml_bcr_patient_barcode,tcgaMani$bcr_patient_barcode)
colnames(tcgaMani) = paste0('rccUpdate_',colnames(tcgaMani))
mDat = cbind(mDat,tcgaMani[m,])
mDat$hasUpdate = !is.na(m)
mDat$UniqueSampleID = rownames(mDat)
mDat$age = mDat$cgc_case_age_at_diagnosis*12
mDat$biopsy = mDat$gdc_cases.samples.sample_type
mDat$tissue = as.character(mDat$xml_tumor_tissue_site)
mDat$subtype = ifelse(is.na(mDat$rccUpdate_PanKidney.Pathology),
                      as.character(mDat$xml_histological_type),
                      mDat$rccUpdate_PanKidney.Pathology
                      )
mDat$subtype[mDat$rccUpdate_Additional.Pathology.Data=='Metabolically Divergent (MD-)ChRCC '] = 'chRCC MD'
mDat$stage = mDat$rccUpdate_pathologic_T
mDat$outcome = ifelse(mDat$cgc_case_vital_status=='Dead',
                      'Dead',
                      mDat$cgc_case_tumor_status
                      )
tmp = c(ccRCC='ccRCC',ChRCC='chRCC','chRCC MD'='chRCC','KIRP CIMP'='pcRCC','PRCC T1'='pcRCC','PRCC T2'='pcRCC','PRCC Unc'='pcRCC')[mDat$subtype]
mDat$disease = ifelse(is.na(tmp),
                      mDat$subtype,
                      tmp
                      )
mDat$disease[mDat$disease=='Kidney Clear Cell Renal Carcinoma'] ='ccRCC'
mDat$disease[mDat$disease=='Kidney Papillary Renal Cell Carcinoma'] ='ccRCC'
mDat$toKeep = mDat$hasUpdate
mDat$source='TCGA'
#tcgaMani = mDat
#maniRCC = read.table('Data/TCGA_BulkMetadata.tsv',sep='\t',header=TRUE)
#maniRCC = maniRCC[which(maniRCC$Tissue=='Kidney'),]
#m = match(maniRCC$UniqueSampleID,gsub('\\.','-',mDat$UniqueSampleID))
#maniRCC$SampleType = mDat$gdc_cases.samples.sample_type[m]
#maniRCC$Disease2 = tcgaMani$rccUpdate_PanKidney.Pathology[m]
#maniRCC$OldDisease = maniRCC$Disease
#maniRCC$Disease = c(ccRCC='ccRCC',ChRCC='chRCC','KIRP CIMP'='chRCC','PRCC T1'='pcRCC','PRCC T2'='pcRCC','PRCC Unc'='pcRCC')[maniRCC$Disease2]
#Add mutations
mDat$nMuts = as.numeric(mDat$rccUpdate_Mutations.Total)
#Group by severity
doubleHit = c('Frame_shift_Del x2',
              'Frame_Shift_Del x2/Missense_Mutation',
              'Frame_Shift_Del/Missense_Mutation',
              'Frame_Shift_Del/Missense_Mutation x2',
              'Frame_Shift_Del/Nonsense_Mutation',
              'Frame_Shift_Del/Splice_Site',
              'Frame_Shift_Ins/Frame_Shift_Del',
              'Frame_Shift_Ins/Splice_Site',
              'In_Frame_Del x2',
              'In_Frame_Del/Missense_Mutation',
              'Missense_Mutation x2',
              'Nonsense_Mutation x2',
              'Nonsense_Mutation x2/Frame_Shift_Del',
              'Nonsense_Mutation/Frame_Shift_Del',
              'Nonsense_Mutation/Missense_Mutation',
              'Nonsense_Mutation/Missense_Mutation x4',
              'Nonsense_Mutation/Splice_Site',
              'Splice_Site/Missense_Mutation',
              'Splice_Site Missense_Mutation')
nonsense = c('Frame_shift_Del',
             'Frame_Shift_Del',
             'Frame_Shift_Ins',
             'Nonsense_Mutation')
splice=c('Splice_Site',
         'Translation_Start_Site')
missense=c('In_Frame_Del',
           'In_Frame_Ins',
           'Missense_Mutation')
rccGenotypeGenes = c('TP53','BAP1','CDKN2A','VHL','PBRM1','SETD2','KDM5C','PTEN','MTOR','PIK3CA','MET','FAT1','NF2','KDM6A','SMARCB1','NFE2L2','STAG2')
for(tgt in rccGenotypeGenes){
  lab = paste0('rccUpdate_',tgt,'.Mutation')
  if(! lab %in% colnames(mDat))
    lab = paste0('rccUpdate_',tgt,'.Mut.')
  tmp = mDat[,lab]
  mDat[,tgt] = ifelse(tmp %in% c(''),
                      'WT',
                      ifelse(tmp %in% c('n/a'),
                             NA,
                             tmp
                             )
                      )
  mDat[,paste0('hitType',tgt)] = ifelse(is.na(mDat[,tgt]),
                                        'NA',
                                        ifelse(mDat[,tgt] %in% doubleHit,
                                               'doubleHit',
                                               ifelse(mDat[,tgt] %in% nonsense,
                                                      'nonsense',
                                                      ifelse(mDat[,tgt] %in% splice,
                                                             'splice',
                                                             ifelse(mDat[,tgt] %in% missense,
                                                                    'missense',
                                                                    ifelse(mDat[,tgt]=='WT',
                                                                           'WT',
                                                                           'other'
                                                                           )
                                                                    )
                                                             )
                                                      )
                                               )
                                        )
}
mDat$tgtFile = file.path('Results/bulkRNAseq/TCGA/',bulkAssayType,paste0(mDat$UniqueSampleID,'.tsv'))
mDat$toKeep = mDat$hasUpdate & mDat$biopsy=='Primary Tumor'
mDat$dataset = 'TCGA'
bulkMani[['TCGA']] = as.data.frame(mDat[,c(cmnCols,setdiff(colnames(mDat),cmnCols))])
######
# MRT
#TARGET MRT and NBL (control)
tgts = list.files(file.path('Results/bulkRNAseq/TARGET-RT',bulkAssayType)) 
tgts = gsub('.tsv$','',tgts)
mDat = read.delim('Data/TARGET_RT_Discovery_ClinicalData_20180107.txt',sep='\t',header=TRUE)
mDat2 = read.delim('Data/TARGET_RT_Validation_ClinicalData_20180107.txt',sep='\t',header=TRUE)
m = match(gsub('.*_(TARGET-[0-9]+-[A-Z]+)-.*','\\1',tgts),mDat$TARGET.USI)
m2 = match(gsub('.*_(TARGET-[0-9]+-[A-Z]+)-.*','\\1',tgts),mDat2$TARGET.USI)
#The 7 samples with no match are cell lines, need to annotate as such
out = list()
for(i in seq_along(tgts)){
  if(!is.na(m[i])){
    out[[tgts[i]]] = mDat[m[i],,drop=FALSE]
  }else{
    if(!is.na(m2[i])){
      out[[tgts[i]]] = mDat2[m2[i],,drop=FALSE]
    }else{
      out[[tgts[i]]] = mDat[NA,,drop=FALSE][1,,drop=FALSE]
    }
  }
}
mDat = do.call(rbind,out)
rownames(mDat) = tgts
mDat$Disease = TGTtumourCodes[gsub('.*_TARGET-([0-9]+)-([A-Z]+)-([0-9]+)(\\.([0-9]+))?([A-Z])(\\.([0-9]+))?-([0-9]+[A-Z])','\\1',tgts)]
mDat$TissueCode = TGTtissueCodes[gsub('.*_TARGET-([0-9]+)-([A-Z]+)-([0-9]+)(\\.([0-9]+))?([A-Z])(\\.([0-9]+))?-([0-9]+[A-Z])','\\3',tgts)]
mDat$CellLineNo = gsub('.*_TARGET-([0-9]+)-([A-Z]+)-([0-9]+)(\\.([0-9]+))?([A-Z])(\\.([0-9]+))?-([0-9]+[A-Z])','\\5',tgts)
mDat$TissuePortion = gsub('.*_TARGET-([0-9]+)-([A-Z]+)-([0-9]+)(\\.([0-9]+))?([A-Z])(\\.([0-9]+))?-([0-9]+[A-Z])','\\6',tgts)
mDat$NucleicAcid = gsub('.*_TARGET-([0-9]+)-([A-Z]+)-([0-9]+)(\\.([0-9]+))?([A-Z])(\\.([0-9]+))?-([0-9]+[A-Z])','\\9',tgts)
mDat$UniqueSampleID = tgts
mDat$source = 'TARGET'
mDat$tissue = 'Kidney'
mDat$disease = 'MRT'
mDat$biopsy = mDat$TissueCode
mDat$subtype = NA
mDat$outcome = mDat$Vital.status
mDat$toKeep = mDat$biopsy=='Primary'
mDat$tgtFile = file.path('Results/bulkRNAseq/TARGET-RT',bulkAssayType,paste0(mDat$UniqueSampleID,'.tsv'))
mDat$dataset = 'MRT'
bulkMani[['MRT']] = mDat[,c(cmnCols,setdiff(colnames(mDat),cmnCols))]
################
# Neuroblastoma
tgts = list.files(file.path('Results/bulkRNAseq/TARGET-NBL',bulkAssayType)) 
tgts = gsub('.tsv$','',tgts)
mDat = read.delim('Data/TARGET_NBL_Discovery_ClinicalData_20170525.txt',sep='\t',header=TRUE)
#Work out what to do with weird ones.  Drop them as they map to different patients.
tgts = grep(',',tgts,invert=TRUE,value=TRUE)
m = match(gsub('.*_(TARGET-[0-9]+-[A-Z]+)-.*','\\1',tgts),mDat$TARGET.USI)
mDat = mDat[m,]
rownames(mDat) = tgts
mDat$TissueCode = TGTtissueCodes[gsub('.*_TARGET-([0-9]+)-([A-Z]+)-([0-9]+)(\\.([0-9]+))?([A-Z])(\\.([0-9]+))?-([0-9]+[A-Z])','\\3',tgts)]
mDat$TissuePortion = gsub('.*_TARGET-([0-9]+)-([A-Z]+)-([0-9]+)(\\.([0-9]+))?([A-Z])(\\.([0-9]+))?-([0-9]+[A-Z])','\\6',tgts)
mDat$NucleicAcid = gsub('.*_TARGET-([0-9]+)-([A-Z]+)-([0-9]+)(\\.([0-9]+))?([A-Z])(\\.([0-9]+))?-([0-9]+[A-Z])','\\9',tgts)
mDat$UniqueSampleID = tgts
mDat$source = 'TARGET'
mDat$tissue = 'Adrenal'
mDat$disease = 'NBL'
mDat$biopsy = mDat$TissueCode
mDat$subtype = mDat$COG.Risk.Group
mDat$toKeep = mDat$biopsy == 'Primary'
mDat$outcome = mDat$Vital.Status
mDat$tgtFile = file.path('Results/bulkRNAseq/TARGET-NBL',bulkAssayType,paste0(mDat$UniqueSampleID,'.tsv'))
mDat$dataset = 'NBL'
bulkMani[['NBL']] = mDat[,c(cmnCols,setdiff(colnames(mDat),cmnCols))]
########
# ATRT
mDat = read.table('Data/bulkRNAseq/ATRTManifest.txt',sep='\t',header=FALSE)
colnames(mDat) = c('Sample','Subtype')
rownames(mDat) = paste0('ATRT_',mDat[,1])
mDat$UniqueSampleID = rownames(mDat)
mDat$source = 'German'
mDat$tissue = 'Brain'
mDat$disease = 'ATRT'
mDat$biopsy = 'Tumour'
colnames(mDat) = gsub('^Subtype$','subtype',colnames(mDat)) 
mDat$outcome = NA
mDat$tgtFile = file.path('Results/bulkRNAseq/ATRT',bulkAssayType,paste0(mDat$UniqueSampleID,'.tsv'))
mDat$toKeep = TRUE
mDat$dataset = 'ATRT'
bulkMani[['ATRT']] = mDat[,c(cmnCols,setdiff(colnames(mDat),cmnCols))]
#################
# Benchmark data
mDat = read.table('Data/benchMetadata.tsv',sep='\t',header=TRUE)
mDat$source = 'immuneSort'
mDat$tissue = 'Blood'
mDat$disease = 'Normal'
mDat$biopsy = 'Normal'
mDat$subtype = gsub(' ','_',mDat$group)
mDat$outcome = NA
mDat$toKeep=TRUE
mDat$UniqueSampleID = paste0('SangerProjectBench_',mDat$SRR)
mDat$tgtFile = file.path('Results/bulkRNAseq/SangerProjectBench',bulkAssayType,paste0(mDat$UniqueSampleID,'.tsv'))
mDat$dataset = 'BenchImmune'
bulkMani[['BenchImmune']] = mDat[,c(cmnCols,setdiff(colnames(mDat),cmnCols))]
######
# ALL
mDat = read.table('Data/TARGET_RNA_Illumina.tsv',sep='\t',header=TRUE)
mDat$UniqueSampleID = paste0('SangerProjectTARGET_ALL_phase1_',mDat$Run)
mDat$tgtFile = file.path('Results/bulkRNAseq/SangerProjectTARGET_ALL_phase1',bulkAssayType,paste0(mDat$UniqueSampleID,'.tsv'))
mDat = mDat[file.exists(mDat$tgtFile),]
tmp = read.table('Data/TARGET_ALL_Phase_I_ClinicalData_20170525.txt',sep='\t',header=TRUE)
m = match(mDat$submitted_subject_id,tmp$TARGET.USI)
mDat$source = 'TARGET'
mDat$tissue = 'Blood'
mDat$disease = 'ALL'
mDat$biopsy = TGTtissueCodes[gsub('TARGET-10-[A-Z0-9]+-([0-9]+).-.*','\\1',mDat$biospecimen_repository_sample_id)]
mDat$subtype = tmp$Cell.of.Origin[m]
mDat$outcome = tmp$Vital.Status[m]
mDat$toKeep = !is.na(m)
mDat$dataset = 'ALL'
bulkMani[['ALL']] = mDat[,c(cmnCols,setdiff(colnames(mDat),cmnCols))]
#######################
# Foetal Kidney normal
mDat = read.table('Results/bulkRNAseq/FoetalMap/Metadata.tsv',sep='\t',header=TRUE)
mDat$dataset = 'FoetalMap'
mDat$source = 'Sanger'
mDat$tissue = 'Kidney'
mDat$disease = 'Normal'
mDat$biopsy = 'Normal'
mDat$subtype = NA
mDat$outcome = NA
mDat$tgtFile = file.path('Results/bulkRNAseq/FoetalMap',bulkAssayType,paste0(mDat$UniqueSampleID,'.tsv'))
mDat$toKeep=TRUE
bulkMani[['FoetalMap']] = mDat[,c(cmnCols,setdiff(colnames(mDat),cmnCols))]
########
# GTeX
rc2 = readRDS('Data/bulkRNAseq/rse_gene_GTeX.RDS')
mDat = colData(rc2)
rownames(mDat) = paste0('GTEX_',gsub('-','.',rownames(mDat)))
mDat = as.data.frame(mDat)
mDat$dataset = 'GTEX'
mDat$source = 'GTEX'
mDat$tissue = mDat$smts
mDat$disease = 'Normal'
mDat$biopsy = 'Normal'
mDat$subtype = mDat$smtsd
mDat$outcome = NA
mDat$UniqueSampleID = rownames(mDat)
mDat$tgtFile = file.path('Results/bulkRNAseq/GTEX',bulkAssayType,paste0(mDat$UniqueSampleID,'.tsv'))
mDat$toKeep = mDat$tissue=='Kidney'
bulkMani[['GTEX']] = mDat
###########
# SickKids
mDat = read.table('Results/bulkRNAseq/SickKids/Metadata.tsv',sep='\t',header=TRUE)
mDat$dataset = 'SickKids'
mDat$source = 'SickKids'
colnames(mDat) = gsub('Tissue','tissue',colnames(mDat))
colnames(mDat) = gsub('Disease','disease',colnames(mDat))
mDat$biopsy = 'Tumour'
mDat$subtype=NA
mDat$outcome = NA
mDat$tgtFile = file.path('Results/bulkRNAseq/SickKids',bulkAssayType,paste0(mDat$UniqueSampleID,'.tsv'))
mDat$toKeep = FALSE
bulkMani[['SickKids']] = mDat
##############
# Misc Sanger
mDat = data.frame(dataset='SangerMisc',
                  source='Sanger',
                  tissue='Kidney',
                  disease='Cryptic',
                  biopsy='Tumour',
                  subtype=NA,
                  outcome=NA,
                  UniqueSampleID='SangerProject1700_PR40309a',
                  toKeep=FALSE)
mDat$tgtFile = file.path('Results/bulkRNAseq/SangerProject1700',bulkAssayType,paste0(mDat$UniqueSampleID,'.tsv'))
bulkMani[['SangerMisc']] = mDat
########
# Merge
tmp = unique(unlist(lapply(bulkMani,colnames)))
bigBulkMani = lapply(bulkMani,function(e) {for(x in tmp){if(!x %in% colnames(e)){e[,x]=NA}};e[,tmp]})
bigBulkMani = do.call(rbind,bigBulkMani)
rownames(bigBulkMani) = bigBulkMani$UniqueSampleID

######################
# Define control set #
######################
controlSet = c()
#TARGET-ALLs, dropping weird ones
controlSet = c(controlSet,bigBulkMani$UniqueSampleID[grep('^B precursor',bigBulkMani$subtype)])
#And the PBMC populations
#B
controlSet = c(controlSet,bigBulkMani$UniqueSampleID[which(bigBulkMani$group5=='B cells')])
#Monocytes
controlSet = c(controlSet,bigBulkMani$UniqueSampleID[which(bigBulkMani$group4=='Monocytes')])
#T CD4
controlSet = c(controlSet,bigBulkMani$UniqueSampleID[which(bigBulkMani$group3=='Tfh+Th')])

#############################
# Common names and ordering #
#############################
#Converts a vector of reference signal (e.g. PBMC_B) to prefered order 
sigOrder = data.frame(dirtyName = c(names(refs$kidPBMCandMast$lrFit),'Intercept'))
#Define grouping
sigOrder$group = gsub('_.*','',sigOrder$dirtyName)
sigOrder$group = c(fKid='devKidney',sciKidRef='matureKidney',PBMC='Leucocytes')[sigOrder$group]
sigOrder$group[grep('MST',sigOrder$dirtyName)]='Leucocytes'
sigOrder$group[grep('Intercept',sigOrder$dirtyName)]='Technical'
sigOrder$group = factor(sigOrder$group,levels=c('devKidney','matureKidney','Leucocytes','Technical'))
#Define names
sigOrder$name = gsub('^[^_]+_','',sigOrder$dirtyName)
#Tidy and fine tune
sigOrder$name = gsub('_cells','',sigOrder$name)
sigOrder$name[grep('MST',sigOrder$name)]='Mast'
sigOrder$name[sigOrder$name=='MSC'] = 'MPC'
sigOrder$name[sigOrder$name=='NPC'] = 'CapMes'
sigOrder$name[sigOrder$name %in% c('AV1','AV2','DV','GE')] = 'End'
sigOrder$name[sigOrder$name=='C1']='IntTypeA'
sigOrder$name[sigOrder$name=='C2']='IntTypeB'
sigOrder$name[sigOrder$name=='D']='Distal'
sigOrder$name[sigOrder$name=='F']='Fibroblast'
sigOrder$name[sigOrder$name=='G']='Glomerulus'
sigOrder$name[sigOrder$name=='H']='LoH'
sigOrder$name[sigOrder$name=='M']='Mesangial'
sigOrder$name[sigOrder$name=='End']='Vascular'
sigOrder$name[sigOrder$name=='Intercept']='Unexplained'
#Order
sigOrder$name = factor(sigOrder$name,levels=c('Vascular','Fibroblast','Mesangial','MPC','ICa','ICb','CapMes','RVCSB','SSBpod','SSBpr','SSBm.d','Pod','ErPrT','PT1','PT3','Glomerulus','DTLH','LoH','Distal','CnT','IntTypeA','IntTypeB','UBCD','B','CD4_T','CD8_T','NK','Dendritic','Monocytes','Mast','Unexplained'))
#Do final ordering
sigOrder = sigOrder[order(sigOrder$group,sigOrder$name),]
rownames(sigOrder) = sigOrder$dirtyName
#Load in the colours
sigOrder$groupCols = c(devKidney=sigCols['Kidney_Foetal','colour'],
                       matureKidney=sigCols['Kidney_Mature','colour'],
                       Leucocytes = sigCols['Leucocytes_Mature','colour'],
                       Technical = sigCols['Intercept_Mature','colour'])[as.character(sigOrder$group)]


##################
# Colour schemes #
##################
colSchemes = function(){
  #Samples
  gg = ggplot(data.frame(x='Type',sampleType=factor(names(sampCols),levels=names(sampCols))),aes(x,sampleType)) + 
    geom_tile(aes(fill=sampleType)) +
    scale_fill_manual(values=sampCols) +
    theme(axis.title.x = element_blank(),
          axis.text.x = element_blank(),
          axis.ticks.x = element_blank())
  plotFun =function(noPlot=FALSE,noFrame=FALSE){
    plot(gg)
  }
  saveFig(file.path(resDir,'colSchemeTumour'),plotFun,height=5)
  #Signals
  for(nom in unique(sigCols$L1)){
    tgts = sigCols[sigCols$L1==nom,]
    tgts = tgts[order(tgts$level),]
    dat = matrix(rownames(tgts),nrow=2,byrow=FALSE)
    colnames(dat) = tgts$name[tgts$development=='Mature']
    rownames(dat) = c('Foetal','Mature')
    dat = t(dat)
    tt = tgts[tgts$development=='Mature',c('L1','L2','level')]
    if(all(is.na(tt$L2)))
      tt = tt[,-which(colnames(tt)=='L2'),drop=FALSE]
    if(all(is.na(tt$level)))
      tt = tt[,-which(colnames(tt)=='level'),drop=FALSE]
    sideAnno = HeatmapAnnotation(df = tt,which='row')
    #Row split by things that are the same
    rSplit = tgts$hue[tgts$development=='Mature']
    rSplit = tgts$name[match(rSplit,tgts$hue)]
    rSplit = factor(rSplit,level=unique(rSplit))
    m = match(rownames(dat),tgts$name)
    hm = Heatmap(dat,
                 col=setNames(tgts$colour,rownames(tgts)),
                 name='Hi',
                 right_annotation=sideAnno,
                 cluster_rows=FALSE,
                 cluster_columns=FALSE,
                 show_heatmap_legend=FALSE,
                 show_column_names=TRUE,
                 #row_order = order(tgts$L1[m],tgts$L2[m],tgts$level[m],tgts$colour[m]),
                 row_split = rSplit,
                 row_title_rot = 0
                 )
    plotFun = function(noFrame=FALSE,noPlot=FALSE){
      draw(hm)
    }
    tmp = sigCols
    tmp$label = rownames(tmp)
    saveFig(file.path(resDir,paste0('colSchemeRef',nom)),plotFun,rawData=tmp,width=7,height=5)
    #pdf(file.path(workDir,writeDir,sprintf('colourDefinitions%s.pdf',nom)))
    #print(hm)
    #dev.off()
  }
}


#####################################
# Schematic reference and benchmark #
#####################################

###########################
# Foetal kidney reference #
###########################
fKidPlot = function(){
  dd = cbind(fKid@meta.data,fKid@reductions$umap@cell.embeddings)
  plotFun = function(noFrame=FALSE,noPlot=FALSE){
    par(mar=c(2,2.5,1,0))
    #Construct colours
    ccs = suppressWarnings(brewer.pal(100,'Set3'))[-c(2,10)]
    ccs = c(NPC=ccs[3],
            RVCSB = ccs[5],
            SSBpod = ccs[10],
            SSBpr = '#ccb400',#ccs[10],
            SSBm.d = '#ffe733',#ccs[10],
            Pod = ccs[4],
            ErPrT = ccs[2],
            DTLH = ccs[1],#Purely custom, to lose the pink ccs[7],
            CnT = '#328173',
            UBCD = '#235c53',#ccs[1],
            End = '#c10b69',#ccs[8],
            ICa = ccs[9],
            ICb = ccs[6],
            MSC = '#69931f'#This is a tweaked version of ccs[6]
            )
    #Alternate
    ccs = c(NPC= sigCols['CapMesenchyme_Foetal','colour'],
            RVCSB = sigCols['PrimitiveVesicle_Foetal','colour'],
            SSBpod = sigCols['EarlyTubule_Foetal','colour'],
            SSBpr = sigCols['EarlyTubule_Foetal','colour'],
            SSBm.d = sigCols['EarlyTubule_Foetal','colour'],
            Pod = sigCols['Vesicle_Foetal','colour'],
            ErPrT = sigCols['Tubules_Foetal','colour'],
            DTLH = sigCols['DistalTubules_Foetal','colour'],
            CnT = sigCols['IntercalatedTypeA_Foetal','colour'],
            UBCD = sigCols['UretericBud_Foetal','colour'],
            End = sigCols['Vascular_Foetal','colour'],
            ICa = sigCols['ICa_Foetal','colour'],
            ICb = sigCols['ICb_Foetal','colour'],
            MSC = sigCols['MSC_Foetal','colour']
            )
    plot(dd$UMAP_1,dd$UMAP_2,
         las=1,
         type='n',
         xlab=ifelse(noFrame,'','UMAP1'),
         ylab=ifelse(noFrame,'','UMAP2'),
         main=ifelse(noFrame,'','Foetal Kidney'),
         xaxt=ifelse(noFrame,'n','s'),
         yaxt=ifelse(noFrame,'n','s'),
         frame.plot=FALSE)
    if(!noPlot){
      #Add density contours
      pl$addDensityContours(dd$UMAP_1,dd$UMAP_2,dd$annot,col=pl$colAlpha('black',0.4))
      points(dd$UMAP_1,dd$UMAP_2,
             col = ccs[dd$annot],
             pch = 19,
             cex=0.01)
    }
    if(!noFrame){
      #Add coloured labels
      mids = aggregate(cbind(UMAP_1,UMAP_2) ~ annot,data=dd,FUN=mean)
      #Position tweaks
      mids[mids$annot=='RVCSB','UMAP_2'] = mids[mids$annot=='RVCSB','UMAP_2'] - 1
      mids[mids$annot=='RVCSB','UMAP_1'] = mids[mids$annot=='RVCSB','UMAP_1'] + 2
      mids[mids$annot=='SSBpod','UMAP_2'] = mids[mids$annot=='SSBpod','UMAP_2'] + 2
      mids[mids$annot=='SSBpr','UMAP_2'] = mids[mids$annot=='SSBpr','UMAP_2'] + 1
      mids[mids$annot=='SSBm.d','UMAP_2'] = mids[mids$annot=='SSBm.d','UMAP_2'] - 1
      mids[mids$annot=='DTLH','UMAP_1'] = mids[mids$annot=='DTLH','UMAP_1'] - 1
      mids[mids$annot=='DTLH','UMAP_2'] = mids[mids$annot=='DTLH','UMAP_2'] + 1
      mids[mids$annot=='CnT','UMAP_1'] = mids[mids$annot=='CnT','UMAP_1'] + 1
      mids[mids$annot=='CnT','UMAP_2'] = mids[mids$annot=='CnT','UMAP_2'] - 1
      boxed.labels(mids$UMAP_1,mids$UMAP_2,
                   labels=mids$annot,
                   bg=ccs[mids$annot],
                   col='black')
    }
  }
  dd$rowName = rownames(dd)
  saveFig(file.path(resDir,'fKidRef'),plotFun,rawData=dd)
  plotFun = function(noFrame=FALSE,noPlot=FALSE){
    #Bar plot of age breakdown
    fKid@meta.data$ageGroup = cut(fKid@meta.data$age,breaks = c(0,12,15,20))
    x = table(fKid@meta.data$ageGroup,fKid@meta.data$annot)
    x = x[,c('NPC','RVCSB','SSBpod','SSBpr','SSBm.d','Pod','ErPrT','DTLH','CnT','UBCD','End','MSC','ICa','ICb')]
    barplot(t(t(x)/colSums(x)),las=2)
  }
  saveFig(file.path(resDir,'fKidRefAge'),plotFun,rawData=dd)
  #Add other plots for supplements
  dd = cbind(fKid@meta.data,fKid@reductions$umap@cell.embeddings)
  cnts = dd$fullAnnot
  cnts[is.na(cnts)]='NA'
  cnts = table(cnts,dd$seurat_clusters)
  x = round(t(t(cnts)/colSums(cnts))*100)
  class(x) = 'matrix'
  x = x[c('NPCa','NPCb','NPCc','NPCd','RVCSBa','RVCSBb','PTA','SSBpod','SSBpr','SSBm.d','Pod','ErPrT','DTLH','CnT','UBCD','ICa','ICb','IPC','End','Mes','Leu','NA'),as.character(c(0,5,11,10,15,7,9,2,4,8,13,14,17,18,16,12,1,3,6,19,20))]
  #Make nice heatmap
  cSplit = ifelse(colnames(x) %in% c(19,20),
                  'Other',
                  ifelse(colnames(x) %in% c(12,1,3,6),
                         'Interstitial',
                         'Nephron'
                         )
                  )
  cSplit = factor(cSplit,levels=c('Nephron','Interstitial','Other'))
  hh = Heatmap(x,
               col=circlize::colorRamp2(seq(0,100,length.out=9),brewer.pal(9,'OrRd')),
               name = 'Percent matching',
               column_split = cSplit,
               cluster_rows = FALSE,
               cluster_column_slices=FALSE,
               cluster_columns = FALSE,
               row_title = 'PLoS reference classes',
               column_gap = unit(5,'mm')
               )
  plotFun = function(noFrame=FALSE,noPlot=FALSE){
    draw(hh,column_title='Seurat clusters',column_title_side='bottom')
  }
  tt = as.data.frame(x)
  tt$rowName = rownames(tt)
  saveFig(file.path(resDir,'fKidRefClasses'),plotFun,width=8,height=5,rawData=tt)
  #The dumb map
  plotFun = function(noFrame=FALSE,noPlot=FALSE){
    plot(DimPlot(fKid,group.by='seurat_clusters',label=TRUE))
  }
  saveFig(file.path(resDir,'fKidRefClusters'),plotFun,width=5,height=5,rawData=dd)
  #Expression of top markers for each annotation group
  annot = fKid@meta.data$annot
  annot[annot=='NPC']='CapMes'
  annot[annot=='MSC']='MPC'
  annot = factor(annot,levels=c('CapMes','RVCSB','SSBpod','Pod','SSBpr','ErPrT','SSBm.d','DTLH','CnT','UBCD','MPC','ICa','ICb','End'))
  hh = annHeatmap(fKid,annot)
  plotFun = function(noFrame=FALSE,noPlot=FALSE){
    draw(hh$hh,column_title='Single cell clusters',row_title='Marker genes')
  }
  tt = as.data.frame(hh$rawData)
  tt$rowName=rownames(tt)
  saveFig(file.path(resDir,'fKidRefGenes'),plotFun,width=7,height=9,rawData=tt)
  #Comparing MPCs to different references
  tgtDat = fKid@assays$RNA@counts[,fKid@meta.data$annot=='MSC']
  rownames(tgtDat)= cmn@misc$geneMapCommon[rownames(tgtDat),'V1']
  tmp = rep('MPC',ncol(tgtDat))
  outKid = lr$predictSimilarity(refs$fKidPBMC$lrFit,tgtDat,classes=tmp)
  outAd = lr$predictSimilarity(refs$fAdPBMC$lrFit,tgtDat,classes=tmp)
  outMouse = lr$predictSimilarity(refs$mouseDevPBMC$lrFit,tgtDat,classes=tmp)
  dd = data.frame(refClass = c(colnames(outKid),colnames(outAd),colnames(outMouse)),
                  refType = rep(c('Kidney','Adrenal','mouseEmbryo'),c(ncol(outKid),ncol(outAd),ncol(outMouse))),
                  value = c(unlist(outKid),unlist(outAd),unlist(outMouse))
                  )
  #Drop duplicate entries (Leucocytes)
  dd = dd[!duplicated(dd$refClass),]
  dd$group = gsub('_.*','',dd$refClass)
  dd$group = factor(dd$group,levels=c('PBMC','fKid','fAd','mouseDev'))
  dd = dd[order(dd$group),]
  plotFun = function(noFrame=FALSE,noPlot=FALSE){
    plot(0,
         type='n',
         xlim=c(0.5,nrow(dd)+0.5),
         ylim=c(-4,4),
         xaxt='n',
         main='MPC compared to other references.',
         xlab='',
         ylab='Similarity score (logit)',
         frame.plot=FALSE,
         las=1
         )
    #Axis line
    abline(h=0)
    for(i in seq(nrow(dd))){
      lines(x=c(i,i),
            y=c(0,dd$value[i]),
            lty=3,
            lwd=0.5,
            xpd=NA
      )
    }
    #The points
    cc = colSet(dd$group,'Greys')
    points(seq(nrow(dd)),dd$value,
           col=cc[as.character(dd$group)],
           pch=19,
           cex=3*((1+exp(-dd$value))**-1),
           xpd=NA
    )
 }
 saveFig(file.path(resDir,'MPCsimilarity'),plotFun,width=7,height=5,rawData=dd)
}


#############
# Schematic #
#############
schematic = function(){
  tgtDat = cmn@assays$RNA@counts
  rownames(tgtDat)= cmn@misc$geneMapCommon[rownames(tgtDat),'V1']
  out = lr$predictSimilarity(refs$fKidPBMC$lrFit,tgtDat)
  #Collapse to reduced class
  out2 = lr$summariseToClass(out,cmn@meta.data$annot)
  colnames(out2) = gsub('(PBMC|fKid)_','',colnames(out2))
  colnames(out2) = gsub('_cells','',colnames(out2))
  #Keep just the positive slice
  out3 = out2[c('B','T','Monocytes','NK','End','Tumour'),c('End','MSC','Monocytes','B','CD4_T')]
  #Convert to greyscale
  out3 = 1/(1+exp(-out3))
  plotFun = function(noFrame=FALSE,noPlot=FALSE){
    hh = lr$similarityHeatmap(out3,
                         show_row_dend=FALSE,
                         show_row_names=TRUE,
                         row_title_rot=0,
                         column_split = ifelse(colnames(out3) %in% c('End','MSC'),'fKid','PBMC'),
                         column_order = colnames(out3),
                         cluster_columns=FALSE
            )
    draw(hh)
  }
  saveFig(file.path(resDir,'lrSchematic'),plotFun)
}

#############
# Benchmark #
#############
bench = function(skipFit){
  tgtSamps = unique(c(controlSet,bigBulkMani$UniqueSampleID[bigBulkMani$dataset %in% c('BenchImmune','ALL')]))
  sigSet = 'benchSet'
  #Make a sample lookup file. 
  srcFile = tempfile()
  mDat = bigBulkMani[bigBulkMani$UniqueSampleID %in% tgtSamps,]
  write.table(normalizePath(mDat$tgtFile),srcFile,row.names=FALSE,col.names=FALSE,quote=FALSE)
  #Load it
  bulkData = loadBulk(mDat$tgtFile)
  ############
  # Our model
  bulkFitCS = runAndLoad(srcFile,refs[[sigSet]]$clustSumFile,file.path(outDir,'immuneBench'),weights=weightVec)
  #########
  # BSeqSC
  dat = bulkData$cnts
  badGenes = getMITRIB()
  gns = gnsMap[!gnsMap[,2] %in% badGenes,]
  gns = gns[gns[,1] %in% rownames(dat),]
  fDat = gns
  rownames(fDat) = fDat[,1]
  colnames(fDat) = c('ENSEMBL','Symbol')
  fDat = AnnotatedDataFrame(fDat)
  pDat = mDat
  rownames(pDat) = pDat$UniqueSampleID
  pDat = AnnotatedDataFrame(pDat)
  eset = as.matrix(dat[gns[,1],])
  eset = ExpressionSet(eset,phenoData=pDat,featureData=fDat)
  #Get markers from single cell data
  tmpA = pbmc@assays$RNA@counts
  tmpB = sciKidRef@assays$RNA@counts[,sciKidRef@meta.data$annot=='PT1']
  classes = c(paste0('PBMC_',pbmc@meta.data$annot),rep('sciKidRef_PT1',ncol(tmpB)))
  tmp = intersect(rownames(tmpA),rownames(tmpB))
  scDat = cbind(tmpA[tmp,],tmpB[tmp,])
  marks = SoupX::quickMarkers(scDat,classes,N=100,expressCut=0.9)
  marks = marks[marks$geneFrequency>0.4 & marks$geneFrequencyOutsideCluster<0.05,]
  marks = split(marks$gene,marks$cluster)
  #Create expression set from single cell
  pDat = data.frame(row.names=colnames(scDat),
                    sampleName = ifelse(grepl('PBMC',colnames(scDat)),'PBMC',gsub('_.*','',colnames(scDat))),
                    cellTypeID = as.numeric(factor(classes)),
                    cellType = classes
                    )
  pDat$sampleID = as.numeric(factor(pDat$sampleName))
  scDat = ExpressionSet(as.matrix(scDat),AnnotatedDataFrame(pDat))
  #Create the silly "Basis" thing
  B = bseqsc_basis(scDat,marks,clusters='cellType',samples='sampleID',ct.scale=TRUE)
  #Now we need cibersort installed
  bseqsc_config('Code/CIBERSORT.R')
  #Actually run the thing
  bsFit = bseqsc_proportions(eset,B,verbose=TRUE)
  tmp = as.data.frame(t(bsFit$stats[,2,drop=FALSE]))
  rownames(tmp)='pR2'
  bulkFitBS = list(extras=tmp,
                   fit=as.data.frame(bsFit$coefficients),
                   norm=as.data.frame(bsFit$coefficients))
  ########
  # MuSiC
  phenoData(scDat)$sampleID = sample(1:2,ncol(scDat),replace=TRUE)
  musFit = music_prop(bulk.eset = eset,sc.eset = scDat,clusters = 'cellType',samples='sampleID',verbose=TRUE)
  bulkFitMS = list(extras=as.data.frame(matrix(musFit$r.squared.full,byrow=FALSE,nrow=1,dimnames=list('pR2',names(musFit$r.squared.full)))),
                   fit=as.data.frame(t(musFit$Est.prop.weighted)),
                   norm=as.data.frame(t(musFit$Est.prop.weighted))
                   )
  ###############
  # Combine them
  #Create merge of everything
  fits = list(CS = bulkFitCS,
              BS = bulkFitBS,
              MS = bulkFitMS)
  cSigs = sort(unique(unlist(lapply(fits,function(e) rownames(e$norm)))))
  cSigs = cSigs[c(which(cSigs!='Intercept'),which(cSigs=='Intercept'))]
  #Exclude non-core if any
  norm = do.call(cbind,lapply(fits,function(e) e$norm[cSigs,mDat$UniqueSampleID]))
  rownames(norm) = cSigs
  #Sort by method, then cell type
  mDat$boxSplit = ifelse(is.na(mDat$group5),
                         'ALL',
                         mDat$group5)
  norm = norm[,order(gsub('\\..*','',colnames(norm)),mDat[gsub('.*\\.','',colnames(norm)),'boxSplit'])]
  norm = as.matrix(norm)
  mm = mDat[gsub('.*\\.','',colnames(norm)),]
  rownames(mm) = colnames(norm)
  mm$method = gsub('\\..*','',rownames(mm))
  mm = mm[,c('method','boxSplit')]
  dd = as.data.frame(t(norm))
  #Grouping mapper
  groups = c(PBMC_B_cells = 'B cells',
             sciKidRef_PT1 = 'Renal PT1',
             Intercept = 'Unexplained',
             PBMC_Monocytes = 'Other leucocytes',
             PBMC_Megakaryocytes = 'Other leucocytes',
             PBMC_Dendritic_cells = 'Other leucocytes',
             PBMC_NK_cells = 'Other leucocytes',
             PBMC_CD4_T_cells = 'Other leucocytes',
             PBMC_CD8_T_cells = 'Other leucocytes'
             )
  dd = do.call(cbind,lapply(split(colnames(dd),groups[colnames(dd)]),function(e) rowSums(dd[,e,drop=FALSE],na.rm=TRUE)))
  dd = as.data.frame(dd)
  #Simplify the groupings
  dd = cbind(method = mm[rownames(dd),'method'],
             cellType = mm[rownames(dd),'boxSplit'],
             sampleID = gsub('.*\\.','',rownames(dd)),
             dd)
  #Decide which subset of data to keep
  #Keep just the B precursor ALL samples
  dd = dd[(!is.na(mDat[dd$sampleID,'subtype']) & mDat[dd$sampleID,'subtype']=="B precursor (Non-T, Non-B ALL)") | dd$cellType %in% c('B cells','Progenitors'),]
  ##################
  # Plot the result
  #Define the colours
  benchCols = c('B cells'="#9BB4DD",
                'Renal PT1'='#24B24B',
                'Other leucocytes' = '#4963A2',
                Unexplained="#BCBEC0")
  #Define the layout
  plotFun = function(noFrame=FALSE,noPlot=FALSE){
    layout(matrix(c(1,2,3),ncol=3),
           widths=table(dd$cellType)[c('B cells','Progenitors','ALL')]/nrow(dd)
    )
    par(mar=c(2,2,1,1))
    #B cells
    tmp = dd[dd$cellType=='B cells',-c(2,3)]
    barplot(t(as.matrix(tmp[,-1])),
            col = benchCols[colnames(tmp)[-1]],
            main = 'Mature normal B cells',
            names.arg = tmp$method,
            space = c(0,diff(as.numeric(factor(gsub('\\..*','',rownames(tmp)))))),
            axes = FALSE,
            las = 2,
            cex.names = 0.5,
            width = 3,
            )
    axis(side=2,
         line=-.9,
         at=seq(0,1,.2),
         labels=c(0,'','','','',1),
         las=2
         )
    axis(side=4,line=-.9,labels=FALSE)
    #Progenitors
    par(mar=c(2,1,1,1))
    tmp = dd[dd$cellType=='Progenitors',-c(2,3)]
    barplot(t(as.matrix(tmp[,-1])),
            col = benchCols[colnames(tmp)[-1]],
            main = 'Progenitors',
            names.arg = tmp$method,
            space = c(0,diff(as.numeric(factor(gsub('\\..*','',rownames(tmp)))))),
            las = 2,
            cex.names = 0.5,
            width = 3,
            axes=FALSE
            )
    axis(side=2,line=0,labels=FALSE)
    axis(side=4,line=0,labels=FALSE)
    #legend (x="topright", legend = colnames(tmp[,-1]), col = benchCols[colnames(tmp)[-1]], pch = 19 ) 
    #ALL
    par(mar=c(2,1,1,1))
    tmp = dd[dd$cellType=='ALL',-c(2,3)]
    barplot(t(as.matrix(tmp[,-1])),
            col = benchCols[colnames(tmp)[-1]],
            main = 'Pre-B cell ALL',
            names.arg = tmp$method,
            space = c(0,diff(as.numeric(factor(gsub('\\..*','',rownames(tmp)))))), #I'm proud of this little bit of magic 
            las = 2,
            axes=FALSE,
            cex.names = 0.5,
            width = 3,
            )
    axis(side=2,line=0,labels=FALSE)
    axis(side=4,line=0,labels=FALSE)
  }
  saveFig(file.path(resDir,'benchCS'),plotFun,width=6.5,height=1.8)
  #legend (x="topright", legend = colnames(tmp[,-1]), col = benchCols[colnames(tmp)[-1]], pch = 19 ) 
  #legend (x="topright", legend = colnames(tmp[,-1]), col = benchCols[colnames(tmp)[-1]], pch = 19 ) 
  #Make nicely formatted table
  dd = as.data.frame(t(norm))
  dd = cbind(method = mm[rownames(dd),'method'],
             cellType = mm[rownames(dd),'boxSplit'],
             sampleID = gsub('.*\\.','',rownames(dd)),
             dd)
  #Decide which subset of data to keep
  #Keep just the B precursor ALL samples
  dd = dd[(!is.na(mDat[dd$sampleID,'subtype']) & mDat[dd$sampleID,'subtype']=="B precursor (Non-T, Non-B ALL)") | dd$cellType %in% c('B cells','Progenitors'),]
  colnames(dd)[1:3] = c('fitMethod','cellType','UniqueSampleID')
  dd$fitMethod = gsub('BS','BSEQ-sc',
                      gsub('CS','CellularSignals',
                           gsub('MS','MuSiC',
                                dd$fitMethod
                                )
                           )
                      )
  dd$cellType = gsub('Progenitors','HematopoeticStemCell',dd$cellType)
  dd$Intercept[dd$fitMethod!='CellularSignals']=NA
  write.table(dd,file.path(resDir,paste0('benchCS_rawData.tsv')),row.names=FALSE,col.names=TRUE,quote=FALSE,sep='\t')
}


########################
# RCC signal benchmark #
########################
benchRCC = function(){ 
  #Define the sample set
  mDat = bigBulkMani[which((bigBulkMani$dataset %in% c('TCGA')) & bigBulkMani$toKeep),]
  mDat = mDat[mDat$biopsy=='Primary Tumor' & mDat$disease=='ccRCC',]
  #Load it
  bulkData = loadBulk(mDat$tgtFile)
  ###############
  # Cell Signals
  srcFile = tempfile()
  write.table(normalizePath(mDat$tgtFile),srcFile,row.names=FALSE,col.names=FALSE,quote=FALSE)
  bulkFitCS = runAndLoad(srcFile,refs$mKidPBMC$clustSumFile,file.path(outDir,'rccBench'),weights=weightVec)
  ##########
  # BSEQsc
  dat = bulkData$cnts
  badGenes = getMITRIB()
  gns = gnsMap[!gnsMap[,2] %in% badGenes,]
  gns = gns[gns[,1] %in% rownames(dat),]
  fDat = gns
  rownames(fDat) = fDat[,1]
  colnames(fDat) = c('ENSEMBL','Symbol')
  fDat = AnnotatedDataFrame(fDat)
  pDat = mDat
  rownames(pDat) = gsub('-','.',pDat$UniqueSampleID)
  pDat = AnnotatedDataFrame(pDat)
  eset = as.matrix(dat[gns[,1],])
  eset = ExpressionSet(eset,phenoData=pDat,featureData=fDat)
  #Get markers from single cell data
  tmpA = pbmc@assays$RNA@counts
  w = sciKidRef@meta.data$annot %in% gsub('sciKidRef_','',mKidAnnots)
  tmpB = sciKidRef@assays$RNA@counts[,w]
  classes = c(paste0('PBMC_',pbmc@meta.data$annot),paste0('sciKidRef_',sciKidRef@meta.data$annot[w]))
  tmp = intersect(rownames(tmpA),rownames(tmpB))
  scDat = cbind(tmpA[tmp,],tmpB[tmp,])
  marks = SoupX::quickMarkers(scDat,classes,N=100,expressCut=0.9)
  marks = marks[marks$geneFrequency>0.4 & marks$geneFrequencyOutsideCluster<0.05,]
  marks = split(marks$gene,marks$cluster)
  #Create expression set from single cell
  pDat = data.frame(row.names=colnames(scDat),
                    sampleName = ifelse(grepl('PBMC',colnames(scDat)),'PBMC',gsub('_.*','',colnames(scDat))),
                    cellTypeID = as.numeric(factor(classes)),
                    cellType = classes
                    )
  pDat$sampleID = as.numeric(factor(pDat$sampleName))
  scDat = ExpressionSet(as.matrix(scDat),AnnotatedDataFrame(pDat))
  #Create the silly "Basis" thing
  B = bseqsc_basis(scDat,marks,clusters='cellType',samples='sampleID',ct.scale=TRUE)
  #Now we need cibersort installed
  bseqsc_config('Code/CIBERSORT.R')
  #Actually run the thing
  bsFit = bseqsc_proportions(eset,B,verbose=TRUE)
  tmp = as.data.frame(t(bsFit$stats[,2,drop=FALSE]))
  rownames(tmp)='pR2'
  bulkFitBS = list(extras=tmp,
                   fit=as.data.frame(bsFit$coefficients),
                   norm=as.data.frame(bsFit$coefficients))
  ########
  # MuSiC
  phenoData(scDat)$sampleID = sample(1:2,ncol(scDat),replace=TRUE)
  musFit = music_prop(bulk.eset = eset,sc.eset = scDat,clusters = 'cellType',samples='sampleID',verbose=TRUE)
  bulkFitMS = list(extras=as.data.frame(matrix(musFit$r.squared.full,byrow=FALSE,nrow=1,dimnames=list('pR2',names(musFit$r.squared.full)))),
                   fit=as.data.frame(t(musFit$Est.prop.weighted)),
                   norm=as.data.frame(t(musFit$Est.prop.weighted))
                   )
  ###################
  # Combine and plot
  dat = bulkFitCS$norm
  colnames(dat) = paste0('CS___',colnames(dat))
  tmp = bulkFitBS$norm[rownames(dat),]
  colnames(tmp) = paste0('BS___',colnames(tmp))
  dat = cbind(dat,tmp)
  tmp = bulkFitMS$norm[rownames(dat),]
  colnames(tmp) = paste0('MS___',colnames(tmp))
  dat = cbind(dat,tmp)
  #Merge scores
  s = rep('Nephron',nrow(dat))
  s[grep('PBMC',rownames(dat))] = 'Leucocytes'
  s[grep('PT1',rownames(dat))]='PT1'
  s[rownames(dat) %in% c('fKid_End','sciKidRef_AV1','sciKidRef_AV2','sciKidRef_DV','sciKidRef_GE')]='Vascular'
  s[rownames(dat) %in% c('sciKidRef_F','sciKidRef_M')]='Fibroblast'
  s[rownames(dat) %in% c('Intercept')]='Unexplained'
  dat = do.call(rbind,lapply(split(rownames(dat),s),function(e) colSums(dat[e,,drop=FALSE])))
  #Define order
  dat = as.data.frame(dat)
  dat = dat[c('Leucocytes','Vascular','Fibroblast','PT1','Nephron','Unexplained'),]
  dat[is.na(dat)]=0
  #Define colours
  cc = setNames(sigCols[paste0(rownames(dat),'_Mature'),'colour'],rownames(dat))
  isControl = c(CS=FALSE,BS=FALSE,MS=FALSE)
  #Make plot
  plotFun = function(noFrame=FALSE,noPlot=FALSE){
    exposurePlot(dat,isControl=isControl,controlColChange=FALSE,ptAlpha=0.4,colPal=cc,labCex=1)
  }
  tt = dat
  tt$rowName = rownames(tt)
  saveFig(file.path(resDir,'BenchRCC'),plotFun,width=12,height=7,rawData=tt)
  #Save the relative abundances
  #write.table(norm,file.path(resDir,sprintf('%s_table.tsv',paste0(figNom,ifelse(nom=='','','_'),nom))),sep='\t',row.names=TRUE,col.names=TRUE,quote=FALSE)
}


#################
# Cryptic bench #
#################
cryptBench = function(){ 
  #Define the sample set
  mDat = bigBulkMani[bigBulkMani$UniqueSampleID %in% c('SickKids_X0043_290136','SangerProject1700_PR40309a'),]
  #Load it
  bulkData = loadBulk(mDat$tgtFile)
  ###############
  # Cell Signals
  srcFile = tempfile()
  write.table(normalizePath(mDat$tgtFile),srcFile,row.names=FALSE,col.names=FALSE,quote=FALSE)
  bulkFitCS = runAndLoad(srcFile,refs$kidPBMC$clustSumFile,file.path(outDir,'cryptBench'),weights=weightVec)
  ##########
  # BSEQsc
  dat = bulkData$cnts
  badGenes = getMITRIB()
  gns = gnsMap[!gnsMap[,2] %in% badGenes,]
  gns = gns[gns[,1] %in% rownames(dat),]
  fDat = gns
  rownames(fDat) = fDat[,1]
  colnames(fDat) = c('ENSEMBL','Symbol')
  fDat = AnnotatedDataFrame(fDat)
  pDat = mDat
  rownames(pDat) = gsub('-','.',pDat$UniqueSampleID)
  pDat = AnnotatedDataFrame(pDat)
  eset = as.matrix(dat[gns[,1],])
  eset = ExpressionSet(eset,phenoData=pDat,featureData=fDat)
  #Get markers from single cell data
  tmpA = pbmc@assays$RNA@counts
  w = sciKidRef@meta.data$annot %in% gsub('sciKidRef_','',mKidAnnots)
  tmpB = sciKidRef@assays$RNA@counts[,w]
  tmpC = fKid@assays$RNA@counts
  rownames(tmpC) = cmn@misc$geneMapCommon[rownames(tmpC),'V1']
  classes = c(paste0('PBMC_',pbmc@meta.data$annot),paste0('sciKidRef_',sciKidRef@meta.data$annot[w]),paste0('fKid_',fKid@meta.data$annot))
  tmp = intersect(intersect(rownames(tmpA),rownames(tmpB)),rownames(tmpC))
  scDat = cbind(tmpA[tmp,],tmpB[tmp,],tmpC[tmp,])
  marks = SoupX::quickMarkers(scDat,classes,N=100,expressCut=0.9)
  marks = marks[marks$geneFrequency>0.4 & marks$geneFrequencyOutsideCluster<0.05,]
  marks = split(marks$gene,marks$cluster)
  #Create expression set from single cell
  pDat = data.frame(row.names=colnames(scDat),
                    sampleName = ifelse(grepl('PBMC',colnames(scDat)),'PBMC',gsub('_.*','',colnames(scDat))),
                    cellTypeID = as.numeric(factor(classes)),
                    cellType = classes
                    )
  pDat$sampleID = as.numeric(factor(pDat$sampleName))
  scDat = ExpressionSet(as.matrix(scDat),AnnotatedDataFrame(pDat))
  #Create the silly "Basis" thing
  B = bseqsc_basis(scDat,marks,clusters='cellType',samples='sampleID',ct.scale=TRUE)
  #Now we need cibersort installed
  bseqsc_config('Code/CIBERSORT.R')
  #Actually run the thing
  bsFit = bseqsc_proportions(eset,B,verbose=TRUE)
  tmp = as.data.frame(t(bsFit$stats[,2,drop=FALSE]))
  rownames(tmp)='pR2'
  bulkFitBS = list(extras=tmp,
                   fit=as.data.frame(bsFit$coefficients),
                   norm=as.data.frame(bsFit$coefficients))
  ########
  # MuSiC
  phenoData(scDat)$sampleID = sample(1:2,ncol(scDat),replace=TRUE)
  musFit = music_prop(bulk.eset = eset,sc.eset = scDat,clusters = 'cellType',samples='sampleID',verbose=TRUE)
  bulkFitMS = list(extras=as.data.frame(matrix(musFit$r.squared.full,byrow=FALSE,nrow=1,dimnames=list('pR2',names(musFit$r.squared.full)))),
                   fit=as.data.frame(t(musFit$Est.prop.weighted)),
                   norm=as.data.frame(t(musFit$Est.prop.weighted))
                   )
  ###################
  # Combine and plot
  dat = bulkFitCS$norm
  colnames(dat) = paste0('CS___',colnames(dat))
  tmp = bulkFitBS$norm[rownames(dat),]
  colnames(tmp) = paste0('BS___',colnames(tmp))
  dat = cbind(dat,tmp)
  tmp = bulkFitMS$norm[rownames(dat),]
  colnames(tmp) = paste0('MS___',colnames(tmp))
  dat = cbind(dat,tmp)
  #Get immaturity score for all methods
  x = colSums(dat[grep('fKid',rownames(dat)),],na.rm=TRUE)
  plotFun = function(noFrame=FALSE,noPlot=FALSE){
    barplot(matrix(x,ncol=2,byrow = TRUE),
            beside = TRUE,
            ylab='Immaturity',
            names.arg = c('Fig6 F-G','Fig6 C-E')
    )
    legend(x='topleft',legend = c('CS','BSeq','MuSiC'),fill = c('black','darkgrey','lightgrey'))
  }
  tt = dat
  tt$rowName = rownames(tt)
  saveFig(file.path(resDir,'BenchCrypt'),plotFun,width=7,height=4,rawData=tt)
  #Save the relative abundances
  #write.table(norm,file.path(resDir,sprintf('%s_table.tsv',paste0(figNom,ifelse(nom=='','','_'),nom))),sep='\t',row.names=TRUE,col.names=TRUE,quote=FALSE)
}


######################
# Immaturity measure #
######################

##############
# Foetalness #
##############
foetalScore = function(){
  #Drop ones that aren't relevant
  mDat = bigBulkMani[!bigBulkMani$dataset %in% c('ALL','ATRT','BenchImmune','NBL') & grepl('Kidney',bigBulkMani$tissue),]
  #Get ones that are labelled as keepers, or are normal.  GTEX samples we care about already pass due to toKeep
  mDat = mDat[mDat$toKeep | (mDat$dataset!='GTEX' & grepl('Normal',mDat$biopsy)),]
  #Drop bilateral wilms and those normal sections with clonal nephrogenesis
  mDat = mDat[-which(mDat$dataset=='WilmsSanger' & (mDat$Bilateral=='Y' | (mDat$Clonal.nephrogenesis=='Yes' & mDat$biopsy=="Normal_tissue"))),]
  #mDat = mDat[!(mDat$dataset=='WilmsSanger' & mDat$biopsy=='Normal_tissue'),]
  #Make age common
  mDat$ageYears = ifelse(!is.na(mDat$age),#TCGA
                         mDat$age/12,
                         ifelse(!is.na(mDat$AgeMonths),#Wilms Sanger
                                 mDat$AgeMonths/12,
                                 ifelse(!is.na(mDat$Age.at.Diagnosis.in.Days),#Wilms TARGET
                                        mDat$Age.at.Diagnosis.in.Days/365,
                                        ifelse(!is.na(mDat$Age.at.Diagnosis.in.days),#CCSK TARGET
                                               mDat$Age.at.Diagnosis.in.days/365,
                                               NA
                                               )
                                        )
                                 )
                         )
  #Make a sample lookup file. 
  srcFile = tempfile()
  write.table(normalizePath(mDat$tgtFile),srcFile,row.names=FALSE,col.names=FALSE,quote=FALSE)
  #Fit the model
  bulkFitCS = runAndLoad(srcFile,refs$kidPBMC$clustSumFile,file.path(outDir,'kidFitAll'),weights=weightVec)
  dat = bulkFitCS$norm
  colnames(dat) = mDat$UniqueSampleID
  #Define the score
  mDat$immScore = colSums(dat[grep('fKid',rownames(dat)),])
  mDat$pR2 = unlist(bulkFitCS$extra['pR2',])
  mDat$intFrac = unlist(dat['Intercept',])
  #Colours
  colBg = "#0000FF70"
  colBgEmph = "#FF000070"
  colPtEmph = "#FF0000FF"
  colPt = "#0000FFFF"
  colShade = '#00009930'
  #################
  # Normal samples
  #Define how we split things
  normSamps = grepl('Normal',mDat$biopsy) 
  normGroups = ifelse(mDat[normSamps,'dataset']=='FoetalMap',
                      'Foetal',
                      'Mature')
                      #ifelse(mDat[normSamps,'dataset']=='WilmsSanger',
                      #       paste0('Wilms',interaction(mDat[normSamps,'Unilateral_or_Bilateral'],mDat[normSamps,'Clonal.nephrogenesis'])),
                      #       'Mature'
                      #       )
                      #)
  dd = mDat[normSamps,]
  dd$split = normGroups
  #Don't include Wilms from TARGET as we don't know which contain clonal nephrogenesis which will make normal not normal
  dd = dd[dd$dataset!='WilmsTARGET',]
  #Define normal range
  normRangeMax = quantile(dd$immScore[dd$dataset!='FoetalMap'],1)
  plotFun = function(noFrame=FALSE,noPlot=FALSE){
    #Set up axis
    plot(0,
         type='n',
         xlim=c(-5,110),
         ylim=c(0,1),
         xaxt='n',
         main='Maturity of normal kidney tissue',
         ylab='Developmental Fraction',
         xlab='Age',
         frame.plot=FALSE,
         las=1
         )
    axis(1,
         col.ticks=1,
         label=c(0,5,16,50,90),
         at = c(0,5,16,50,90)
         )
    axis(1,labels='Fetal',at=-5)
    axis(1,labels='No age',at=105)
    #Add points
    points(dd$ageYears,dd$immScore,
           pch=21,
           bg= colBg,
           col = colPt,
           cex=1.5)
    #Foetal samples
    w = which(dd$dataset=='FoetalMap')
    points(-5+rnorm(length(w),0,1),dd$immScore[w],
           pch=21,
           bg = colBgEmph,
           col = colPtEmph,
           cex=1.5)
    #Undefined samples
    w = which(dd$dataset!='FoetalMap' & is.na(dd$ageYears))
    beeswarm(dd$immScore[w],
             add=TRUE,
             col = colPt,
             cex = 1,
             pch = 19,
             at = 105)
    #Add normal range rectange
    rect(xleft=-5,
         xright=120,
         ybottom=0,
         ytop=normRangeMax,
         col = colShade,
         border = 0)
  }
  #Two versions, to exagerate the 0-5 range in one
  saveFig(file.path(resDir,'immScoreNormalPaed'),plotFun,width=35,height=4)
  tt = dd[,cmnCols]
  tt$rowName=rownames(tt)
  saveFig(file.path(resDir,'immScoreNormal'),plotFun,width=6,height=4,rawData=tt)
  ################
  # Adult tumours
  adultTumSamps = !grepl('Normal',mDat$biopsy) & mDat$source=='TCGA'
  adultTumGroups = mDat[adultTumSamps,'subtype']
  #Rename them and define order
  rccRemap = c('ccRCC'='ccRCC',
               'KIRP CIMP'='pRCC CIMP',
               'PRCC T1'='pRCC T1',
               'PRCC T2'='pRCC T2',
               'PRCC Unc'='pRCC Unc',
               'ChRCC'='chRCC',
               'chRCC MD'='chRCC MD')
  #Define plot order
  adultTumGroups = factor(rccRemap[adultTumGroups],levels=rccRemap)
  dd = mDat[adultTumSamps,]
  dd$split = adultTumGroups
  plotFun = function(noFrame=FALSE,noPlot=FALSE){
    width=30
    plot(0,
         type='n',
         frame.plot=FALSE,
         xlim=c(-width/2,nlevels(dd$split)*width-width/2),
         ylim=c(0,1),
         xaxt='n',
         xlab='Tumour Type',
         ylab = 'Developmental Fraction'
         )
    #Work out where to shove things
    for(i in seq_along(levels(dd$split))){
      nom = levels(dd$split)[i]
      tmp = dd$immScore[dd$split==nom]
      beeswarm(tmp,
               add=TRUE,
               at = (i-1)*width,
               bg=colBg,
               col=colPt,
               pch=21,
               cex=1,
               corral='wrap',
               corralWidth=width*2/3,
               method='hex')
      rect(xleft=(i-1)*width-(width/2*2/3),
           xright=(i-1)*width+(width/2*2/3),
           ybottom=median(tmp)-0.002,
           ytop=median(tmp)+0.002,
           col='white',
           border='black')
      #points((i-1)*width,median(tmp),pch=15,cex=1.5)
      #text((i-1)*width,1,length(tmp))
    }
    #Add normal range rectange
    rect(xleft=-width/2,
         xright=nlevels(dd$split)*width-width/2,
         ybottom=0,
         ytop=normRangeMax,
         col = colShade,
         border = 0)
    #Re-add axis
    axis(1,
         col=0,
         col.ticks=1,
         labels=levels(dd$split),
         at = (seq_along(levels(dd$split))-1)*width,
         las = 2,
         cex.axis=1)
  }
  tt = dd[,cmnCols]
  tt$rowName = rownames(tt)
  saveFig(file.path(resDir,'immScoreAdult'),plotFun,width=6,height=4,rawData=tt)
  ####################
  # Childhood tumours
  childTumSamps = !grepl('Normal',mDat$biopsy) & mDat$source!='TCGA'
  childTumGroups = ifelse(mDat[childTumSamps,'disease']=='Wilms',
                          ifelse(mDat[childTumSamps,'dataset']=='WilmsTARGET',
                                 'WilmsUntreated',
                                 'WilmsTreated'
                                 ),
                          mDat[childTumSamps,'disease']
                          )
  #Rename them and define order
  paedRemap = c('MRT'='MRT',
               'CMN'='CMN',
               'CCSK'='CCSK',
               'WilmsUntreated'='Wilms',
               'WilmsTreated'='Wilms + Chemo'
               )
  #Define plot order
  childTumGroups = factor(paedRemap[childTumGroups],levels=paedRemap)
  dd = mDat[childTumSamps,]
  dd$split = childTumGroups
  plotFun = function(noFrame=FALSE,noPlot=FALSE){
    width=30
    plot(0,
         type='n',
         frame.plot=FALSE,
         xlim=c(-width/2,nlevels(dd$split)*width-width/2),
         ylim=c(0,1),
         xaxt='n',
         xlab='Tumour Type',
         ylab = 'Developmental Fraction'
         )
    #Work out where to shove things
    for(i in seq_along(levels(dd$split))){
      nom = levels(dd$split)[i]
      tmp = dd$immScore[dd$split==nom]
      beeswarm(tmp,
               add=TRUE,
               at = (i-1)*width,
               bg=colBgEmph,
               col=colPtEmph,
               pch=21,
               cex=1,
               corral='wrap',
               corralWidth=width*2/3,
               method='hex')
      rect(xleft=(i-1)*width-(width/2*2/3),
           xright=(i-1)*width+(width/2*2/3),
           ybottom=median(tmp)-0.002,
           ytop=median(tmp)+0.002,
           col='white',
           border='black')
      #points((i-1)*width,median(tmp),pch=15,cex=1.5)
      #text((i-1)*width,1,length(tmp))
    }
    #Add normal range rectange
    rect(xleft=-width/2,
         xright=nlevels(dd$split)*width-width/2,
         ybottom=0,
         ytop=normRangeMax,
         col = colShade,
         border = 0)
    #Re-add axis
    axis(1,
         col=0,
         col.ticks=1,
         labels=levels(dd$split),
         at = (seq_along(levels(dd$split))-1)*width,
         las = 2,
         cex.axis=1)
  }
  tt = dd[,cmnCols]
  tt$rowName = rownames(tt)
  saveFig(file.path(resDir,'immScorePaed'),plotFun,width=4,height=4,rawData=tt)
  ######################
  # Genotype comparison
  #adultTumSamps = !grepl('Normal',mDat$biopsy) & mDat$source=='TCGA'
  adultTumSamps = mDat$source=='TCGA'
  adultTumGroups = ifelse(mDat[adultTumSamps,'biopsy']=='Primary Tumor',mDat[adultTumSamps,'subtype'],'Normal')
  #Rename them and define order
  rccRemap = c(Normal='Normal',
               'ccRCC'='ccRCC',
               'KIRP CIMP'='pRCC CIMP',
               'PRCC T1'='pRCC T1',
               'PRCC T2'='pRCC T2',
               'PRCC Unc'='pRCC Unc',
               'ChRCC'='chRCC',
               'chRCC MD'='chRCC MD')
  #Define plot order
  adultTumGroups = factor(rccRemap[adultTumGroups],levels=rccRemap)
  dd = mDat[adultTumSamps,]
  dd$split = adultTumGroups
  #Get original Nature paper (https://www.nature.com/articles/nature12222) groupings
  natClus = read.table('Data/mRNA_miRNA_clusters.txt',sep='\t',header=TRUE)
  m = match(dd$xml_bcr_patient_barcode,natClus$Patient)
  dd$mRNA = natClus$mRNA_cluster[m]
  dd$mRNA[dd$mRNA=='NA']=NA
  dd$mRNA = factor(dd$mRNA)
  dd$miRNA = natClus$microRNA_cluster[m]
  dd$miRNA[dd$miRNA=='NA']=NA
  dd$miRNA = factor(dd$miRNA)
  #Construct limited covariate table
  tgts = paste0('hitType',rccGenotypeGenes)
  tmp = dd[,c('immScore','split','mRNA','miRNA',tgts)]
  #Add extra covariates
  ##mRNA/miRNA clusters
  #tmp$mRNA2 = dd$rccUpdate_mRNA.Cluster
  #tmp$miRNA2 = dd$rccUpdate_miRNA.Cluster
  #tmp$miRNA2[tmp$miRNA2=='n/a'] = NA
  #tmp$meth = dd$rccUpdate_Methylation.Cluster
  #tmp$meth[tmp$meth=='n/a']=NA
  tmp$grade = dd$rccUpdate_neoplasm_histologic_grade
  tmp$grade[!grepl('^G',tmp$grade)]=NA
  tmp$stage = dd$rccUpdate_pathologic_stage
  tmp$stage[!grepl('Stage',tmp$stage)]=NA
  tmp$pathT = dd$rccUpdate_pathologic_N
  #tmp$sarcomatoid = dd$rccUpdate_presence_of_sarcomatoid_features
  #tmp$sarcomatoid = ifelse(tmp$sarcomatoid %in% c('YES','NO'),tmp$sarcomatoid,NA)
  #Splice mutations are a bit ambiguous, plus there are hardly any, drop them.
  #tmp = tmp[apply(tmp[,tgts],1,function(e) sum(e=='splice'))==0,]
  #Simplify mutational categories
  #tmp$split = factor(as.character(tmp$split))
  for(tgt in tgts){
    x = tmp[,tgt]
    x[x %in% c('missense','nonsense','splice')]='singleHit'
    tmp[,tgt] = factor(x,c('WT','singleHit','doubleHit'))
  }
  ####Drop the NAs
  ####tmp = tmp[apply(tmp[,tgts],1,function(e) sum(e=='NA'))==0,]
  ####Summary tables for each tumour type
  ###for(i in c('ccRCC','pRCC T1','pRCC T2','chRCC')){
  ###  #Check for association with mRNA subtype
  ###  tt = tmp[tmp$split==i,c('mRNA')]
  ###  lmFit
  ###  #Just want to check on the genotype effect
  ###  tt = tmp[tmp$split==i,-which(colnames(tmp)%in% c('split','mRNA2','miRNA2','miRNA','meth')),]
  ###  lmFit = lm(immScore ~ .,data=tt[,which(lengths(apply(tt,2,table))>1)])
  ###  x = split(tt$immScore,factor(tt$miRNA,levels=1:6))
  ###  #boxplot(x,ylim=c(0,1))
  ###  #title(i)
  ###  #rect(xleft=0,
  ###  #     xright=7,
  ###  #     ybottom=0,
  ###  #     ytop=normRangeMax,
  ###  #     col = colShade,
  ###  #     border = 0)
  ###  x = summary(lmFit)
  ###  print(x)
  ###  x = x$coefficients
  ###  x = cbind(covariate = rownames(x),as.data.frame(x))
  ###  write.table(x$coefficients,file.path(resDir,sprintf('lmSummary_%s.tsv',gsub(' ','_',i))),row.names=FALSE,col.names=TRUE,sep='\t',quote=FALSE)
  ###}
  #Save covariate table
  cTab = tmp
  #Model 1 - Mutations, all samples
  fit1 = lm(immScore ~ .,cTab[,-which(colnames(cTab) %in% c('mRNA','miRNA','grade','stage','pathT','sarcomatoid'))])
  tt = summary(fit1)$coefficients
  tt = as.data.frame(tt)
  tt$qVal = p.adjust(tt[,4],method='BH')
  tt$coefficient = rownames(tt)
  tt = tt[,c(6,1:5)]
  write.table(tt,file.path(resDir,'immScore_lm_genotype.tsv'),row.names=FALSE,col.names=TRUE,sep='\t',quote=FALSE)
  #saveFig(file.path(resDir,'immScore_ccRCC_mRNA'),plotFun,width=4,height=4,rawData=tt)
  #Model 2 - Other meta-data, ccRCC spcefic 
  x = cTab[cTab$split=='ccRCC',-which(colnames(cTab) %in% c('split','pathT','sarcomatoid',grep('hitType',colnames(cTab),value=TRUE)))]
  x$mRNA = factor(ifelse(is.na(x$mRNA),'0',as.character(x$mRNA)))
  x$miRNA = factor(ifelse(is.na(x$miRNA),'0',as.character(x$miRNA)))
  fit2 = lm(immScore ~ .,x)
  tt = summary(fit2)$coefficients
  tt = as.data.frame(tt)
  tt$qVal = p.adjust(tt[,4],method='BH')
  tt$coefficient = rownames(tt)
  tt = tt[,c(6,1:5)]
  write.table(tt,file.path(resDir,'immScore_lm_meta.tsv'),row.names=FALSE,col.names=TRUE,sep='\t',quote=FALSE)
  ###################################
  # Make the ccRCC mRNA cluster plot
  #Define plot order
  dd = cTab[cTab$split=='ccRCC',]
  dd$split = (dd$mRNA)
  dd= dd[!is.na(dd$split),]
  plotFun = function(noFrame=FALSE,noPlot=FALSE){
    width=30
    plot(0,
         type='n',
         frame.plot=FALSE,
         xlim=c(-width/2,nlevels(dd$split)*width-width/2),
         ylim=c(0,1),
         xaxt='n',
         xlab='mRNA cluster',
         ylab = 'Developmental Fraction'
         )
    #Work out where to shove things
    for(i in seq_along(levels(dd$split))){
      nom = levels(dd$split)[i]
      tmp = dd$immScore[dd$split==nom]
      beeswarm(tmp,
               add=TRUE,
               at = (i-1)*width,
               bg=colBg,
               col=colPt,
               pch=21,
               cex=1,
               corral='wrap',
               corralWidth=width*2/3,
               method='hex')
      rect(xleft=(i-1)*width-(width/2*2/3),
           xright=(i-1)*width+(width/2*2/3),
           ybottom=median(tmp)-0.002,
           ytop=median(tmp)+0.002,
           col='white',
           border='black')
      #points((i-1)*width,median(tmp),pch=15,cex=1.5)
      #text((i-1)*width,1,length(tmp))
    }
    abline(h=median(cTab$immScore[cTab$split=='ccRCC']),col='red')
    #Add normal range rectange
    rect(xleft=-width/2,
         xright=nlevels(dd$split)*width-width/2,
         ybottom=0,
         ytop=normRangeMax,
         col = colShade,
         border = 0)
    #Re-add axis
    axis(1,
         col=0,
         col.ticks=1,
         labels=levels(dd$split),
         at = (seq_along(levels(dd$split))-1)*width,
         las = 2,
         cex.axis=1)
    title('ccRCC mRNA Clusters')
  }
  tt = dd
  tt$rowName = rownames(tt)
  saveFig(file.path(resDir,'immScore_ccRCC_mRNA'),plotFun,width=4,height=4,rawData=tt)
  #####################
  # Make plot for PTEN
  #Define plot order
  dd = cTab[cTab$split!='Normal',]
  #Reduce to WT, single, double 
  dd$split = (dd$hitTypePTEN)
  dd= dd[!is.na(dd$split),]
  plotFun = function(noFrame=FALSE,noPlot=FALSE){
    width=30
    plot(0,
         type='n',
         frame.plot=FALSE,
         xlim=c(-width/2,nlevels(dd$split)*width-width/2),
         ylim=c(0,1),
         xaxt='n',
         xlab='PTEN status',
         ylab = 'Developmental Fraction'
         )
    #Work out where to shove things
    for(i in seq_along(levels(dd$split))){
      nom = levels(dd$split)[i]
      tmp = dd$immScore[dd$split==nom]
      beeswarm(tmp,
               add=TRUE,
               at = (i-1)*width,
               bg=colBg,
               col=colPt,
               pch=21,
               cex=1,
               corral='wrap',
               corralWidth=width*2/3,
               method='hex')
      rect(xleft=(i-1)*width-(width/2*2/3),
           xright=(i-1)*width+(width/2*2/3),
           ybottom=median(tmp)-0.002,
           ytop=median(tmp)+0.002,
           col='white',
           border='black')
      #points((i-1)*width,median(tmp),pch=15,cex=1.5)
      #text((i-1)*width,1,length(tmp))
    }
    #Add normal range rectange
    rect(xleft=-width/2,
         xright=nlevels(dd$split)*width-width/2,
         ybottom=0,
         ytop=normRangeMax,
         col = colShade,
         border = 0)
    #Re-add axis
    axis(1,
         col=0,
         col.ticks=1,
         labels=levels(dd$split),
         at = (seq_along(levels(dd$split))-1)*width,
         las = 2,
         cex.axis=1)
    title('ccRCC PTEN')
  }
  tt = dd
  tt$rowName = rownames(tt)
  saveFig(file.path(resDir,'immScore_RCC_PTEN'),plotFun,width=3,height=4,rawData=tt)
  #################
  #### KM Survival
  ###mani = mDat
  ###mani$futime = mani$gdc_cases.diagnoses.days_to_death
  ###mani$futime[is.na(mani$futime)] <- mani$gdc_cases.diagnoses.days_to_last_follow_up[is.na(mani$futime)]
  ###mani$fustat <- mani$cgc_case_vital_status %in% "Dead" +0
  ####Drop MD-ChRCCs and PTEN muts
  ###mani = mani[-match(rownames(cTab)[which(cTab$hitTypePTEN=='doubleHit')],rownames(mani)),]
  ###mani = mani[mani$subtype!='chRCC MD',]
  ###mani = mani[mani$disease %in% c('ccRCC','chRCC','pcRCC') & !grepl('Normal',mani$biopsy),]
  ####Define a low immaturity score cut-off
  ###mani$dedif = ifelse(mani$immScore < quantile(mani$immScore,0.25),'low','!low')
  #### Create Kaplen Meier curves
  ###surv_object <- Surv(time = mani$futime, event = mani$fustat)
  ###fit1 <- survfit(surv_object ~ dedif, data = mani)
  ###plotFun = function(noFrame=FALSE,noPlot=FALSE){
  ###  gg = ggsurvplot(fit1, 
  ###                  data = mani, 
  ###                  pval = TRUE,
  ###                  pval.method=TRUE,
  ###                  color='dedif',
  ###                  palette=c('#828282','#232323'))
  ###  plot(gg)
  ###}
  ###saveFig(file.path(resDir,'immScore_KM'),plotFun,width=5,height=4)
}


##############
# CMN Figure #
##############

###############
# CMN signals #
###############
csCMN = function(){
  tgts = which(bigBulkMani$disease %in% c('CMN') & bigBulkMani$toKeep)
  mDat = bigBulkMani[c(which(bigBulkMani$UniqueSampleID %in% c(controlSet)),tgts),]
  #Nice genotype information
  mDat$gtype = ifelse(grepl('BRAF',mDat$subtype),
                      'BRAF',
                      ifelse(grepl('NTRK1',mDat$subtype),
                             'NTRK1',
                             ifelse(grepl('EGFR',mDat$subtype),
                                    'EGFR',
                                    'Other'
                                    )
                             )
                      )
  #Define how to split, group, and order
  mDat$boxSplit = ifelse(mDat$dataset=='BenchImmune',
                         'Immune',
                         mDat$dataset)
  isControl = c(CMN=FALSE,ALL=TRUE,Immune=TRUE)
  #Write reference
  srcFile = tempfile()
  write.table(normalizePath(mDat$tgtFile),srcFile,row.names=FALSE,col.names=FALSE,quote=FALSE)
  #Fit CS
  bulkFitCS = runAndLoad(srcFile,refs$fKidPBMC$clustSumFile,file.path(outDir,'CMN_fKid'),weights=weightVec)
  #Define groupings
  dat = bulkFitCS$norm
  s = rep('Nephron',nrow(dat))
  s[grep('PBMC',rownames(dat))]='Leucocytes'
  s[rownames(dat) %in% c('fKid_End')]='Vascular'
  s[rownames(dat) %in% c('Intercept')]='Unexplained'
  s[rownames(dat) %in% c('fKid_ICa','fKid_ICb')]='Interstitial'
  s[rownames(dat) %in% c('fKid_NPC')]='CapMesenchyme'
  s[rownames(dat) %in% c('fKid_MSC')]='MSC'
  dat = do.call(rbind,lapply(split(rownames(dat),s),function(e) colSums(dat[e,,drop=FALSE])))
  #Define order
  dat = dat[c('Leucocytes','Vascular','Interstitial','MSC','CapMesenchyme','Nephron','Unexplained'),]
  #Add in sample grouping
  colnames(dat) = paste0(mDat$boxSplit,'___',colnames(dat))
  #Define signal colours
  cc = setNames(sigCols[paste0(rownames(dat),'_Foetal'),'colour'],rownames(dat))
  #Make plot
  plotFun = function(noFrame=FALSE,noPlot=FALSE){
    exposurePlot(dat,isControl=isControl,controlColChange=TRUE,ptAlpha=0.4,colPal=cc)
  }
  tt = as.data.frame(dat)
  tt$rowName = rownames(tt)
  saveFig(file.path(resDir,'CMN_fKid'),plotFun,width=8,height=5,rawData=dat)
  #################
  # Heamap version
  ss = !isControl[mDat$boxSplit]
  norm = bulkFitCS$norm[,ss]
  extras = bulkFitCS$extras[,ss]
  o = order(factor(s,levels=rownames(dat)),sigOrder[rownames(norm),'name'])
  norm = norm[o,]
  #Construct the annotation for the signals
  sigAnno = HeatmapAnnotation(signalGroup = s[o],
                              col = list(signalGroup = cc),
                              which='row',
                              show_annotation_name=FALSE
  )
  hh = csHeatmap(norm,extras,
            topAnno = list(genotype = mDat$gtype[ss],
                           col = list(genotype=colSet(mDat$gtype[ss],'Set1'))),
            row_labels = as.character(sigOrder[rownames(norm),'name']),
            column_labels = gsub('.*_','',colnames(norm)),
            show_column_names=TRUE,
            row_split = sigOrder[rownames(norm),'group'],
            right_annotation = sigAnno)
  plotFun = function(noFrame=FALSE,noPlot=FALSE){
    draw(hh,row_title='Reference cell signals')
  }
  dd = as.data.frame(norm)
  dd$pR2 = unlist(extras['pR2',])
  dd$rowName=rownames(dd)
  saveFig(file.path(resDir,'CMN_fKid_heatmap'),plotFun,width=12,height=7,rawData=dd)
}


#############################
# Somatic CMN genes in fKid #
#############################
genesCMN = function(){
  #Aggregate in clusters
  tmp = do.call(cbind,lapply(split(colnames(fKid),fKid@meta.data$annot),function(e) rowMeans(fKid@assays$RNA@data[somaticGenes$CMN,e,drop=FALSE])))
  tmp = tmp[c('NTRK3','EGFR'),]
  #Order and rename columns
  colnames(tmp) = gsub('End','Vascular',colnames(tmp))
  tmp = tmp[,c('ICa','ICb','MSC','Vascular','NPC','RVCSB','SSBpod','Pod','SSBpr','ErPrT','SSBm.d','DTLH','CnT','UBCD')]
  ss = ifelse(colnames(tmp) %in% c('ICa','ICb','MSC'),
              'Interstitial',
              ifelse(colnames(tmp)=='Vascular',
                     'Vascular',
                     'Nephron'
                     )
              )
  ss = factor(ss,levels=c('Interstitial','Vascular','Nephron'))
  #Colours
  col = circlize::colorRamp2(seq(-2,2,length.out=length(divColsAlt)),divColsAlt)
  plotFun=function(noFrame=FALSE,noPlot=FALSE){
    hh = Heatmap(t(scale(t(tmp))),
                 col=col,
                 name='Expression\nz-score',
                 cluster_rows=FALSE,
                 cluster_columns=FALSE,
                 column_split = ss
                 )
    draw(hh)
  }
  tt = as.data.frame(tmp)
  tt$rowName=rownames(tt)
  saveFig(file.path(resDir,'CMN_genes'),plotFun,width=4,height=2,rawData=tt)
}


#######################################
# Mature and developmental comparison #
#######################################
csCMN_devComp = function(){
  tgts = which(bigBulkMani$disease %in% c('CMN') & bigBulkMani$toKeep)
  mDat = bigBulkMani[c(which(bigBulkMani$UniqueSampleID %in% c(controlSet)),tgts),]
  #Define how to split, group, and order
  mDat$boxSplit = ifelse(mDat$dataset=='BenchImmune',
                         'Immune',
                         mDat$dataset)
  isControl = c(CMN=FALSE,ALL=TRUE,Immune=TRUE)
  #Write reference
  srcFile = tempfile()
  write.table(normalizePath(mDat$tgtFile),srcFile,row.names=FALSE,col.names=FALSE,quote=FALSE)
  #Fit CS
  bulkFitCS = runAndLoad(srcFile,refs$kidPBMC$clustSumFile,file.path(outDir,'CMN_fKid_devComp'),weights=weightVec)
  #Define groupings
  dat = bulkFitCS$norm
  dat = dat[,grep('SangerProject1716',colnames(dat))]
  #Get the comparison between F and MSC
  dd = data.frame(MSC = unlist(dat['fKid_MSC',]),
                  'F' = unlist(dat['sciKidRef_F',]),
                  sample = colnames(dat)
                  )
  #Determine colours
  cc = setNames(sigCols[c('MSC_Foetal','Fibroblast_Mature'),'colour'],c('MSC','F'))
  plotFun = function(noFrame=FALSE,noPlot=FALSE){
    par(mar=c(2,2,0,0))
    #Make the plot
    plot(0,
         type='n',
         xlab='',
         ylab='SignalFrac',
         xaxt='n',
         xlim=c(0.5,2.5),
         ylim=c(0,1),
         frame.plot=FALSE,
         )
    axis(1,
         at=c(1,2),
         label=c('MSC','F')
         )
    #Points
    dd$xMSC = 1+rnorm(nrow(dd),0,0.05)
    dd$xF = 2+rnorm(nrow(dd),0,0.05)
    #Lines
    for(i in seq(nrow(dd))){
      lines(c(dd$xMSC[i],dd$xF[i]),
            y=c(dd$MSC[i],dd$F[i]),
            col='grey',
            lwd=0.01)
    }
    points(dd$xMSC,dd$MSC,
           col=cc['MSC'],
           pch=15,
           cex=1)
    points(dd$xF,dd$F,
           col=cc['F'],
           pch=19,
           cex=1)
  }
  tt = as.data.frame(dd)
  tt$rowName=rownames(tt)
  saveFig(file.path(resDir,'CMN_devComp'),plotFun,width=3,height=4,rawData=tt)
}


############
# CMN UMAP #
############
umapCMN = function(){
  #Construct colours
  #ccs = setNames(brewer.pal(3,'Accent'),c('Organoid','Fresh','Nuclear'))
  #Make object
  dd = cbind(cmn@meta.data,cmn@reductions$umap@cell.embeddings,EGFR=cmn@assays$RNA@data['EGFR',],NTRK3=cmn@assays$RNA@data['NTRK3',])
  plotFun = function(noFrame=FALSE,noPlot=FALSE){
    par(mfcol=c(1,2))
    for(tgt in c('NTRK3','EGFR')){
      par(mar=c(2,2.5,0,2.0))
      plot(dd$UMAP_1,dd$UMAP_2,
           las=1,
           type='n',
           xlab=ifelse(noFrame,'','UMAP1'),
           ylab=ifelse(noFrame,'','UMAP2'),
           xaxt=ifelse(noFrame,'n','s'),
           yaxt=ifelse(noFrame,'n','s'),
           frame.plot=FALSE)
      if(!noPlot){
        #Add density contours
        pl$addDensityContours(dd$UMAP_1,dd$UMAP_2,dd$annot,col=pl$colAlpha('black',0.4),nSplits=10)
        points(dd$UMAP_1,dd$UMAP_2,
               col = ifelse(dd[,tgt]>0,pl$colAlpha('darkred',1),pl$colAlpha('black',0.03)),
               pch = 19,
               cex=0.3)
      }
      if(!noFrame){
        #Add coloured labels
        mids = aggregate(cbind(UMAP_1,UMAP_2) ~ annot,data=dd,FUN=mean)
        mids[mids$annot=='Adrenal',c('UMAP_1','UMAP_2')] = c(7,-16)
        mids[mids$annot=='B',c('UMAP_1','UMAP_2')] = c(-12,10)
        mids[mids$annot=='End',c('UMAP_1','UMAP_2')] = c(-5,-8)
        mids[mids$annot=='Monocytes',c('UMAP_1','UMAP_2')] = c(-12,0)
        mids[mids$annot=='NK',c('UMAP_1','UMAP_2')] = c(6,5)
        mids[mids$annot=='T',c('UMAP_1','UMAP_2')] = c(11,-5)
        mids[mids$annot=='Tumour',c('UMAP_1','UMAP_2')] = c(-7,-15)
        boxed.labels(mids$UMAP_1,mids$UMAP_2,
                     labels=mids$annot,
                     #bg=ccs[mids$type],
                     col='black')
      }
    }
  }
  dd$rowName=rownames(dd)
  saveFig(file.path(resDir,'CMN_UMAP'),plotFun,rawData=dd)
  #Expression of top markers for each annotation group
  annot = cmn@meta.data$annot
  annot[annot=='End']='Endo'
  annot = factor(annot,levels=c('Tumour','B','T','NK','Monocytes','Endo','Adrenal'))
  hh = annHeatmap(cmn,annot)
  plotFun = function(noFrame=FALSE,noPlot=FALSE){
    draw(hh$hh,column_title='Single cell clusters',row_title='Marker genes')
  }
  tt = as.data.frame(hh$rawData)
  tt$rowName=rownames(tt)
  saveFig(file.path(resDir,'CMNRefGenes'),plotFun,width=7,height=9,rawData=tt)
}

##########
# CMN LR #
##########
lrCMN = function(){
  tgtDat = cmn@assays$RNA@counts
  rownames(tgtDat)= cmn@misc$geneMapCommon[rownames(tgtDat),'V1']
  out = lr$predictSimilarity(refs$fKidPBMC$lrFit,tgtDat)
  colnames(out) = gsub('(PBMC|fKid)_','',colnames(out))
  colnames(out) = gsub('_cells','',colnames(out))
  #Collapse to class
  out2 = lr$summariseToClass(out,cmn@meta.data$annot)
  colnames(out2) = gsub('(PBMC|fKid)_','',colnames(out2))
  colnames(out2) = gsub('_cells','',colnames(out2))
  #Keep and order a subset
  toKeep = c('Monocytes','CD4_T','B','NK','ICa','ICb','MSC','NPC','End')
  out2=out2[,toKeep]
  rSplit = ifelse(rownames(out2) == 'Adrenal',
                                            'Adrenal',
                                            ifelse(rownames(out2) == 'End',
                                                   'Endothelium',
                                                   ifelse(rownames(out2) == 'Tumour',
                                                          'Tumour',
                                                          'Immune'
                                                          )
                                                   )
                                            )
  rSplit = factor(rSplit,levels=c('Tumour','Adrenal','Endothelium','Immune'))
  cSplit = ifelse(colnames(out2) %in% c('CD4_T','B','NK','Monocytes'),
                                               'Immune',
                                               ifelse(colnames(out2) %in% c('ICa','ICb','MSC','NPC'),
                                                      'Mesenchyme',
                                                      'Vascular'
                                                      )
                                               )
  cSplit = factor(cSplit,levels=c('Immune','Mesenchyme','Vascular'))
  plotFun = function(noFrame=FALSE,noPlot=FALSE){
    hh = lr$similarityHeatmap(out[,toKeep],
                              show_row_dend=FALSE,
                              row_split = factor(cmn@meta.data$annot,c('Tumour','Monocytes','T','B','NK','End','Adrenal')),
                              column_split = cSplit,
                              show_row_names=FALSE,
                              cluster_row_slices=FALSE,
                              cluster_columns=FALSE
                              )
    #hh = lr$similarityHeatmap(out2,
    #                     show_row_dend=FALSE,
    #                     show_row_names=TRUE,
    #                     row_title_rot=0,
    #                     row_split = rSplit,
    #                     column_split = cSplit,
    #                     #column_order = c('NPC','RVCSB','SSBpod','Pod','SSBpr','ErPrT','SSBm.d','DTLH','CnT','UBCD','End','MSC','ICa','ICb','B','CD4_T','CD8_T','NK','Monocytes','Dendritic'),
    #                     cluster_columns=FALSE
    #        )
    draw(hh)
  }
  tt = as.data.frame(out)
  tt$rowName=rownames(tt)
  saveFig(file.path(resDir,'lrCMN_fKid'),plotFun,height=3.5,width=3,rawData=tt)
}

#############
# NTRK Bone #
#############
boneCMN = function(){
  #Load bone data
  srat = readRDS('Data/boneSC.RDS')
  srat = scQC$quickCluster(srat@assays$RNA@counts)
  #Get similarity to MSC
  tgtDat = srat@assays$RNA@counts
  rownames(tgtDat) = gnsMap$V1
  fit = lr$predictSimilarity(refs$fKidPBMC$lrFit,tgtDat)#,classes=as.character(srat@meta.data$seurat_clusters))
  #NTRK expression and highest expressing cluster
  m = match(rownames(srat@meta.data),rownames(fit))
  dd = cbind(srat@meta.data,srat@reductions$umap@cell.embeddings,NTRK=srat@assays$RNA@data['NTRK2',],sim=fit[m,'fKid_MSC'])
  cc = circlize::colorRamp2(c(0,2),c('lightgrey','blue'))
  plotFun = function(noFrame=FALSE,noPlot=FALSE){
    plot(dd$UMAP_1,dd$UMAP_2,
         las=1,
         type='n',
         xlab=ifelse(noFrame,'','UMAP1'),
         ylab=ifelse(noFrame,'','UMAP2'),
         xaxt=ifelse(noFrame,'n','s'),
         yaxt=ifelse(noFrame,'n','s'),
         frame.plot=FALSE)
    if(!noPlot){
      #Add density contours for cluster 6
      tt = dd[dd$seurat_clusters=='6',]
      points(dd$UMAP_1,dd$UMAP_2,
             col = cc(dd$NTRK),
             pch = 19,
             cex=0.1)
      pl$addDensityContours(tt$UMAP_1,tt$UMAP_2,tt$seurat_clusters,col='black',lwd=2)
    }
    if(!noFrame){
      pl$addColorBar('topright',col=cc(seq(0,2,.1)),breaks=seq(0,2,.1))
    }
  }
  saveFig(file.path(resDir,'CMN_fBone_NTRK'),plotFun,height=7,width=7)
  #Similarity score for same.
  cc = circlize::colorRamp2(c(0,1),c('lightgrey','blue'))
  cc = circlize::colorRamp2(seq(-4,4,length.out=6),c('#8c510a','#d8b365','#f6e8c3','#c7eae5','#5ab4ac','#01665e'))
  plotFun = function(noFrame=FALSE,noPlot=FALSE){
    plot(dd$UMAP_1,dd$UMAP_2,
         las=1,
         type='n',
         xlab=ifelse(noFrame,'','UMAP1'),
         ylab=ifelse(noFrame,'','UMAP2'),
         xaxt=ifelse(noFrame,'n','s'),
         yaxt=ifelse(noFrame,'n','s'),
         frame.plot=FALSE)
    if(!noPlot){
      #Add density contours for cluster 6
      tt = dd[dd$seurat_clusters=='6',]
      points(dd$UMAP_1,dd$UMAP_2,
             col = cc(dd$sim),
             pch = 19,
             cex=0.1)
      pl$addDensityContours(tt$UMAP_1,tt$UMAP_2,tt$seurat_clusters,col='black',lwd=2)
    }
    if(!noFrame){
      tmp = seq(-4,4,length.out=10)
      pl$addColorBar('topright',col=cc(tmp),breaks=tmp)
    }
  }
  tt = dd[,c('UMAP_1','UMAP_2','seurat_clusters','NTRK','sim')]
  saveFig(file.path(resDir,'CMN_fBone_sim'),plotFun,height=7,width=7,rawData=tt)
}

#########################
# Wilms and CCSK figure #
#########################

#########
# Wilms #
#########
csWilms = function(){
  mDat = bigBulkMani[c(which(bigBulkMani$UniqueSampleID %in% controlSet),which(bigBulkMani$disease %in% c('Wilms','CCSK') & bigBulkMani$toKeep)),]
  #Drop bilateral
  mDat = mDat[-which(mDat$Bilateral=='Y'),]
  #Define how to split, group, and order
  mDat$boxSplit = ifelse(mDat$dataset=='BenchImmune',
                         'Immune',
                         ifelse(mDat$dataset=='WilmsSanger',
                                'Wilms+Chemo',
                                ifelse(mDat$dataset=='WilmsTARGET',
                                       'Wilms',
                                       ifelse(mDat$dataset=='CCSK',
                                              'CCSK',
                                              'ALL'
                                              )
                                       )
                                )
                         )
  isControl = c(Wilms=FALSE,'Wilms+Chemo'=FALSE,CCSK=FALSE,ALL=TRUE,Immune=TRUE)
  #Write reference
  srcFile = tempfile()
  write.table(normalizePath(mDat$tgtFile),srcFile,row.names=FALSE,col.names=FALSE,quote=FALSE)
  #Fit CS
  bulkFitCS = runAndLoad(srcFile,refs$fKidPBMC$clustSumFile,file.path(outDir,'Wilms_fKid'),weights=weightVec)
  #Define groupings
  dat = bulkFitCS$norm
  #Grouping if fitting with fKidPBMC
  s = rep('Nephron',nrow(dat))
  s[grep('PBMC',rownames(dat))]='Leucocytes'
  s[rownames(dat) %in% c('fKid_End')]='Vascular'
  s[rownames(dat) %in% c('Intercept')]='Unexplained'
  s[rownames(dat) %in% c('fKid_ICa','fKid_ICb','fKid_MSC')]='Interstitial'
  s[rownames(dat) %in% c('fKid_MSC')]='MSC'
  s[rownames(dat) %in% c('fKid_NPC','fKid_RVCSB')]='CapMesenchyme'
  #s[rownames(dat) %in% c('fKid_RVCSB')]='PrimitiveVesicle'
  s[grep('SSB',rownames(dat))]='CapMesenchyme'
  dat = do.call(rbind,lapply(split(rownames(dat),s),function(e) colSums(dat[e,,drop=FALSE])))
  #Define order
  dat = dat[c('Leucocytes','Vascular','Interstitial','MSC','CapMesenchyme','Nephron','Unexplained'),]
  #Add in sample grouping
  colnames(dat) = paste0(mDat$boxSplit,'___',colnames(dat))
  #Define signal colours
  cc = setNames(sigCols[paste0(rownames(dat),'_Foetal'),'colour'],rownames(dat))
  ###Grouping if fitting with kidPBMC
  ##s = rep('DevNephron',nrow(dat))
  ##s[grep('PBMC',rownames(dat))]='Leucocytes'
  ##s[rownames(dat) %in% c('fKid_End','sciKidRef_AV1','sciKidRef_AV2','sciKidRef_DV','sciKidRef_GE')]='Vascular'
  ##s[rownames(dat) %in% paste0('sciKidRef_',c('D','G','H','P','PT1','PT2','PT3','U1','U2','U3','U4','C1','C2'))]='MatureNephron'
  ##s[rownames(dat) %in% c('Intercept')]='Unexplained'
  ##s[rownames(dat) %in% c('fKid_ICa','fKid_ICb','fKid_MSC')]='Mesenchyme'
  ##s[rownames(dat) %in% c('sciKidRef_F','sciKidRef_M')]='Fibroblasts'
  ##s[rownames(dat) %in% c('fKid_NPC','fKid_RVCSB')]='CapMesenchyme'
  ##dat = do.call(rbind,lapply(split(rownames(dat),s),function(e) colSums(dat[e,,drop=FALSE])))
  ###Define order
  ##dat = dat[c('Leucocytes','Vascular','Mesenchyme','Fibroblasts','CapMesenchyme','DevNephron','MatureNephron','Unexplained'),]
  ###Add in sample grouping
  ##colnames(dat) = paste0(mDat$boxSplit,'___',colnames(dat))
  ###Define signal colours
  ##cc = setNames(sigCols[paste0(rownames(dat),'_Foetal'),'colour'],rownames(dat))
  ##cc['Fibroblasts']=sigCols['Fibroblast_Mature','colour']
  ##cc['DevNephron']=sigCols['Nephron_Foetal','colour']
  ##cc['MatureNephron']=sigCols['Nephron_Mature','colour']
  #Make plot
  plotFun = function(noFrame=FALSE,noPlot=FALSE){
    exposurePlot(dat,isControl=isControl,controlColChange=TRUE,ptAlpha=0.4,colPal=cc)
  }
  tt = as.data.frame(dat)
  tt$rowName = rownames(tt)
  saveFig(file.path(resDir,'csWilms_fKid'),plotFun,height=5,width=8,rawData=tt)
  #Create the relapse fisher's exact test table
  dd = mDat
  dd$CM = unlist(dat['CapMesenchyme',])
  #Keep only those that have definitive relapse information
  dd = dd[which(dd$Relapsed %in% c('Y','N')),]
  cnts = table(ifelse(dd$Relapsed=='Y','Relapsed','NoEvent'),ifelse(dd$CM>0.2,'CM+','CM-'))
  class(cnts)='matrix'
  cnts = as.data.frame(cnts)
  cnts = cnts[c('Relapsed','NoEvent'),]
  #################
  # Heamap version
  ss = !isControl[mDat$boxSplit]
  norm = bulkFitCS$norm[,ss]
  extras = bulkFitCS$extras[,ss]
  o = order(factor(s,levels=rownames(dat)),sigOrder[rownames(norm),'name'])
  norm = norm[o,]
  #Construct the annotation for the signals
  sigAnno = HeatmapAnnotation(signalGroup = s[o],
                              col = list(signalGroup = cc),
                              which='row',
                              show_annotation_name=FALSE
  )
  splitter = names(ss)[ss]
  splitter[grep('Chemo',splitter)]='Wilms\n(Chemo)'
  hh = csHeatmap(norm,extras,
            row_labels = as.character(sigOrder[rownames(norm),'name']),
            column_labels = gsub('.*_','',colnames(norm)),
            column_title = NULL,
            row_split = sigOrder[rownames(norm),'group'],
            column_split = factor(splitter,levels=c('Wilms','Wilms\n(Chemo)','CCSK')),
            right_annotation = sigAnno)
  plotFun = function(noFrame=FALSE,noPlot=FALSE){
    draw(hh,row_title='Reference cell signals',column_title='Bulk Transcriptomes')
  }
  tt = as.data.frame(norm)
  tt['pR2',] = unlist(extras['pR2',])
  tt$rowName = rownames(tt)
  saveFig(file.path(resDir,'csWilms_fKid_heatmap'),plotFun,width=12,height=7,rawData=tt)
}


##############
# Wilms UMAP #
##############
umapWilms = function(){
  #Construct colours
  #ccs = setNames(brewer.pal(3,'Accent'),c('Organoid','Fresh','Nuclear'))
  #Make object
  dds = list(wilms=cbind(wilms@meta.data,wilms@reductions$umap@cell.embeddings))
  dds$CCSK = cbind(ccsk@meta.data,ccsk@reductions$umap@cell.embeddings)
  plotFun = function(noFrame=FALSE,noPlot=FALSE){
    par(mfcol=c(1,2))
    ccs = setNames(brewer.pal(3,'Greys'),c('End','Leucocytes','Tumour'))
    for(nom in names(dds)){
      dd = dds[[nom]]
      dd$annot = gsub('Tumour.','Tumour',dd$annot)
      par(mar=c(2,2.5,1,2.0))
      plot(dd$UMAP_1,dd$UMAP_2,
           las=1,
           type='n',
           xlab=ifelse(noFrame,'','UMAP1'),
           ylab=ifelse(noFrame,'','UMAP2'),
           main=ifelse(noFrame,'',nom),
           xaxt=ifelse(noFrame,'n','s'),
           yaxt=ifelse(noFrame,'n','s'),
           frame.plot=FALSE)
      if(!noPlot){
        #Add density contours
        pl$addDensityContours(dd$UMAP_1,dd$UMAP_2,dd$annot,col=pl$colAlpha('black',0.4),nSplits=50,splitsToUse=c(2,3,4))
        points(dd$UMAP_1,dd$UMAP_2,
               col = ccs[dd$annot],
               pch = 19,
               cex=0.3)
      }
      if(!noFrame){
        #Add coloured labels
        mids = aggregate(cbind(UMAP_1,UMAP_2) ~ annot,data=dd,FUN=mean)
        #mids[mids$annot=='TumourA',c('UMAP_1','UMAP_2')] = c(4,-8)
        #mids[mids$annot=='TumourB',c('UMAP_1','UMAP_2')] = c(-4,0)
        #mids[mids$annot=='Leucocytes',c('UMAP_1','UMAP_2')] = c(12,-4)
        #mids[mids$annot=='End',c('UMAP_1','UMAP_2')] = c(0,10)
        boxed.labels(mids$UMAP_1,mids$UMAP_2,
                     labels=mids$annot,
                     #bg=ccs[mids$type],
                     col='black')
      }
    }
  }
  tt = dds[[1]]
  tt$rowName=rownames(tt)
  write.table(tt,file.path(resDir,'Wilms_umap_leftPan_rawData.tsv'),quote=FALSE,sep='\t',row.names=FALSE,col.names=TRUE)
  tt = dds[[2]]
  tt$rowName=rownames(tt)
  write.table(tt,file.path(resDir,'Wilms_umap_rightPan_rawData.tsv'),quote=FALSE,sep='\t',row.names=FALSE,col.names=TRUE)
  saveFig(file.path(resDir,'Wilms_umap'),plotFun,width=4,height=3)
  #Genes to justify annotation.  Two groups, Wilms genes, Leucoycte genes. 
  gns=c('PTPRC','WT1','IGF2','SIX1')
  plotFun = function(noFrame=FALSE,noPlot=FALSE){
    plot(FeaturePlot(wilms,gns))
  }
  dd = as.data.frame(cbind(wilms@reductions$umap@cell.embeddings,t(wilms@assays$RNA@data[gns,])))
  dd$rowName = rownames(dd)
  saveFig(file.path(resDir,'WilmsRefGenes'),plotFun,width=7,height=6,rawData=dd)
  #Phase plot
  plotFun = function(noFrame=FALSE,noPlot=FALSE){
    plot(DimPlot(wilms,group.by='Phase'))
  }
  dd = as.data.frame(cbind(wilms@meta.data,wilms@reductions$umap@cell.embeddings))
  dd$rowName = rownames(dd)
  saveFig(file.path(resDir,'WilmsRefPhase'),plotFun,width=7,height=6,rawData=dd)
  #Same deal for CCSK
  gns = c('PTPRC','PECAM1','BCOR')
  plotFun = function(noFrame=FALSE,noPlot=FALSE){
    plot(FeaturePlot(ccsk,gns))
  }
  dd = as.data.frame(cbind(ccsk@reductions$umap@cell.embeddings,t(ccsk@assays$RNA@data[gns,])))
  dd$rowName = rownames(dd)
  saveFig(file.path(resDir,'CCSKRefGenes'),plotFun,width=7,height=6,rawData=dd)
}

####################
# Wilms similarity #
####################
simWilms = function(){
  tgtDat = wilms@assays$RNA@counts
  gns = wilms@misc$geneMapCommon
  rownames(tgtDat) = gns[match(rownames(tgtDat),gns$V2),'V1']
  out = lr$predictSimilarity(refs$fKidPBMC$lrFit,tgtDat)
  #Add CCSK
  tgtDat = ccsk@assays$RNA@counts
  gns = wilms@misc$geneMapCommon
  rownames(tgtDat) = gns[match(rownames(tgtDat),gns$V2),'V1']
  out2 = lr$predictSimilarity(refs$fKidPBMC$lrFit,tgtDat,minGeneMatch=0)
  #Make a big matrix and annotate it
  mat = rbind(out,out2)
  colnames(mat) = gsub('(PBMC|fKid)_','',colnames(mat))
  colnames(mat) = gsub('_cells','',colnames(mat))
  cSplit = gsub('_.*','',colnames(out))
  rSplit = rep(c('Wilms','CCSK'),c(nrow(out),nrow(out2)))
  annot = gsub('Tumour.','Tumour',c(wilms@meta.data$annot,ccsk@meta.data$annot))
  rSplit = interaction(rSplit,annot,drop=TRUE)
  ccs = setNames(brewer.pal(3,'Greys'),c('End','Leucocytes','Tumour'))
  lAnno = HeatmapAnnotation(annot=annot,col=list(annot=ccs),which='row')
  toKeep = c('NPC','RVCSB','SSBpod','SSBpr','SSBm.d','MSC','ICa','ICb','End','Monocytes')
  plotFun = function(noFrame=FALSE,noPlot=FALSE){
    hh = lr$similarityHeatmap(mat[,toKeep],
                         show_row_dend=FALSE,
                         show_row_names=FALSE,
                         row_title_rot=0,
                         left_annotation = lAnno,
                         row_split= rSplit,
                         column_split = cSplit[match(toKeep,colnames(mat))],
                         #column_order = c('NPC','RVCSB','SSBpod','Pod','SSBpr','ErPrT','SSBm.d','DTLH','CnT','UBCD','End','MSC','ICa','ICb','B','CD4_T','CD8_T','NK','Monocytes','Dendritic'),
                         cluster_columns=FALSE
            )
    draw(hh)
  }
  tt = as.data.frame(mat)
  tt$rowName=tt
  saveFig(file.path(resDir,'lrWilms_fKid'),plotFun,width=5,height=3,rawData=tt)
}


###############################
# All fits for GOF comparison #
###############################
gof = function(){
  #Drop ones that aren't relevant
  mDat = bigBulkMani[!bigBulkMani$dataset %in% c('ALL','ATRT','BenchImmune','NBL') & grepl('Kidney',bigBulkMani$tissue),]
  #Get ones that are labelled as keepers, or are normal.  GTEX samples we care about already pass due to toKeep
  mDat = mDat[mDat$toKeep | (mDat$dataset!='GTEX' & grepl('Normal',mDat$biopsy)),]
  #Drop bilateral wilms and those normal sections with clonal nephrogenesis
  mDat = mDat[-which(mDat$dataset=='WilmsSanger' & (mDat$Bilateral=='Y' | (mDat$Clonal.nephrogenesis=='Yes' & mDat$biopsy=="Normal_tissue"))),]
  #Add the inappropriate samples
  mDat = rbind(mDat,bigBulkMani[bigBulkMani$group4 %in% c('B Naive+Memory','T CD4 Memory'),])
  #######################################################
  # Fit the mature kidney things.  Exclude foetal normal
  tmp = mDat[grepl('Normal',mDat$biopsy) | (mDat$disease %in% c('ccRCC','pcRCC','chRCC','Normal')),]
  tmp = tmp[tmp$dataset!='FoetalMap',]
  tmp = tmp[tmp$dataset!='BenchImmune',]
  srcFile = tempfile()
  write.table(normalizePath(tmp$tgtFile),srcFile,row.names=FALSE,col.names=FALSE,quote=FALSE)
  mKidFit = runAndLoad(srcFile,refs$mKidPBMC$clustSumFile,file.path(outDir,'mKidFitAll'),weights=weightVec)
  ####################################################
  # Fit the foetal things.  Include the foetal normal
  tmp = mDat[(mDat$dataset=='FoetalMap') | (mDat$disease %in% c('CCSK','Wilms','MRT','CMN') & !grepl('Normal',mDat$biopsy)),]
  srcFile = tempfile()
  write.table(normalizePath(tmp$tgtFile),srcFile,row.names=FALSE,col.names=FALSE,quote=FALSE)
  fKidFit = runAndLoad(srcFile,refs$fKidPBMC$clustSumFile,file.path(outDir,'fKidFitAll'),weights=weightVec)
  #####################
  # CCSK Mature Kidney
  tmp = mDat[mDat$dataset=='CCSK',]
  srcFile = tempfile()
  write.table(normalizePath(tmp$tgtFile),srcFile,row.names=FALSE,col.names=FALSE,quote=FALSE)
  ccsk_mKid = runAndLoad(srcFile,refs$mKidPBMC$clustSumFile,file.path(outDir,'mKidCCSK'),weights=weightVec)
  ######################
  # CCSK Foetal Adrenal
  tmp = mDat[mDat$dataset=='CCSK',]
  srcFile = tempfile()
  write.table(normalizePath(tmp$tgtFile),srcFile,row.names=FALSE,col.names=FALSE,quote=FALSE)
  ccsk_fAd = runAndLoad(srcFile,refs$fAdPBMC$clustSumFile,file.path(outDir,'fAdCCSK'),weights=weightVec)
  #############
  # CCSK Mouse
  tmp = mDat[mDat$dataset=='CCSK',]
  srcFile = tempfile()
  write.table(normalizePath(tmp$tgtFile),srcFile,row.names=FALSE,col.names=FALSE,quote=FALSE)
  ccsk_mouseDev = runAndLoad(srcFile,refs$mouseDevPBMC$clustSumFile,file.path(outDir,'mouseDevCCSK'),weights=weightVec)
  ###########
  # CCSK HCL
  tmp = mDat[mDat$dataset=='CCSK',]
  srcFile = tempfile()
  write.table(normalizePath(tmp$tgtFile),srcFile,row.names=FALSE,col.names=FALSE,quote=FALSE)
  ccsk_HCL = runAndLoad(srcFile,refs$mouseDevPBMC$clustSumFile,file.path(outDir,'mouseDevHCL'),weights=weightVec)
  ###########################
  # Inappriopriate reference
  tmp = mDat[mDat$dataset=='BenchImmune',]
  srcFile = tempfile()
  write.table(normalizePath(tmp$tgtFile),srcFile,row.names=FALSE,col.names=FALSE,quote=FALSE)
  inappro = runAndLoad(srcFile,refs$fKidOnly$clustSumFile,file.path(outDir,'inappropriate'),weights=weightVec)
  #####################
  # Merge, split, plot
  dd = data.frame(intercept = c(unlist(mKidFit$norm['Intercept',]),unlist(fKidFit$norm['Intercept',]),unlist(ccsk_mKid$norm['Intercept',]),unlist(ccsk_fAd$norm['Intercept',]),unlist(ccsk_mouseDev$norm['Intercept',]),unlist(ccsk_HCL$norm['Intercept',]),unlist(inappro$norm['Intercept',])),
                  pR2 = c(unlist(mKidFit$extras['pR2',]),unlist(fKidFit$extras['pR2',]),unlist(ccsk_mKid$extras['pR2',]),unlist(ccsk_fAd$extras['pR2',]),unlist(ccsk_mouseDev$extras['pR2',]),unlist(ccsk_HCL$extras['pR2',]),unlist(inappro$extras['pR2',])),
                  samples = c(colnames(mKidFit$norm),colnames(fKidFit$norm),colnames(ccsk_mKid$norm),colnames(ccsk_fAd$norm),colnames(ccsk_mouseDev$norm),colnames(ccsk_HCL$norm),colnames(inappro$norm)),
                  refs = rep(c('mKid','fKid','mKid','fAd','mouseDev','HCL','fKid'),c(ncol(mKidFit$norm),ncol(fKidFit$norm),ncol(ccsk_mKid$norm),ncol(ccsk_fAd$norm),ncol(ccsk_mouseDev$norm),ncol(ccsk_HCL$norm),ncol(inappro$norm)))
                  )
  m = match(gsub('-','.',dd$samples),gsub('-','.',mDat$UniqueSampleID))
  dd = cbind(dd,mDat[m,cmnCols]) 
  dd$split = ifelse(dd$disease %in% c('ccRCC','chRCC','pcRCC'),
                    dd$subtype,
                    dd$disease)
  dd$split[grepl('Normal',dd$biopsy)] = 'Normal'
  w = dd$disease == 'Wilms'
  dd$split[w] = ifelse(dd$dataset[w]=='WilmsSanger','Wilms+Chemo','Wilms')
  w = dd$disease == 'CCSK'
  dd$split[w] = paste0(dd$split[w],'_',dd$refs[w])
  dd$split[dd$dataset=='BenchImmune'] = 'Inappropriate'
  #Set the order of the splits
  dd$split = factor(dd$split,levels=c('Normal','ccRCC','PRCC T1','PRCC T2','KIRP CIMP','PRCC Unc','ChRCC','chRCC MD','CMN','MRT','Wilms','Wilms+Chemo','CCSK_fKid','CCSK_mKid','CCSK_fAd','CCSK_mouseDev','CCSK_HCL','Inappropriate'))
  dd = dd[order(dd$split,dd$intercept),]
  #Plot
  dat = as.matrix(rbind(dd$intercept,1-dd$intercept))
  cnts = table(dd$split)
  spacing = rep(0,nrow(dd))
  spacing[which(!duplicated(dd$split))]=10
  plotFun = function(noFrame=FALSE,noPlot=FALSE){
    mids = barplot(dat,
                   width=(1/cnts)[dd$split],
                   border=NA,
                   space=spacing
                   )
    #Create axis
    mids = split(mids,dd$split)
    for(nom in names(mids)){
      axis(1,
           at=c(min(mids[[nom]]),mean(mids[[nom]]),max(mids[[nom]])),
           labels=c(NA,nom,NA),
           lwd.ticks=NA,
           las=2)
      lines(quantile(mids[[nom]],c(0.1,0.9)),rep(median(dd$intercept[dd$split==nom]),2),col='red',lwd=2)
      lines(rep(mean(mids[[nom]]),2),quantile(dd$intercept[dd$split==nom],c(0.25,0.75)),col='red',lwd=2)
    }
  }
  tt = as.data.frame(dat)
  tt$rowName = rownames(tt)
  saveFig(file.path(resDir,'GOF'),plotFun,height=3.5,rawData=tt)
}

########################
# Wilms and CCSK genes #
########################
genesWilms = function(){
  #Extra GO based CCSK Neural genes
  gnsExtra = c('NPTX2','ECEL1','LHX2','NGFR','NTRK3')
  #Aggregate in clusters
  tmp = do.call(cbind,lapply(split(colnames(fKid),fKid@meta.data$annot),function(e) rowMeans(fKid@assays$RNA@data[c(somaticGenes$Wilms,somaticGenes$CCSK,gnsExtra),e,drop=FALSE])))
  tmp = tmp[c('WT1','SIX1','SIX2','IGF2',gnsExtra),]
  #Order and rename columns
  colnames(tmp) = gsub('End','Vascular',colnames(tmp))
  tmp = tmp[,c('ICa','ICb','MSC','Vascular','NPC','RVCSB','SSBpod','Pod','SSBpr','ErPrT','SSBm.d','DTLH','CnT','UBCD')]
  ss = ifelse(colnames(tmp) %in% c('ICa','ICb','MSC'),
              'Interstitial',
              ifelse(colnames(tmp)=='Vascular',
                     'Vascular',
                     'Nephron'
                     )
              )
  ss = factor(ss,levels=c('Interstitial','Vascular','Nephron'))
  #Colours
  col = circlize::colorRamp2(seq(-2,2,length.out=length(divColsAlt)),divColsAlt)
  plotFun = function(noFrame=FALSE,noPlot=FALSE){
    hh = Heatmap((scale(t(tmp))),
                 col=col,
                 name='Expression\nz-score',
                 cluster_rows=FALSE,
                 cluster_columns=FALSE,
                 row_split = ss,
                 column_split = ifelse(rownames(tmp) %in% gnsExtra,'CCSK','Wilms')
                 )
    draw(hh)
  }
  tt = as.data.frame(tmp)
  tt$rowName = rownames(tt)
  saveFig(file.path(resDir,'wilmsAndCCSK_genes'),plotFun,height=3.5,rawData=tt)
}

#############
# MRT Plots #
#############

#######################
# MRT Signal Analysis #
#######################
csMRT = function(){
  tgts = which(bigBulkMani$disease %in% c('MRT') & bigBulkMani$toKeep)
  mDat = bigBulkMani[c(which(bigBulkMani$UniqueSampleID %in% c(controlSet)),tgts),]
  #Define how to split, group, and order
  mDat$boxSplit = ifelse(mDat$dataset=='BenchImmune',
                         'Immune',
                         mDat$dataset)
  isControl = c(MRT=FALSE,ALL=TRUE,Immune=TRUE)
  #Write reference
  srcFile = tempfile()
  write.table(normalizePath(mDat$tgtFile),srcFile,row.names=FALSE,col.names=FALSE,quote=FALSE)
  #Fit CS
  bulkFitCS = runAndLoad(srcFile,refs$fKidPBMC$clustSumFile,file.path(outDir,'MRT_fKid'),weights=weightVec)
  #Define groupings
  dat = bulkFitCS$norm
  s = rep('Nephron',nrow(dat))
  s[grep('PBMC',rownames(dat))]='Leucocytes'
  s[rownames(dat) %in% c('fKid_End')]='Vascular'
  s[rownames(dat) %in% c('Intercept')]='Unexplained'
  s[rownames(dat) %in% c('fKid_ICa','fKid_ICb')]='Interstitial'
  s[rownames(dat) %in% c('fKid_NPC')]='CapMesenchyme'
  s[rownames(dat) %in% c('fKid_MSC')]='MSC'
  dat = do.call(rbind,lapply(split(rownames(dat),s),function(e) colSums(dat[e,,drop=FALSE])))
  #Define order
  dat = dat[c('Leucocytes','Vascular','Interstitial','MSC','CapMesenchyme','Nephron','Unexplained'),]
  #Add in sample grouping
  colnames(dat) = paste0(mDat$boxSplit,'___',colnames(dat))
  #Define signal colours
  cc = setNames(sigCols[paste0(rownames(dat),'_Foetal'),'colour'],rownames(dat))
  #Make plot
  plotFun = function(noFrame=FALSE,noPlot=FALSE){
    exposurePlot(dat,isControl=isControl,controlColChange=TRUE,ptAlpha=0.4,colPal=cc)
  }
  tt = as.data.frame(dat)
  tt$rowName = rownames(tt)
  saveFig(file.path(resDir,'MRT_fKid'),plotFun,width=6,height=4,rawData=tt)
  #################
  # Heamap version
  ss = !isControl[mDat$boxSplit]
  norm = bulkFitCS$norm[,ss]
  extras = bulkFitCS$extras[,ss]
  o = order(factor(s,levels=rownames(dat)),sigOrder[rownames(norm),'name'])
  norm = norm[o,]
  #Construct the annotation for the signals
  sigAnno = HeatmapAnnotation(signalGroup = s[o],
                              col = list(signalGroup = cc),
                              which='row',
                              show_annotation_name=FALSE
  )
  hh = csHeatmap(norm,extras,
            row_labels = as.character(sigOrder[rownames(norm),'name']),
            column_labels = gsub('.*_','',colnames(norm)),
            row_split = sigOrder[rownames(norm),'group'],
            right_annotation = sigAnno)
  plotFun = function(noFrame=FALSE,noPlot=FALSE){
    draw(hh,row_title='Reference cell signals')
  }
  tt = as.data.frame(norm)
  tt['PR2',] = unlist(extras['pR2',])
  tt$rowName = rownames(tt)
  saveFig(file.path(resDir,'MRT_fKid_heatmap'),plotFun,width=12,height=7,rawData=tt)
}

###############
# MRT sc Plot #
###############
scMRT_plot = function(){
  #Construct colours
  ccs = setNames(brewer.pal(3,'Greys'),c('Organoid','Fresh','Nuclear'))
  #Make object
  dd = cbind(mrt@meta.data,mrt@reductions$umap@cell.embeddings)
  #Reduced annotation
  dd$annot = ifelse(grepl('^T',dd$annot),'Tumour',ifelse(grepl('^L',dd$annot),'Leucocytes','Stroma'))
  plotFun = function(noFrame=FALSE,noPlot=FALSE){
    par(mar=c(2,2.5,1,2.0))
    plot(dd$UMAP_1,dd$UMAP_2,
         las=1,
         type='n',
         xlab=ifelse(noFrame,'','UMAP1'),
         ylab=ifelse(noFrame,'','UMAP2'),
         main=ifelse(noFrame,'','scMRT'),
         xaxt=ifelse(noFrame,'n','s'),
         yaxt=ifelse(noFrame,'n','s'),
         frame.plot=FALSE)
    if(!noPlot){
      #Add density contours
      pl$addDensityContours(dd$UMAP_1,dd$UMAP_2,dd$annot,col=pl$colAlpha('black',0.4),nSplits=100)
      points(dd$UMAP_1,dd$UMAP_2,
             col = ccs[dd$type],
             pch = 19,
             cex=0.01)
    }
    if(!noFrame){
      #Add coloured labels
      mids = aggregate(cbind(UMAP_1,UMAP_2) ~ annot,data=dd,FUN=mean)
      cnts = table(dd$type,dd$annot)
      cnts = setNames(rownames(cnts)[apply(cnts,2,which.max)],colnames(cnts))
      mids$type = cnts[mids$annot]
      mids[mids$annot=='L3','UMAP_2'] = mids[mids$annot=='L3','UMAP_2'] + 2
      boxed.labels(mids$UMAP_1,mids$UMAP_2,
                   labels=mids$annot,
                   bg=ccs[mids$type],
                   col='black')
      legend(x=9.5,
             y=15,
             legend=names(ccs),
             col=ccs,
             pch=19,
             cex=1,
             bty='n',
             title='Assay Type',
             xpd=NA
             )
    }
  }
  tt = as.data.frame(dd)
  tt$rowNames = rownames(tt)
  saveFig(file.path(resDir,'scMRT'),plotFun,width=2.5,rawData=tt)
  #Annotation justification
  gns=c('PTPRC','SMARCB1','SMARCA4','SMARCA2')
  plotFun = function(noFrame=FALSE,noPlot=FALSE){
    plot(FeaturePlot(mrt,gns))
  }
  tt = as.data.frame(cbind(mrt@reductions$umap@cell.embeddings,t(mrt@assays$RNA@data[gns,])))
  tt$rowName = rownames(tt)
  saveFig(file.path(resDir,'MRTRefGenes'),plotFun,width=7,height=6,rawData=tt)
  #TWIST1 expression
  dd = cbind(TWIST1=mrt@assays$RNA@counts['TWIST1',],mrt@reductions$umap@cell.embeddings,mrt@meta.data)
  dd = as.data.frame(dd)
  dd = dd[order(dd$TWIST1),]
  dd$annot = gsub('[0-9]','',dd$annot)
  dd$expr = ifelse(dd$TWIST1>0,'expr','noExpr')
  ccs = c(noExpr='black',expr='red')
  plotFun = function(noFrame=FALSE,noPlot=FALSE){
    plot(dd$UMAP_1,dd$UMAP_2,
         las=1,
         type='n',
         xlab=ifelse(noFrame,'','UMAP1'),
         ylab=ifelse(noFrame,'','UMAP2'),
         xaxt=ifelse(noFrame,'n','s'),
         yaxt=ifelse(noFrame,'n','s'),
         frame.plot=FALSE)
    if(!noPlot){
      #Add density contours
      pl$addDensityContours(dd$UMAP_1,dd$UMAP_2,dd$annot,col=pl$colAlpha('black',0.4),nSplits=100)
      points(dd$UMAP_1,dd$UMAP_2,
             col = ccs[dd$expr],
             pch = 19,
             cex=0.3)
    }
    if(!noFrame){
      ##Add coloured labels
      #mids = aggregate(cbind(UMAP_1,UMAP_2) ~ annot,data=dd,FUN=mean)
      #cnts = table(dd$type,dd$annot)
      #cnts = setNames(rownames(cnts)[apply(cnts,2,which.max)],colnames(cnts))
      #mids$type = cnts[mids$annot]
      #boxed.labels(mids$UMAP_1,mids$UMAP_2,
      #             labels=mids$annot,
      #             bg=ccs[mids$type],
      #             col='black')
      legend(x=9.5,
             y=15,
             legend=names(ccs),
             col=ccs,
             pch=19,
             cex=1,
             bty='n',
             title='TWIST1 status',
             xpd=NA
             )
    }
  }
  tt =as.data.frame(dd)
  tt$rowName = rownames(tt)
  saveFig(file.path(resDir,'MRTscTWIST1'),plotFun,width=7,height=6,rawData=tt)
}

##########
# MRT LR #
##########
scMRT_LR = function(){
  #Convert to ENS matrix
  tgtDat = mrt@assays$RNA@counts
  rownames(tgtDat)= mrtFresh@misc$geneMapCommon[rownames(tgtDat),'V1']
  ccs = setNames(brewer.pal(3,'Greys'),c('Organoid','Fresh','Nuclear'))
  #Decide on major ordering of cells
  mDat = mrt@meta.data
  mDat$annotSub = gsub('[0-9]','',mDat$annot)
  mDat$sorter = as.character(interaction(mDat$annotSub,mDat$type))
  o = order(mDat$sorter)
  tgtDat = tgtDat[,o]
  mDat = mDat[o,]
  #######
  # fKid
  out = lr$predictSimilarity(refs$fKidPBMC$lrFit,tgtDat)
  cSplit = gsub('_.*','',colnames(out))
  out2 = lr$summariseToClass(out,mDat$annot)
  out2 = lr$summariseToClass(out,gsub('[0-9]+','',mDat$annot))
  colnames(out2) = gsub('(PBMC|fKid)_','',colnames(out2))
  colnames(out2) = gsub('_cells','',colnames(out2))
  colnames(out) = gsub('(PBMC|fKid)_','',colnames(out))
  colnames(out) = gsub('_cells','',colnames(out))
  #Drop most non-matching columns to save space
  toKeep = c('NPC','RVCSB','SSBpod','MSC','ICa','ICb','Pod','CD4_T','CD8_T','NK','Monocytes')
  #Drop stroma
  toKeepCell = c('L','T')
  plotFun = function(noFrame=FALSE,noPlot=FALSE){
    m = match(toKeep,colnames(out))
    out = out[,toKeep]
    mm = mDat$annotSub %in% toKeepCell
    out = out[mm,]
    mDat = mDat[mm,]
    o = unlist(lapply(split(seq(nrow(out)),mDat$sorter),function(e) {
                  tmp = out[e,]
                  tmp = hclust(dist(tmp))
                  tmp$labels[tmp$order]}))
    o = match(o,rownames(mDat))
    out = out[o,]
    mDat = mDat[o,]
    #Sort within each grouping
    #Type annotation
    lAnno = HeatmapAnnotation(type=mDat$type,name='AssayType',col=list(type=ccs),which='row')
    hh = lr$similarityHeatmap(out,
                         show_row_dend=FALSE,
                         show_row_names=FALSE,
                         row_title_rot=0,
                         row_split = mDat$annotSub,
                         column_split = cSplit[m],
                         column_order = toKeep,
                         left_annotation=lAnno,
                         cluster_columns=FALSE,
                         cluster_rows=FALSE
            )
    draw(hh)
  }
  tt = as.data.frame(out)
  tt$rowNames = rownames(tt)
  saveFig(file.path(resDir,'lrMRT_fKid'),plotFun,width=3.5,rawData=tt)
  ######
  # fAd
  out = lr$predictSimilarity(refs$fAdPBMC$lrFit,tgtDat)
  cSplit = gsub('_.*','',colnames(out))
  rSplit = gsub('[0-9]','',mrt@meta.data$annot)
  out2 = lr$summariseToClass(out,mrt@meta.data$annot)
  out2 = lr$summariseToClass(out,gsub('[0-9]+','',mrt@meta.data$annot))
  colnames(out2) = gsub('(PBMC|fAd)_','',colnames(out2))
  colnames(out2) = gsub('_cells','',colnames(out2))
  colnames(out) = gsub('(PBMC|fAd)_','',colnames(out))
  colnames(out) = gsub('_cells','',colnames(out))
  #Drop most non-matching columns to save space
  toKeep = c('SCPs','Bridge','Chromaffin','Sympathoblastic','Cortex','Mesenchyme','Endothelium','B','CD4_T','CD8_T','NK','Monocytes','Dendritic')
  #Drop stroma
  toKeepCell = c('L','T')
  plotFun = function(noFrame=FALSE,noPlot=FALSE){
    m = match(toKeep,colnames(out))
    out = out[,toKeep]
    mm = rSplit %in% toKeepCell
    out = out[mm,]
    #Type annotation
    lAnno = HeatmapAnnotation(type=mrt@meta.data$type[mm],name='AssayType',col=list(type=ccs),which='row')
    hh = lr$similarityHeatmap(out,
                         show_row_dend=FALSE,
                         show_row_names=FALSE,
                         row_title_rot=0,
                         row_split = rSplit[mm],
                         column_split = cSplit[m],
                         column_order = toKeep,
                         left_annotation=lAnno,
                         cluster_columns=FALSE
            )
    draw(hh)
  }
  tt = as.data.frame(out)
  tt$rowNames = rownames(tt)
  saveFig(file.path(resDir,'lrMRT_fAd'),plotFun,width=3.5,rawData=tt)
  ###########
  # MouseDev
  out = lr$predictSimilarity(refs$mouseDevPBMC$lrFit,tgtDat)
  cSplit = gsub('_.*','',colnames(out))
  out2 = lr$summariseToClass(out,mDat$annotSub)
  colnames(out2) = gsub('(PBMC|mouseDev)_','',colnames(out2))
  colnames(out2) = gsub('_cells','',colnames(out2))
  colnames(out) = gsub('(PBMC|mouseDev)_','',colnames(out))
  colnames(out) = gsub('_cells','',colnames(out))
  #Drop most non-matching columns to save space
  gastPops = c('Caudal_Mesoderm','Caudal_neurectoderm','Def_endoderm','Intermediate_mesoderm','Mixed_mesoderm','Nascent_mesoderm','Neural_crest','NMP','Paraxial_mesoderm','Parietal_endoderm','Pharyngeal_mesoderm','Rostral_neurectoderm','Somitic_mesoderm','Surface_ectoderm')
  postGastPops = c('Blood_progenitors_1','Blood_progenitors_2','Cardiomyocytes','Endothelium','Erythroid1','Erythroid2','Erythroid3','Forebrain_Midbrain_Hindbrain','Gut','Haematoendothelial_progenitors','Mesenchyme','Notochord','Spinal_cord')
  cSplit = ifelse(cSplit=='PBMC',
                  'PBMC',
                  ifelse(colnames(out) %in% gastPops,
                         'Gast',
                         'PostGast'
                         )
                  )
  #Keep only a handful
  toKeep  = colnames(out)
  toKeep = c('Neural_crest','Paraxial_mesoderm','Parietal_endoderm','Monocytes','Notochord','Erythroid1','Mesenchyme')
  #Drop stroma
  toKeepCell = c('L','T')
  plotFun = function(noFrame=FALSE,noPlot=FALSE){
    m = match(toKeep,colnames(out))
    out = out[,toKeep]
    mm = mDat$annotSub %in% toKeepCell
    out = out[mm,]
    mDat = mDat[mm,]
    o = unlist(lapply(split(seq(nrow(out)),mDat$sorter),function(e) {
                  tmp = out[e,]
                  tmp = hclust(dist(tmp))
                  tmp$labels[tmp$order]}))
    o = match(o,rownames(mDat))
    out = out[o,]
    mDat = mDat[o,]
    #Sort within each grouping
    #Type annotation
    lAnno = HeatmapAnnotation(type=mDat$type,name='AssayType',col=list(type=ccs),which='row')
    hh = lr$similarityHeatmap(out,
                         show_row_dend=FALSE,
                         show_row_names=FALSE,
                         row_title_rot=0,
                         row_split = mDat$annotSub,
                         column_split = cSplit[m],
                         column_order = toKeep,
                         column_label = ifelse(colnames(out)=='Monocytes','Monocytes',substr(colnames(out),1,7)),
                         left_annotation=lAnno,
                         cluster_columns=FALSE,
                         cluster_rows=FALSE
            )
    draw(hh)
  }
  tt = as.data.frame(out)
  tt$rowNames = rownames(tt)
  saveFig(file.path(resDir,'lrMRT_mouseDev'),plotFun,width=5,rawData=tt)
}


#######################################
# MRT Signal analysis against Adrenal #
#######################################
csMRT_fAd = function(){
  tgts = which(bigBulkMani$disease %in% c('MRT','NBL') & bigBulkMani$toKeep)
  mDat = bigBulkMani[c(which(bigBulkMani$UniqueSampleID %in% c(controlSet)),tgts),]
  #Define how to split, group, and order
  mDat$boxSplit = ifelse(mDat$dataset=='BenchImmune',
                         'Immune',
                         mDat$dataset)
  isControl = c(MRT=FALSE,NBL=FALSE,ALL=TRUE,Immune=TRUE)
  #Write reference
  srcFile = tempfile()
  write.table(normalizePath(mDat$tgtFile),srcFile,row.names=FALSE,col.names=FALSE,quote=FALSE)
  #Fit CS
  bulkFitCS = runAndLoad(srcFile,refs$fAdPBMC$clustSumFile,file.path(outDir,'MRT_fAd'),weights=weightVec)
  #Define groupings
  dat = bulkFitCS$norm
  s = rep('Leucocytes',nrow(dat))
  s[rownames(dat) %in% c('fAd_SCPs','fAd_Bridge','fAd_Chromaffin','fAd_Sympathoblastic')] = 'SCPs'
  #s[rownames(dat) %in% c('fAd_SCPs')] = 'SCPs'
  #s[rownames(dat) %in% c('fAd_Bridge')] = 'Bridge'
  #s[rownames(dat) %in% c('fAd_Chromaffin')] = 'Chromaffin'
  #s[rownames(dat) %in% c('fAd_Sympathoblastic')] = 'Sympathoblastic'
  s[rownames(dat) %in% c('fAd_Cortex')] = 'Cortex'
  s[rownames(dat) %in% c('fAd_Endothelium')] = 'Vascular'
  s[rownames(dat) %in% c('fAd_Mesenchyme')] = 'MSC'
  s[rownames(dat) %in% c('Intercept')]='Unexplained'
  dat = do.call(rbind,lapply(split(rownames(dat),s),function(e) colSums(dat[e,,drop=FALSE])))
  #Define order
  dat = dat[c('Leucocytes','Vascular','Cortex','SCPs','MSC','Unexplained'),]
  #Add in sample grouping
  colnames(dat) = paste0(mDat$boxSplit,'___',colnames(dat))
  #Define signal colours
  cc = setNames(sigCols[paste0(rownames(dat),'_Foetal'),'colour'],rownames(dat))
  #Make plot
  plotFun = function(noFrame=FALSE,noPlot=FALSE){
    exposurePlot(dat,isControl=isControl,controlColChange=TRUE,ptAlpha=0.4,colPal=cc)
  }
  tt = as.data.frame(dat)
  tt$rowName = rownames(tt)
  saveFig(file.path(resDir,'MRT_fAd'),plotFun,width=7,height=4,rawData=tt)
}

#######################
# Adult tumour figure #
#######################

################
# RCC big fit  #
################
rccBigFit = function(){
  #RCCs
  mDat = bigBulkMani[which((bigBulkMani$dataset %in% c('TCGA')) & bigBulkMani$toKeep),]
  #Normal tissue
  tmp = bigBulkMani[which(bigBulkMani$dataset %in% c('GTEX','TCGA') & grepl('Normal',bigBulkMani$biopsy) & bigBulkMani$tissue=='Kidney'),]
  mDat = rbind(mDat,tmp)
  #Define split
  mDat$boxSplit = ifelse(grepl('Normal',mDat$biopsy),'Normal',mDat$disease)
  isControl = c(Normal=FALSE,ccRCC=FALSE,pcRCC=FALSE,chRCC=FALSE)
  #Do the fit
  srcFile = tempfile()
  write.table(normalizePath(mDat$tgtFile),srcFile,row.names=FALSE,col.names=FALSE,quote=FALSE)
  fit = runAndLoad(srcFile,refs$mKidPBMC$clustSumFile,file.path(outDir,'rccBigFit'),weights=weightVec)
  #Create plot 
  dat = fit$norm
  s = rep('Nephron',nrow(dat))
  s[grep('PBMC',rownames(dat))]='Leucocytes'
  s[rownames(dat) %in% paste0('sciKidRef_',c('AV1','AV2','DV','GE'))]='Vascular'
  s[rownames(dat) %in% c('sciKidRef_M','sciKidRef_F')] = 'Fibroblast'
  s[rownames(dat) %in% c('Intercept')]='Unexplained'
  #s[rownames(dat) %in% c('sciKidRef_C1')] = 'TypeA'
  #s[rownames(dat) %in% c('sciKidRef_C2')] = 'TypeB'
  dat = do.call(rbind,lapply(split(rownames(dat),s),function(e) colSums(dat[e,,drop=FALSE])))
  #Define order
  dat = dat[c('Leucocytes','Vascular','Fibroblast','Nephron','Unexplained'),]
  #Add in sample grouping
  colnames(dat) = paste0(mDat$boxSplit,'___',colnames(dat))
  #Define signal colours
  cc = setNames(sigCols[paste0(rownames(dat),'_Mature'),'colour'],rownames(dat))
  #Make plot
  plotFun = function(noFrame=FALSE,noPlot=FALSE){
    exposurePlot(dat,isControl=isControl,controlColChange=TRUE,ptAlpha=0.4,colPal=cc)
  }
  tt = as.data.frame(dat)
  tt$rowName = rownames(tt)
  saveFig(file.path(resDir,'allRCC_mKid'),plotFun,height=5,width=8,rawData=tt)
  #################
  # Heamap version
  ss = !isControl[mDat$boxSplit] & mDat$biopsy=='Primary Tumor' 
  norm = fit$norm[,ss]
  extras = fit$extras[,ss]
  o = order(factor(s,levels=rownames(dat)),sigOrder[rownames(norm),'name'])
  norm = norm[o,]
  #Collapse vascular by mapping sum onto first match
  tmp = as.character(sigOrder[rownames(norm),'name'])
  toKeep = !duplicated(tmp)
  norm[match('Vascular',tmp),] = colSums(norm[tmp=='Vascular',])
  norm = norm[toKeep,]
  sAlt = s[o]
  sAlt = sAlt[toKeep]
  #Construct the annotation for the signals
  sigAnno = HeatmapAnnotation(signalGroup = sAlt,
                              col = list(signalGroup = cc),
                              which='row',
                              show_annotation_name=FALSE
  )
  #Subtypes
  sTypes = mDat$subtype[ss]
  sTypes = ifelse(sTypes=='chRCC MD',
                  'Metabolically Divergent',
                  ifelse(grepl('PRCC T.',sTypes),
                         gsub('PRCC T([0-9])','Type\\1',sTypes),
                         ifelse(sTypes=='KIRP CIMP',
                                'CIMP',
                                NA
                                )
                         )
                  )
  hh = csHeatmap(norm,extras,
            row_labels = as.character(sigOrder[rownames(norm),'name']),
            column_labels = gsub('.*_','',colnames(norm)),
            column_title=NULL,
            row_split = sigOrder[rownames(norm),'group'],
            column_split = factor(mDat$disease[ss],levels=c('ccRCC','pcRCC','chRCC')),
            topAnno = list(subtype = sTypes,
                           col=list(subtype=colSet(sTypes,'Set1'))),
            right_annotation = sigAnno)
  plotFun = function(noFrame=FALSE,noPlot=FALSE){
    draw(hh,row_title='Reference cell signals',column_title='Bulk transcriptomes')
  }
  tt = as.data.frame(norm)
  tt['pR',] = unlist(extras['pR2',])
  tt$rowName = rownames(tt)
  saveFig(file.path(resDir,'allRCC_mKid_heatmap'),plotFun,width=12,height=7,rawData=tt)
}

##############
# Mast scRCC #
##############
scRCCmast = function(){
  #Map ENSEMBL ID to experiment
  expMap = 
    c(`4602STDY7018923` = "Wilms3_Kid_N_C_ldc_1_1", `4602STDY7018924` = "Wilms3_Kid_N_M_ldc_1_1", 
      `4602STDY7018925` = "Wilms3_Kid_N_P_ldc_1_1", `4602STDY7018920` = "Wilms3_Kid_T_ldc_1_1", 
      `4602STDY7018921` = "Wilms3_Kid_T_ldc_1_2", `4602STDY7018922` = "Wilms3_Kid_T_ldc_1_3", 
      `4602STDY7018927` = "Wilms3_Ure_N_ldc_1_1", `4602STDY7090428` = "Wilms2_Kid_N_C_ldc_1_1", 
      `4602STDY7090429` = "Wilms2_Kid_N_M_ldc_1_1", `4602STDY7090430` = "Wilms2_Kid_N_P_ldc_1_1", 
      `4602STDY7090425` = "Wilms2_Kid_T_ldc_1_1", `4602STDY7090426` = "Wilms2_Kid_T_ldc_1_2", 
      `4602STDY7090427` = "Wilms2_Kid_T_ldc_1_3", `4602STDY7090431` = "Wilms2_Ure_N_ldc_1_1", 
      `4602STDY6949193` = "Wilms1_Kid_R_ldc_1_1", `4602STDY6949196` = "Wilms1_Kid_R_ldc_1_2", 
      `4602STDY6949198` = "Wilms1_Kid_R_ldc_1_3", `4602STDY6949192` = "Wilms1_Kid_N_ldc_1_1", 
      `4602STDY6949195` = "Wilms1_Kid_N_ldc_1_2", `4602STDY6949191` = "Wilms1_Kid_T_ldc_1_1", 
      `4602STDY6949194` = "Wilms1_Kid_T_ldc_1_2", `4602STDY6949197` = "Wilms1_Kid_T_ldc_1_3", 
      `4602STDY6949178` = "VHL_Kid_N_ldc_1_1", `4602STDY6949179` = "VHL_Kid_N_ldc_1_2", 
      `4602STDY6949180` = "VHL_Kid_N_ldc_2_1", `4602STDY6949181` = "VHL_Kid_N_ldc_2_2", 
      `4602STDY6949182` = "VHL_Kid_T_ldc_1_1", `4602STDY6949183` = "VHL_Kid_T_ldc_1_2", 
      `4602STDY7018628` = "RCC3_Kid_N_ldc_1_1", `4602STDY7018629` = "RCC3_Kid_N_ldc_1_2", 
      `4602STDY7018630` = "RCC3_Kid_N_ldc_2_1", `4602STDY7018624` = "RCC3_Kid_T_ldc_1_1", 
      `4602STDY7018625` = "RCC3_Kid_T_ldc_1_2", `4602STDY7018626` = "RCC3_Kid_T_ldc_1_3", 
      `4602STDY7018627` = "RCC3_Kid_T_ldc_1_4", `4602STDY7018631` = "RCC3_Ure_N_ldc_1_1", 
      `4602STDY6976426` = "RCC2_Kid_N_ldc_1_1", `4602STDY6976427` = "RCC2_Kid_N_ldc_1_2", 
      `4602STDY6976428` = "RCC2_Kid_N_ldc_1_3", `4602STDY6976422` = "RCC2_Kid_T_ldc_1_1", 
      `4602STDY6976423` = "RCC2_Kid_T_ldc_1_2", `4602STDY6976424` = "RCC2_Kid_T_ldc_2_1", 
      `4602STDY6976425` = "RCC2_Kid_T_ldc_2_2", `4602STDY6930852` = "RCC1_Kid_N_ldc_1_1", 
      `4602STDY6930853` = "RCC1_Kid_N_ldc_1_2", `4602STDY6930856` = "RCC1_Kid_N_ldc_2_1", 
      `4602STDY6930850` = "RCC1_Kid_T_ldc_1_1", `4602STDY6930851` = "RCC1_Kid_T_ldc_1_2", 
      `4602STDY6930854` = "RCC1_Kid_T_ldc_2_1", `4602STDY6930855` = "RCC1_Kid_T_ldc_2_2", 
      `4602STDY6930857` = "RCC1_Ure_N_ldc_1_1", `4602STDY6795897` = "pRCC_Kid_N_ldc_1_1", 
      `4602STDY6795898` = "pRCC_Kid_N_ldc_1_2", `4602STDY6795899` = "pRCC_Kid_T_ldc_1_1", 
      `4602STDY6795900` = "pRCC_Kid_T_ldc_1_2", `4602STDY6949184` = "Trans_Kid_N_L_C_ldc_1_1", 
      `4602STDY6949185` = "Trans_Kid_N_L_C_ldc_2_1", `4602STDY6949187` = "Trans_Kid_N_L_M_ldc_1_1", 
      `4602STDY6949188` = "Trans_Kid_N_L_M_ldc_2_1", `4602STDY6949186` = "Trans_Kid_N_R_C_ldc_1_1", 
      `4602STDY6949189` = "Trans_Kid_N_R_M_ldc_1_1", `4834STDY7002875` = "F16_Kid_N_pos_1_1", 
      `4834STDY7002876` = "F16_Kid_N_neg_1_1", `4834STDY7002881` = "F17_Kid_N_ldf_1_1", 
      `4834STDY7002885` = "F17_Kid_N_pos_1_1", `4834STDY7002886` = "F17_Kid_N_neg_1_1"
      )
  sciKidRef@meta.data$experiment = expMap[as.character(sciKidRef@meta.data$orig.ident)]
  sciKidRef@meta.data$samp = gsub('_.*','',sciKidRef@meta.data$experiment)
  #Make the count table
  cnts = with(sciKidRef@meta.data,table(annot,samp))
  frac = colSums(cnts[grep('MST',rownames(cnts)),])/colSums(cnts)
  frac = rbind(frac,1-frac)
  #Make plot
  plotFun = function(noFrame=FALSE,noPlot=FALSE){
    barplot(frac[,c('pRCC','RCC1','RCC2','RCC3','VHL')])
  }
  tt = as.data.frame(frac)
  tt$rowName = rownames(tt)
  saveFig(file.path(resDir,'scMast'),plotFun,height=4,width=3,rawData=tt)
}

################
# Mast in pRCC #
################
mastRCC = function(){
  #RCCs
  mDat = bigBulkMani[which((bigBulkMani$dataset %in% c('TCGA')) & bigBulkMani$toKeep),]
  #Normal tissue
  tmp = bigBulkMani[which(bigBulkMani$dataset %in% c('GTEX','TCGA') & grepl('Normal',bigBulkMani$biopsy) & bigBulkMani$tissue=='Kidney'),]
  mDat = rbind(mDat,tmp)
  #Define split
  mDat$boxSplit = ifelse(grepl('Normal',mDat$biopsy),'Normal',mDat$disease)
  mDat$boxSplit = ifelse(mDat$boxSplit=='Normal',
                         'Normal',
                         mDat$subtype)
  mDat$boxSplit[mDat$boxSplit == 'chRCC MD'] = 'ChRCC'
  mDat$boxSplit = factor(mDat$boxSplit,levels=rev(c('ChRCC','PRCC Unc','PRCC T1','PRCC T2','KIRP CIMP','ccRCC','Normal')))
  #Do the fit
  srcFile = tempfile()
  write.table(normalizePath(mDat$tgtFile),srcFile,row.names=FALSE,col.names=FALSE,quote=FALSE)
  fit = runAndLoad(srcFile,refs$mKidPBMCandMast$clustSumFile,file.path(outDir,'rccBigFitWithMast'),weights=weightVec)
  #Create plot 
  dat = fit$norm
  mstFrac = colSums(dat[grep('MST',rownames(dat)),])
  toUse = c('Normal','ccRCC','PRCC T1','PRCC T2','ChRCC')
  plotFun = function(noFrame=FALSE,noPlot=FALSE){
    plot(0,
         xlim=c(-0.5,length(toUse)-0.5),
         ylim=c(-7.5,-2),
         xaxt='n',
         type='n',
         frame.plot=FALSE,
         ylab='mast Fraction'
         )
    i=0
    for(nom in toUse){
      w = as.character(mDat$boxSplit)==nom
      boxplot(log10(mstFrac[w]),
              at=i,
              outline=FALSE,
              boxwex=0.95,
              add=TRUE)
      beeswarm(log10(mstFrac[w]),
               add=TRUE,
               at = i,
               pch=19,
               cex=0.8,
               col=pl$colAlpha('black',1/20),
               corral='wrap',
               corralWidth=0.4,
               method='hex'
               )
      i=i+1
    }
    #Re-add axis
    axis(1,
         col=0,
         col.ticks=1,
         labels=toUse,
         at = seq_along(toUse)-1,
         las = 2,
         cex.axis=1)
  }
  tt = as.data.frame(mstFrac)
  tt$rowName = rownames(mstFrac)
  saveFig(file.path(resDir,'mastFracBulk'),plotFun,height=4,width=3,rawData=tt)
  ####Now load the bastards
  ###bDat = loadBulk(mDat$tgtFile)
  ####Convert to TPM
  ###dat = bDat$cnts/bDat$lens
  ###dat = t(t(dat)/colSums(dat))*1e6
  ####Get the genes we're interested in
  ###tgts = c(TPSB2 = 'ENSG00000197253',
  ###         TPSD1 = 'ENSG00000095917',
  ###         TPSAB1 = 'ENSG00000172236'
  ###         )
  ###dat = dat[tgts,]
  ###rownames(dat) = names(tgts)
  ####Boxplots on their side, plus jittered points
  ###plotFun = function(noFrame=FALSE,noPlot=FALSE){ 
  ###  #Make two of them
  ###  par(mfrow=c(1,3))
  ###  ####################################
  ###  # Type A.  This has the axis labels
  ###  plot(0,0,
  ###       type='n',
  ###       xlab='log10(TPM)',
  ###       ylab='Sample Type',
  ###       xlim=c(-3,ceiling(max(log10(dat[1,])))),
  ###       yaxt='n',
  ###       ylim=range(as.numeric(mDat$boxSplit))+c(-0.5,0.5),
  ###       frame.plot=FALSE,
  ###       main=rownames(dat)[1],
  ###       las=1)
  ###  abline(v=0,col='red')
  ###  boxplot(split(log10(dat[1,]),mDat$boxSplit),
  ###          outline=FALSE,
  ###          las=1,
  ###          boxwex=0.95,
  ###          horizontal=TRUE,
  ###          add=TRUE)
  ###  points(log10(dat[1,]),
  ###         as.numeric(mDat$boxSplit) + rnorm(nrow(mDat),0,0.1),
  ###         cex=0.2,
  ###         pch=19,
  ###         col=alpha('black',0.5)
  ###         )
  ###  ########
  ###  # TypeB
  ###  plot(0,0,
  ###       type='n',
  ###       xlab='log10(TPM)',
  ###       ylab='Sample Type',
  ###       xlim=c(-3,ceiling(max(log10(dat[2,])))),
  ###       yaxt='n',
  ###       ylim=range(as.numeric(mDat$boxSplit))+c(-0.5,0.5),
  ###       frame.plot=FALSE,
  ###       main=rownames(dat)[2],
  ###       las=1)
  ###  abline(v=0,col='red')
  ###  boxplot(split(log10(dat[2,]),mDat$boxSplit),
  ###          outline=FALSE,
  ###          las=1,
  ###          boxwex=0.95,
  ###          horizontal=TRUE,
  ###          add=TRUE)
  ###  points(log10(dat[2,]),
  ###         as.numeric(mDat$boxSplit) + rnorm(nrow(mDat),0,0.1),
  ###         cex=0.2,
  ###         pch=19,
  ###         col=alpha('black',0.5)
  ###         )
  ###  ########
  ###  # TypeC
  ###  plot(0,0,
  ###       type='n',
  ###       xlab='log10(TPM)',
  ###       ylab='Sample Type',
  ###       xlim=c(-3,ceiling(max(log10(dat[3,])))),
  ###       yaxt='n',
  ###       ylim=range(as.numeric(mDat$boxSplit))+c(-0.5,0.5),
  ###       frame.plot=FALSE,
  ###       main=rownames(dat)[3],
  ###       las=1)
  ###  abline(v=0,col='red')
  ###  boxplot(split(log10(dat[3,]),mDat$boxSplit),
  ###          outline=FALSE,
  ###          las=1,
  ###          boxwex=0.95,
  ###          horizontal=TRUE,
  ###          add=TRUE)
  ###  points(log10(dat[3,]),
  ###         as.numeric(mDat$boxSplit) + rnorm(nrow(mDat),0,0.1),
  ###         cex=0.2,
  ###         pch=19,
  ###         col=alpha('black',0.5)
  ###         )
  ###}
  ###saveFig(file.path(resDir,'RCCmast'),plotFun)
}

###############
# smFISH Mast #
###############
smMast = function(){
  dat = list.files('Data/smFishMastRCC',full.names=TRUE)
  dat = lapply(dat,read.table,sep='\t',header=TRUE)
  dat = do.call(rbind,dat)
  dat$tType = ifelse(grepl('TM_RCC_00FS',dat$Image),'pRCC2',ifelse(grepl('TM_RCC_00FY',dat$Image),'ccRCC','pRCC1'))
  #Number of cells with mast marker compared to monocytes
  dd = list()
  for(nom in unique(dat$tType)){
    tmp = dat[dat$tType==nom & dat$Name=='PathCellObject',]
    #For each 
    dd[[nom]] = data.frame(tType=nom,
                           nCells=nrow(tmp),
                           nMast = sum(tmp$Subcellular..Channel.4..Num.single.spots>1,na.rm=TRUE) #This is TPSB2, the mast cell marker
                           )
  }
  dd = do.call(rbind,dd)
  dd$mastFrac = dd$nMast/dd$nCells
  #Save table. If Kwasi completes the quantification for other samples, this can be converted to boxplots.
  write.table(dd,file.path(resDir,'smFishQuant.tsv'),sep='\t',row.names=FALSE,col.names=TRUE,quote=FALSE)
}

#########
# chRCC #
#########
chRCC = function(){
  #RCCs
  mDat = bigBulkMani[c(which(bigBulkMani$UniqueSampleID %in% controlSet),which(bigBulkMani$disease=='chRCC' & bigBulkMani$toKeep)),]
  #Define how to split, group, and order
  mDat$boxSplit = ifelse(mDat$dataset=='BenchImmune',
                         'Immune',
                         ifelse(mDat$dataset=='TCGA',
                                'chRCC',
                                'ALL'
                                )
                         )
  isControl = c(chRCC=FALSE,ALL=TRUE,Immune=TRUE)
  #Do the fit
  srcFile = tempfile()
  write.table(normalizePath(mDat$tgtFile),srcFile,row.names=FALSE,col.names=FALSE,quote=FALSE)
  fit = runAndLoad(srcFile,refs$mKidPBMC$clustSumFile,file.path(outDir,'chRCCFit'),weights=weightVec)
  #Create plot 
  dat = fit$norm
  s = rep('Nephron',nrow(dat))
  s[grep('PBMC',rownames(dat))]='Leucocytes'
  s[rownames(dat) %in% paste0('sciKidRef_',c('AV1','AV2','DV','GE'))]='Vascular'
  s[rownames(dat) %in% c('sciKidRef_M','sciKidRef_F')] = 'Fibroblast'
  s[rownames(dat) %in% c('Intercept')]='Unexplained'
  s[rownames(dat) %in% c('sciKidRef_C1')] = 'TypeA'
  s[rownames(dat) %in% c('sciKidRef_C2')] = 'TypeB'
  dat = do.call(rbind,lapply(split(rownames(dat),s),function(e) colSums(dat[e,,drop=FALSE])))
  #Define order
  dat = dat[c('Leucocytes','Vascular','Fibroblast','Nephron','TypeA','TypeB','Unexplained'),]
  #Add in sample grouping
  colnames(dat) = paste0(mDat$boxSplit,'___',colnames(dat))
  #Define signal colours
  cc = setNames(sigCols[paste0(rownames(dat),'_Mature'),'colour'],rownames(dat))
  #Make plot
  plotFun = function(noFrame=FALSE,noPlot=FALSE){
    exposurePlot(dat,isControl=isControl,controlColChange=TRUE,ptAlpha=0.4,colPal=cc)
  }
  tt = as.data.frame(dat)
  tt$rowName = tt
  saveFig(file.path(resDir,'chRCC_mKid'),plotFun,height=5,width=8,rawData=tt)
  #################
  # Heamap version
  ss = !isControl[mDat$boxSplit] & mDat$biopsy=='Primary Tumor' 
  norm = fit$norm[,ss]
  extras = fit$extras[,ss]
  o = order(factor(s,levels=rownames(dat)),sigOrder[rownames(norm),'name'])
  norm = norm[o,]
  #Collapse vascular by mapping sum onto first match
  tmp = as.character(sigOrder[rownames(norm),'name'])
  toKeep = !duplicated(tmp)
  norm[match('Vascular',tmp),] = colSums(norm[tmp=='Vascular',])
  norm = norm[toKeep,]
  sAlt = s[o]
  sAlt = sAlt[toKeep]
  #Construct the annotation for the signals
  sigAnno = HeatmapAnnotation(signalGroup = sAlt,
                              col = list(signalGroup = cc),
                              which='row',
                              show_annotation_name=FALSE
  )
  #Subtypes
  sTypes = mDat$subtype[ss]
  sTypes = ifelse(sTypes=='chRCC MD','Metabolically Divergent',NA)
  hh = csHeatmap(norm,extras,
            row_labels = as.character(sigOrder[rownames(norm),'name']),
            row_split = sigOrder[rownames(norm),'group'],
            topAnno = list(subtype = sTypes,
                           col=list(subtype=c('Metabolically Divergent'='DarkBlue'))),
            right_annotation = sigAnno)
  plotFun = function(noFrame=FALSE,noPlot=FALSE){
    draw(hh,row_title='Reference cell signals')
  }
  tt = as.data.frame(norm)
  tt['pR2',] = unlist(extras['pR2',])
  tt$rowName = rownames(tt)
  saveFig(file.path(resDir,'chRCC_mKid_heatmap'),plotFun,width=12,height=7,rawData=tt)
}


###################
# typeA/B in Bulk #
###################
typeAB = function(){
  #RCCs
  mDat = bigBulkMani[which((bigBulkMani$dataset %in% c('TCGA')) & bigBulkMani$toKeep),]
  #Normal tissue
  tmp = bigBulkMani[which(bigBulkMani$dataset %in% c('GTEX','TCGA') & grepl('Normal',bigBulkMani$biopsy) & bigBulkMani$tissue=='Kidney'),]
  mDat = rbind(mDat,tmp)
  #Define split
  mDat$split = ifelse(grepl('Normal',mDat$biopsy),'Normal',mDat$disease)
  w = mDat$split=='pcRCC'
  mDat$split[w] = mDat$subtype[w]
  cMap = c(chRCC='chRCC',
           'PRCC Unc'='pRCC Unc',
           'KIRP CIMP'='pRCC CIMP',
           'PRCC T1'='pRCC T1',
           'PRCC T2'='pRCC T2',
           ccRCC = 'ccRCC',
           Normal = 'Normal Tissue')
  mDat$split = factor(cMap[mDat$split],levels=cMap)
  #Now load the bastards
  bDat = loadBulk(mDat$tgtFile)
  #Convert to TPM
  dat = bDat$cnts/bDat$lens
  dat = t(t(dat)/colSums(dat))*1e6
  #Get the genes we're interested in
  gns = wilms@misc$geneMapCommon
  genes = c(typeA='SLC4A1',typeB='SLC26A4')
  dat = dat[gns[match(genes,gns[,2]),1],]
  rownames(dat) = names(genes)
  #Boxplots on their side, plus jittered points
  plotFun = function(noFrame=FALSE,noPlot=FALSE){ 
    #Make two of them
    par(mfrow=c(1,2))
    ####################################
    # Type A.  This has the axis labels
    plot(0,0,
         type='n',
         xlab='log10(TPM)',
         ylab='Sample Type',
         xlim=c(-3,ceiling(max(log10(dat[1,])))),
         yaxt='n',
         ylim=range(as.numeric(mDat$split))+c(-0.5,0.5),
         frame.plot=FALSE,
         main=rownames(dat)[1],
         las=1)
    abline(v=0,col='red')
    boxplot(split(log10(dat[1,]),mDat$split),
            outline=FALSE,
            las=1,
            boxwex=0.95,
            horizontal=TRUE,
            add=TRUE)
    points(log10(dat[1,]),
           as.numeric(mDat$split) + rnorm(nrow(mDat),0,0.1),
           cex=0.2,
           pch=19,
           col=alpha('black',0.5)
           )
    ########
    # TypeB
    plot(0,0,
         type='n',
         xlab='log10(TPM)',
         ylab='Sample Type',
         xlim=c(-3,ceiling(max(log10(dat[2,])))),
         yaxt='n',
         ylim=range(as.numeric(mDat$split))+c(-0.5,0.5),
         frame.plot=FALSE,
         main=rownames(dat)[2],
         las=1)
    abline(v=0,col='red')
    boxplot(split(log10(dat[2,]),mDat$split),
            outline=FALSE,
            las=1,
            boxwex=0.95,
            horizontal=TRUE,
            add=TRUE)
    points(log10(dat[2,]),
           as.numeric(mDat$split) + rnorm(nrow(mDat),0,0.1),
           cex=0.2,
           pch=19,
           col=alpha('black',0.5)
           )
  }
  tt = as.data.frame(dat)
  tt$rowName = rownames(tt)
  saveFig(file.path(resDir,'typeAB'),plotFun,width=12,height=7,rawData=tt)
}


####################
# Clinical utility #
####################

###############################
# Sensitivity of core signals #
###############################
sands = function(){
  ###############
  # Prep samples
  mDat = bigBulkMani[!bigBulkMani$dataset %in% c('ALL','ATRT','BenchImmune','NBL') & grepl('Kidney',bigBulkMani$tissue),]
  #Get ones that are labelled as keepers, or are normal.  GTEX samples we care about already pass due to toKeep
  mDat = mDat[mDat$toKeep | (mDat$dataset!='GTEX' & grepl('Normal',mDat$biopsy)),]
  #Drop bilateral wilms and those normal sections with clonal nephrogenesis
  mDat = mDat[-which(mDat$dataset=='WilmsSanger' & (mDat$Bilateral=='Y' | (mDat$Clonal.nephrogenesis=='Yes' & mDat$biopsy=="Normal_tissue"))),]
  ####Drop ones that aren't relevant
  ###mDat = bigBulkMani[!bigBulkMani$dataset %in% c('ALL','ATRT','BenchImmune','NBL','WilmsSanger') & grepl('Kidney',bigBulkMani$tissue),]
  ####Get ones that are labelled as keepers, or are normal.  GTEX samples we care about already pass due to toKeep
  ###mDat = mDat[mDat$toKeep ,]
  ###mDat = mDat[!grepl('Normal',mDat$biopsy),]
  #Make age common
  mDat$ageYears = ifelse(!is.na(mDat$age),#TCGA
                         mDat$age/12,
                         ifelse(!is.na(mDat$AgeMonths),#Wilms Sanger
                                 mDat$AgeMonths/12,
                                 ifelse(!is.na(mDat$Age.at.Diagnosis.in.Days),#Wilms TARGET
                                        mDat$Age.at.Diagnosis.in.Days/365,
                                        ifelse(!is.na(mDat$Age.at.Diagnosis.in.days),#CCSK TARGET
                                               mDat$Age.at.Diagnosis.in.days/365,
                                               NA
                                               )
                                        )
                                 )
                         )
  #Update "disease" to split normal categories by age
  mDat$reducedDisease = mDat$disease
  w = grep('Normal',mDat$biopsy)
  mDat$reducedDisease[w] = ifelse(mDat$dataset[w]=='FoetalMap',
                                  'NormalFoetal',
                                  ifelse(!is.na(mDat$ageYears[w]),
                                         ifelse(mDat$ageYears[w]>18,
                                                'NormalAdult',
                                                'NormalPaed'
                                                ),
                                         'Normal'
                                         )
                                  )
  #Define grouping
  mDat$sampleClass = ifelse(mDat$reducedDisease %in% c('ccRCC','pcRCC','chRCC'),
                          'AdultTumour',
                          ifelse(grepl('Normal',mDat$reducedDisease),
                                 'Normal',
                                 'PaedTumour'
                                 )
                          )
  #Split wilms
  w = which(mDat$reducedDisease=='Wilms')
  mDat$reducedDisease[w] = ifelse(mDat$dataset[w]=='WilmsTARGET','Wilms','Wilms+Chemo')
  #Define order and drop Normal that we can't assign to age group.  Also drop MD chRCC
  mDat$reducedDisease = factor(mDat$reducedDisease,levels=c('NormalAdult','NormalPaed','NormalFoetal','ccRCC','chRCC','pcRCC','CCSK','CMN','Wilms','Wilms+Chemo','MRT'))
  #Compare exposures by class
  classes = list(PT1 = 'sciKidRef_PT1',
                 Vascular = c(paste0('sciKidRef_',c('AV1','AV2','DV','GE'))),#,'fKid_End'),
                 Intercalated = c('sciKidRef_C1','sciKidRef_C2'),
                 #NehprogenicCells = paste0('fKid_',c('CnT','DTLH','ErPrT','NPC','Pod','SSBpr','RVCSB','SSBm.d','SSBpod','UBCD')),
                 NehprogenicCells = paste0('fKid_',c('NPC','RVCSB','SSBm.d','SSBpod','SSBpr')),
                 MPC = 'fKid_MSC'
                 #Interstitial = paste0('fKid_',c('ICa','ICb'))
                 )
  #Map class to disease
  cMap = list(PT1=c('ccRCC','pcRCC'),
           Vascular = 'ccRCC',
           Intercalated = 'chRCC',
           NehprogenicCells = 'Wilms',
           MPC = 'CMN'
           )
  ######
  # Fit
  srcFile = tempfile()
  write.table(normalizePath(mDat$tgtFile),srcFile,row.names=FALSE,col.names=FALSE,quote=FALSE)
  #Fit the model
  fitBoth = runAndLoad(srcFile,refs$kidPBMC$clustSumFile,file.path(outDir,'sandsAll'),weights=weightVec)
  fitDev = runAndLoad(srcFile,refs$fKidPBMC$clustSumFile,file.path(outDir,'sandsDev'),weights=weightVec)
  fitMat = runAndLoad(srcFile,refs$mKidPBMC$clustSumFile,file.path(outDir,'sandsMat'),weights=weightVec)
  ##########
  # Process
  bulkFitCS = fitBoth
  #Decide if fetal or not
  devScore = colSums(fitBoth$norm[grep('fKid_',rownames(fitBoth$norm)),])
  isFetal = devScore>0.25
  #Get intercept score
  intVal = ifelse(isFetal,fitDev$norm['Intercept',],fitMat$norm['Intercept',])
  extras = bulkFitCS$extras
  norm = bulkFitCS$norm
  fit = bulkFitCS$fit
  roc = list()
  med = list()
  for(cNom in names(classes)){
    tgts = classes[[cNom]]
    score = ifelse(isFetal,colSums(fitDev$norm[tgts,,drop=FALSE]),colSums(fitMat$norm[tgts,,drop=FALSE]))
    #intVal = norm['Intercept',]
    df= data.frame(score=score,
                   disease=mDat$reducedDisease,
                   sampleClass = mDat$sampleClass
                   )
    #Drop NAs, low goodness of fit, and weird subtypes
    df = df[!is.na(df$score) & !is.na(df$disease) & intVal<0.4 & (is.na(mDat$subtype) | mDat$subtype!='chRCC MD') & df$disease!='Wilms+Chemo',]
    df$disease = factor(df$disease,levels=setdiff(levels(df$disease),'Wilms+Chemo'))
    med[[cNom]] = df
    gg = ggplot(df,aes(disease,score)) +
      geom_boxplot(aes(colour=disease),outlier.shape=NA,alpha=0.5) +
      geom_jitter(position=position_jitter(width=.1, height=0)) +
      ylim(0,1) + 
      theme(axis.text.x = element_text(angle=45,hjust=1)) +
      scale_colour_manual(values=sampCols[levels(df$disease)]) +
      ggtitle(sprintf("Specificity of %s signal",cNom))
    x = df[df$disease %in% cMap[[cNom]],]
    x = x[order(x$score),]
    #The numerator is number of tumours that are below cut-off (most of which are true negatives) minus the number in this group below threshold (false negatives) giving the number of true negatives
    #The denominator is the number of negatives in total population
    x$specificity = sapply(x$score,function(e) (sum(df$score[df$sampleClass!='Normal']<e)-sum(x$score<e))/(nrow(df[df$sampleClass!='Normal',])-nrow(x)))
    #Becauase the list is sorted, the seq is the number of true positives
    x$sensitivity = seq(nrow(x),1)/nrow(x)
    #Add start and end points
    x = x[c(1,seq(nrow(x)),nrow(x)),]
    x[1,c('sensitivity','specificity','score')] = c(1,0,0)
    x[nrow(x),c('sensitivity','specificity','score')] = c(0,1,1)
    roc[[cNom]] = x
    #pdf(file.path(resDir,sprintf('sands_%s.pdf',cNom)),width=21)
    #plot(gg)
    #dev.off()
  }
  df = do.call(rbind,med)
  df$scoreType = gsub('\\..*','',rownames(df))
  #Relevel to remove empty groups
  df$disease = factor(df$disease,levels=intersect(levels(df$disease),as.character(df$disease)))
  gg = ggplot(df,aes(disease,score)) +
    geom_jitter(position=position_jitter(width=.1, height=0),size=0.2) +
    geom_boxplot(aes(colour=disease),outlier.shape=NA,alpha=0.5) +
    ylim(0,1) + 
    theme(axis.text.x = element_text(angle=45,hjust=1)) +
    scale_colour_manual(values=sampCols[levels(df$disease)]) +
    facet_wrap(~scoreType) + 
    xlab('Sample Type') +
    ylab('Sample specific score') +
    guides(colour=FALSE) + 
    ggtitle(sprintf("Sample Type distribution of sample specific signals"))
  plotFun = function(noFrame=FALSE,noPlot=FALSE){
    plot(gg)
  }
  tt = as.data.frame(df)
  tt$rowName = rownames(tt)
  saveFig(file.path(resDir,'sands_fullBoxplots'),plotFun,width=14,height=7,rawData=tt)
  dat = do.call(rbind,roc)
  dat$scoreType = gsub('\\..*','',rownames(dat))
  #Define colours for each grouping
  cc = c(PT1 = '#2179B4',
         MPC = '#C369AA',
         Vascular = '#E65E5C',
         NehprogenicCells= '#B4D88B',
         Intercalated = '#00A79D')
  plotFun = function(noFrame=FALSE,noPlot=FALSE){
    par(mar=c(7.1, 4.1, 4.1, 9.1))
    plot(dat$specificity,
         dat$sensitivity,
         pch=19,
         cex=0.1,
         col=cc[dat$scoreType],
         xlab='Specificity',
         ylab='Sensitivity')
    #Plot each score type
    for(sType in unique(dat$scoreType)){
      x = dat[dat$scoreType==sType,]
      for(i in seq_len(nrow(x)-1)){
        lines(c(x$specificity[i],x$specificity[i+1]),
              c(x$sensitivity[i],x$sensitivity[i+1]),
            col=cc[sType],
            lwd=1+x$score[i+1]*2)
      }
    }
    abline(1,-1,lty=2)
    legend(x=1.1,
           y=1,
           legend=unique(dat$scoreType),
           col = cc[unique(dat$scoreType)],
           lty = 1,
           lwd = 2,
           xpd=TRUE)
  }
  tt = as.data.frame(dat)
  tt$rowName = rownames(tt)
  saveFig(file.path(resDir,'sands_roc'),plotFun,width=5,height=4,rawData=tt)
  #And a plot using just the medians
  agg = do.call(rbind,med)
  agg$scoreType = gsub('\\..*','',rownames(agg))
  agg= aggregate(score ~ interaction(disease,scoreType),data=agg,FUN=median)
  agg$disease = gsub('\\..*','',agg[,1])
  agg$scoreType = gsub('.*\\.','',agg[,1])
  agg = agg[,c('scoreType','disease','score')]
  agg$disease = factor(agg$disease,levels=names(sampCols))
  #Make plot
  gg = ggplot(agg,aes(scoreType,score,colour=disease)) + 
    #geom_text_repel(aes(label=disease)) + 
    geom_point(size=10) + 
    scale_colour_manual(values = sampCols)
  plotFun = function(noFrame=FALSE,noPlot=FALSE){
    plot(gg)
  }
  tt = as.data.frame(agg)
  tt$rowName = rownames(tt)
  saveFig(file.path(resDir,'sands_meds'),plotFun,width=5,height=4,rawData=tt)
  #par(mar=c(7.1, 4.1, 4.1, 9.1))
  #plot(0,0,
  #     type='n',
  #     xlab='Score Type',
  #     ylab='Median score',
  #     xlim=c(0,nlevels(factor(agg$scoreType))),
  #     ylim=c(0,1),
  #     frame.plot=FALSE,
  #     las=1
  #     )
  #for(i in seq_len(nrow(agg))){
  #  x = as.numeric(factor(agg$scoreType))[i]
  #  lines(c(x-0.3,x+0.3),c(agg$score[i],agg$score[i]),col=sampCols[agg$disease[i]],lwd=2)
  #}
  #agg$disease = factor(agg$disease,levels=unlist(sampColsName))
  #legend(x=5.1,
  #       y=1,
  #       legend = levels(agg$disease),
  #       col = sampCols[levels(agg$disease)],
  #       lty = 1,
  #       lwd = 2,
  #       xpd=TRUE)
  #dev.off()
}

#################################
# Clinically interesting sample #
#################################
clinic = function(){
  #Just want the mystery Wilms and the childhood RCC
  mDat = bigBulkMani[bigBulkMani$UniqueSampleID %in% c('SickKids_X0043_290136','SangerProject1700_PR40309a'),]
  #Do the fit against everything
  srcFile = tempfile()
  write.table(normalizePath(mDat$tgtFile),srcFile,row.names=FALSE,col.names=FALSE,quote=FALSE)
  fit = runAndLoad(srcFile,refs$kidPBMC$clustSumFile,file.path(outDir,'clinicFit'),weights=weightVec)
  extras = fit$extras
  norm = fit$norm
  fit = fit$fit
  imFrac = colSums(norm[grep('^PBMC_',rownames(norm)),])
  matureFrac = colSums(norm[grep('^sciKidRef_',rownames(norm)),])
  devFrac = colSums(norm[grep('^fKid_',rownames(norm)),])
  #Include intercept or not?  This looks at the relative fraction amongst those that are fit 
  tmp = imFrac+matureFrac+devFrac
  intFrac = 1-tmp
  dd = data.frame(devFrac = devFrac,
                  imFrac = imFrac,
                  matureFrac = matureFrac,
                  intFrac = intFrac,
                  disease = mDat$disease,
                  pR2 = t(extras)[,'pR2']
                  )
  #Write table of developmental scores
  #write.table(df,sprintf('Results/%s_foetalScores.tsv',figNom),sep='\t',row.names=TRUE,col.names=TRUE,quote=FALSE)
  #Do detailed fits
  devSamps = mDat[which(dd$devFrac>dd$matureFrac),,drop=FALSE]
  matSamps = mDat[which(dd$devFrac<dd$matureFrac),,drop=FALSE]
  if(nrow(devSamps)>0){
    srcFile = tempfile()
    write.table(normalizePath(devSamps$tgtFile),srcFile,row.names=FALSE,col.names=FALSE,quote=FALSE)
    devFits = runAndLoad(srcFile,refs$fKidPBMC$clustSumFile,file.path(outDir,'clinicFitDev'),weights=weightVec)
  }
  if(nrow(matSamps)>0){
    srcFile = tempfile()
    write.table(normalizePath(matSamps$tgtFile),srcFile,row.names=FALSE,col.names=FALSE,quote=FALSE)
    matFits = runAndLoad(srcFile,refs$mKidPBMC$clustSumFile,file.path(outDir,'clinicFitMat'),weights=weightVec)
  }
  #Save pretty table
  tt = devFits$norm
  tt = tt[order(rownames(tt)),,drop=FALSE]
  tt = data.frame(uniqueSampleID = colnames(tt),
                  cellSignal = rownames(tt),
                  cellSignalContribution = tt[,1])
  tmp = tt
  tt = matFits$norm
  tt = tt[order(rownames(tt)),,drop=FALSE]
  tt = data.frame(uniqueSampleID = colnames(tt),
                  cellSignal = rownames(tt),
                  cellSignalContribution = tt[,1])
  tt = rbind(tmp,tt)
  #write.table(tt,file.path(resDir,paste0(figNom,'_raw.tsv')),sep='\t',row.names=TRUE,col.names=TRUE,quote=FALSE)
  #Plot each of them
  nDots=100
  nRow = 5
  labx=2
  mar = c(1.5,0,1.5,0)
  pch=19
  cex=10
  for(i in seq_len(nrow(mDat))){
    nom = mDat$UniqueSampleID[i]
    score = t(as.matrix(dd[nom,c('devFrac','matureFrac','imFrac','intFrac')]))
    #Now make the dot matrix for the appropriate fit
    if(score['devFrac',]>score['matureFrac',]){
      fit = devFits
      #Merge things and round
      norm = fit$norm
      s = rep('Leucocytes',nrow(norm))
      s[grepl('fKid_',rownames(norm))]='Nephron'
      s[rownames(norm) %in% c('fKid_End','sciKidRef_AV1','sciKidRef_AV2','sciKidRef_DV','sciKidRef_GE')]='Vascular'
      s[rownames(norm) %in% paste0('fKid_',c('MSC'))]='MSC'
      s[rownames(norm) %in% paste0('fKid_',c('ICa','ICb'))]='Mesenchyme'
      s[rownames(norm) %in% paste0('fKid_',c('NPC','RVCSB'))] = 'PrimitiveVesicle'
      s[grepl('_SSB',rownames(norm))] = 'PrimitiveVesicle'
      s[rownames(norm)=='Intercept']='Intercept'
      norm = do.call(rbind,lapply(split(rownames(norm),s),function(e) colSums(norm[e,,drop=FALSE])))
      norm = round(norm*100)
      #Drop things that are zero
      norm = norm[norm[,1]>0,,drop=FALSE]
      #Order
      norm = norm[c('PrimitiveVesicle','Nephron','MSC','Vascular','Leucocytes','Intercept'),,drop=FALSE]
      #Colours
      sCols = sigCols[paste0(rownames(norm),'_Foetal'),'colour']
      names(sCols) = rownames(norm)
    }else{
      fit = matFits
      #Merge things and round
      norm = fit$norm
      s = rep('Leucocytes',nrow(norm))
      s[rownames(norm) %in% c('fKid_End','sciKidRef_AV1','sciKidRef_AV2','sciKidRef_DV','sciKidRef_GE')]='Vascular'
      s[rownames(norm) %in% c('sciKidRef_PT1')]='PT1'
      s[rownames(norm) %in% paste0('sciKidRef_',c('D','G','H','P','PT2','PT3','U1','U2','U3','U4','C1','C2'))]='MatureNephron'
      s[rownames(norm) %in% c('sciKidRef_F','sciKidRef_M')]='Fibroblast'
      s[rownames(norm)=='Intercept']='Intercept'
      norm = do.call(rbind,lapply(split(rownames(norm),s),function(e) colSums(norm[e,,drop=FALSE])))
      norm = round(norm*100)
      #Drop things that are zero
      norm = norm[norm[,1]>0,,drop=FALSE]
      #Order
      norm = norm[c('PT1','Leucocytes','Vascular','Fibroblast','Intercept'),,drop=FALSE]
      #Colours
      sCols = sigCols[paste0(rownames(norm),'_Mature'),'colour']
      names(sCols) = rownames(norm)
    }
    #Dump non-foetalness values
    score[which(rownames(score)!='devFrac'),]=0
    #Make the plot
    barWid = 2
    dotWid = ceiling(nDots/nRow)
    legWid = labx*ceiling(sum(norm>0)/nRow)
    plotFun = function(noFrame=FALSE,noPlot=FALSE){
      layout(matrix(1:3,nrow=1),widths=c(barWid,dotWid,legWid))
      par(mar=c(0,6,0,0))
      tmp = score
      colnames(tmp)=NULL
      barplot(score,
              beside=FALSE,
              ylab='Foetalness',
              ylim=c(0,1),
              cex.axis=3,
              cex.lab=3
              )
      #Create dot-matrix representation
      dotDat = rep(rownames(norm),norm[,1])
      dotCols = sCols
      x = seq(0,nDots-1)
      xx = x%/%nRow
      yy = -(x%%nRow)
      #Margins
      #pdf('~/trueHome/temp.pdf',width=ceiling(nDots/nRow),height=nRow)
      par(mar=mar)
      plot(0,0,type='n',xlab='',ylab='',ylim=range(yy),xlim=range(xx),xaxt='n',yaxt='n',frame.plot=FALSE,las=1)
      points(xx,yy,col=dotCols[dotDat],pch=ifelse(i==1,19,15),cex=cex,xpd=TRUE)
      abline(v=max(xx)+0.5,xpd=TRUE,lwd=2)
      #Manually build legend
      par(mar=mar)
      x = seq(0,length(dotCols)-1)
      xx = x%/%nRow
      yy = -(x%%nRow)
      #How much space for labels and points
      xx = xx*labx
      xMax = labx*ceiling(length(dotCols)/nRow)
      #pdf('~/trueHome/temp.pdf',width=xMax,height=nRow)
      par(mar=mar)
      plot(0,0,type='n',xlab='',ylab='',ylim=range(yy),xlim=c(0,xMax),xaxt='n',yaxt='n',frame.plot=FALSE,las=1)
      points(xx+0.1,yy,col=dotCols,pch=ifelse(i==1,19,15),cex=cex,xpd=TRUE)
      text(xx+0.5,yy,gsub('^[^_]+_','',names(dotCols)),xpd=TRUE,cex=2,pos=4)
    }
    tt = as.data.frame(norm)
    tt$rowName = rownames(tt)
    saveFig(file.path(resDir,sprintf('clin_%s',nom)),plotFun,width=barWid+dotWid+legWid,height=nRow,rawData=tt)
  }
}


################
# Extra tables #
################

############################
# All bulk RNA-seq samples #
############################
allBulk = function(resDir=writeDir) {
  ############
  # Benchmark
  tgts = list()
  tgts[['bench']] = unique(c(controlSet,bigBulkMani$UniqueSampleID[bigBulkMani$dataset %in% c('BenchImmune','ALL')]))
  ############
  # Bench RCC
  mDat = bigBulkMani[which((bigBulkMani$dataset %in% c('TCGA')) & bigBulkMani$toKeep),]
  mDat = mDat[mDat$biopsy=='Primary Tumor' & mDat$disease=='ccRCC',]
  tgts[['benchRCC']]=mDat$UniqueSampleID
  #############
  # Foetalness
  mDat = bigBulkMani[!bigBulkMani$dataset %in% c('ALL','ATRT','BenchImmune','NBL') & grepl('Kidney',bigBulkMani$tissue),]
  #Get ones that are labelled as keepers, or are normal.  GTEX samples we care about already pass due to toKeep
  mDat = mDat[mDat$toKeep | (mDat$dataset!='GTEX' & grepl('Normal',mDat$biopsy)),]
  #Drop bilateral wilms and those normal sections with clonal nephrogenesis
  mDat = mDat[-which(mDat$dataset=='WilmsSanger' & (mDat$Bilateral=='Y' | (mDat$Clonal.nephrogenesis=='Yes' & mDat$biopsy=="Normal_tissue"))),]
  tgts[['foetalScore']] = mDat$UniqueSampleID
  ######
  # CMN
  tgts[['csCMN']] = bigBulkMani$UniqueSampleID[which(bigBulkMani$disease %in% c('CMN') & bigBulkMani$toKeep)]
  ########
  # Wilms
  mDat = bigBulkMani[c(which(bigBulkMani$UniqueSampleID %in% controlSet),which(bigBulkMani$disease %in% c('Wilms','CCSK') & bigBulkMani$toKeep)),]
  #Drop bilateral
  mDat = mDat[-which(mDat$Bilateral=='Y'),]
  tgts[['csWilms']] = mDat$UniqueSampleID
  ######
  # gof
  mDat = bigBulkMani[!bigBulkMani$dataset %in% c('ALL','ATRT','BenchImmune','NBL') & grepl('Kidney',bigBulkMani$tissue),]
  #Get ones that are labelled as keepers, or are normal.  GTEX samples we care about already pass due to toKeep
  mDat = mDat[mDat$toKeep | (mDat$dataset!='GTEX' & grepl('Normal',mDat$biopsy)),]
  #Drop bilateral wilms and those normal sections with clonal nephrogenesis
  mDat = mDat[-which(mDat$dataset=='WilmsSanger' & (mDat$Bilateral=='Y' | (mDat$Clonal.nephrogenesis=='Yes' & mDat$biopsy=="Normal_tissue"))),]
  #Add the inappropriate samples
  mDat = rbind(mDat,bigBulkMani[bigBulkMani$group4 %in% c('B Naive+Memory','T CD4 Memory'),])
  tgts[['gof']]=mDat$UniqueSampleID
  ######
  # MRT
  tgts[['csMRT']] = bigBulkMani$UniqueSampleID[which(bigBulkMani$disease %in% c('MRT') & bigBulkMani$toKeep)]
  tgts[['csMRT_fAd']] = bigBulkMani$UniqueSampleID[which(bigBulkMani$disease %in% c('MRT','NBL') & bigBulkMani$toKeep)]
  ######
  # RCC
  mDat = bigBulkMani[which((bigBulkMani$dataset %in% c('TCGA')) & bigBulkMani$toKeep),]
  #Normal tissue
  tmp = bigBulkMani[which(bigBulkMani$dataset %in% c('GTEX','TCGA') & grepl('Normal',bigBulkMani$biopsy) & bigBulkMani$tissue=='Kidney'),]
  mDat = rbind(mDat,tmp)
  tgts[['rccBigFit']] = mDat$UniqueSampleID
  mDat = bigBulkMani[c(which(bigBulkMani$UniqueSampleID %in% controlSet),which(bigBulkMani$disease=='chRCC' & bigBulkMani$toKeep)),]
  tgts[['chRCC']] = mDat$UniqueSampleID
  ########
  # SANDS
  mDat = bigBulkMani[!bigBulkMani$dataset %in% c('ALL','ATRT','BenchImmune','NBL') & grepl('Kidney',bigBulkMani$tissue),]
  #Get ones that are labelled as keepers, or are normal.  GTEX samples we care about already pass due to toKeep
  mDat = mDat[mDat$toKeep | (mDat$dataset!='GTEX' & grepl('Normal',mDat$biopsy)),]
  #Drop bilateral wilms and those normal sections with clonal nephrogenesis
  mDat = mDat[-which(mDat$dataset=='WilmsSanger' & (mDat$Bilateral=='Y' | (mDat$Clonal.nephrogenesis=='Yes' & mDat$biopsy=="Normal_tissue"))),]
  tgts[['sands']] = mDat$UniqueSampleID
  #########
  # clinic
  mDat = bigBulkMani[bigBulkMani$UniqueSampleID %in% c('SickKids_X0043_290136','SangerProject1700_PR40309a'),]
  tgts[['clinic']] = mDat$UniqueSampleID
  ###########
  # Finalise
  tab = bigBulkMani[unique(unlist(tgts)),]
  tab = tab[,cmnCols[!cmnCols %in% c('toKeep')]]
  #Add extra columns
  tab$SampleSource = tab$source
  tab$ProcessingMethod = 'Rail-RNA'
  tab$ProcessingMethod[tab$source=='TARGET']='TARGET'
  ##############
  # Flow bench #
  ##############
  w = grep('SangerProjectBench',tab$UniqueSampleID)
  tab$SampleSource[w] = 'https://www.cell.com/cell-reports/fulltext/S2211-1247(19)30059-2'
  ###########
  # Cryptic #
  ###########
  w = which(tab$source=='SickKids')
  tab$ProcessingMethod[w] = 'SickKids'
  ##########
  # Normal #
  ##########
  w = grep('Normal',tab$biopsy)
  tab$biopsy[w] = 'Normal'
  tab$SampleSource[w] = gsub('FoetalMap','Sanger',tab$SampleSource[w])
  tab$ProcessingMethod[w] = ifelse(grepl('Foetal',tab$source[w]),'STAR','Rail-RNA')
  ###############
  # Save result #
  ###############
  #Create folder with all bulk data
  dir.create(file.path(resDir,'bulkRNAseq'),showWarnings=FALSE,recursive=FALSE)
  i=1
  for(tgtFile in tab$tgtFile){
    message(tgtFile)
    tgtFolder = gsub('Results/','',dirname(dirname(tgtFile)))
    dir.create(file.path(resDir,tgtFolder),showWarnings=FALSE,recursive=FALSE)
    file.copy(tgtFile,file.path(resDir,tgtFolder,basename(tgtFile)),overwrite=TRUE)
    tab$tgtFile[i] = file.path(tgtFolder,basename(tgtFile))
    i=i+1
  }
  write.table(tab,file.path(resDir,'bulkMani.tsv'),sep='\t',row.names=FALSE,col.names=TRUE,quote=FALSE)
}

###################
# scRNA  manifest #
###################
sampleManifest = function() {
  #Foetal Kidney
  fKid = c("4834STDY7002873", "4834STDY7002874", "4834STDY7002875", "4834STDY7002876","4834STDY7002881", "4834STDY7002885", "4834STDY7002886", "FCAImmP7462242","FCAImmP7462243", "FCAImmP7528292", "FCAImmP7528293", "FCAImmP7555849","FCAImmP7555850", "FCAImmP7579214", "FCAImmP7579215", "w09","w11", "w13", "w16", "w18")
  #Add tumour samples
  srcDir = '~/trueHome/scratch/HCA/CMN1/'
  cmn = list.files('Data/scTumours/CMN/GOSH016/')
  mrtOrg = grep('114[12]$',list.files('Data/scTumours/MRT/Organoids/60'),value=TRUE)
  mrtFresh = gsub('.*CG_SB_([^_]+)_GRCh38.*$','\\1',list.files('Data/scTumours/MRT/GOSH029'))
  mrtNuc
  mrt = c('5640STDY7891141','5640STDY7891142')
  df = data.frame(channelLabel = c(fKid,cmn,mrt),
                  patientLabel = NA,
                  organ = 'Kidney',
                  sampleType = 'Normal',
                  disease = rep(c('Normal','CMN','MRT'),c(length(fKid),length(cmn),length(mrt))),
                  age = NA,
                  sort = NA,
                  dataSource = 'NewData',
                  assay = "10X 3'")
  df$patientLabel[grep('4834STDY700287[34]',df$channelLabel)] = 'Foetus15'
  df$patientLabel[grep('4834STDY700287[56]',df$channelLabel)] = 'Foetus16'
  df$patientLabel[grep('4834STDY700288[156]',df$channelLabel)] = 'Foetus17'
  df$patientLabel[grep('FCAImmP746224[23]',df$channelLabel)] = 'Foetus35'
  df$patientLabel[grep('FCAImmP752829[23]',df$channelLabel)] = 'Foetus38'
  df$patientLabel[grep('FCAImmP75558',df$channelLabel)] = 'Foetus41'
  df$patientLabel[grep('FCAImmP757921',df$channelLabel)] = 'Foetus45'
  df$patientLabel[grep('w[0-9]+',df$channelLabel)] = grep('w[0-9]+',df$channelLabel,value=TRUE)
  df$patientLabel[grep('4602STDY768533',df$channelLabel)] = 'CMN'
  df$patientLabel[grep('5640STDY789114',df$channelLabel)] = 'MRT1'
  df$sampleType[grep('4602STDY768533',df$channelLabel)] = 'Tumour'
  df$sampleType[grep('5640STDY7891141',df$channelLabel)] = 'NormalOrganoid'
  df$sampleType[grep('5640STDY7891142',df$channelLabel)] = 'TumourOrganoid'
  df$sort[grep('4834STDY7002873|4834STDY7002875|4834STDY7002885|FCAImmP7462242|FCAImmP7528292|FCAImmP7555849|FCAImmP7579214',df$channelLabel)]='CD45+'
  df$sort[grep('4834STDY7002874|4834STDY7002876|4834STDY7002886|FCAImmP7462243|FCAImmP7528293|FCAImmP7555850|FCAImmP7579215',df$channelLabel)]='CD45-'
  df$sort[grep('4834STDY7002881',df$channelLabel)]='None'
  df$sort[grep('4602STDY768533',df$channelLabel)] = 'None'
  df$sort[grep('5388STDY',df$channelLabel)] = 'None'
  df$sort[grep('5640STDY',df$channelLabel)] = 'None'
  df$age[grep('4834STDY700287[34]',df$channelLabel)] = '12 PCW'
  df$age[grep('4834STDY700287[56]',df$channelLabel)] = '8+1 PCW'
  df$age[grep('4834STDY700288[156]',df$channelLabel)] = '9+1 PCW'
  df$age[grep('FCAImmP746224[23]',df$channelLabel)] = '7+6 PCW'
  df$age[grep('FCAImmP752829[23]',df$channelLabel)] = '13+6 PCW'
  df$age[grep('FCAImmP75558',df$channelLabel)] = '16+6 PCW'
  df$age[grep('FCAImmP757921[45]',df$channelLabel)] = '13+6 PCW'
  #Need ages for F38,41,45
  df$dataSource[grep('4834STDY',df$channelLabel)] = 'DOI: 10.1126/science.aat1699'
  df$dataSource[grep('w[0-9]',df$channelLabel)] = 'DOI: 10.1371/journal.pbio.3000152'
  #Add Wilms 
  df = rbind(df,data.frame(
                           channelLabel = c("NB8350521", "NB8350522", "NB8350523", "NB8368876", "NB8368877"),
                           patientLabel = 'Wilms1',
                           organ = 'Kidney',
                           sampleType = 'Tumour',
                           disease = 'Wilms',
                           age = NA,
                           sort = c('None','CD45-','CD45-','None','CD45-'),
                           dataSource = 'NewData',
                           assay = "10X 3'"))
  #CCSK
  df = rbind(df,data.frame(
                           channelLabel = c("NB8711738",'NB8711739','NB8711740','NB8711741','NB8711742','TM211','TM212'),
                           patientLabel = c('CCSK1','CCSK1','CCSK2','CCSK2','CCSK3','1015T1','1015T1'),
                           organ = 'Kidney',
                           sampleType = 'Tumour',
                           disease = 'CCSK',
                           age = NA,
                           sort = NA,
                           dataSource = 'NewData',
                           assay = rep(c("Nuclei 10X 3'",'CEL-Seq2'),c(5,2))))
  #MRT fresh
  df = rbind(df,data.frame(
                           channelLabel = c("NB8350518", "NB8350519", "NB8350520", "NB8368300", "NB8368301"),
                           patientLabel = 'MRT2',
                           organ = 'Kidney',
                           sampleType = 'Tumour',
                           disease = 'MRT',
                           age = NA,
                           sort = c('None','CD45-','CD45-','None','CD45-'),
                           dataSource = 'NewData',
                           assay = "10X 3'"))
  #MRT Nuclear
  df = rbind(df,data.frame(
                           channelLabel = c("NB8711736",'NB8711737'),
                           patientLabel = 'MRT3',
                           organ = 'Kidney',
                           sampleType = 'Tumour',
                           disease = 'MRT',
                           age = NA,
                           sort = NA,
                           dataSource = 'NewData',
                           assay = "Nuclei 10X 3'"))
  #Save table at last
  write.table(df,file.path(resDir,'sampleMani.tsv'),row.names=FALSE,sep='\t',col.names=TRUE,quote=FALSE)
}


#################################
# Figure and table construction #
#################################

#####################################
# Figure1 and Supplementary Figure 1/2
fKidPlot()
schematic()
bench()
benchRCC()

#########################
# Figure 2 and Table 3/4
foetalScore()

########################################
# Figure 3 and Supplementary Figure 3/4
csCMN()
genesCMN()
csCMN_devComp()
umapCMN()
lrCMN()

########################################
# Figure 4 and Supplementary Figure 5-7
csWilms()
umapWilms()
simWilms()
gof()

#########################################
# Figure 5 and Supplementary Figure 8-10
csMRT()
scMRT_plot()
scMRT_LR()

##########################################
# Figure 6 and Supplementary Figure 11-14
rccBigFit()
scRCCmast()
mastRCC()
chRCC()
typeAB()

##########
# Table 5
smMast()

##########################################
# Figure 7 and Supplementary Figure 14/15
sands()
clinic()

##########################
# Supplementary Figure 17
colSchemes()

###########################
# Data underlying figures #
###########################
#Specify which file contains the data for which figure
tabMap = c(Fig1B = 'fKidRef_rawData.tsv',
           Fig1C = 'fKidRefAge_rawData.tsv',
           Fig1D = 'benchCS_rawData.tsv',
           Fig2A = 'immScoreNormal_rawData.tsv',
           Fig2B = 'immScoreAdult_rawData.tsv',
           Fig2C = 'immScorePaed_rawData.tsv',
           Fig2D = 'immScore_ccRCC_mRNA_rawData.tsv',
           Fig2E = 'immScore_RCC_PTEN_rawData.tsv',
           Fig3A = 'CMN_fKid_rawData.tsv',
           Fig3B = 'CMN_genes_rawData.tsv',
           Fig3C = 'CMN_devComp_rawData.tsv',
           Fig3D = 'CMN_UMAP_rawData.tsv',
           Fig3E = 'lrCMN_fKid_rawData.tsv',
           Fig4A = 'csWilms_fKid_rawData.tsv',
           Fig4BLeft = 'Wilms_umap_leftPan_rawData.tsv',
           Fig4BRight = 'Wilms_umap_rightPan_rawData.tsv',
           Fig4C = 'lrWilms_fKid_rawData.tsv',
           Fig4D = 'GOF_rawData.tsv',
           Fig5A = 'MRT_fKid_rawData.tsv',
           Fig5B = 'scMRT_rawData.tsv',
           Fig5CHuman = 'lrMRT_fKid_rawData.tsv',
           Fig5CMouse = 'lrMRT_mouseDev_rawData.tsv',
           Fig6A = 'allRCC_mKid_rawData.tsv',
           Fig6B = 'scMast_rawData.tsv',
           Fig6C = 'mastFracBulk_rawData.tsv',
           Fig6E = 'chRCC_mKid_rawData.tsv',
           Fig7A = 'sands_roc_rawData.tsv',
           Fig7B = 'sands_meds_rawData.tsv',
           Fig7DE = 'clin_SangerProject1700_PR40309a_rawData.tsv',
           Fig7FG = 'clin_SickKids_X0043_290136_rawData.tsv',
           SupFig1A = 'fKidRefClusters_rawData.tsv',
           SupFig1B = 'fKidRefClasses_rawData.tsv',
           SupFig1C = 'fKidRefGenes_rawData.tsv',
           SupFig1D = 'MPCsimilarity_rawData.tsv',
           SupFig2 = 'BenchRCC_rawData.tsv',
           SupFig3 = 'CMN_fKid_heatmap_rawData.tsv',
           SupFig4 = 'CMNRefGenes_rawData.tsv',
           SupFig5 = 'csWilms_fKid_heatmap_rawData.tsv',
           SupFig6A = 'WilmsRefGenes_rawData.tsv',
           SupFig6B = 'WilmsRefPhase_rawData.tsv',
           SupFig7 = 'CCSKRefGenes_rawData.tsv',
           SupFig8 = 'MRT_fKid_heatmap_rawData.tsv',
           SupFig9 = 'MRTRefGenes_rawData.tsv',
           SupFig10 = 'MRTscTWIST1_rawData.tsv',
           SupFig11 = 'allRCC_mKid_heatmap_rawData.tsv',
           SupFig12 = 'typeAB_rawData.tsv',
           SupFig13 = 'chRCC_mKid_heatmap_rawData.tsv',
           SupFig14 = 'sands_fullBoxplots_rawData.tsv'
           )
#Rename them and put them all in a folder
tgtFolder = file.path(resDir,'dataForFigures')
dir.create(tgtFolder,showWarnings=FALSE,recursive=FALSE)
for(nom in names(tabMap)){
  message(nom)
  file.copy(file.path(resDir,tabMap[[nom]]),file.path(tgtFolder,paste0(nom,'.tsv')),overwrite=TRUE)
}


    
```
