## Supplementary material for "Single cell derived mRNA signals across human kidney tumors": Source Code: scQC.R

```
#' Mostly basic QC and filtering functions.  Depends on scrublet being installed.  Also has some other general single cell helper functions.

#############
# Libraries #
#############
library(ggplot2)
library(reshape2)
library(Seurat)
library(ComplexHeatmap)
library(Matrix)

#############
# Functions #
#############

#' Parse and extract a set of prefixes.
#'
#' Used by fPlot.  Pass the features and prefixes are extracted and returned if valid.
#'
#' @param features A set of features from which prefixes are to be extracted.
#' @param splitChr Character to split on.
parsePrefix = function(features,splitChr=';'){
  tmp = strsplit(features,splitChr)
  out = vector(mode='list',length=length(tmp))
  for(i in seq_along(tmp)){
    #Can we even split it?
    if(length(tmp[[i]])==1){
      out[[i]]$valid=FALSE
      prefix=''
      feature=tmp[[i]]
    }else{
      out[[i]]$valid = TRUE
      #Get the first bit as the prefix
      prefix = tmp[[i]][1]
      feature = paste(tmp[[i]][-1],collapse=splitChr)
    }
    out[[i]]$feature = feature
    out[[i]]$prefix = prefix
    out[[i]]$flags = list()
    #Now we have a prefix, work with it
    patterns = c(log='log$',log2='log2$',log10='log10$')
    for(nom in names(patterns)){
      pattern = patterns[nom]
      out[[i]]$flags[[nom]] = grepl(prefix,pattern)
      prefix = gsub(pattern,'',prefix)
    }
    #It's only valid if we've consumed the whole prefix
    out[[i]]$valid = prefix=='' && out[[i]]$valid
    #If it's not valid, reset all flags
    if(!out[[i]]$valid)
      for(j in seq_along(out[[i]]$flags))
        out[[i]]$flags[j] = FALSE
  }
  return(out)
}


#' Reimplementation of FeaturePlot/DimPlot
#'
#' @param srat Seurat object
#' @param features Things to plot.  Can prefix with 'log' to show log base 10 of value.
#' @param reduction Which reduction to use.
#' @param dims Which dimensions to plot.
#' @param colScheme Options are logit, gene, default.  If default, nothing is specified and user can over-write.
#' @param ptSize Size of points.
#' @param ... Passed to geom_point
#' @return ggplot2 object
fPlot =function(srat,features,reduction='umap',dims=c(1,2),colScheme=c('gene','logit'),cex=0.3,pch=19,zLims=c(NA,NA),nrow=NULL,ncol=NULL,catLab=TRUE,catLeg=FALSE,...){
  #Get the colour scheme
  colScheme = match.arg(colScheme)
  #Get the data
  labs = paste0(Key(srat[[reduction]]), dims)
  data = FetchData(srat, vars = c(labs, "ident", features))
  #Check if there are any missing that we can try and get
  modifiers = parsePrefix(features)
  isVal = sapply(modifiers,function(e) e$valid)
  w = !(features %in% colnames(data)) & isVal
  idxFeats = features
  idxFeats[w] = unlist(lapply(modifiers[w],function(e) e$feature))
  #Try and get the modified ones
  if(any(w))
    data = cbind(data,FetchData(srat,vars=idxFeats[w]))
  #Drop ones that aren't available
  w = which(idxFeats %in% colnames(data))
  idxFeats = idxFeats[w]
  modifiers = modifiers[w]
  #Get number of plots
  nPlots = wrap_dims(sum(idxFeats %in% colnames(data)),nrow=nrow,ncol=ncol)
  #Set the bastards
  par(mfrow=nPlots)
  #Get a vector of those that are numeric
  isNum = sapply(idxFeats,function(e) is.numeric(data[,e]))
  #Rearrange to put the non-numeric things first
  o = order(isNum)
  idxFeats = idxFeats[o]
  isNum = isNum[o]
  modifiers = modifiers[o]
  #Get the zlims if not given
  if(any(isNum) && (is.na(zLims[1]) || is.na(zLims[2]))){
    tmp = range(data[,idxFeats[isNum]],na.rm=TRUE)
    zLims[is.na(zLims)] = tmp[is.na(zLims)]
  }
  #Now do the plotting one at a time.
  for(i in seq_along(idxFeats)){
    idxFeat = idxFeats[i]
    #Decide if we need left/bottom axes
    leftAxes = i%%nPlots[2]==1
    bottomAxes = i> prod(nPlots)-nPlots[2]
    #Set the margins
    par(mar=c(bottom=ifelse(bottomAxes,5,1),
              left=ifelse(leftAxes,4,1),
              top=1,
              right=2))
    #Make the plot area
    plot(data[,labs[1]],data[,labs[2]],
         type='n',
         frame.plot=FALSE,
         xlab=ifelse(bottomAxes,labs[1],''),
         ylab=ifelse(leftAxes,labs[2],''),
         main=idxFeat,
         yaxt=ifelse(leftAxes,'s','n'),
         xaxt=ifelse(bottomAxes,'s','n')
         )
    #Decide what type it is.
    x = data[,idxFeat]
    if(isNum[i]){
      #Decide on the colour scheme
      if(colScheme=='logit'){
        ccCont = circlize::colorRamp2(seq(-4,4,length.out=6),c('#8c510a','#d8b365','#f6e8c3','#c7eae5','#5ab4ac','#01665e'))
      }else if(colScheme=='gene'){
        ccCont = circlize::colorRamp2(zLims,c('grey','blue'))
      }
      cc = ccCont
    }else{
      #Now add the non-ones.
      if(!is.factor(x)){
        x = factor(x)
      }
      #Make a colour scheme
      cBase = suppressWarnings(RColorBrewer::brewer.pal(nlevels(x),'Set1'))
      if(nlevels(x)==length(cBase)){
        cc = function(e) setNames(cBase,levels(x))[e]
      }else{
        cNew = circlize::colorRamp2(seq(0,1,length.out=length(cBase)),cBase)(seq(0,1,length.out=nlevels(x)))
        cc = function(e) setNames(cNew,levels(x))[e]
      }
    }
    points(data[,labs[1]],
           data[,labs[2]],
           col=cc(x),
           pch=pch,
           cex=cex)
    #Extra stuff for category plots
    if(!isNum[i]){
      tmp = cbind(data[,labs],x)
      colnames(tmp) = c('RD1','RD2','x')
      mids = aggregate(cbind(RD1,RD2) ~ x,FUN=mean,data=tmp)
      if(catLab)
        text(mids[,2],mids[,3],mids[,1])
      #Legend
      if(catLeg){
        legend(x=max(data[,labs[1]])*1.1,
               y=max(range(data[,labs[2]])),
               legend = levels(x),
               col = cc(levels(x)),
               pch=pch,
               bty='n',
               y.intersp=0.5,
               xpd=NA
        )
      }
    }
    #Add the numeric colour bar at the end of any row with numeric things.
    if(any(isNum) && (i%%nPlots[2]==0 || i==length(idxFeats)) && isNum[i])
      pp$addColorBar(x=max(data[,labs[1]])*1.1,
                     y=max(range(data[,labs[2]])),
                     col=rgb(attributes(ccCont)$colors),
                     breaks=rev(attributes(ccCont)$breaks),
                     ticks=5,
                     barFrac=0.8,
                     title='value'
      )
  }
  return(NULL)
}


#' Feature plot, but less retarded.
#'
#' @param srat Seurat object
#' @param features Things to plot.
#' @param reduction Which reduction to use.
#' @param dims Which dimensions to plot.
#' @param colScheme Options are logit, gene, default.  If default, nothing is specified and user can over-write.
#' @param ptSize Size of points.
#' @param ... Passed to geom_point
#' @return ggplot2 object
fPlot =function(srat,features,reduction='umap',dims=c(1,2),colScheme=c('gene','logit','default'),ptSize=0.3,...){
  colScheme = match.arg(colScheme)
  #Get the data
  labs = paste0(Key(srat[[reduction]]), dims)
  data = FetchData(srat, vars = c(labs, "ident", features))
  dd = melt(data,id.vars=c(labs,'ident'))
  #Now make the plots
  gg = ggplot(dd,aes_string(labs[1],labs[2])) +
    geom_point(aes(colour=value),size=ptSize,...) +
    theme_classic() +
    facet_wrap(~variable)
  #Logit cols
  if(colScheme=='logit'){
    gg = gg +
      scale_colour_gradientn(colours= c('#8c510a','#d8b365','#f6e8c3','#c7eae5','#5ab4ac','#01665e'),limits=c(-4,4))
  }else if(colScheme=='gene'){
    gg = gg +
      scale_colour_gradient(low='grey',high='blue')
  }
  return(gg)
}

#' Quick default proccessing with seurat.
#'
#' @param dat Raw filtered count matrix.
#' @param numPCs How many PCs to use.
#' @param clusteringRes Clustering parameter to use.
#' @return Seurat object.
quickCluster = function(dat,numPCs=50,clusteringRes=1.0){
  ss = CreateSeuratObject(dat)
  ss = NormalizeData(ss,verbose=FALSE)
  ss = FindVariableFeatures(ss,verbose=FALSE)
  ss = ScaleData(ss,verbose=FALSE)
  ss = RunPCA(ss,npcs=numPCs,approx=FALSE,verbose=FALSE)
  ss = FindNeighbors(ss,dims=seq(numPCs),verbose=FALSE)
  ss = FindClusters(ss,res=clusteringRes,verbose=FALSE)
  ss = RunUMAP(ss,dims=seq(numPCs),verbose=FALSE)
  return(ss)
}

#' Compare fraction of overlapping barcodes
#'
#' Takes a collection of filtered cell directories (that can be Read10X'ed) and calculates the overlapping fraction between them.
#'
#' @param srcDirs Directories that cell barcodes are to be loaded from.  Should be a filtered folder.  Ideally named.
#' @param spliter Groups samples by this when plotting.
compBarcodes = function(srcDirs,splitter=NULL){
  if(is.null(names(srcDirs)))
    names(srcDirs) = paste0('SeuratObject',seq_along(srcDirs))
  dat = Read10X(srcDirs)
  barcodes = data.frame(row.names=colnames(dat),
                        barcode = gsub('.*_','',colnames(dat)),
                        source = gsub('_[ACGT]+(-[0-9]+)?$','',colnames(dat)))
  #Now split and do the comparison
  comp = list()
  breaks = factor(barcodes$source)
  for(left in levels(breaks)){
    for(right in levels(breaks)){
      comp[[length(comp)+1]] = data.frame(left = left, 
                                          right=right, 
                                          leftCnt = sum(breaks==left),
                                          rightCnt = sum(breaks==right),
                                          overlap = sum(barcodes$barcode[breaks==left] %in% barcodes$barcode[breaks==right])
                                          )
    }
  }
  comp = do.call(rbind,comp)
  comp$frac = round(comp$overlap/comp$leftCnt *100,2)
  #Make a matrix and heatmap
  mat = acast(comp,left ~ right,value.var='frac')
  mat = mat[names(srcDirs),names(srcDirs),drop=FALSE]
  #Make another with the overlap to stick on the heatmap
  matOlap = acast(comp,left ~ right,value.var='overlap')
  matOlap = matOlap[names(srcDirs),names(srcDirs),drop=FALSE]
  col = suppressWarnings(RColorBrewer::brewer.pal(1000,'Spectral'))
  col = circlize::colorRamp2(seq(0,100,length.out=length(col)),rev(col))
  hh = Heatmap(mat,
            name='% Olap w/ Left',
            col=col,
            cell_fun = function(j,i,x,y,width,height,fill) grid.text(sprintf('%d',matOlap[i,j]),x,y,gp = gpar(fontsize=10)),
            row_split = splitter,
            column_split = splitter,
            row_title_rot=0,
            column_title_rot=90,
            cluster_rows=FALSE,
            cluster_columns=FALSE)
  return(list(comp=comp,mat=mat,hmap=hh))
}


#' Initial filtering and metadata
#'
#' Calculates metadata, drop cells based on QC metrics.  Doesn't require the pre-processing structure of loading things into unswappedCounts and cleanCounts, along with the corresponding automatically generated files.  But if these are given, the extra files and meta-data will be used and loaded into the resulting Seurat object.
#'
#' @param dataDirs Data directories to load, passed to Read10X.
#' @param excludeGenes Genes that will be dropped from the count matrix.
#' @param geneSets Sets of genes to aggregate counts for and store as metadata.  Name by list element names.  Must include an entry named "mtGenes".
#' @param sPhaseGenes Genes used to identify S phase genes.  If NULL, will be loaded from Seurat.
#' @param g2mPhaseGenes Genes used to identify G2M phase genes.  If NULL, will be loaded from Seurat.
#' @param maxMT Maximum fraction of expression from Mitochondrial genes allow per cell.
#' @param minGenes Minimum number of genes for a cell to express and pass QC.
#' @param minUMIs Minimum number of UMIs for a cell to have and pass QC.
#' @param maxBadFrac Maximum fraction of cells not passing QC in a cluster before we dump the whole cluster.
#' @param skipScrub Should we skip running scrublet, even if it's not available.
#' @param scrubScoreMax Mark a cell as a doublet if its scrublet score exceeds this value, even if scrublet does not call it as such.
#' @param numPCs Number of PCs to use for the rough clustering and doublet identification.  This is just used for generating very fine grained clustering and it shouldn't matter much what you set it to.
#' @param clusteringRes Resolution for fine clustering.  Should be very large.
#' @param scrubPath Relative path to file containing scrublet run.  If NULL, scrublet will be run on data. 
#' @param scPath Path to where SoupChannel objects created by SoupX are saved in RDS format.  If not found, ignored.
#' @param cellCallPath Path to file detailing the reason each barcode is called as a cell. 
#' @param doPlot Make some QC summary plots.
#' @param verbose Shut up or not?
#' @param ... Parameters passed to \code{Read10X}
#' @return A Seurat object with cells dropped that fail the QC filters.
basicQC = function(dataDirs,excludeGenes = c(),geneSets=list(mtGenes=c("MT-ND1", "MT-ND2", "MT-CO1", "MT-CO2", "MT-ATP8", "MT-ATP6","MT-CO3", "MT-ND3", "MT-ND4L", "MT-ND4", "MT-ND5", "MT-ND6","MT-CYB")),sPhaseGenes=NULL,g2mPhaseGenes=NULL,maxMT=0.2,minGenes=300,minUMIs=1000,maxBadFrac=0.5,skipScrub=FALSE,scrubScoreMax = 0.5,numPCs=75,clusteringRes=10.0,scrubPath='../cleanCounts/scrubletScores.tsv',scPath = './SoupChannelOjbect.RDS',cellCallPath = '../cleanCounts/filteredBarcodes.tsv',doPlot=TRUE,verbose=TRUE,...){
  if(is.null(dataDirs)){
    warning("Data directories not named, generating automatic names.")
    names(dataDirs) = paste0('DataSource',seq_along(dataDirs))
  }
  if(verbose)
    message('Loading data.')
  srat = Read10X(dataDirs,...)
  srat = CreateSeuratObject(srat)
  #Get first bonus bit of meta-data, what method(s) passed this barcode as a cell.
  tgts = file.path(dataDirs,cellCallPath)
  if(all(file.exists(tgts))){
    if(verbose)
      message("Loading barcode filtering to cell meta-data.")
    keepCols = paste0('pass',c('Knee','Inf','ED','CR',''))
    for(keepCol in keepCols)
      srat@meta.data[,gsub('pass','filt',keepCol)]=NA
    for(i in seq_along(dataDirs)){
      filtDat = read.table(tgts[i],sep='\t',header=TRUE)
      #Make barcodes like with the load function
      if(all(gsub('.*-','',filtDat$barcode)=='1'))
        filtDat$barcode = gsub('-1','',filtDat$barcode)
      #Save the relevant information in meta.data
      m = match(paste0(names(dataDirs[i]),'_',filtDat$barcode),rownames(srat@meta.data))
      o = !is.na(m)
      for(keepCol in keepCols)
        srat@meta.data[m[o],gsub('pass','filt',keepCol)] = filtDat[o,keepCol]
    }
    srat@meta.data$reasonForFilt = with(srat@meta.data,
                                        ifelse(xor(filtKnee,filtED),
                                               ifelse(filtKnee,
                                                      'Knee',
                                                      'EmptyDrops'
                                                      ),
                                               'Both'
                                               )
                                        )
  }
  #Check for SoupX objects.
  tgts = file.path(dataDirs,scPath)
  srat@misc$sc=list()
  srat@misc$soupFrac = data.frame(dataDir=dataDirs,soupFrac=NA)
  for(i in seq_along(dataDirs)){
    if(file.exists(tgts[i])){
      srat@misc$sc[[i]] = readRDS(tgts[i])
      srat@misc$soupFrac[i,'soupFrac'] = srat@misc$sc[[i]]$metaData$rho[1]
    }else{
      srat@misc$sc[[i]]=NA
    }
  }
  #Load and store the gene objects
  if(verbose)
    message("Loading gene meta-data.")
  srat@misc$geneMap = lapply(dataDirs,function(e) {
                               if(file.exists(file.path(e,'features.tsv.gz'))){
                                 x = read.table(file.path(e,'features.tsv.gz'),sep='\t',header=FALSE)
                               }else{
                                 x = read.table(file.path(e,'genes.tsv'),sep='\t',header=FALSE)
                               }
                               #Do the Seurat gene name conversion.  Assume column 2.
                               x[,2] = make.unique(gsub('_','-',x[,2]))
                               return(x)})
  #Check if they're all the same.
  srat@misc$geneMapCommon=TRUE
  for(i in seq_along(dataDirs)){
    if(i==1){
      base = srat@misc$geneMap[[i]]
    }else{
      tst = srat@misc$geneMap[[i]]
      #Check they have the same dimensions
      if(!all(dim(base)==dim(tst))){
        srat@misc$geneMapCommon=FALSE
        next
      }
      #Check they're all the same
      for(j in seq(ncol(base)))
        if(!all(base[,j]==tst[,j]))
          srat@misc$geneMapCommon=FALSE
    }
  }
  #If they're all the same, store the same one
  if(srat@misc$geneMapCommon){
    srat@misc$geneMapCommon = base
    rownames(srat@misc$geneMapCommon) = base[,2]
  }else{
    srat@misc$geneMapCommon = NULL
  }
  if(is.null(names(geneSets))){
    warning("No names given for gene sets.  Setting to uninformative auto-generated values.")
    names(geneSets) = paste0('geneSet',seq_along(geneSets))
  }
  if(is.null(sPhaseGenes) | is.null(g2mPhaseGenes)){
    data('cc.genes.updated.2019',package='Seurat')
  }
  if(is.null(sPhaseGenes))
    sPhaseGenes = cc.genes.updated.2019$s.genes
  if(is.null(g2mPhaseGenes))
    g2mPhaseGenes = cc.genes.updated.2019$g2m.genes
  #Get extra meta-data things
  if(verbose)
    message("Generating extra meta-data.")
  for(nom in names(geneSets)){
    tmp = geneSets[[nom]]
    tmp = tmp[tmp %in% rownames(srat@assays$RNA@data)]
    if(length(tmp)==0)
      next
    srat@meta.data[,nom] = colSums(srat@assays$RNA@counts[tmp,,drop=FALSE])/srat@meta.data$nCount_RNA
  }
  #Get cell cycle
  srat = NormalizeData(srat,verbose=FALSE)
  #The seeed parameter magic is needed because the Seurat authors are dicks
  srat = CellCycleScoring(srat, s.features=sPhaseGenes,g2m.features=g2mPhaseGenes,seed=sample(1e9,1))
  #Call doublets
  if(verbose)
    message("Loading or running scrublet.")
  srat@meta.data$scrubScore=NA
  srat@meta.data$scrubCall=NA
  srat@meta.data$scrubSim1=NA
  srat@meta.data$scrubSim2=NA
  if(!is.null(scrubPath)){
    scrubRun = rep(FALSE,length(dataDirs))
  }else{
    scrubRun = !file.exists(file.path(dataDirs,scrubPath))
  }
  for(i in seq_along(dataDirs)){
    if(scrubRun[i]){
      if(verbose)
        message(sprintf('Running for channel %s',dataDirs[i]))
      scrubScores = runScrublet(dataDirs[i],nPCs=numPCs)
      #Make barcodes like with the load function
      if(all(gsub('.*-','',scrubScores$barcode)=='1'))
        scrubScores$barcode = gsub('-1','',scrubScores$barcode)
    }else{
      scrubScores = read.table(file.path(dataDirs[i],scrubPath),sep='\t',header=TRUE)
    }
    #Save the relevant information in meta.data
    m = match(paste0(names(dataDirs[i]),'_',scrubScores$barcode),rownames(srat@meta.data))
    o = !is.na(m)
    srat@meta.data$scrubScore[m[o]] = scrubScores$score[o]
    srat@meta.data$scrubCall[m[o]] = scrubScores$call[o]
    srat@meta.data$scrubSim1[m[o]] = scrubScores$simScore1[o]
    srat@meta.data$scrubSim2[m[o]] = scrubScores$simScore2[o]
  }
  #Do basic processing and clustering
  if(verbose)
    message("Basic processing to complete QC.")
  srat = FindVariableFeatures(srat,verbose=FALSE)
  srat = ScaleData(srat,verbose=FALSE)
  srat = RunPCA(srat,npcs=numPCs,approx=FALSE,verbose=FALSE)
  srat = FindNeighbors(srat,dims=seq(numPCs),verbose=FALSE)
  srat = FindClusters(srat,res=clusteringRes,verbose=FALSE)
  srat = RunUMAP(srat,dims=seq(numPCs),verbose=FALSE)
  #Set the basic filters.  Default to TRUE if NA
  srat@meta.data$PASS_MT = !(srat@meta.data$mtGenes >= maxMT)
  srat@meta.data$PASS_nGenes = !(srat@meta.data$nFeature_RNA <= minGenes)
  srat@meta.data$PASS_nCounts = !(srat@meta.data$nCount_RNA <= minUMIs)
  srat@meta.data$PASS_doublet = !(!is.na(srat@meta.data$scrubCall) & (srat@meta.data$scrubCall | srat@meta.data$scrubScore > scrubScoreMax))
  #Work out which ones don't pass because of clustering
  tt = aggregate(!PASS_MT | !PASS_nGenes | !PASS_nCounts | !PASS_doublet ~ seurat_clusters,FUN=mean,data=srat@meta.data)
  srat@meta.data$fracBadClust = tt[match(srat@meta.data$seurat_clusters,tt[,1]),2]
  srat@meta.data$PASS_cluster = !(srat@meta.data$fracBadClust > maxBadFrac)
  #Work out the final filter
  srat@meta.data$PASS = with(srat@meta.data,PASS_MT & PASS_nGenes & PASS_nCounts & PASS_doublet & PASS_cluster)
  #Make a reason for fail variable
  nBasicFail = with(srat@meta.data,4 - (PASS_MT + PASS_nGenes + PASS_nCounts + PASS_doublet))
  tmp = rep('Pass',length(nBasicFail))
  tmp[!srat@meta.data$PASS_MT] = 'highMT'
  tmp[!srat@meta.data$PASS_nGenes] = '#genesLow'
  tmp[!srat@meta.data$PASS_nCounts] = '#countsLow'
  tmp[!srat@meta.data$PASS_doublet] = 'doublet'
  tmp[nBasicFail>1] = 'multiple'
  #Don't want "multiple" to encompass cluster otherwise there'll be heaps where cluster + bad obscures the true reason
  tmp[tmp=='Pass' & !srat@meta.data$PASS_cluster] = 'badCluster'
  srat@meta.data$reasonForFail = tmp
  #Make some QC plots before dropping things
  if(doPlot){
    if(verbose)
      message("Making diagnostic plots.")
    #Plot distributions of basic things
    df = srat@meta.data
    par(mfrow=c(2,2))
    plot(density(df$mtGenes),main='MT',xlab='MT Frac')
    abline(v=maxMT,col='red')
    plot(density(log10(df$nFeature_RNA)),main='#Genes',xlab='log10(No Genes)')
    abline(v=log10(minGenes),col='red')
    plot(density(log10(df$nCount_RNA)),main='#UMIs',xlab='log10(No UMIs)')
    abline(v=log10(minUMIs),col='red')
    plot(density(df$scrubScore),main='Scrublet Score',xlab='Scrub Score')
    #Standard QC things
    plot(DimPlot(srat,group.by='seurat_clusters',label=TRUE,repel=TRUE)+guides(colour=FALSE))
    #A hack for another of Seurat's stupid decisions
    if('labels' %in% names(as.list(args(DimPlot)))){
      plot(DimPlot(srat,group.by=c('Phase','orig.ident'),labels=c('Phase','Source')))
    }else{
      gg = DimPlot(srat,group.by=c('Phase','orig.ident'),combine=FALSE)
      labs = c('Phase','Source')
      for(i in seq_along(labs)){
        gg[[i]] = gg[[i]]  + ggtitle(labs[i])
      }
      plot(patchwork::wrap_plots(gg))
    }
    plot(FeaturePlot(srat,c('mtGenes','nFeature_RNA','nCount_RNA','fracBadClust')))
    #Which cells pass based on what
    if('labels' %in% names(as.list(args(DimPlot)))){
      plot(DimPlot(srat,group.by=c('PASS_MT','PASS_nGenes','PASS_nCounts','PASS_doublet','PASS_cluster','PASS'),labels=c('Pass MT Frac?','Pass #Genes?','Pass #Counts?','Pass Doublet?','Pass cluster','PASS'),legend='none'))
    }else{
      gg = DimPlot(srat,group.by=c('PASS_MT','PASS_nGenes','PASS_nCounts','PASS_doublet','PASS_cluster','PASS'),combine=FALSE)
      labs =c('Pass MT Frac?','Pass #Genes?','Pass #Counts?','Pass Doublet?','Pass cluster','PASS')
      for(i in seq_along(labs)){
        gg[[i]] = gg[[i]] +guides(colour=FALSE) + ggtitle(labs[i])
      }
      plot(patchwork::wrap_plots(gg))
    }
    plot(DimPlot(srat,group.by=c('reasonForFail')))
    #srat@meta.data$isClustBad = ifelse(srat@meta.data$fracBadClust >maxBadFrac | !srat@meta.data$PASS,'Drop','Keep')
    #plot(DimPlot(srat,group.by='isClustBad',label=FALSE))
    #srat@meta.data$isClustBad=NULL
    #srat@meta.data$fracBadClust=NULL
  }
  #Filter
  if(verbose)
    message("Filtering and finalising.")
  w = which(srat@meta.data$PASS)
  #w = which(!(as.character(srat@meta.data$seurat_clusters) %in% as.character(tt[tt[,2]>maxBadFrac,1]) | !srat@meta.data$PASS))
  #srat@meta.data$clusterPASS = TRUE
  #srat@meta.data$clusterPASS[w] = FALSE
  #Make the new version, store the old one in it
  sratOld = srat
  #Restart without them
  mDat = srat@meta.data[w,]
  #Drop old clustering
  mDat = mDat[,!colnames(mDat) %in% c('seurat_clusters',grep('^RNA_snn_res',colnames(mDat),value=TRUE))]
  #And drop genes that we don't care about
  genesToKeep = rownames(srat@assays$RNA@counts)
  genesToKeep = genesToKeep[!genesToKeep %in% excludeGenes]
  srat = CreateSeuratObject(srat@assays$RNA@counts[genesToKeep,w])
  #Merge in metadata
  srat@meta.data = cbind(srat@meta.data,mDat[,!(colnames(mDat) %in% colnames(srat@meta.data)),drop=FALSE])
  srat = NormalizeData(srat,verbose=FALSE)
  srat = FindVariableFeatures(srat,verbose=FALSE)
  srat@misc$preQC = sratOld
  srat@misc$geneMap = sratOld@misc$geneMap
  srat@misc$geneMapCommon = sratOld@misc$geneMapCommon
  return(srat)
}


#' Run scrublet
#' 
#' Uses a crude call to python to run scrublet and pull in the results.
#'
#' @param dat10X Directory containing 10X matrix and genes files.
#' @param nPCs Number of PCs to use.
#' @return A data.frame with the scrublet results
runScrublet = function(dat10X,nPCs){
  fNom = tempfile()
  cmd = sprintf(
  "import scrublet as scr
import scipy.io
import numpy as np
import os
import gzip
os.chdir('%s')
input_dir = os.path.expanduser('%s')
output_file = '%s'
try:
  counts_matrix = scipy.io.mmread(input_dir + '/matrix.mtx').T.tocsc()
  genes = np.array(scr.load_genes(input_dir + '/genes.tsv', delimiter='\\t', column=1))
except FileNotFoundError:
  counts_matrix = scipy.io.mmread(input_dir + '/matrix.mtx.gz').T.tocsc()
  #Copy relevant part of scr.load_genes
  gene_list = []
  gene_dict = {}
  with gzip.open(input_dir + '/features.tsv.gz','rt') as f:
    for l in f:
      gene = l.strip('\\n').split('\\t')[1]
      if gene in gene_dict:
        gene_dict[gene] += 1
        gene_list.append(gene + '__' + str(gene_dict[gene]))
        if gene_dict[gene] == 2:
          i = gene_list.index(gene)
          gene_list[i] = gene + '__1'
      else: 
        gene_dict[gene] = 1
        gene_list.append(gene)
  genes = np.array(gene_list)
#Convert to integers.  Will do nothing most of the time.
counts_matrix.data = np.round(counts_matrix.data).astype('int32')
scrub = scr.Scrublet(counts_matrix)
doublet_scores, predicted_doublets = scrub.scrub_doublets(min_counts=2, 
                                                        min_cells=3, 
                                                        min_gene_variability_pctl=85, 
                                                        n_prin_comps=%d)
with open(output_file,'w') as f:
  #Header row
  f.write('score\\tcall\\tsimScore1\\tsimScore2\\n')
  #As nSim = 2*nObs, we can split into two columns 
  offset = len(doublet_scores)
  f.write('\\n'.join([str(x)+'\\t'+str(predicted_doublets[i]).upper()+'\\t'+str(scrub.doublet_scores_sim_[i])+'\\t'+str(scrub.doublet_scores_sim_[offset+i]) for i,x in enumerate(doublet_scores)]))
  f.close()",getwd(),dat10X,fNom,nPCs)
  system(paste0('python -c "',cmd,'"'))
  out = read.table(fNom,header=TRUE,sep='\t')
  if(file.exists(file.path(dat10X,'barcodes.tsv'))){
    tmp = read.table(file.path(dat10X,'barcodes.tsv'),header=FALSE)
  }else{
    tmp = read.table(file.path(dat10X,'barcodes.tsv.gz'),header=FALSE)
  }
  out$barcode = tmp[,1]
  return(out)
}
```
